# Identification of 5-HT_2A_ Receptor Signaling Pathways Responsible for Psychedelic Potential

**DOI:** 10.1101/2023.07.29.551106

**Authors:** Jason Wallach, Andrew B. Cao, Maggie M. Calkins, Andrew J. Heim, Janelle K. Lanham, Emma M. Bonniwell, Joseph J. Hennessey, Hailey A. Bock, Emilie I. Anderson, Alexander M. Sherwood, Hamilton Morris, Robbin de Klein, Adam K. Klein, Bruna Cuccurazzu, James Gamrat, Tilka Fannana, Randy Zauhar, Adam L. Halberstadt, John D. McCorvy

## Abstract

Serotonergic psychedelics possess considerable therapeutic potential. Although 5-HT_2A_ receptor activation mediates psychedelic effects, prototypical psychedelics activate both 5-HT_2A_-Gq/11 and β-arrestin2 signaling, making their respective roles unclear. To elucidate this, we developed a series of 5-HT_2A_-selective ligands with varying Gq efficacies, including β-arrestin-biased ligands. We show that 5-HT_2A_-Gq but not 5-HT_2A_-β-arrestin2 efficacy predicts psychedelic potential, assessed using head-twitch response (HTR) magnitude in male mice. We further show that disrupting Gq-PLC signaling attenuates the HTR and a threshold level of Gq activation is required to induce psychedelic-like effects, consistent with the fact that certain 5-HT_2A_ partial agonists (e.g., lisuride) are non-psychedelic. Understanding the role of 5-HT_2A_-Gq efficacy in psychedelic-like psychopharmacology permits rational development of non-psychedelic 5-HT_2A_ agonists. We also demonstrate that β-arrestin-biased 5-HT_2A_ receptor agonists induce receptor downregulation and tachyphylaxis, and have an anti-psychotic-like behavioral profile. Overall, 5-HT_2A_ receptor signaling can be fine-tuned to generate ligands with properties distinct from classical psychedelics.

## Introduction

Classical (serotonergic) psychedelics have undergone a resurgence of interest due to their potential to produce rapid and sustained therapeutic effects^1, 2^. The therapeutic properties of psychedelics are being explored both preclinically and clinically, but studies have focused on a limited number of compounds^3, 4^. The therapeutic use of psychedelics may be limited by their hallucinogenic effects, which can cause confusion and anxiety in some patients, necessitating close clinical supervision. Recent preclinical work, however, suggests it may be possible to disentangle the psychedelic and therapeutic properties^5, 6^. Nevertheless, fundamental questions exist regarding which receptors and signaling pathways mediate the effects of psychedelics, limiting the rational design of new molecules.

Serotonergic psychedelics are derived from multiple chemical scaffolds (e.g., tryptamines, phenethylamines, and lysergamides), all which activate the 5-HT_2A_ receptor (5-HT_2A_R), a G protein-coupled receptor (GPCR). 5-HT_2A_Rs appear to primarily mediate psychedelic experiences as evident with studies with the 5-HT_2A_R antagonist ketanserin, which attenuates subjective effects of psilocybin and LSD in humans^7, 8^. Responses to psychedelics has also been studied in preclinical behavioral models, including the head-twitch response (HTR), which is a 5-HT_2A_R-mediated involuntary head movement in mice that predicts human psychedelic activity^9^. Multiple receptors, however, appear to contribute to the behavioral effects of existing psychedelics, including psilocybin^10^ and LSD^11^, which adds to their complicated psychopharmacology.

Ligands targeting GPCRs can stabilize certain receptor conformations that energetically favor coupling to subsets of transducer proteins^12, 13^. Ligand-dependent bias has important implications for drug development and clinical pharmacology^14, 15^. For example, the G protein-biased μ-opioid receptor (MOR) agonist oliceridine reportedly has a correspondingly improved tolerability profile^16^, although alternative explanations exist^17^. As existing psychedelics activate both Gq and β-arrestin2 via 5-HT_2A_R^18, 19^, the role these pathways play in the effects of psychedelics is unclear. Although putative non-psychedelic 5-HT_2A_R agonists exist^5, 6^, an adequate explanation for their lack of psychedelic action does not exist. A clear and defined pharmacological signaling mechanism explaining why certain 5-HT_2A_R agonists lack psychedelic effects is thus needed.

Herein, we utilized structure-inspired design to develop 5-HT_2A_R-selective ligands acting as biased agonists and leveraged these compounds to develop a mechanistic and molecular explanation of biased 5-HT_2A_R signaling. Furthermore, these compounds were used to probe the relationship between 5-HT_2A_R-Gq versus 5-HT_2A_R-β-arrestin activity and psychedelic potential *in vivo*. Our goal was to identify the 5-HT_2A_R-coupled signaling pathways mediating psychedelic activity for the design of next-generation 5-HT_2A_R agonists.

## Results

### Psychedelics exhibit similar Gq and β-arrestin2 activity at 5-HT_2A_R

To investigate the 5-HT_2A_R signaling profiles of classical psychedelics, we used bioluminescence resonance energy transfer approaches (BRET, Fig. 1A,B, Supplementary Table 1), which provide a proximity measure of intracellular transducer engagement. BRET is used extensively to quantify GPCR biased agonism and has the advantage of not being susceptible to second messenger amplification or receptor reserve issues obfuscating GPCR signaling preference determinations^20^. BRET has been used to measure 5-HT_2A_R-β-arrestin2 and 5-HT_2A_R-Gq activity directly^11^, and to confirm 5-HT_2A_R G protein coupling preferences^18^. In our assay platform, 5-HT_2A_R strongly couples to Gq/11 and β-arrestin2 over all other G protein subtypes and β-arrestin1 (Fig. 1C).

**Figure 1.**
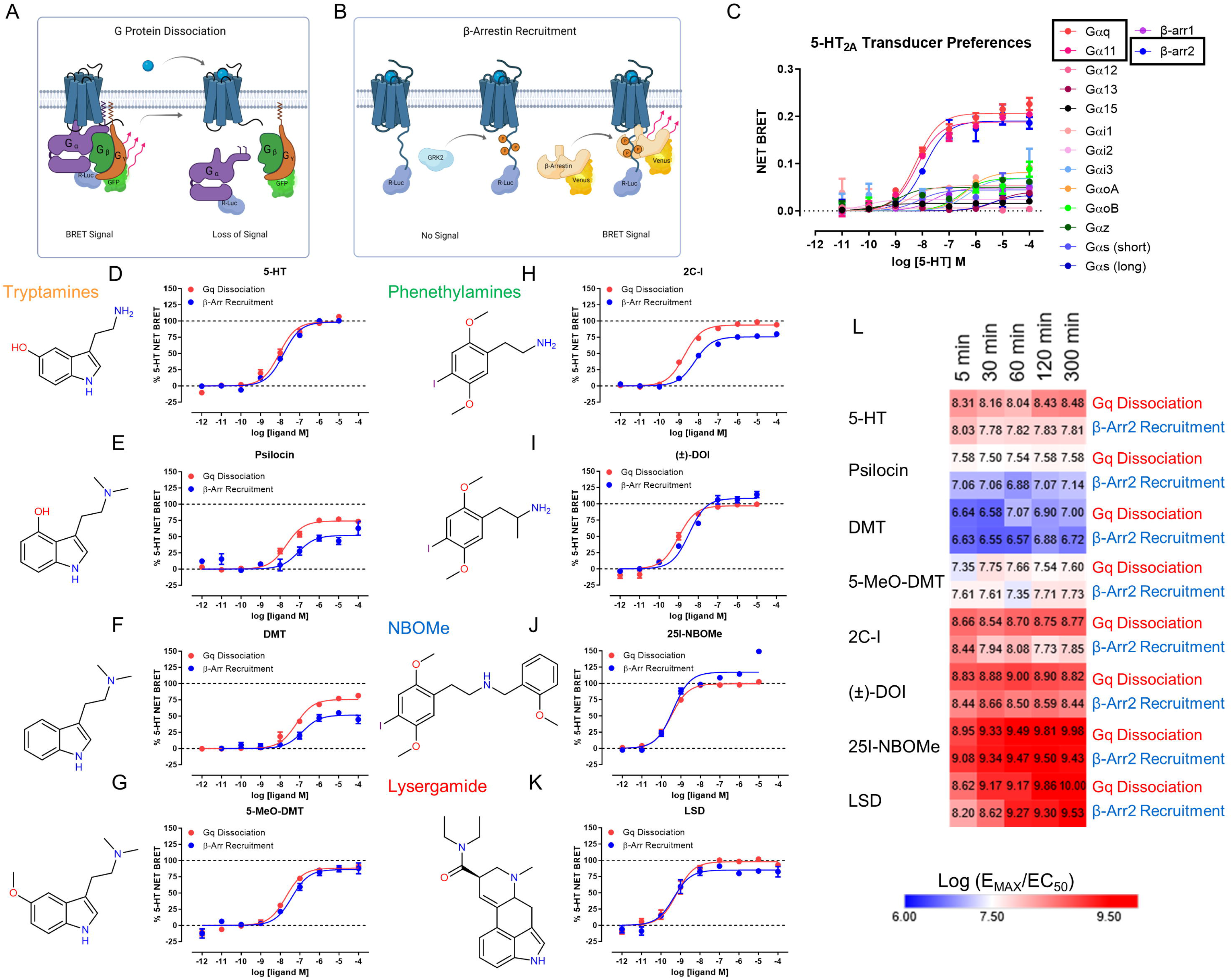
Psychedelics exhibit similar Gq and β-arrestin2 activity at 5-HT_2A_R. Schematic of 5-HT_2A_ receptor G protein dissociation **(A)** and β-arrestin recruitment **(B)** to determine signaling preferences. **(C)** Determination of 5-HT_2A_ receptor G protein-wide and β-arrestin1/2 transducer preferences as estimated by magnitude net BRET for each transducer as stimulated by 5-HT. Data represent the mean and SEM from three independent experiments, which were performed at 37°C with 60 minute compound incubations. **(D-K)** Comparison of 5-HT_2A_ receptor Gq dissociation (red) and β-arrestin2 recruitment (blue) for 5-HT **(D)** and several classes of psychedelics: **(E)** Psilocin, **(F)** DMT, **(G)** 5-MeO-DMT, **(H)** 2C-I, **(I)** DOI, **(J)** 25I-NBOMe, **(K)** LSD. Data represent the mean and SEM from three independent experiments, which were performed at 37°C with 60 minute compound incubations. **(L)** Heat map of Gq dissociation and β-arrestin2 recruitment kinetics displayed as log (E_MAX_/EC_50_) for the psychedelics tested. Data represent the mean and SEM from three independent experiments, which were performed at 37°C at the indicated compound incubation time points.

Next, we tested classical psychedelics from multiple chemical classes and examined their effects on Gq and β-arrestin2 (Fig. 1D-K). Kinetic issues can confound studies of ligand-directed bias^21^, and slow ligand kinetics can delay full receptor occupancy, as in the case for LSD at 5-HT_2A_R^19^. Therefore, we thoroughly assessed Gq and β-arrestin2 activities at various time points at 37°C to ensure full receptor occupancy and confirm that transducer signaling preferences do not substantially change when compared at the same time point. Our results show that psychedelics exhibit dynamic, time-dependent profiles of Gq and β-arrestin activity that in some cases exceeds the activity of 5-HT (i.e., superagonism) at longer time points (e.g., 300 minutes, Supplementary Fig. 1A-H), which is not surprising given the dynamic temporal nature of GPCR signaling^22^. For all tested psychedelics, however, effects on Gq and β-arrestin were strikingly similar at equivalent time points and closely mirrored the pathway-balanced endogenous ligand 5-HT (Heat map, Fig. 1L), indicating no strong preferences for one of the transducers. Psilocin, DMT and 2C-I showed slightly less β-arrestin2 efficacy compared to Gq activity, but the difference was not substantial when both transducers were compared at each respective time point (Supplementary Fig. 1B,C,E). Importantly, all tested psychedelics lacked a clear preference for Gq or β-arrestin2, demonstrating these compounds are not substantially biased for either transducer.

### Rational design of a 5-HT_2A_-selective agonist template

To engineer a series of biased agonists for interrogating the 5-HT_2A_R-coupled signaling pathways responsible for psychedelic potential, a scaffold exhibiting some degree of selectivity for 5-HT_2A_R over other 5-HT receptors is required. Most psychedelics are not selective for 5-HT_2A_R, exhibiting complex polypharmacology^11^. 5-HT_2A_R shares considerable homology with 5-HT_2B_R and 5-HT_2C_R, making it challenging to develop selective 5-HT_2A_R agonists. To date, few 5-HT_2A_R-selective agonists have been discovered, but the *N*-benzyl-phenethylamines 25CN-NBOH^23^ and DMBMPP^24^ are purported examples. Importantly, recently discovered “non-psychedelic” 5-HT_2A_R agonists show little selectivity for 5-HT_2A_R^5, 6^, complicating attempts to interpret their psychopharmacology.

To develop selective 5-HT_2A_R biased ligands, we focused on the phenethylamine scaffold, which tends to have high selectivity for 5-HT_2_ subtypes. 25N or 2C-N (1) was selected as the core phenethylamine based on previous reports of potential 5-HT_2A_R biased agonism^25, 26^. We confirmed 25N (1) is a high-affinity, potent 5-HT_2_ agonist with weak selectivity for 5-HT_2A_R and 5-HT_2C_R over 5-HT_2B_R (Fig. 2A, Supplementary Table 8). Because *N*-benzylation can increase affinity and potency of phenethylamines at 5-HT_2A_R ^27^, we converted 25N (1) to 25N-NB (2) (see Supplement for all synthetic schemes), which reduced 5-HT_2B_R efficacy substantially (E_MAX_ = 32% of 5-HT, Fig. 2A). Unfortunately, 25N-NB (2) retained potent 5-HT_2C_R activity (EC_50_ = 8.3 nM), resulting in a weakly (4-fold) 5-HT_2A_R-selective ligand.

**Figure 2.**
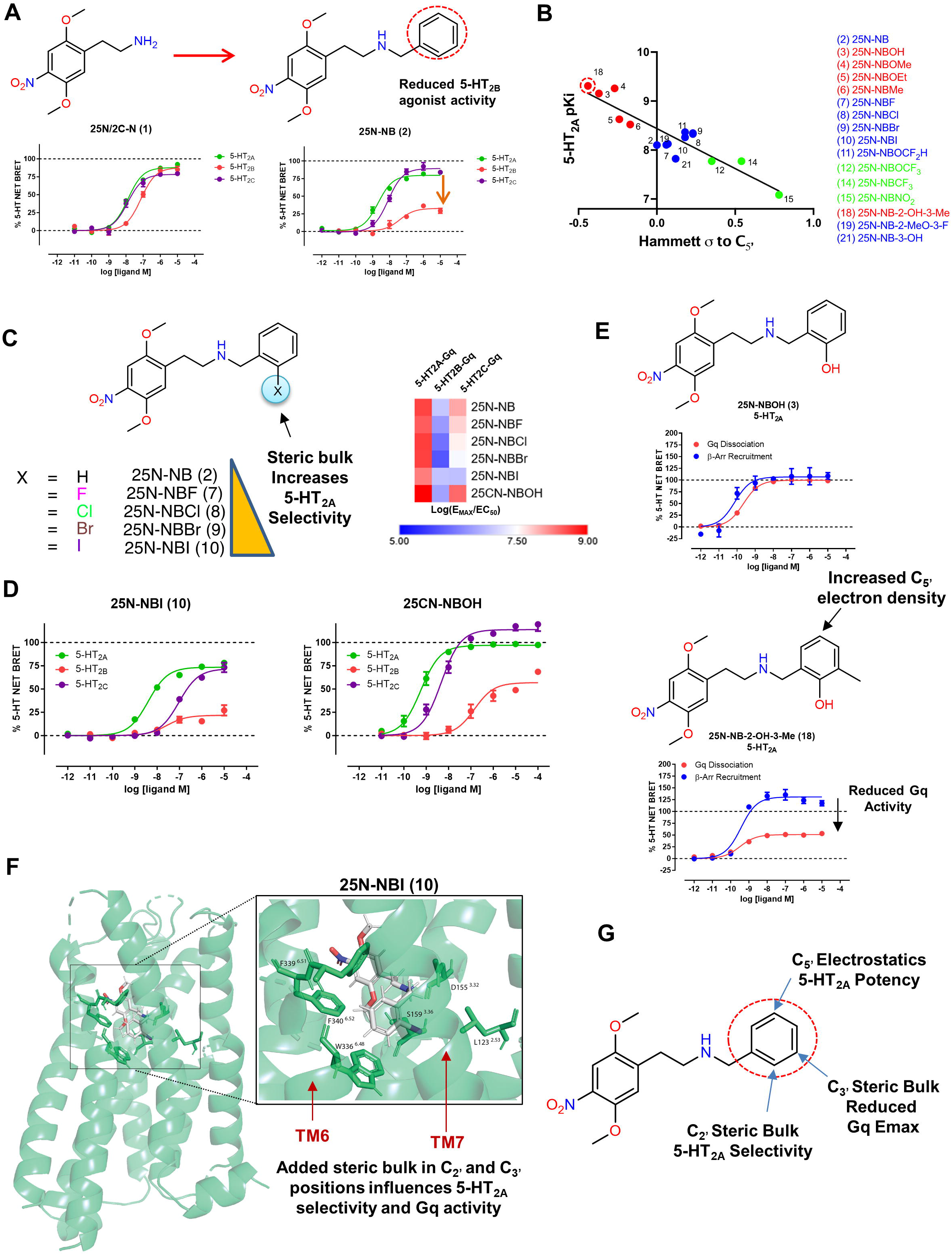
Rational design of a 5-HT_2A_-selective agonist template. **(A)** *N*-Benzylation of 25N (1) to 25N-NB (2) leads to reduced 5-HT_2B_ receptor efficacy, as measured by Gq dissociation by BRET. Data represent the mean and SEM from three independent experiments performed at 37°C with 60 minute compound incubation. **(B)** Role of *N*-benzyl ring electrostatics in 5-HT_2A_ receptor potency leading to development of 25N-NB-2-OH-3-Me (18) using QSAR. Correlation between 5-HT_2A_ receptor p*K*_i_ and Hammett constant values (Pearson’s *R* = -0.9078, *R*^2^ = 0.8241, *p*<0.0001, *N* = 16). **(C)** The relationship between steric bulk and 5-HT_2A/2C_ receptor selectivity is shown for the halogen series, leading to the identification of the 5-HT_2A_ receptor-selective agonist 25N-NBI (10) (*left*). Also shown is a 5-HT_2A/2C_ receptor selectivity heatmap comparing the 25N halogen series to 25CN-NBOH (*right*). **(D)** Comparison of 5-HT_2A_ receptor (green) 5-HT_2B_ receptor (red) and 5-HT_2C_ receptor (purple) Gq dissociation activities for 25N-NBI (10) (*left*) and 25CN-NBOH (*right*). Data represent mean and SEM from three independent experiments, which were performed at 37°C with 60 minute compound incubation **(E)** 5-HT_2A_ receptor Gq dissociation and β-arrestin2 BRET concentration response curves for 25N-NBOH (3, *top*) and 25N-NB-2-OH-3-Me (18, *bottom*) showing addition of a 3-methyl group leads to reduced Gq efficacy. **(F)** 25N-NBI (10) induced fit docking (IFD) with orthosteric site residue side chains displayed. The window shows a zoom-in view illustrating key ligand-residue interactions within the orthosteric site and illustrating the close proximity of the 2’- and 3’-positions to TM6 and TM7, which are known to influence ligand bias. **(G)** Summary of structure-activity relationships (SAR) for the 25N series encompassing key effects on electrostatics, 5-HT_2A_ receptor selectivity, and reduced Gq E_MAX_.

To optimize 5-HT_2A_R affinity and selectivity, we synthesized a series of 25N-NB analogs designed to modify the electrostatic properties of the *N*-benzyl ring system (Supplementary Fig. 2; Supplementary Table 2). The increased 5-HT_2A_R affinity of *N*-benzyl-phenethylamines is thought to result in part from hydrogen bonding between the *N*-benzyl 2-position and residues in 5-HT_2A_R^28^, but we developed an alternate hypothesis that ring-electrostatics (i.e. the effect of increasing π-electron density in portions of the *N*-benzyl-ring, quantified as ring-substituents with negative Hammett constants relative to C_5’_) drives 5-HT_2A_R affinity (Fig. 2B, Supplementary Fig. 3; Supplementary Table 3). Using the 25N series, we found that estimates of increasing electron density around the *N*-benzyl C_5’_ position (para to the 2’-position) increase 5-HT_2A_R binding affinity and agonist potency (Supplementary Fig. 3; Supplementary Tables 3-8). To confirm the importance of this ring-region, we tested the effect of adding a methoxy group to the C_5’_ position of 25N-NBOMe (4), which this analog 25N-NB-2,5-DiMeO (20) reduced affinity 400-fold, suggesting steric clash and/or altered electronics disrupted optimal electrostatic interaction (Supplementary Fig. 3J). The electrostatic-relationship was supported by correlations with additional potency estimates, experimental NMR chemical shifts, and *in silico* Hirshfeld surface analyses (Supplementary Fig. 3A-I, K-P, Supplementary Table 4-6). This resulted in the discovery of several high-affinity 5-HT_2A_R agonists within the *N*-benzyl-phenethylamine class of psychedelics (Supplementary Table 8).

We then examined the effect of *N*-benzyl 2-postion substitutions on 5-HT_2A_R selectivity for the 25N series. In the 2-halogen series, a larger/bulkier 2-bromo or 2-iodo atom on the *N*-benzyl ring reduced 5-HT_2C_R and 5-HT_2B_R activity substantially but retained potent 5-HT_2A_R activity, resulting in increased 5-HT_2A_R selectivity (Fig. 2C, Supplementary Fig. 4A). We compared 25N-NBI (10), the most selective 5-HT_2A_R agonist from this series, to the purported 5-HT_2A_R-selective agonist 25CN-NBOH^29^ and determined 25N-NBI (10) showed 23-fold selectivity for 5-HT_2A_R over 5-HT_2C_R, whereas 25CN-NBOH shows only 7-fold selectivity (Fig. 2D). We confirmed the superior 5-HT_2A_R selectivity of 25N-NBI (10) using an orthogonal assay measuring Gq-mediated calcium flux (Supplementary Fig. 4B; Supplementary Table 7), confirming in two functional assays that 25N-NBI (10) has greater 5-HT_2A_R selectivity than 25CN-NBOH. 25N-NBI (10) also showed selectivity for 5-HT_2A_R over 5-HT_2B_R and 5-HT_2C_R in competitive binding studies (Supplementary Table 8). Furthermore, when tested at >40 other 5-HT receptors and off-targets, 25N-NBI (10) exhibited weak micromolar affinity for all targets screened (Supplementary Table 9). Competitive radioligand binding studies showed high selectivity for 5-HT_2_ subtypes across the series (Supplementary Table 8-10). Finally, 25N-NBI (10) induced the HTR in mice (ED_50_ = 10.9 μmol/kg), confirming selective 5-HT_2A_R agonists possess psychedelic potential (Supplementary Fig. 4C).

Although a 2-iodo *N*-benzyl substitution yielded a 5-HT_2A_R-selective agonist with psychedelic potential, 25N-NBI (10) did not show a preference for either 5-HT_2A_R-Gq or 5-HT_2A_R-β-arrestin2 activity (Supplementary Fig. 4D). Therefore, we focused on another 25N analog, 25N-NB-2-OH-3-Me (18; Fig. S2A), which was rationally designed to maximize C_5’_ electron density (Supplementary Fig. 3A, Supplementary Tables 3,4) and confirmed to have the highest 5-HT_2A_R affinity in the 25N series (Fig. 2B,E). 25N-NB-2-OH-3-Me (18) showed a selective reduction in 5-HT_2A_R-Gq efficacy but not 5-HT_2A_R-β-arrestin2 efficacy compared to 25N-NBOH (3) (Fig. 2E), which lacks 3-methyl substitution, providing a critical clue that steric effects in the 3-position influence 5-HT_2A_R biased-agonist activity.

To gain additional insights into the binding mode of the 25N series, we docked 25N-NBI (10), 25N-NB-2-OH-3-Me (18), and other analogs into the Gq-bound 5-HT_2A_R cryo-EM structure (6WHA)^18^ using induced fit docking (IFD) (Fig. 2F; Supplementary Fig. 5). Consistent with other studies^18, 27^, the *N*-benzyl groups formed an edge-to-face π-π stacking interaction with F339^6.51^ and F340^6.52^, known drivers of high 5-HT_2A_R affinity^27, 30^ (Fig. 2D and Supplementary Table 20). We validated this docking pose and the 5-HT_2A_R model using a series of binding pocket mutants. Examination of mutants in transmembrane 6 (TM6), extracellular loop 2 (EL2), TM3, and TM7 indicate the 25N analog binding poses are consistent with the 25CN-NBOH-bound cryo-EM structure. Specifically, mutants of key aromatic anchor residues, F339^6.51^L and F340^6.52^L, showed complete loss of 25N-NBI (10) Gq activity (Supplementary Fig. 4E), suggesting the *N*-benzyl moiety is positioned near TM6. In summary, we found that 5-HT_2A_R affinity, selectivity, and potentially biased agonism can be modulated in the 25N series, and that additional bulk in the 2- and 3-positions of the *N*-benzyl ring may drive 5-HT_2A_R selectivity and ligand bias, respectively (Fig. 2E-G).

### Structure-based design of β-arrestin-biased 5-HT_2A_ agonists

Based on the promising results obtained with the initial 25N series that led to increased 5-HT_2A_R selectivity, we explored whether it is possible to reduce Gq signaling further by replacing the *N*-benzyl ring with bulkier bi-aryl ring systems such as *N*-naphthyl (25N-N1-Nap (16)) and *N*-biphenyl (25N-NBPh (17)). Steric interactions within GPCR binding pockets are an important driver of agonist, antagonist, and biased-agonist conformational states, especially when they involve residues in the extended binding pocket encompassing TM6 and TM7^12, 31^. Strikingly, 25N-N1-N (16) and 25N-NBPh (17) showed a substantial reduction in 5-HT_2A_R-Gq efficacy yet preserved 5-HT_2A_R-β-arrestin efficacy, thus exhibiting β-arrestin-biased agonism at 5-HT_2A_R compared to the balanced agonist 25N-NBOMe (4) (Fig. 3A; Supplementary Table 13). We further explored the SAR by synthesizing *N*-benzyl analogs containing oth and 3-substituents and this also results in β-arrestin bias (Supplementary Fig. 6A). Similar to our findings with 25N NBI (10), the bulky analogs 25N-N1-Nap (16) and 25N-NBPh (17) showed weaker Gq activity at 5-HT_2B_R and 5-HT_2C_R, retaining degrees of 5-HT_2A_R selectivity (Supplementary Fig. 6B; Supplementary Tables 13 and 14). Bindin affinities confirmed these compounds exhibit weak affinities for other 5-HT receptors and off-targets (Supplementa Table 8-10). To test whether the β-arrestin bias was specific for 5-HT_2A_R, we measured β-arrestin2 activity at 5-HT and 5-HT_2C_R and found substantially weaker β-arrestin2 recruitment at these subtypes (Supplementary Fig. 6C), confirming a lack of biased agonism at these receptors. Furthermore, weak activity was confirmed in 5-HT_2A_-G11 dissociation, 5-HT_2A/2B/2C_-Gq/11-mediated calcium flux activity, and at all human 5-HT GPCRs measuring G protein agonism by BRET confirmed with 25N-N1-Nap (16) and 25N-NBPh (17) (Supplementary Fig. 6D,E, F). Interesting the β-arrestin-biased agonists showed partial antagonism of 5-HT_2A_-Gq activity, similarly found with the non-psychedelic 2-Br-LSD^11^, and also showed baseline inhibition in 5-HT_2A_R Gq/11-mediated calcium flux traces at long time points, suggestive of potential β-arrestin2-mediated desensitization in this cell system (Supplementary Fig. 6 G). We also confirmed and validated β-arrestin2 bias using our previously established kinetic BRET assay, which consistently showed robust β-arrestin2 and weak Gq activities for 25N-N1-Nap (16) and 25N-NBPh (17) at all time points tested (Supplementary Fig. 6H).

**Figure 3.**
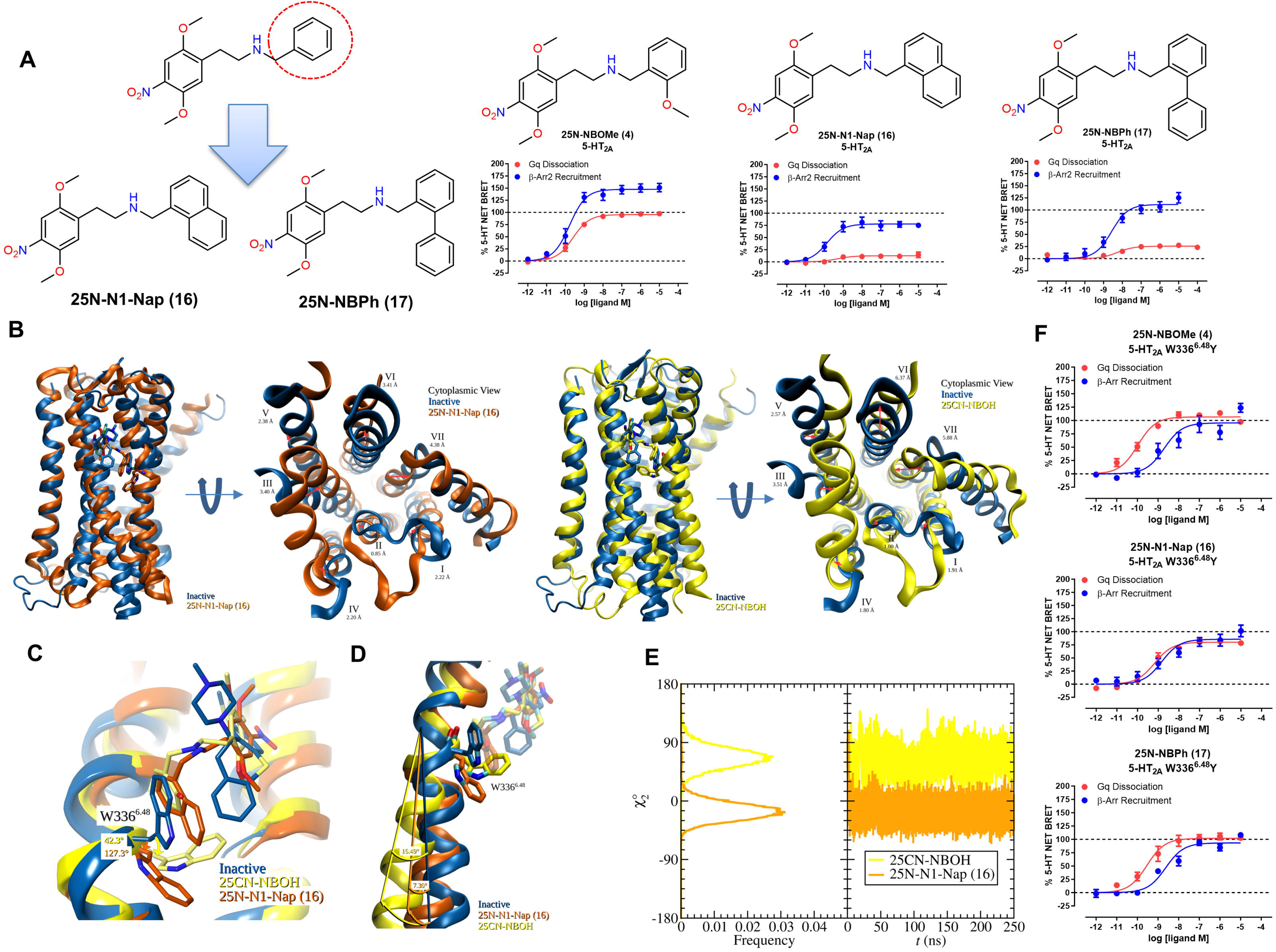
Structure-based design of β-arrestin-biased 5-HT_2A_ agonists. **(A)** Effect of the larger *N*-substituted 25N analogs 25N-N1-Nap (16) and 25N-NBPh (17) on 5-HT_2A_ receptor Gq dissociation (red) and β-arrestin2 (blue) recruitment. Data represent the mean and SEM from three independent experiments performed at 37°C with 60 minute compound incubation. **(B)** Outward pivot of TM6 for 25CN-NBOH and 25N-N1-Nap (16) MD simulations relative to the inactive 5-HT_2A_ receptor structure with final frame shown as representative. Note the intermediate TM6 tilt angle with 25N-N1-Nap (16) relative to that of 25CN-NBOH. **(C)** Change in W366^6.48^ toggle switch χ^2^ angle. χ^2^ angle given is the absolute difference in peak angle relative to the inactive state. Final frame shown as representative. **(E)** Distribution and time dependence W366^6.48^ χ^2^ angle of 25CN-NBOH (yellow) and 25N-N1-Nap (16) (orange) simulations. **(E)** Effect of 5-HT_2A_ receptor W336^6.48^Y mutation on 25N-NBOMe (4), 25N-NBPh (17), and 25N-N1-Nap (16) Gq dissociation (red) versus β-arrestin2 (blue) recruitment activities. Data represent the mean and SEM from three independent experiments performed at 37°C with 60 minute compound incubation.

Encouraged by the progress made identifying β-arrestin-biased 5-HT_2A_R agonists, we examined the binding mode of 25N-N1-Nap (16) at the Gq-bound cryo-EM 5-HT_2A_R structure (PDB: 6WHA) using IFD (Supplementary Fig. 5). Binding poses using RMSD as a metric were clustered to reveal key interactions with the conserved salt bridge to Asp155^3.32^ placed 25N-N1-Nap (16) in a similar pose as the 25CN-NBOH structure (Supplementary Table 20; Supplementary Fig. 5). Importantly, in the flexible docking, the *N*-naphthyl ring of 25N-N1-Nap (16) is wedged near W336^6.48^, a highly conserved residue in class A GPCRs (Supplementary Fig. 5). W336^6.48^, sometimes called the “toggle switch”, is implicated in GPCR activation and signaling bias, with distinct W6.48 rotamer conformations occurring in activated and non-activated GPCR structures^32–35^, including the 25CN-NBOH cryo-EM structure^18^. Given the close proximity and added steric bulk observed in this region with biased agonists like 25N-N1-Nap (16), and based on the electrostatic SAR, we hypothesized the weak Gq activity and β-arrestin bias are dependent, in part, on interactions with W336^6.48^.

GPCR crystal and cryo-EM structures only show a “snapshot” of activation and are highly dependent on their ternary complex composition and intracellular binding partners. Therefore, we performed molecular dynamics (MD) to investigate interactions between 25N-N1-Nap (16) and 5-HT_2A_R, focusing especially on residue W336^6.48^, and compared the results to simulations performed with 25CN-NBOH in the cryo-EM structure. Across 250 ns simulations, time-dependent intermolecular interaction energies were maintained between each ligand and the anchoring orthosteric residues D155^3.32^, F339^6.51^ and F340^6.52^ (Supplementary Fig. 7, 8). Comparison of the trajectories reveals that the 25N-N1-Nap simulation has features intermediate between the inactive (6WH4) and active (25CN-NBOH) conformations (Fig. 3B,C). For example, in the final frame of the simulation, the outward swing of TM6, with 25N-N1-Nap (16), relative to the inactive state (6WH4) is lesser than that of 25CN-NBOH (7.30° vs. 15.49°) (Fig. 3D). A clear difference in the W336^6.48^ χ_2_ angle frequency was observed between the 25CN-NBOH and 25N-N1-Nap simulations, where 25N-N1-Nap (16) showed a preference for W336^6.48^ χ_2_ angle down (centered at -15.6°), and 25CN-NBOH showed a preference for W336^6.48^ χ_2_ angle up (66.2°) in the active rotamer conformation (Fig. 3E), similar to the active state cryo-EM structure^18^. Interestingly, in the 25N-N1-Nap simulation, W336^6.48^ initially toggles further downward into the pocket and mostly remains in this orientation throughout the simulation, likely to accommodate the larger *N*-naphthyl ring in 25N-N1-Nap (16), and is distinct from the W336^6.48^ χ_2_ angles in active- (6WHA) and inactive-state 5-HT_2A_R structures (Fig. 3B,E; e.g., PDB structures 6WGT, 6A94, 6A93, 6WH4; compare Supplementary Tables 21 and 22).

To verify the role of W336^6.48^ in 5-HT_2A_R biased signaling, we constructed conservative mutations, W336^6.48^Y and W336^6.48^L, designed to increase space and accommodate the larger substituents in 25N β-arrestin-biased ligands. Although the W336^6.48^L mutant exhibited reduced Gq dissociation and impaired β-arrestin recruitment (Fig. S3F), preventing further experimental use, the W336^6.48^Y mutant showed robust Gq and β-arrestin recruitment and was used in subsequent experiments (Supplementary Fig. 6I). When 25N-N1-Nap (16) and 25N-NBPh (17) were tested at the W336^6.48^Y mutant, the Gq efficacy recovered substantially, resulting in balanced agonist activity similar to 25N-NBOMe (4) (Fig. 3F). The recovery of Gq activity likely results from the increased space created by a smaller size Tyr residue at position 6.48, allowing the “toggle switch” to move dynamically to initiate Gq-bound activation states. The mutagenesis data support the hypothesis that the W366^6.48^ toggle switch plays an important role in the signaling bias of 25N-N1-Nap (16) and 25N-NBPh (17), with steric bulk on the *N*-benzyl ring forcing W336^6.48^ into a unique rotamer conformation, making it less likely that Gq-bound conformations will arise while preserving β-arrestin-preferring conformational states.

### β-Arrestin-biased 5-HT_2A_ agonists lack psychedelic potential

The HTR is commonly used as a behavioral proxy for psychedelic effects because non-psychedelic 5-HT_2A_R agonists do not induce head twitches^36^, and because there is a robust correlation between HTR activity in mice and potency to induce psychedelic effects in humans and discriminative stimulus effects in rats^9^. To assess psychedelic potential for the 25N series, seventeen compounds were tested in the HTR assay based on their diverse range of Gq and β-arrestin efficacies. Male mice were used to maintain consistency with the validation experiments supporting use of the HTR as a cross-species readout of psychedelic potential^9, 36^ and to leverage the large dataset of HTR dose-response data previously generated in male mice. When tested in C57BL/6J mice, eleven of the compounds increased HTR counts significantly over baseline levels (Supplementary Tables 15 and 16). Not surprisingly, 25N-NBOMe (4), which acts as a psychedelic in humans^37^, produced a potent HTR response consistent with this effect (Fig. 4A). By contrast, the β-arrestin2-biased compounds 25N-N1-Nap (16), 25N-NBPh (17) and 25N-NB-2-OH-3-Me (18) failed to induce the HTR (Fig. 4B-D; Supplementary Tables 15,16), suggesting Gq efficacy is necessary for psychedelic potential. To confirm the β-arrestin2-biased agonists can partition into the brain and engage central 5-HT_2A_R necessary for the HTR, we pretreated mice with those compounds, which subsequently blocked the HTR induced by psychedelic 5-HT_2A_R agonist DOI (Fig. 4E-H). 25N-NBPh (17) had lower potency than 25N-N1-Nap (16) in the blockade experiments, which we believe is due to pharmacokinetic differences limiting its CNS distribution, potentially reflecting its higher cLogP (4.8 vs. 4.5, respectively). We thus focused on 25N-N1-Nap (16) in subsequent experiments. As expected, 25N-N1-Nap (16) and 25N-NBPh (17) antagonized 5-HT_2A_R-Gq signaling in vitro with potency similar to the 5-HT_2A_R-selective antagonist M100907 (Fig. 4I). Given the potential for species differences, which is evident for rodent versus human 5-HT_2A_R^38, 39^, we verified that the β-arrestin bias exhibited by 25N-N1-Nap (16) and 25N-NBPh (17) is preserved at mouse 5-HT_2A_R (Supplementary Fig. 9A).

**Figure 4.**
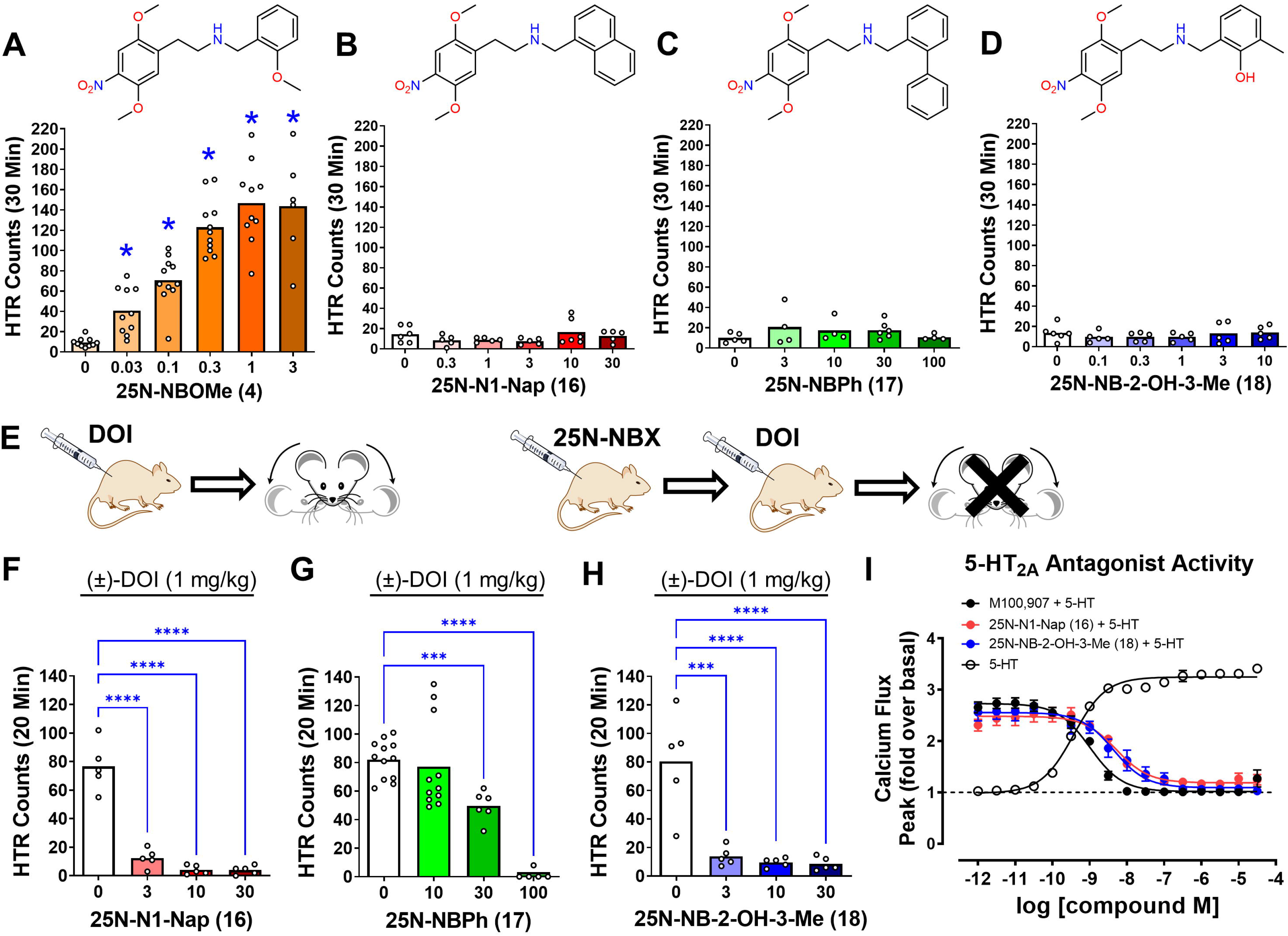
β-Arrestin-biased 5-HT_2A_ agonists lack psychedelic potential. **(A)** Effect of 25N-NBOMe (4) on the head-twitch response (HTR). \**p*<0.05, significant difference vs. vehicle control (Dunnett’s T3 test). **(B)** Effect of 25N-N1-Nap (16) on the HTR. **(C)** Effect of 25N-NBPh (17) on the HTR. **(D)** Effect of 25N-NB-2-OH-3-Me (18) on the HTR. **(E)** Illustration showing the procedures used to confirm that compounds tested in panels F–H are brain penetrant and capable of engaging 5-HT_2A_ receptors in the CNS of mice. Mice were pretreated with vehicle or drug, (±)-DOI (1 mg/kg IP) was injected 10 minutes later, and then HTR activity was assessed for 20 minutes. **(F)** Pretreatment with 25N-N1-Nap (16) blocks the HTR induced by (±)-DOI (*F*_3,16_ = 68.43, *p*<0.0001). **(G)** Pretreatment with 25N-NBPh (17) blocks the HTR induced by (±)-DOI (*W*_3,14.64_ = 144.6, *p*<0.0001). **(H)** Pretreatment with 25N-NB-2-OH-3-Me (18) blocks the HTR induced by (±)-DOI (*F*_3,16_ = 18.39, *p*<0.0001). \**p*<0.05, \*\*\**p*<0.001, \*\*\*\**p*<0.0001, significant difference between groups (Tukey’s test or Dunnett’s T3 test). HTR counts from individual male C57BL/6J mice as well as group means are shown. Drug doses are presented as mg/kg. **(I)** 5-HT_2A_ receptor Gq-mediated calcium flux activity comparing 5-HT (open circles) antagonist activity for 25N-N1-Nap (16) (red), 25N-NB-2-OH-3-Me (18) to M100,907 (black circles). BRET data represent the mean and SEM from three independent experiments.

To confirm that β-arrestin-biased 5-HT_2A_R agonists do not induce the HTR and that this profile is not specific to the 25N series, we synthesized and tested *N*-naphthyl and *N*-biphenyl derivatives of other phenethylamine 5-HT_2A_R agonists. Similar to the previous β-arrestin-biased compounds, these compounds exhibited weak Gq efficacy but had robust efficacy for β-arrestin2 recruitment (Supplementary Fig. 9B; Supplementary Table 13 and 14). When evaluated in mice, 25O-N1-Nap (28) and 2C2-N1-Nap (29) failed to induce head twitches (Supplementary Fig. 9C) but fully blocked the HTR induced by DOI (Supplementary Fig. 9D), indicating brain penetration. In summary, five different β-arrestin2-biased 5-HT_2A_R agonists did not induce the HTR when tested at doses that block the response to a balanced agonist, confirming this strategy can consistently generate β-arrestin2-biased 5-HT_2A_R agonists devoid of psychedelic potential.

### 5-HT2A-Gq signaling predicts psychedelic potential

Although the 5-HT_2A_R signaling pathways responsible for psychedelic potential have been previously investigated, the results were inconclusive^40–42^. To assess further the involvement of 5-HT_2A_R Gq and β-arrestin2 signaling in the HTR, correlation analyses were performed using the 25N series, which contains a mixture of HTR-active and inactive compounds. For the 25N compounds with Gq and β-arrestin2 data (*n*=14), there was robust correlation between HTR magnitude (the maximum number of HTR induced by each drug in counts/minute) and 5-HT_2A_R-Gq efficacy (%5-HT E_MAX_; *R*_S_ = 0.8242, *p* = 0.0005; Fig. 5A). By contrast, HTR magnitude was not correlated with β-arrestin2 recruitment (*R*_S_ = -0.01538, *p* = 0.9638; Fig. 5B). Notably, the relationship between 5-HT_2A_R-Gq efficacy and HTR magnitude was nonlinear and none of the 25N derivatives with Gq E_MAX_ values <70% induced the HTR, potentially indicating that 5-HT_2A_R efficacy must exceed a strong Gq agonist threshold level in order to induce the behavior. Similar to the BRET Gq data, the magnitude of the HTR induced by 25N derivatives is correlated with their 5-HT_2A_R Gq/11-mediated calcium flux response (*R*_S_ = 0.8175, *p* = 0.0002; Supplementary Fig. 10A) and only the compounds with Gq-mediated calcium flux efficacy exceeding 70% 5-HT E_MAX_ induced head twitches, providing further evidence that 5-HT_2A_R-Gq efficacy is necessary for psychedelic potential.

**Figure 5.**
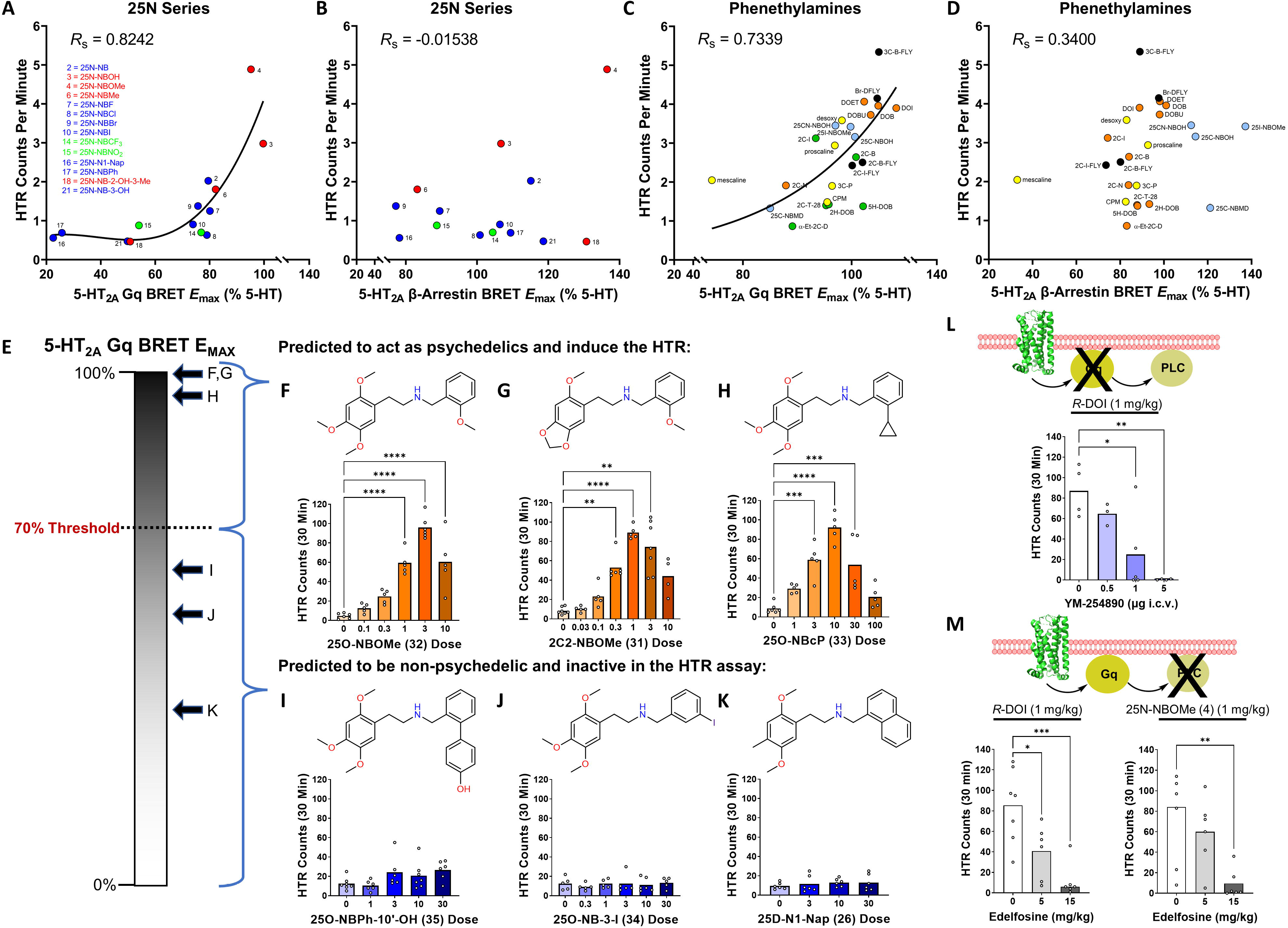
5-HT_2A_-Gq signaling predicts psychedelic potential. **(A)** Scatter plot showing the relationship between 5-HT_2A_ receptor Gq efficacy (E_MAX_ values) and head-twitch response (HTR) magnitude (maximum counts per minute induced by the compound) for 14 members of the 25N series. Spearman’s rank correlation coefficient *R*_s_ is shown. The regression line generated by fitting the data using non-linear regression is included as a visual aid. **(B)** Scatter plot showing the relationship between 5-HT_2A_ receptor β-arrestin2 recruitment efficacy (E_MAX_ values) and HTR magnitude for 14 members of the 25N series. **(C)** Scatter plot showing the relationship between 5-HT_2A_ receptor Gq efficacy (E_MAX_ values) and HTR magnitude for 24 phenethylamine psychedelics. To illustrate the non-linear nature of the relationship, the regression line generated by fitting the data using non-linear regression is shown. **(D)** Scatter plot showing the relationship between 5-HT_2A_ receptor β-arrestin2 recruitment efficacy (E_MAX_ values) and HTR magnitude for 24 phenethylamine psychedelics. **(E-K)** Activity in the HTR assay can be predicted based on 5-HT_2A_ receptor Gq efficacy. As predicted, 25O-NBOMe (32) (*F*_5,26_ = 37.01, *p*<0.0001), 2C2-NBOMe (31) (*W*_6,12.67_ = 96.28, *p*<0.0001), and 25O-NBcP (33) (*F*_5,25_ = 20.57, *p*<0.0001) induced head twitches, whereas 25O-NBPh-10’-OH (35) (*F*_4,27_ = 2.58, *p*=0.0601), 25O-NB-3-I (34) (*F*_5,25_ = 0.26, *p*=0.9288), and 25D-N1-Nap (26) (*F*_3,20_ = 0.37, *p*=0.7748) did not induce head twitches. **(L)** Pretreatment with the Gαq/11 inhibitor YM-254890 blocks the HTR induced by *R*-(-)-DOI (*F*_3,12_ = 8.69, *p*=0.0025). Mice were treated ICV with YM-254890 or vehicle and then all of the animals received *R*-(-)-DOI (1 mg/kg IP) 15 minutes later. **(M)** Pretreatment with the phospholipase C (PLC) inhibitor edelfosine blocks the HTR induced by *R*-(-)-DOI (*F*_2,16_ = 11.34, *p*=0.0009) and 25N-NBOMe (4) (*F*_2,16_ = 6.40, *p*=0.0091). Mice received three consecutive injections of vehicle or the indicated dose of edelfosine at 20-minute intervals and then a HTR-inducing drug was administered 10 minutes after the third injection. HTR counts from individual male C57BL/6J mice as well as group means are shown. \**p*<0.05, \*\**p*<0.01, \*\*\**p*<0.001, \*\*\*\**p*<0.0001, significant difference vs. vehicle control (Tukey’s test or Dunnett’s T3 test).

We also tested whether a similar relationship exists for drug potencies (ED_50_) in HTR experiments. Although the *in vivo* potencies of psychedelic drugs are known to be correlated with their *in vitro* potencies at 5-HT_2A_R^43–45^, it is not clear whether a similar relationship exists for the HTR. Notably, we observed robust and highly significant correlations between potencies in the HTR assay and *in vitro* 5-HT_2A_R potency measures, including binding affinities (*K*_i_) measured using [^3^H]-ketanserin, and functional potencies (EC_50_) for activating Gq and β-arrestin2 via 5-HT_2A_R (Supplementary Fig. 10B-F). Similar correlation results were obtained for 5-HT_2A_R Gq and β-arrestin2 pathways, which is not surprising because the rank-order potencies of 5-HT_2A_R agonists for Gq and β-arrestin2 are similar, whereas agonist efficacies often diverge across those two pathways. Overall, these results are consistent with the known role of 5-HT_2A_R in the HTR and show a good correlation between *in vitro* and *in vivo* potency measures, but they do not link the HTR to a particular transducer pathway because they fail to account for 5-HT_2A_R efficacy at each respective transducer.

Encouraged by results obtained with the 25N series, we examined whether the magnitude of the HTR produced by other psychedelics is correlated with their 5-HT_2A_R-Gq efficacy. To test this hypothesis, we compared 5-HT_2A_R efficacies and HTR magnitude for 24 phenethylamine psychedelics (Supplementary Fig. 11; Supplementary Tables 11, 12 and 18), which tend to have greater 5-HT_2_ selectivity compared to other scaffolds^10^. Our results show a significant positive relationship between HTR magnitude and 5-HT_2A_R-Gq efficacy (*R*_S_ = 0.7339, *p*<0.0001; Fig. 5C). Conversely, there is not a significant correlation between HTR magnitude and 5-HT_2A_R-β-arrestin2 efficacy (*R*_S_ = 0.34, *p*=0.104; Fig. 5D). Consistent with the 25N series data, all of the tested psychedelics activated Gq with relatively high efficacy and the trendline between HTR magnitude and Gq efficacy indicates a minimum of 70% 5-HT_2A_R-Gq efficacy (%5-HT E_MAX_). In summary, our results indicate there is a threshold level of 5-HT_2A_R-Gq efficacy required for psychedelic-like activity.

To test the hypothesis that HTR activity and psychedelic potential can be predicted based on the existence of a 5-HT_2A_R-Gq efficacy threshold, we synthesized *N*-benzyl derivatives of other phenethylamines and made predictions about their HTR activity based on their 5-HT_2A_R-Gq efficacy (Fig. 5E). We synthesized and tested 2C2-NBOMe (31) (E_MAX_ = 98.4%), 25O-NBOMe (32) (E_MAX_ = 99.9%), and 25O-NBcP (33) (E_MAX_ = 96.3%), which are highly efficacious 5-HT_2A_R-Gq agonists (Supplementary Fig. 10G-I) and would be predicted to induce the HTR. By contrast, 25D-N1-Nap (26) (E_MAX_ = 34.3%), 25O-NB-3-I (34) (E_MAX_ = 51.7%), and 25O-NBPh-10’-OH (35) (E_MAX_ = 60.8%) have efficacy <70% (Supplementary Fig. 10J-L) and were predicted to be inactive in HTR. Indeed, robust HTR responses were measured for 2C2-NBOMe (31), 25O-NBOMe (32), and 25O-NBcP (33), but not for weaker Gq agonists, 25D-N1-Nap (26), 25O-NB-3-I (34), and 25O-NBPh-10’-OH (35) (Fig. 5F-K; Supplementary Table 19). These results show that the ability of 5-HT_2A_R agonists to induce head twitches can be predicted based on their 5-HT_2A_R-Gq efficacy.

To directly test the involvement of 5-HT_2A_R-Gq signaling in the HTR, we evaluated whether inhibition of Gq signaling can block the HTR. Because the use of signaling inhibitors can potentially be confounded by off-target effects, we targeted two different proteins within the Gq-PLC effector pathway to generate convergent evidence. YM-254890 selectively inhibits Gq/11^46^ and is effective in mice when administered systemically or centrally^47^. Intracranial (ICV) pretreatment with YM-254890 blocked the HTR induced by DOI (Fig. 5L). We also tested the phosphoinositide-selective PLC inhibitor edelfosine^48^, which is brain penetrant in mice, with a brain/plasma ratio of ∼0.5 after systemic administration^49^. Edelfosine blocked the HTR induced by 25N-NBOMe (4) and DOI (Fig. 5M). These results strongly support the conclusion that the HTR induced by psychedelics is dependent on activation of the Gq-PLC pathway. This dependence on 5-HT_2A_-Gq signaling is supported by the lack of a HTR with 25N-N1-Nap (16) and other β-arrestin2-biased 5-HT_2A_R agonists, and it does not appear that activation of β-arrestin2 by 5-HT_2A_R agonists is sufficient to induce head twitches. Although YM-254890 inhibited the response to DOI to a greater degree than Gq gene deletion^42^, the ability of both manipulations to dampen the response to DOI strongly supports our conclusions that 5-HT_2A_R-Gq signaling is necessary for the HTR and therefore psychedelic potential.

### Non-psychedelics do not achieve 5-HT_2A_ Gq-signaling efficacy threshold

The existence of an efficacy threshold for activity is intriguing because it potentially explains why certain 5-HT_2A_R agonists such as lisuride fail to induce head twitches in mice and psychedelic effects in humans. To evaluate that possibility, we compared the activity of four psychedelic and four non-psychedelic 5-HT_2A_R agonists in BRET assays and HTR experiments. Clinical studies have confirmed that lisuride^50–53^, 2-Br-LSD ^54,55,56^, and 6-F-DET ^57^ lack psychedelic effects in humans; 6-MeO-DMT has not been evaluated clinically but does not substitute in rats trained to discriminate a psychedelic drug^58^. The psychedelic 5-HT_2A_R agonists LSD, psilocin, 5-MeO-DMT, and DET activate 5-HT_2A_R-Gq signaling with E_MAX_ ranging from 74.1–98.8% (Fig. 6A) and induce the HTR (Fig. 6B). Conversely, the non-psychedelic 5-HT_2A_R agonists lisuride, 2-Br-LSD, 6-F-DET, and 6-MeO-DMT activate 5-HT_2A_R-Gq signaling with E_MAX_ <70% (Fig. 6A) and do not induce head twitches (Fig. 6B). For the eight compounds, there was a significant positive correlation (*R* = 0.8948, *p* = 0.0027) between HTR magnitude and 5-HT_2A_R-Gq E_MAX_ (Fig. 6C). These results are consistent with the predicted activity threshold and demonstrate that the relationship between HTR magnitude and 5-HT_2A_R-Gq efficacy extends to tryptamines and lysergamides. In summary, lisuride, 2-Br-LSD, 6-F-DET, and 6-MeO-DMT are likely non-psychedelic because they have relatively weak Gq efficacy at 5-HT_2A_R.

**Figure 6.**
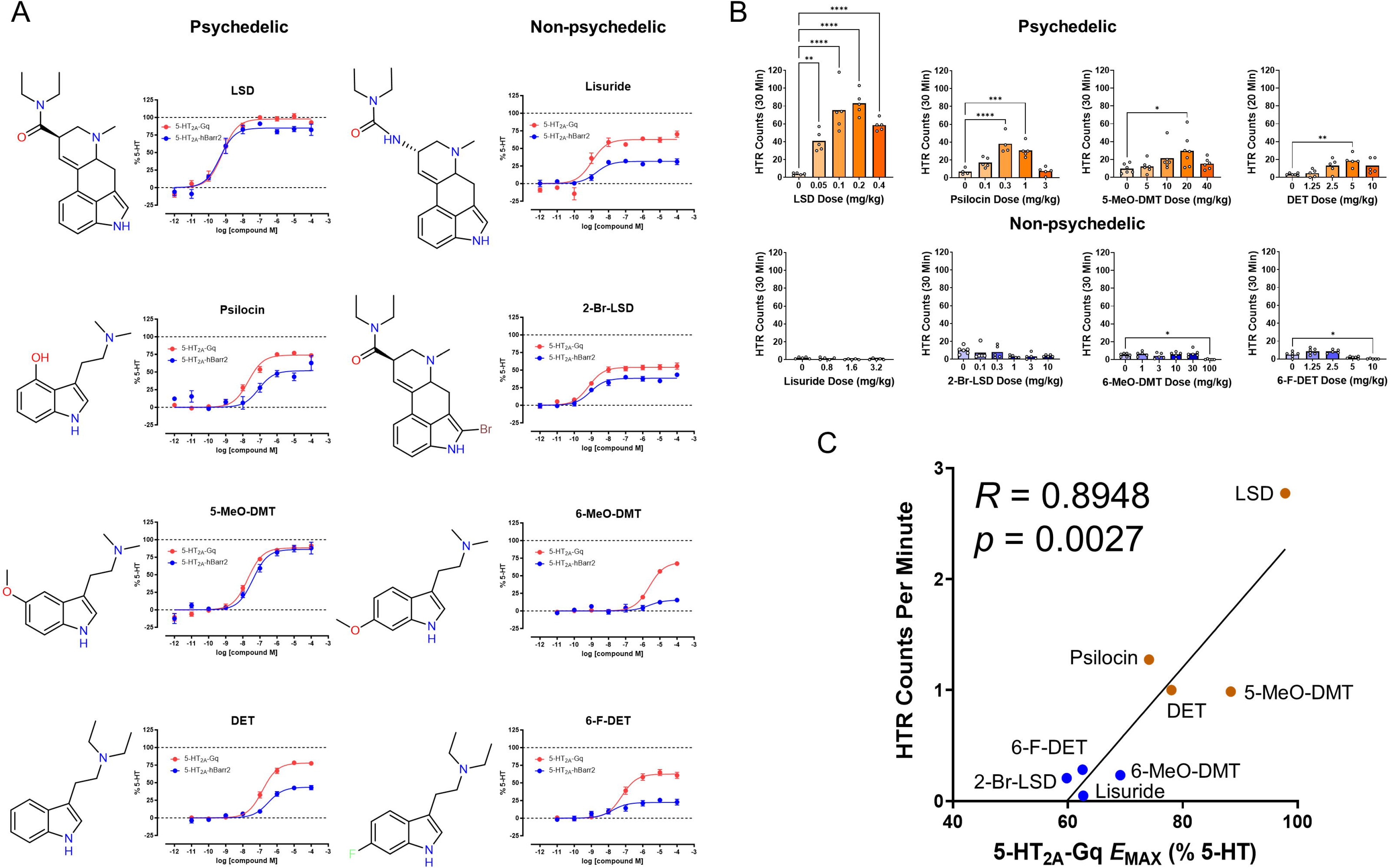
Non-psychedelics do not achieve 5-HT_2A_ Gq-signaling efficacy threshold. **(A)** Effect of (+)-lysergic acid diethylamide (LSD), psilocin, 5-methoxy-*N,N*-dimethyltryptamine (5-MeO-DMT), *N,N*-diethyltryptamine (DET), lisuride, (+)-2-bromolysergic acid diethylamide (2-Br-LSD), 6-methoxy-*N,N*-dimethyltryptamine (6-MeO-DMT), and 6-fluoro-*N,N*-diethyltryptamine (6-F-DET) on 5-HT_2A_ receptor Gq dissociation (red) and β-arrestin2 (blue) recruitment. The BRET data for LSD, psilocin, and 5-MeO-DMT are from Fig. 1; the data for 2-Br-LSD were published previously^11^. Data represent the mean and SEM from three independent experiments. **(B)** Effect of LSD, psilocin, 5-MeO-DMT (*F*_4,26_ = 3.09, *p*=0.0331), DET, lisuride, 2-Br-LSD, 6-MeO-DMT (*F*_5,26_ = 4.19, *p*=0.0063), and 6-F-DET (*F*_4,23_ = 13.49, *p*<0.0001) on the head-twitch response (HTR) in mice. The HTR data for LSD, DET, psilocin, lisuride, and 2-Br-LSD were re-analyzed from published experiments ^9,11, 80, 89^. HTR counts from individual male C57BL/6J mice as well as group means are shown. \**p*<0.05, \*\**p*<0.01, \*\*\**p*<0.001, \*\*\*\**p*<0.0001, significant difference between groups (Tukey’s test). **(C)** Scatter plot and linear regression showing the correlation between 5-HT_2A_ receptor Gq efficacy (E_MAX_ values) and HTR magnitude (maximum counts per minute induced by each drug). Pearson’s correlation coefficient *R* is shown.

### *In vitro* and *in vivo* effects of β-arrestin-biased 5-HT_2A_ agonists

An intriguing question raised by our results is how β-arrestin-biased 5-HT_2A_R agonists differ from 5-HT_2A_R antagonists. One hallmark of β-arrestin-biased agonists is the induction of arrestin-dependent internalization and downregulation, which contributes to drug tolerance^59^. To determine whether 25N-N1-Nap (16) and 25N-NBPh (17) induce β-arrestin2-dependent receptor internalization, which would further support their β-arrestin-biased agonist profile, we conducted NanoBit internalization assays^60, 61^ that utilize a minimally tagged N-terminal HiBiT-tagged 5-HT_2A_R and a membrane impermeable LgBit Nanoluc complementation fragment to measure surface expression loss (Fig. 7A). In experiments where untagged β-arrestin2 was co-expressed (similar conditions to β-arrestin2 BRET assays), 5-HT and DOI induced a strong internalization after 60 minutes. 25N-N1-Nap (16) and 25N-NBPh (17) also induced robust internalization, similar to their potencies in β-arrestin2 recruitment assays (Fig. 7B). By contrast, the 5-HT_2A_R antagonist/inverse agonist pimavanserin (PIM) did not induce internalization, suggesting that β-arrestin-biased 5-HT_2A_R agonists are distinct from pimavanserin in their ability to downregulate 5-HT_2A_R. Previous work in whole cell and *in vivo* systems revealed that some 5-HT_2A_R antagonists cause atypical 5-HT_2A_R internalization, with several proposed mechanisms including direct 5-HT_2A_R mediated, downstream, and off-target effects^62, 63^, though interestingly other 5-HT_2A_R antagonists such as eplivanserin (SR 46349B) upregulate 5-HT_2A_R in mice^64^. Thus, how these 5-HT_2A_R ligands affect *in vivo* receptor expression may be influenced by several factors and requires further study.

**Figure 7.**
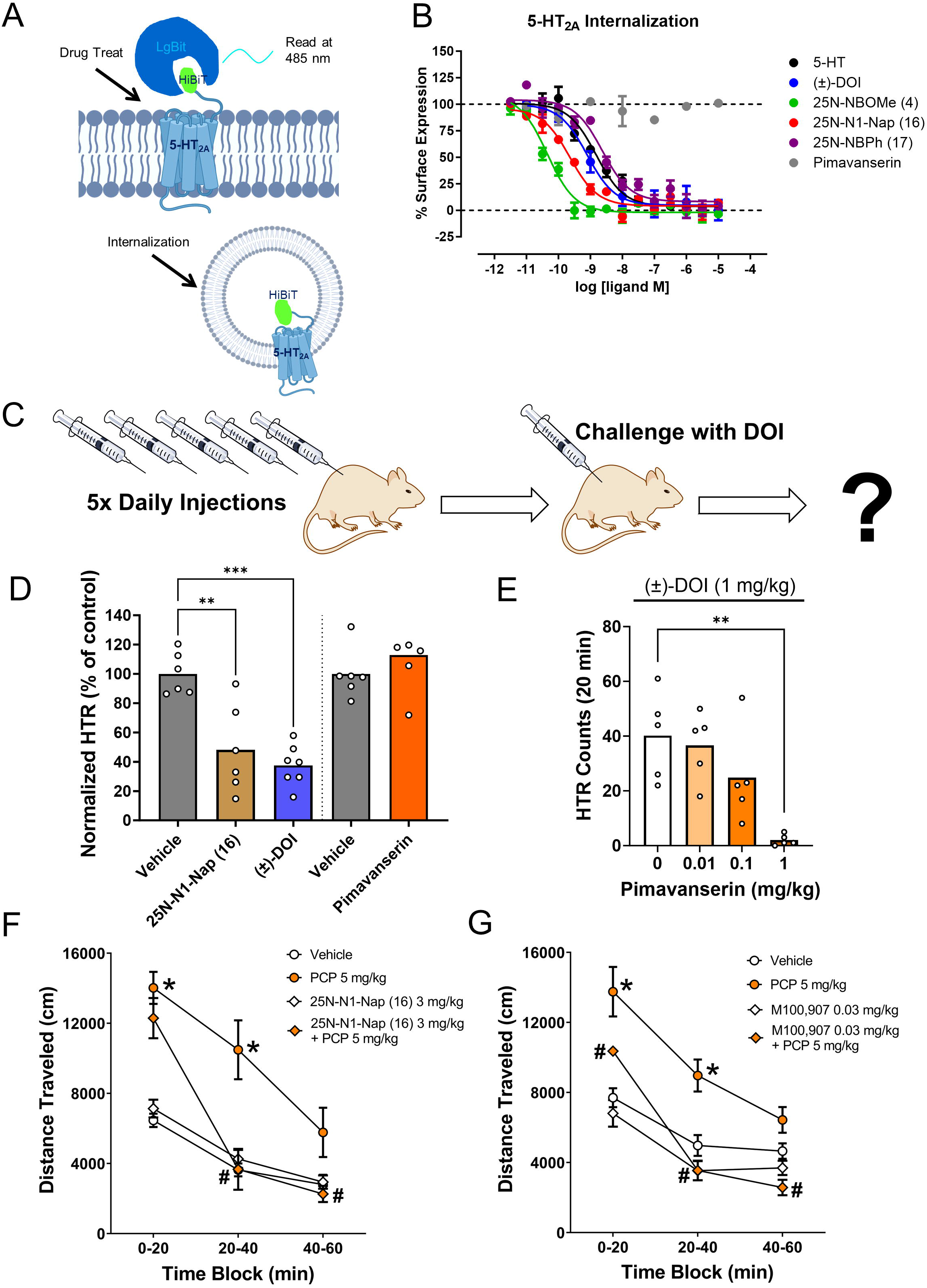
*In vitro* and *in vivo* effects of β-arrestin-biased 5-HT_2A_ agonists. **(A)** Scheme of NanoBit Internalization Assay for 5-HT_2A_ receptor measuring loss of surface expression with the membrane impermeable LgBit. **(B)** 5-HT_2A_ receptor internalization concentration response as calculated as percent basal surface expression and normalized to 0% of max 5-HT response. Data represent mean and SEM from 3 independent experiments with 5-HT (black), DOI (blue), pimavanserin (grey), 25N-NBOMe (4) (green), 25N-N1-Nap (16) (red) and 25N-NBPh (17) (purple). **(C)** Cartoon showing the procedures used to induce tolerance in the head-twitch response (HTR) assay. Mice were injected with vehicle or drug once daily for 5 consecutive days and then challenged with (±)-DOI (1 mg/kg/day IP) 24 hours after the last injection. **(D)** Repeated daily administration of (±)-DOI and 25N-N1-Nap (16), but not pimavanserin, induces a tolerance to the HTR induced by DOI. One set of mice (*n*=6-7/group) was treated with vehicle, 25N-N1-Nap (16) (20 mg/kg/day SC), or (±)-DOI (10 mg/kg/day SC) once daily for 5 consecutive days. A second set of mice (*n*=6/group) was treated with vehicle or pimavanserin (1 mg/kg/day SC) once daily for 5 consecutive days. HTR counts are expressed as a percentage of the response in the respective vehicle control group; data from individual male C57BL/6J mice as well as group means are shown. \*\**p*<0.01, \*\*\**p*<0.001, significant difference between groups (Tukey’s test). **(E)** Pretreatment with the 5-HT_2A_ receptor antagonist pimavanserin blocks the HTR induced by (±)-DOI (*F*_3,16_ = 8.19, *p*=0.0016). HTR counts from individual mice as well as group means are shown. \*\**p*<0.01, significant difference between groups (Tukey’s test). **(F)** Pretreatment with 25N-N1-Nap (16) attenuates phencyclidine (PCP)-induced locomotor hyperactivity (25N-N1-Nap (16) × PCP: *F*_1,20_ = 7.09, *p*=0.015; 25N-N1-Nap (16) × PCP × time: *F*_2,40_ = 11.03, *p*=0.0002). Locomotor activity was measured as distance traveled in cm. \**p*<0.05, significant difference vs. vehicle control; ^#^*p*<0.05, significant difference vs. PCP alone (Tukey’s test). **(G)** M100907 pretreatment attenuates PCP-induced locomotor hyperactivity (M100907 × PCP: *F*_1,20_ = 7.41, *p*=0.0131). \**p*<0.05, significant difference vs. vehicle control; ^#^*p*<0.05, significant difference vs. PCP alone (Tukey’s test). Distance traveled in male C57BL/6J mice are presented as group means ± SEM. Drug doses are presented as mg/kg.

5-HT_2A_R agonists are known to cause receptor downregulation and tolerance or tachyphylaxis^65^. We tested whether β-arrestin2-biased 5-HT_2A_R agonists induce tolerance *in vivo*, based on the rationale that tolerance could potentially act as an *in vivo* readout of β-arrestin2 recruitment. Our hypothesis was that repeated treatment with 25N-N1-Nap (16) would induce tolerance similar to the pathway-balanced 5-HT_2A_R agonist DOI, whereas an inverse agonist like PIM which lacks β-arrestin2 recruitment (Fig 7A) would not. The results (Fig. 7D) were normalized to the vehicle control group to facilitate comparison of the amount of tachyphylaxis induced by the three 5-HT_2A_R ligands. Once daily administration of 25N-N1-Nap (16) or DOI to mice for five consecutive days reduced the ability of a challenge dose of DOI administered 24 h later to induce the HTR (Fig. 7C,D). Conversely, the response to DOI was not altered in mice treated repeatedly with a dose of the 5-HT_2A_R antagonist PIM that is capable of blocking the acute behavioral response to DOI (Fig. 7E). Hence, repeated administration of 25N-N1-Nap (16), DOI, and PIM produced effects in mice that closely parallel the *in vitro* internalization data. Based on these results, β-arrestin2 may play a role in the tachyphylaxis that occurs after repeated administration of 5-HT_2A_R agonists.

Finally, to assess whether β-arrestin-biased 5-HT_2A_R agonists may have similar therapeutic potential as 5-HT_2A_R antagonists/inverse agonists, we tested whether 25N-N1-Nap (16) can antagonize PCP-induced hyperactivity in mice. 5-HT_2A_R antagonists are highly effective at blocking the hyperlocomotion induced by NMDA receptor antagonists such as PCP and MK-801^17, 66^. 25N-N1-Nap (16) blocked the hyperactivity induced by PCP in mice when tested at a dose that had no effect on baseline activity (Fig. 7F). We confirmed that the 5-HT_2A_-selective antagonist M100907 has a similar effect on reducing PCP-induced hyperactivity in mice (Fig. 7G) consistent with an antipsychotic-like profile.

## Discussion

Our ultimate aim was to elucidate which 5-HT_2A_R-coupled transducer pathway is responsible for psychedelic activity. Here we show that rational and structure-based design can be leveraged to develop 5-HT_2A_R-selective compounds with a wide-range of functional activities, including β-arrestin2-biased agonists. Docking, MD, and mutagenesis revealed that W336^6.48^ plays a key role in the mechanism of β-arrestin2-biased agonists. Finally, β-arrestin2-biased 5-HT_2A_R agonists were used to probe the involvement of specific signaling pathways in the psychedelic potential of 5-HT_2A_R agonists. The ability of 5-HT_2A_R agonists to induce the HTR was found to be correlated with Gq efficacy but not β-arrestin2 recruitment. Compounds exhibiting a signaling bias for β-arrestin2 did not induce psychedelic-like behavioral effects, but were capable of blocking psychedelic-like behaviors *in vivo*. Overall, we identified multiple structural and chemical features of psychedelics that can be targeted to fine-tune 5-HT_2A_R activity, potentially allowing the therapeutic efficacy and tolerability of 5-HT_2A_R ligands to be optimized for various therapeutic indications including psychosis.

Although the specific signaling cascades mediating the HTR have not been conclusively identified, Gq and β-arrestin2 are clear candidates. Previous studies in constitutive knockout (KO) mice have attempted to address the role of Gq and β-arrestin2 in the HTR but did not yield conclusive results. Knocking out Gq attenuated but did not eliminate the HTR induced by DOI^42^, potentially because other Gq protein subtypes such as G11 may also be involved. HTR studies in β-arrestin2 KO mice have variously reported that the response to DOI is unaltered^41, 67^, the response to 5-MeO-DMT is enhanced^68^, and the response to LSD is attenuated^40^. Although the reason for these discrepancies is unknown, it could reflect altered membrane receptor trafficking in β-arrestin2 KO mice, leading to constitutive receptor desensitization and other adaptations. Given the lack of an effect of 25N-N1-Nap (16) and other β-arrestin2-biased 5-HT_2A_R agonists in the HTR paradigm, activation of β-arrestin2 by 5-HT_2A_R agonists does not appear to be sufficient to induce head twitches. Although β-arrestin2 may not be involved in the HTR, our findings do implicate β-arrestin2 in the tachyphylaxis that occurs after repeated administration of 5-HT_2A_R agonists. Mice treated repeatedly with psychedelic drugs develop tolerance and cross-tolerance, potentially reflecting 5-HT_2A_R downregulation^67^. Similarly, repeated treatment with 25N-N1-Nap (16) induced a tolerance to a subsequent challenge dose of DOI, indicating β-arrestin2-biased agonists may also be capable of inducing 5-HT_2A_R downregulation. Although it was recently reported that β-arrestin2 KO mice do show tolerance to DOI-induced HTR after repeated treatment^67^, the use of constitutive KOs to address the mechanisms of GPCR tolerance is not ideal because of potential adaptations that may occur to offset the loss of β-arrestin2 in a global genetic knock-out model.

One important result of these studies is the discovery of an apparent 5-HT_2A_R efficacy threshold for induction of the HTR in mice. In addition to finding a robust and highly significant correlation between the magnitude of the HTR and multiple readouts of 5-HT_2A_R-Gq efficacy, we found that compounds with a Gq E_MAX_ <70% do not induce the HTR. 5-HT_2A_R efficacy was found to be correlated with HTR magnitude in previous studies of mixed 5-HT_2A/2C_ agonists^69, 70^, but those studies did not test enough compounds to identify a clear efficacy threshold for HTR activity. Since mice exhibit a baseline level of spontaneous HTR that is driven by basal 5-HT_2A_R activation^71^, the existence of a threshold for HTR activity is not surprising because compounds with low efficacy may act as partial antagonists relative to endogenous 5-HT basal stimulation. A similar threshold may exist for psychedelic effects in humans because all of the psychedelics we tested (including LSD and tryptamines) activated 5-HT_2A_R-Gq signaling with E_MAX_ >70%, whereas non-psychedelic analogs have lower efficacy (E_MAX_ <70%) and likely explain why the latter molecules do not induce the HTR in mice or psychedelic effects in humans. Consistent with our findings, lisuride (E_MAX_ = 48.6%) was reported to have substantially lower efficacy than LSD (Emax = 84.6%) or DOI (E_MAX_ = 81.3%) in a 5-HT_2A_R-Gq calcium mobilization assay^72^. Although we did not test the putative non-psychedelic 5-HT_2A_R agonist tabernathalog, it reportedly has E_MAX_ = 57% in a 5-HT_2A_R calcium flux assay^5^, which may be too low to induce the HTR. These data have implications for drug development as it should be possible to identify 5-HT_2A_R-Gq partial agonists that do not induce the HTR and lack strong psychedelic effects in humans but retain sufficient efficacy to induce therapeutic neurophysiological effects via 5-HT_2A_R (e.g., induction of neuroplasticity). 2-Br-LSD seems to have a profile consistent with this hypothesis because it induces neuroplasticity via 5-HT_2A_R even though its 5-HT_2A_R-Gq E_MAX_ is subthreshold to induce the HTR^11^. Based on these results, it should be possible to rationally design non-psychedelic 5-HT_2A_R agonists with therapeutic potential by fine-tuning their 5-HT_2A_R-Gq efficacy. In effect, partial 5-HT_2A_R-Gq agonists may act as mixed agonist-antagonists, similar to buprenorphine and other opioid receptor partial agonists^73^. The partial agonism of buprenorphine is believed to contribute to its superior safety profile and greater tolerability^17^. Future studies will also need to examine partial 5-HT_2A_R-Gq agonists in preclinical disease models to assess whether they possess therapeutic-like activity, for example as rapid-acting antidepressants. The potential therapeutic utility of β-arrestin-biased 5-HT_2A_R agonists needs further study, but our findings suggest they mimic psychedelics in some ways (induction of cross-tolerance and 5-HT_2A_R downregulation) but also produce antagonist-like effects (e.g. blockade of PCP hyperlocomotion and DOI-induced HTR). Potentially, β-arrestin-biased ligands may mimic the therapeutic effects of 5-HT_2A_R antagonists with less potential to disrupt cognition, possibly improving their efficacy and tolerability.

Based on our results, ligand interactions with residue W336^6.48^ are an important predictor of agonists with weaker Gq efficacy but do not fully explain the β-arrestin bias exhibited by these compounds. Residue W6.48 is known to be critical in propagating conformational changes by altering the position of TM6 to impact the cytosolic transducer binding pockets and influence bias^34^, but the change in W336^6.48^ in our 25N-N1-Nap (16) simulation can likely influence other trigger motifs (PIF and NPxxY), which may be important to preserving GPCR-arrestin recruitment. For example, an influence of W6.48 on the PIF/PIW motif and corresponding changes in TM6 orientation were observed in MD studies with bias at β_2_-adreneregic^32^, rhodopsin^74^, MOR^34^, 5-HT_2C_R^75^, S1PR1^76^, and other GPCRs^35^. Moreover, our MD results with the “partially active” TM6 orientation are intriguing given several arrestin-1 GPCR structures, including LSD-bound 5-HT_2B_R structures^77^, show a larger TM6 outward movement relative to the G protein-bound state, though differences may exist in the initial conformation recognized by the transducer and the resulting complex. Therefore, further structural studies are needed to identify other conformational changes involved in Gq versus arrestin-modulation of the binding pocket, especially with respect to these discovered biased ligands.

In conclusion, we have shown that psychedelic drugs activate both 5-HT_2A_R-Gq and β-arrestin2 transducers, which led to us to design 5-HT_2A_R-selective biased agonists to probe the signaling pathways necessary for psychedelic potential. These results indicate that a threshold level of Gq activation is necessary to produce psychedelic-like effects, as measured by the HTR, and that it is possible to predict psychedelic potential based on the degree of 5-HT_2A_R-Gq efficacy. This study has implications for understanding the neurobiological basis of psychedelic effects and reveals strategies for designing non-psychedelic 5-HT_2A_R agonists that can potentially be used as therapeutics.

## METHODS

### Synthesis Materials and Methods

*N*-benzyl-compounds were synthesized by reductive amination (using NaBH_4_) of the pre-formed imine (in solution) obtained by treating the primary amine (e.g., 25N) with the respective aldehyde in dry (3Å molecular sieves) methanol and THF in the presence of 3 Å molecular sieves in the dark under an argon atmosphere for at least 48 hours. Descriptions, including yields and analytical data including ^1^H and ^13^C NMR chemical shift assignments, ^1^H and ^13^C NMR spectra, HPLC traces are provided in the Supplementary document.

Intermediates and reagents for synthesis were purchased from Sigma-Aldrich (St Louis, MO, USA), AKSci, and Alfa Aesar. Reagents were generally 95% pure or greater. 200 proof ethyl alcohol was obtained from Pharmaco (Greenfield Global, CT, USA). Silica gel flash column chromatography was performed using Merck silica gel grade 9385 (230-400 mesh, 60 Å). Melting points were measured using a Digimelt A160 SRS digital melting point apparatus (Stanford Research Systems, Sunnyvale, CA, USA) using a ramp rate of 2 °C/min.

### High Performance Liquid Chromatography (HPLC)

HPLC analyses were performed on an Agilent 1260 Infinity system that includes a 1260 quaternary pump VL, a 1260 ALS autosampler, a 1260 Thermostatted Column Compartment, and a DAD Multiple Wavelength Detector (Agilent Technologies, Santa Clara, CA, USA). Detection wavelengths were set at 220, 230, 254, and 280 nm but only 220 nm was used for analysis. A Zorbax Eclipse XDB-C18 analytical column (5 µm, 4.6 x 150 mm) from Agilent Technologies was used. Mobile phase A consisted of 10 mM aqueous ammonium formate buffer titrated to pH 4.5. Mobile phase B consisted of acetonitrile. The injection volume of samples was 10 µL, flow rate was 1.0 mL/min, and the column temperature was set at 25°C. Samples were prepared by preparing a 1 mg/mL solution in 1:1 A:B. All samples were injected in duplicate with a wash in between each run. Run time was 10 minutes with a mobile phase ratio (isocratic) of 1:1 for A:B. Chromatograms were analyzed using the Agilent ChemStation Software (Agilent Technologies). Purity values were calculated from area under the curves of the absorbance at 220 nm of any resulting peaks.

### High Resolution Mass Spectrometry (HRMS)

HRMS data were obtained on a Thermo Orbitrap Exactive Mass Spectrometer with an Orbitrap mass analyzer. The instrument was calibrated using electrospray ionization with Pierce^TM^ LTQ ESI Positive Ion Calibration Solution from ThermoFisher Scientific. Samples were introduced into the instrument and ionized via an Atmospheric Solids Analysis Probe (ASAP). Data was analyzed in the Thermo Xcalibur Qual Browser software and identity was confirmed if <5 ppm error. Measurement parameters were as follows: Aux gas flow rate-8, Spray Voltage-3.50 kV, Capillary temperature-275°C, Capillary Voltage-25.00 V, Tube Lens Voltage-65.00 V, Skimmer Voltage-14.00 V, Heater Temperature-100°C.

### Elemental Analysis

Elemental analysis (C, H, N) was run on select compounds by Galbraith Laboratories, Inc. (Knoxville, TN).

### Nuclear Magnetic Resonance

^1^H (400 MHz) and ^13^C NMR spectra (101 MHz) were obtained on the hydrochloride salts as a solution in anhydrous d_6_-DMSO (∼20 mg/mL) (>99.9% D, Sigma-Aldrich) on a Bruker Avance III with PA BBO 400S1 BBF-H-D-05 Z plus probe (Bruker Corporation, Billerica, MA, USA). Solvent (δ = 2.50 and 39.52 ppm for ^1^H and ^13^C spectra respectively) was used for internal chemical shift references. ^19^F (376.5 MHz) NMR was run as described above using ∼100 µL of trichlorofluoromethane (99%+, Sigma-Aldrich) as internal reference (δ = 0.0 ppm). Full NMR chemical shift assignments were made using chemical shift position, splitting patterns, ^13^C and ^13^C PENDANT or APT and hetero- and homo-2-D experiments including HMQC or HSQC, HMBC and COSY (45° pulse tilt). A background water concentration of solvent (determined as integration ratio relative to solvent shift) was also determined to check for water content (indicative of a hydrate) within the final salts for 25N compounds. Evidence of hydrates was not observed.

### Hammett σ Constants

Hammett σ constants are provided in Supplementary Table 3.

### Receptor Binding Experiments

Competition based receptor binding studies were performed by the National Institute of Mental Health Psychoactive Drug Screening Program (NIMH PDSP) using described methods ^78^. Target compounds were dissolved in DMSO and an initial screen performed to assess displacement of the radioligand at target receptors using a concentration of 10 µM. Those compounds which showed >50% displacement of specific radioligand binding for a given receptor then underwent secondary screenings at varying concentrations to determine pK_i_ values. For the K_i_ experiments, compounds were run in triplicate on separate plates. Each plate contained a known ligand of the receptor as a positive control. Full experimental details are available in the NIMH PDSP assay protocol book. For all 5-HT subtypes, an N=3-4 was performed except where noted (Supplementary Table 8) and a mean pK_i_ and SEM was calculated using these replicate experiments. Affinities at off-targets are provided in Supplementary Table 9,10.

### Gq-Dissociation and β-arrestin2 recruitment BRET assays

To measure 5-HT receptor-mediated β-Arrestin2 recruitment as measured by BRET^1^, HEK293T cells (ATCC CRL-11268; mycoplasma-free) were subcultured in DMEM supplemented with 10% dialyzed FBS (Omega Scientific) and were co-transfected in a 1:15 ratio with human or mouse 5-HT receptors containing C-terminal fused *renilla* luciferase (*R*Luc8), and a Venus-tagged N-terminal β-arrestin2 using 3:1 ratio of TransiT-2020 (Mirus). 5-HT_2A_ receptor mutants were designed and performed using Q5 mutagenesis kit (New England Biolabs). All DNA was sequence verified using Sanger nucleotide sequencing (Retrogen, San Diego, CA). To measure 5-HT receptor-mediated Gq activation via Gq/γ1 dissociation as measured by BRET^2^, HEK293T cells were subcultured in DMEM supplemented with 10% dialyzed FBS and were co-transfected in a 1:1:1:1 ratio with *R*Luc8-fused human Gαq (Gαq-*R*Luc8), a GFP^2^-fused to the C-terminus of human Gγ1(Gγ1-GFP^2^), human Gβ1, and 5-HT receptor using TransiT-2020, as described previously ^79^. After at least 18-24 hours, transfected cells were plated in poly-lysine coated 96-well white clear bottom cell culture plates in DMEM containing 1% dialyzed FBS at a density of 25-40,000 cells in 200 µl per well and incubated overnight. The next day, media was decanted and cells were washed with 60 µL of drug buffer (1× HBSS, 20 mM HEPES, pH 7.4), then 60 µL of drug buffer was added per well. Cells were pre-incubated at in a humidified atmosphere at 37°C before receiving drug stimulation. Drug stimulation utilized 30 µL addition of drug (3X) diluted in McCorvy buffer (1× HBSS, 20 mM HEPES, pH 7.4, supplemented with 0.3% BSA fatty acid free, 0.03% ascorbic acid) and plates were incubated at indicated time and temperatures. Substrate addition occurred 15 minutes before reading and utilized 10 µL of the *R*Luc substrate, either coelenterazine h for β-Arrestin2 recruitment BRET^1^ or coelenterazine 400a for Gq dissociation BRET^2^ (Prolume/Nanolight, 5 µM final concentration) and was added per well. Plates were read for luminescence at 485 nm and fluorescent eYFP emission at 530 nm for BRET^1^ and at 400 nm and fluorescent GFP^2^ emission at 510 nm for BRET^2^ at 1 second per well using a Mithras LB940 (Berthold) or a PheraStar FSX (BMGLabTech). The BRET ratios of fluorescence/luminescence were calculated per well and were plotted as a function of drug concentration using Graphpad Prism 5 or 9 (Graphpad Software Inc., San Diego, CA). Data were normalized to % 5-HT stimulation and analyzed using nonlinear regression “log(agonist) vs. response” to yield Emax and EC_50_ parameter estimates.

### Calcium Flux Assays

Stably expressing 5-HT receptor Flp-In 293 T-Rex Tetracycline inducible system (Invitrogen, mycoplasma-free) were used for calcium flux assays, as described and utilized previously ^80^. Cell lines were maintained in DMEM containing 10% FBS, 10 µg/mL Blasticidin (Invivogen), and 100 µg/mL Hygromycin B (GoldBio). Day before the assay, receptor expression was induced with tetracycline (2µL/mL) and seeded into 384-well poly-L-lysine-coated black plates at a density of 7,500 cells/well in DMEM containing 1% dialyzed FBS. On the day of the assay, the cells were incubated with Fluo-4 Direct dye (Invitrogen, 20 µl/well) for 1 h at 37°C, which was reconstituted in drug buffer (20 mM HEPES-buffered HBSS, pH 7.4) containing 2.5 mM probenecid. After dye load, cells were allowed to equilibrate to room temperature for 15 minutes, and then placed in a FLIPR^TETRA^ fluorescence imaging plate reader (Molecular Devices). Drug dilutions were prepared at 5X final concentration in drug buffer (20 mM HEPES-buffered HBSS, pH 7.4) supplemented with 0.3% BSA fatty-acid free and 0.03% ascorbic acid. Drug dilutions were aliquoted into 384-well plastic plates and placed in the FLIPR^TETRA^ for drug stimulation. Fluorescence for the FLIPR^TETRA^ were programmed to read baseline fluorescence for 10 s (1 read/s), and afterward 5 µl of drug per well was added and read for a total of 5-10 min (1 read/s). Fluorescence in each well was normalized to the average of the first 10 reads for baseline fluorescence, and then either maximum-fold peak increase over basal or area under the curve (AUC) was calculated. Either peak or AUC was plotted as a function of drug concentration, and data were normalized to percent 5-HT stimulation. Data was plotted and non-linear regression was performed using “log(agonist) vs. response” in Graphpad Prism 9 to yield Emax and EC_50_ parameter estimates.

### Surface Expression/Internalization Experiments

Surface expression was measured using a HiBit-tagged 5-HT_2A_ receptor and the Nano-Glo HiBit Extracellular Detection System (Promega). N-terminal HiBit-tagged human 5-HT_2A_ receptor was cloned into pcDNA3.1 using Gibson Assembly. HEK293T cells (ATCC CRL-11268; mycoplasma-free) were transfected into 10-cm tissue culture dishes in a 1:15 ratio of HiBit-tagged human 5-HT_2A_ receptor: human β-Arrestin2 (cDNA Resource Center; www.cDNA.org). Cells were transfected in DMEM 10% dFBS and the next day, cells were plated into either poly-L-lysine-coated 96-well white assay plates (Grenier Bio-One). On the day of the assay, plates were decanted and HEPES-buffered DMEM without phenol-red (Invitrogen) was added per well. Plates were allowed to equilibrate at 37°C in a humidified incubator before receiving drug stimulation. Compounds (including 5-HT as control) were serially diluted in McCorvy buffer (20 mM HEPES-buffered HBSS, pH 7.4 supplemented with 0.3% BSA fatty-acid free and 0.03% ascorbic acid), and dilutions were added to plates in duplicate (96). Plates were allowed to incubate at 37°C for 1 hour in a humidified incubator or a specified time point. Approximately 15 minutes before reading, LgBit and coelenterazine h (5 uM final concentration) were added to each well. Plates were sealed to prevent evaporation and read on either a PheraStar FSX (BMB Lab Tech) or Mithras LB940 (Berthold Technologies) at 485 nm at 37°C for time-capture quantification of internalization or loss of surface expression. Luminescence was plotted as a function of drug concentration using Graphpad Prism 5 or 9 (Graphpad Software Inc., San Diego, CA). Data were analyzed using nonlinear regression “log(agonist) vs. response” to yield Emax and EC_50_ parameter estimates and normalized to % 5-HT surface expression, which a full concentration-response curve was present on every plate.

### Induced fit docking

Docking simulations of six of the 25N ligands, selected to assess the binding modes and provide potential insight into the SAR around signaling or receptor selectivity, were carried out against the active state human 5-HT_2A_ receptor (PDB: 6WHA) using the Induced-Fit Docking (IFD) protocol ^81^ of the Schrodinger Suite (2020a). IFD was run with extended sampling enabled, using the default setting of residues within 5 Å of the experimental ligand (25N-NBCN) and other parameters maintained at their default values, and ligand structures processed using the LIGPREP tool (target pH of 7) of the Schrödinger (release 2020c).

Poses generated in the IFD runs were analyzed using a custom Python script with PyMol visualization application (The PyMOL Molecular Graphics System, Version 2.0 Schrödinger, LLC).

### Estimation of ring substituent effects on molecular electrostatic potential

A series of model compounds were constructed that included only the benzylamine or naphthylamine portion of the corresponding ligand structure. Quantum mechanical optimization of the model compounds, in both neutral and cationic forms, was carried out using density functional theory (DFT) with the Jaguar tool of the Schrodinger suite using the B3LYP functional and 6-311g**++ basis set, with the exceptions of the models for 25N-NBI, where the halogen atom was not supported, and a smaller basis (3-21g*++) was substituted. We computed Hirshfeld partial atomic charges (Supplementary Table 4) for the model compounds using the optimized wavefunctions as a proxy for overall MEP. Hirshfeld charges are computed from a spatial partitioning of the electron density, and have been shown to correlate well with a variety of properties of aromatic molecules ^82^.

### Molecular Dynamics (MD) Simulations

Simulations were performed for each of two ligands (25CN-NBOH or 25N-N1-Nap (16) in complex with the human 5-HT_2A_ serotonin receptor in a lipid bilayer of 1,2-dipalmitoyl-sn-glycero-3-phosphocholine (DPPC) and SPC water with enough Cl^-^ and Na^+^ ions to neutralize the system at biologic salinity (0.15 M ionic strength). For the 25CN-NBOH system a model of the protein-ligand part of the system was made from chain A of the 6WHA PDB entry, downloaded from the OPM server ^83^. Missing extracellular loops were filled in with those from chain A of PDB entry 6WGT. Schrödinger’s Protein Preparation Wizard ^84^ was employed to use Prime ^85^ to fill in missing side chains, and to use Epik ^86^ to select the appropriate tautomers and protonation states of the protein and ligand. Due to anticipated difficulty modeling the long ICL3, the helices of TM5 and TM6 were capped, along with the N and C termini, with polar terminating residues: COOH for residue Q262^5.66^ and NH_2_ for I315^6.27^. During pilot simulations, entanglement of polar sidechains in the membrane occurred. To prevent this, it was found necessary that the rotamers for the loops and capped termini whose sidechains would otherwise interact with the polar heads of the lipid molecules be manually adjusted to ensure adequate water solvation at the beginning of the simulation. This was achieved by maximizing the magnitude of atomic coordinate z of terminal side-chain atoms, where the z-axis is perpendicular to the membrane and *z* = 0 locates the middle of the bilayer. Another issue identified during preliminary simulations was that DPPC tails would intrude into the orthosteric pocket between helices TM4 and TM5, causing the nitrile end of the ligand to project up toward the extracellular side of the pocket. On careful inspection, it was discovered that there is a stable position for a water molecule among residues D120^2.50^, S162^3.39^, and N376^7.49^. Manual introduction of a solvent molecule into this void volume successfully prevented lipid intrusion.

A disulfide bond between C148^3.25^ and C227^45.50 18^ was added to the topology, as well one between C349^6.61^ and C353^ECL3^. Each ligand was protonated at the basic nitrogen (as confirmed by Epik) and topology parameter files needed for subsequent dynamics simulation were determined using the ATB server [https://atb.uq.edu.au/]. Partial atomic charges for the ligands were computed via Schrödinger’s Jaguar tool using the density-functional method (B3LYP-D3 functional, 6-31G** basis), with discrete charges derived from the geometry-optimized wavefunction using the Hirshfeld approach (Supplementary Table 4). The total protein comprised 278 residues with 2,849 atoms. The remaining system consisted of 6,933 water molecules, 29 Cl^-^ ions, 20 Na^+^ ions, and 88 DPPC molecules. 25CN-NBOH has 44 atoms. The total number of atoms in the system is 28,137. The orthorhombic initial system dimensions were 61.319 × 60.911 × 97.412 Å^3^.

The protein-ligand complex for 25N-N1-Nap (16) starts with the structure from the top-ranked pose determined by induced fit docking (IFD). A chimeric homology model was created in Maestro using the IFD structure as the template everywhere except at the missing loops; these are taken from the 25CN-NBOH-system. This resulting protein-25N-N1-Nap (16) homology model was superposed (using the positions of the TM C_α_s) onto the protein-ligand complex in the full (water-ion-lipid-protein-ligand) unrelaxed construct built for the 25CN-NBOH simulation. The protein-25CN-NBOH complex was then removed and the 25N-N1-Nap (16)-protein complex substituted in its place. As the IFD docking involved minimal changes to backbone conformation, the resulting model for the 25N-N1-Nap (16) ligand had helix and loop positions nearly identical to those of the 25CN-NBOH model, permitting a direct structural substitution. That said, prior to any further computations, a restrained minimization was carried out with only the membrane/solvent system fully mobile, to remove any high-energy interatomic overlaps that might have been inadvertently produced by the alignment and substitution. The 25N-N1-Nap (16) simulation comprised a total of 28,143 atoms (50 atoms for the ligand and otherwise identical to the 25CN-NBOH system). The orthorhombic initial system dimensions were 61.864 × 61.453 × 96.12 Å^3^.

### Simulation protocols and analysis

Both sets of simulations had three phases: minimization (unrestrained steepest decent), equilibration, and a production run. The equilibration phases for both sets comprise a restrained NVT simulation followed by a restrained NPT one. In both, the protein and ligand atoms were restrained close to their initial positions by a harmonic potential with force constant *k* = 1000 kJ/mol/nm (23.9 kcal/mol/Å). Both production simulations were 250-ns NPT unrestrained MD simulations. The force field used was GROMOS96 54a7^87^. The MD engine used was GROMACS v2021.2. The energy parameters in common between the 25CN-NBOH and the 25N-N1-Nap (16) simulations can be found in Supplementary Table 23. The parameters that vary between stages and systems can be found in Supplementary Table 24.

The chi_2_ dihedral data were measured by GROMACS’s chi command every 10 ps. The histogram bin width was one degree. Plots were made in Grace v5.1.25. The time-series heat maps of the pocket-ligand nonbonded interaction energy were made with Seaborn and Matplotlib. The data were calculated by GROMACS’s mdrun command. The pocket residues were determined by Schrödinger’s Maestro (all residues within 5 Å of the ligand). Protein cartoons and geometric measurements were made in VMD ^88^ and Maestro.

### Animal Behavioral Experiments

Male C57BL/6J mice (6–8 weeks old) from Jackson Labs (Bar Harbor, ME, USA) were used for the behavioral experiments. The mice were housed on a reversed light-dark cycle (lights on at 1900 h, off at 0700 h,) in an AALAC-approved vivarium at the University of California San Diego. Mice were housed up to four per cage in a climate-controlled room and with food and water provided ad libitum except during behavioral testing. Testing was performed between 1000 and 1800 h (during the dark phase of the light-dark cycle). The studies were conducted in accordance with National Institutes Health (NIH) guidelines and were approved by the University of California San Diego Institutional Animal Care and Use Committee.

The drug solutions used for the behavioral experiments were prepared as follows: pimavanserin hemitartrate (MedChemExpress), 25C-NBOH hydrochloride, 3,4-dimethoxy-4-methylphenethylamine hydrochloride (desoxy), and 4-cyclopropyl-3,5-dimethoxyphenethylamine hydrochloride (CPM) were dissolved in sterile water; 6-fluoro-*N,N*-diethyltryptamine (6-F-DET; donated by the Usona Institute, Fitchburg, WI, USA) was dissolved in sterile water acidified with HCl to pH 5; YM-254,890 (FUJIFILM Wako Chemicals USA, Richmond, VA, USA) was dissolved in 100% dimethyl sulfoxide (DMSO); edelfosine (Tocris Bioscience, Minneapolis, MN, USA), (±)-2,5-dimethoxy-4-iodoamphetamine hydrochloride ((±)-DOI; Cayman Chemical, Ann Arbor, MI, USA), *R*-(–)-2,5-dimethoxy-4-iodoamphetamine hydrochloride (*R*-(–)-DOI; donated by the National Institute on Drug Abuse, Rockville, MD, USA), phencyclidine hydrochloride (PCP; Sigma-Aldrich), *N,N*-diethyltryptamine fumarate (DET; donated by the National Institute on Drug Abuse, Rockville, MD, USA), 5-methoxy-*N,N*-dimethyltryptamine hemifumarate (5-MeO-DMT; donated by the National Institute on Drug Abuse, Rockville, MD, USA), 6-methoxy-*N,N*-dimethyltryptamine hydrochloride (6-MeO-DMT), 25N-NBOMe hydrochloride, and 25N-NB hydrochloride were dissolved in isotonic saline; 25C-NBMD hydrochloride was dissolved in sterile water containing 2% Tween-80 (v/v); 25N-NBOEt hydrochloride, 25N-NB-2-OH-3-Me hydrochloride, 25N-NBPh hydrochloride, 25O-N1-Nap hydrochloride, 2C2-N1-Nap hydrochloride, 25O-NBOMe hydrochloride, 25O-NBPh-10’-OH hydrochloride, 25O-NB-3-I hydrochloride, 25O-NBcP hydrochloride, 2C-N hydrochloride, and M100907 were dissolved in sterile water containing 5% Tween-80 (v/v); 25N-NBNO_2_ hydrochloride was dissolved in sterile water containing 20% hydroxypropyl-β-cyclodextran (w/v); 25N-NBBr hydrochloride was dissolved in sterile water containing 5% Tween-80 (v/v) and 20% hydroxypropyl-β-cyclodextran (w/v); for the remaining 25N derivatives, the hydrochloride salts were dissolved in sterile water containing 1% Tween-80 (v/v). 25N derivatives, 25O-NBOMe, 25O-NBPh-10’-OH, 25O-NB-3-I, 25O-NBcP, 25O-N1-Nap, 2C2-NBOMe, 2C2-N1-Nap, and 25D-N1-Nap were injected SC (5 mL/kg or 10 mL/kg) to avoid first-pass effects and ensure stable pharmacokinetics; M100907 was injected SC (5 mL/kg); pimavanserin, edelfosine, *R*-(–)-DOI, (±)-DOI, 25C-NBOH, desoxy, CPM, 25C-NBMD, DET, 6-F-DET, 5-MeO-DMT, 6-MeO-DMT, and PCP were injected IP (5 mL/kg); YM-254,890 was injected into the lateral ventricle (2 µL) over 1 min. Slightly different procedures were used for the for the tolerance experiments (see below). HTR experiments with 1 mg/kg DOI were performed using either the racemate or the *R*-enantiomer as specified.

### Assessment of the Head-Twitch Response

The head-twitch response (HTR) was assessed using a head-mounted neodymium magnet and a magnetometer detection coil, as described previously ^89^. The mice were allowed to recover from the magnet implantation surgeries for at least 1 week prior to behavioral testing. Mice were tested in multiple HTR experiments, with at least 7 days between studies to avoid carryover effects. HTR experiments were conducted in a well-lit room, and the mice were allowed to habituate to the room for at least 1 h prior to testing. Mice were tested in a 12.5-cm diameter glass cylinder surrounded by a magnetometer coil. Coil voltage was low-pass filtered (2 kHz), amplified, and digitized (20-kHz sampling rate) using a Powerlab (model /8SP or 8/35) with LabChart software (ADInstruments, Colorado Springs, CO, USA). Head twitches were identified in the recordings off-line by their waveform characteristics ^90^ or using artificial intelligence ^91^. HTR counts were analyzed using one-way ANOVAs or one-way Welch ANOVAs (in cases where groups showed unequal variances). Tukey’s test or Dunnett’s T3 multiple comparisons test was used for *post hoc* comparisons. Significance was demonstrated by surpassing an α level of 0.05. Median effective doses (ED_50_ values) and 95% confidence intervals for dose-response experiments were calculated by nonlinear regression (Prism 9.02, GraphPad Software, San Diego, CA, USA).

### Assessment of PCP-Induced Locomotor Activity

The mouse behavioral pattern monitor (BPM) was used to assess locomotor activity ^92^. Each mouse BPM chamber (San Diego Instruments, San Diego, CA, USA) is a transparent Plexiglas box with an opaque 30 × 60 cm floor, enclosed in a ventilated isolation box. The position of the mouse in *x,y* coordinates is recorded by a grid of 12 × 24 infrared photobeams located 1 cm above the floor. A second row of 16 photobeams (parallel to the long axis of the chamber, located 2.5 cm above the floor) is used to detect rearing behavior. Holepoking behavior is detected by 11 1.4-cm holes that are situated in the walls (3 holes in each long wall, 2 holes in each short wall) and the floor (3 holes); each hole is equipped with an infrared photobeam. The status of each photobeam is sampled every 55 ms and recorded for offline analysis. Mice were allowed to habituate to the testing room for at least 1 h prior to testing. For the BPM experiments, mice (*n*=6/group) were pretreated SC with test drug or vehicle, PCP (5 mg/kg) or vehicle was injected IP 10 minutes later, and then the mice were placed in the BPM chambers 10 minutes after the second injection and activity was recorded for 60 minutes. Locomotor activity was quantified as distance traveled, which was analyzed in 20-minute blocks using a three-way ANOVA, with pretreatment and treatment as between subject variables and time as a within-subject variable. Tukey’s test was used for *post hoc* comparisons. Significance was demonstrated by surpassing an α level of 0.05.

### Effect of Repeated Treatment with 5-HT_2A_ receptor ligands on Head-Twitch Response

In the first experiment, mice (n=6-7/group, 19 total) were injected SC (5 mL/kg) once daily with vehicle (water containing 5% Tween-80), 25N-N1-Nap hydrochloride (20 mg/kg), or (±)-DOI hydrochloride (10 mg/kg) for five consecutive days. Twenty-four hours later, all of the mice were challenged with an IP injection of 1 mg/kg (±)-DOI hydrochloride and then HTR activity was recorded for 40 minutes. In the second experiment, mice (n=6/group, 12 total) were injected IP (5 mL/kg) once daily with vehicle (water) or pimavanserin hemitartrate (1 mg/kg) for five consecutive days. Twenty-four hours later, all of the mice were challenged with an IP injection of 1 mg/kg (±)-DOI hydrochloride and then HTR activity was recorded for 40 minutes.

## DATA AVAILABILITY

All data generated in this study are included in this article and the Supplementary information. Source data are provided with this paper.

## CODE AVAILABILITY

No custom code was generated in this study.

## ACKNOWLEDGEMENTS

This work was supported by Medical College of Wisconsin Research Affairs Counsel Pilot grant (JDM), National Institutes of Health General Medical Sciences grant (NIGMS R35GM133421 to JDM) and National Institutes of Health Drug Abuse (NIDA R01DA041336 to ALH), as well as by the Veteran’s Administration VISN 22 Mental Illness Research, Education, and Clinical Center. We thank technical assistance from Lisa McNally.

## AUTHOR CONTRIBUTIONS

The study was conceived by JW, ALH, and JDM. Compounds were synthesized and analytically characterized by JW, HM, JG, and TF. BRET, calcium flux, binding, mutagenesis, kinetics, and internalization experiments were performed by ABC, MMC, JKL, HAB, EMB, JJH, and EIA under the supervision of JDM. *In vivo* behavioral experiments were performed by ALH, RK, AKK, and BC. RZ and AJH conducted the docking and molecular dynamics simulations and helped write docking and MD sections. JW, ALH and JDM drafted and edited the original manuscript. All authors reviewed the manuscript.

