## Supplemental Material for "Identification of 5-HT_2A_ Receptor Signaling Pathways Responsible for Psychedelic Potential"

**Supplementary Figure 1. 5-HT$_{2A}$ receptor BRET Gq dissociation and β-arrestin2 recruitment kinetics.** Concentration response curves (*left*) and plots of relative activity, log ($E_{MAX}/EC_{50}$) across time (*right*) for 5-HT **(A)** and several classes of psychedelics: **(B)** Psilocin, **(C)** DMT, **(D)** 5-MeO-DMT, **(E)** 2C-I, **(F)** DOI, **(G)** 25I-NBOMe, and **(H)** LSD. Data represent the mean and SEM from three independent experiments, which were performed at 37°C at the indicated compound incubation time points.

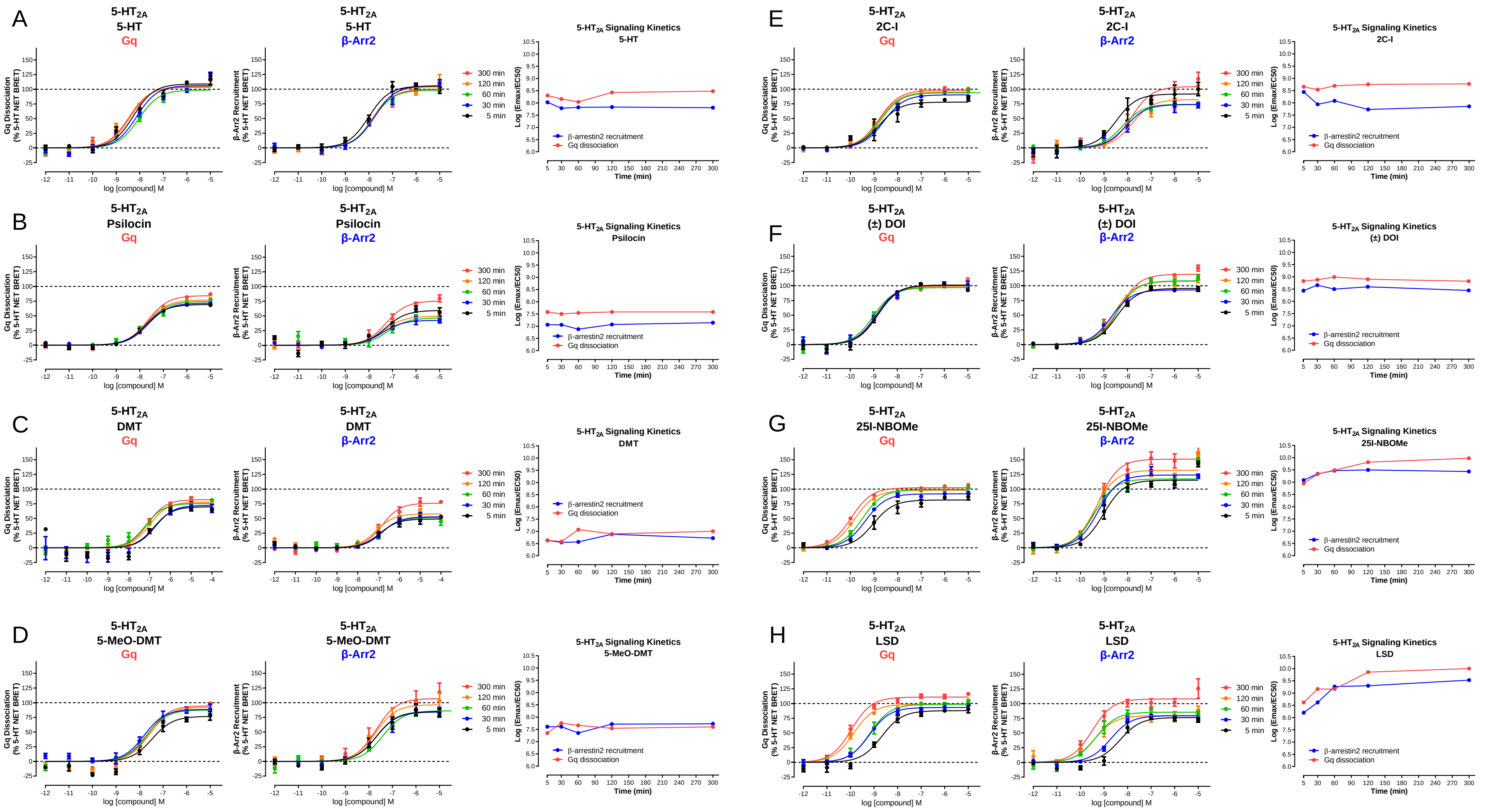

**Supplementary Figure 2. Structures of Target Compounds**

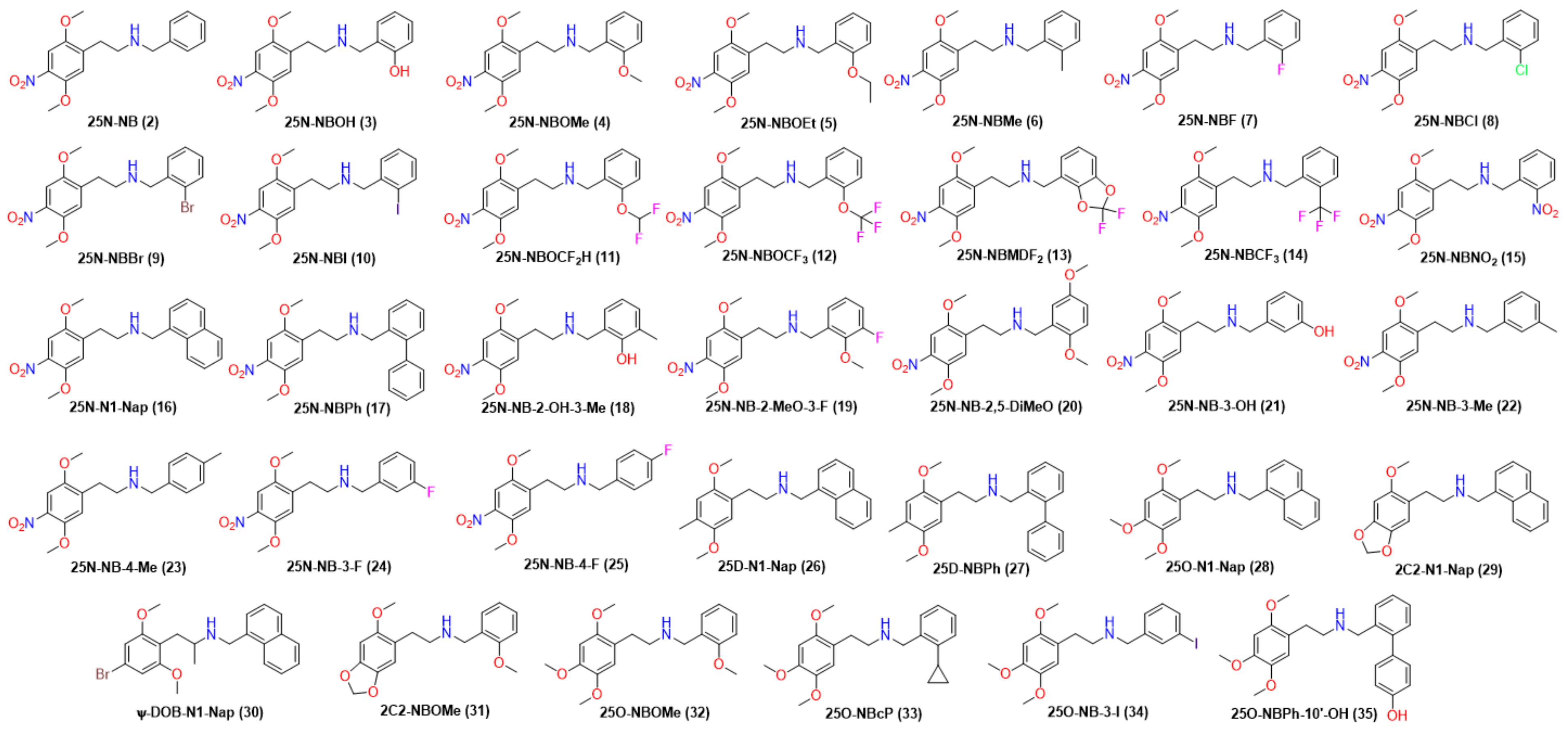

**Supplementary Figure 3. *N*-benzyl SAR electron density and electrostatics. (A)** Comparison of the *N*-benzyl electron density of 25N-NB-$NO_2$ (15) and 25N-NB-2-OH-3-Me (18) and their effect on 5-$HT_{2A}$ receptor affinity (expressed as pKi). (B) Model *N*-benzylamines used to calculate Hirshfeld charges. (C) Numbering scheme for *N*-benzylamine groups. (D) Electrostatic surface for 2-$NO_2$-benzylamine (15b). (E) Hirshfeld charges for 2-$NO_2$-benzylamine (15b). (F) Electrostatic surface for 2-HO-3-Me-benzylamine (18b). (G) Hirshfeld charges for 2-HO-3-Me-benzylamine (18b). (H) Electrostatic surface for 25N-NB$NO_2$ (15). (I) Electrostatic surface for 25N-NB-2-OH-3-Me (18). (J) Impact of C5' substituent steric Clash with optimal 5-$HT_{2A}$ receptor binding. (K) Correlation between $C_{5'}$ Hirshfeld charges and 5-$HT_{2A}$ $pK_i$ values. (L) Correlation between $H_{5'}$ $^1H$ NMR chemical shifts and 5-$HT_{2A}$ $pK_i$ values. (M) Correlation between $C_{5'}$ $^{13}C$ NMR chemical shifts and 5-$HT_{2A}$ $pK_i$ values. (N) Correlation between Hammett σ constant relative to $C_{5'}$ and $Ca^{2+}$ FLUX $pEC_{50}$ potency estimates. (O) Correlation between Hammett σ constant relative to $C_{5'}$ and BRET Gq $pEC_{50}$ potency estimates. (OP) Correlation between Hammett σ constant relative to $C_{5'}$ and BRET Arrestin $pEC_{50}$ potency estimates. Corresponding statistics in Supplementary Tables 5 and 6.

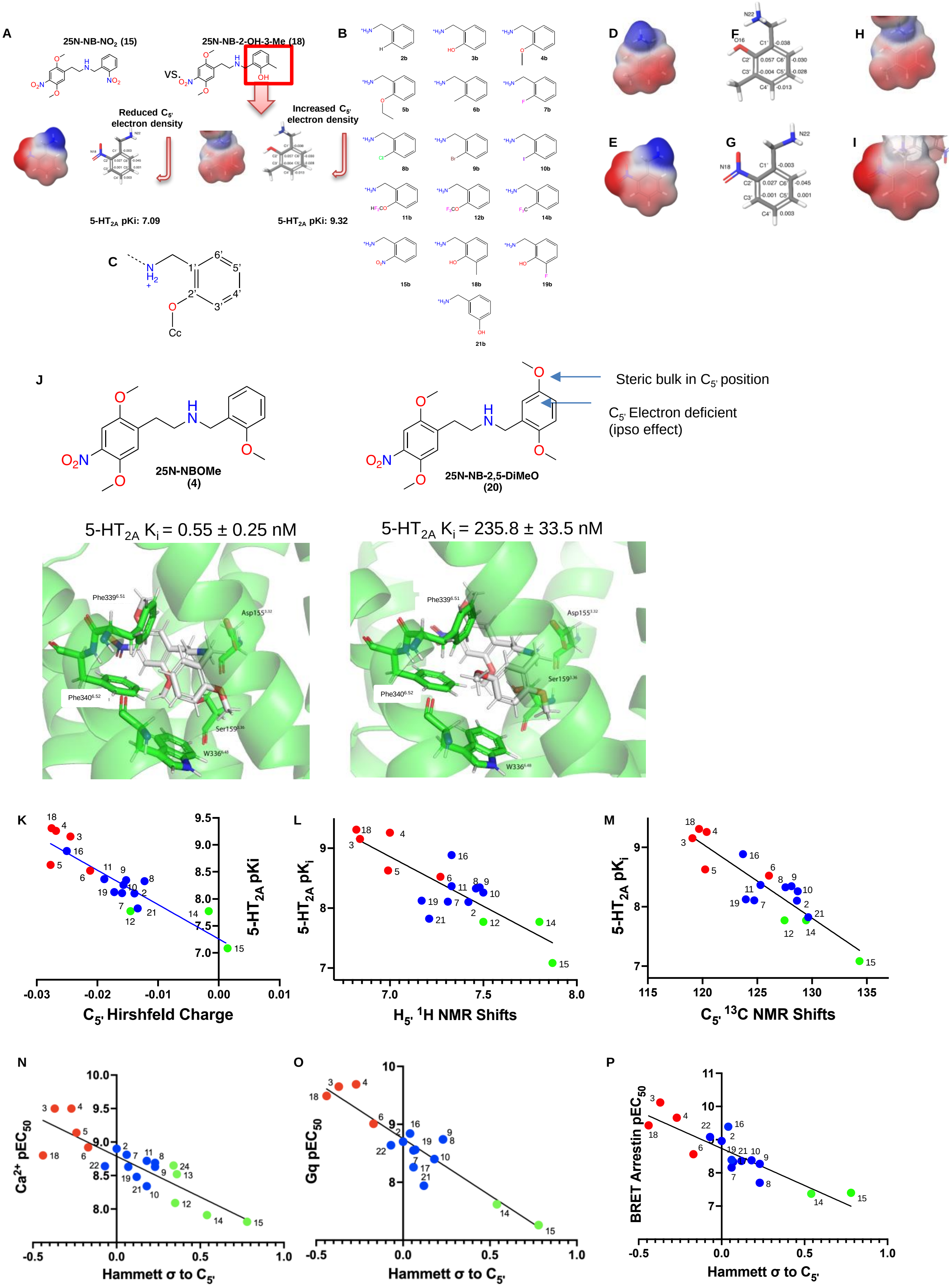

A
25N-NB-NO2 (15)
25N-NB-2-OH-3-Me (18)
VS.
Reduced C5' electron density
Increased C5' electron density
5-HT2A pKi: 7.09
5-HT2A pKi: 9.32
B
C
D
E
F
G
H
I
J
Steric bulk in C5' position
C5' Electron deficient (ipso effect)
25N-NBOMe (4)
25N-NB-2,5-DiMeO (20)
5-HT2A Ki = 0.55 ± 0.25 nM
5-HT2A Ki = 235.8 ± 33.5 nM
K
C5' Hirshfeld Charge
5-HT2A pKi
L
H5' 1H NMR Shifts
5-HT2A pKi
M
C5' 13C NMR Shifts
5-HT2A pKi
N
Hammett σ to C5'
Ca2+ pEC50
O
Hammett σ to C5'
Gq pEC50
P
Hammett σ to C5'
BRET Arrestin pEC50

**Supplementary Figure 4. Orthogonal assays and mutagenesis validation (A)** Comparison of 5-$HT_{2A}$ receptor and 5-$HT_{2C}$ receptor Gq dissociation for the 2-substituted halogen series indicating increasing substitution size reduces 5-$HT_{2C}$ receptor potency. Data represent the mean and SEM from three independent experiments performed at 37°C with 60 minute compound incubation. **(B)** Comparison of 5-$HT_{2A}$ receptor (green) 5-$HT_{2B}$ receptor (red) and 5-$HT_{2C}$ receptor (purple) Gq-mediated calcium flux responses for 25N-NBI (10) and 25CN-NBOH. Data represent the mean and SEM from three independent experiments. **(C)** Effect of 25N-NBI (10) on the HTR, measured as the number of responses recorded over a 30-min time period ($F_{5,24}$ = 14.48, $p$<0.0001). The estimated potency for 25I-NBI (10) is $ED_{50}$ = 10.9 µmol/kg. HTR counts from individual male C57BL/6J mice as well as group means are shown. ***$p$<0.001, ****$p$<0.0001, significant difference vs. vehicle control (Tukey's test). **(D)** 5-$HT_{2A}$ receptor Gq dissociation (red) and β-arrestin2 (blue) recruitment for the 5-$HT_{2A}$ receptor-selective agonist 25N-NBI (10). **(E)** Comparison of the Gq dissociation activity of 25N-NBI (10) at 5-$HT_{2A}$ receptor mutants as expressed as net BRET. Data represent the mean and SEM from three independent experiments performed at 37°C with 60 minute compound incubation.

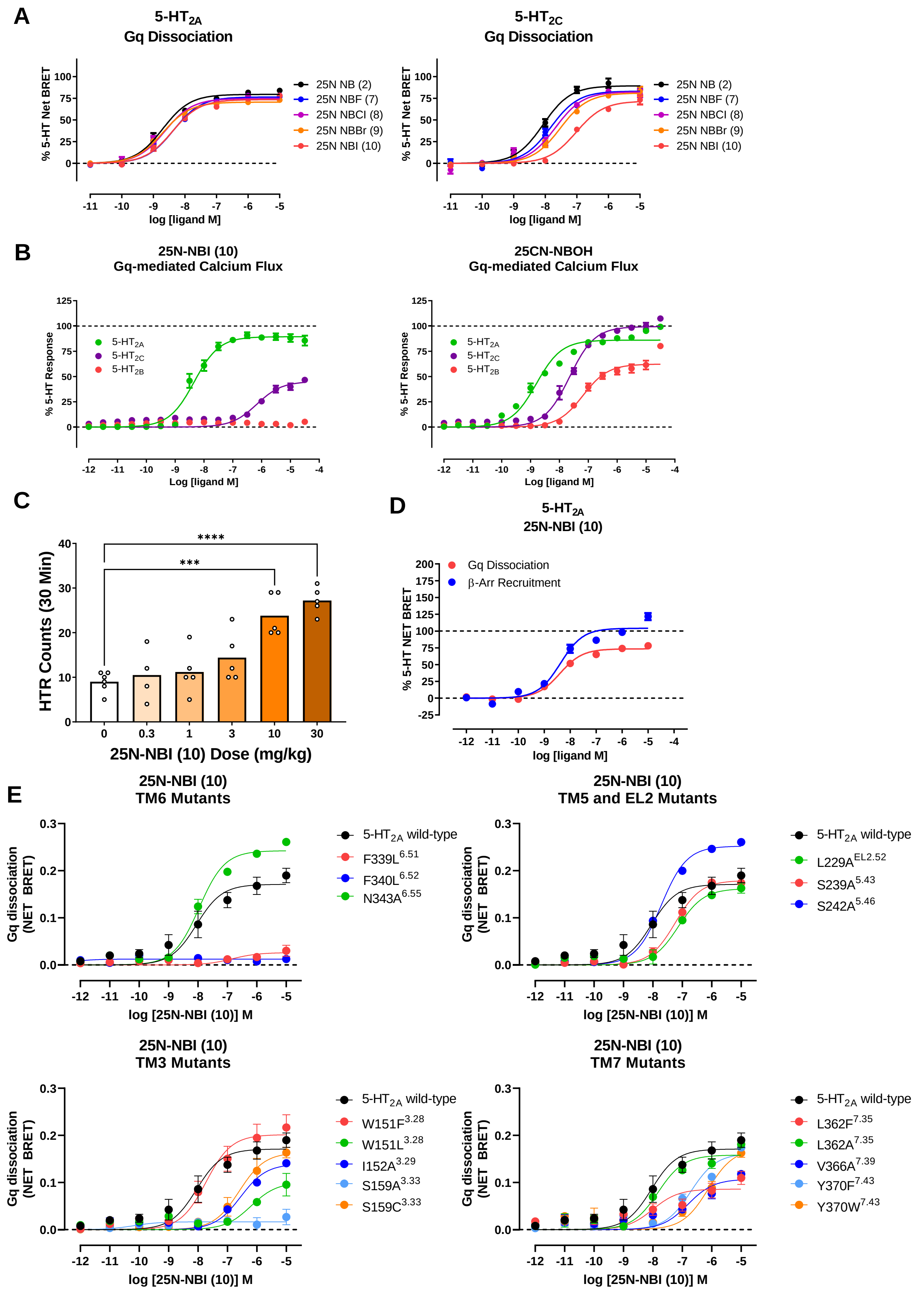
A
5-HT2A Gq Dissociation
5-HT2C Gq Dissociation
% 5-HT Net BRET
25N NB (2)
25N NBF (7)
25N NBCl (8)
25N NBBr (9)
25N NBI (10)
log [ligand M]
B
25N-NBI (10) Gq-mediated Calcium Flux
25CN-NBOH Gq-mediated Calcium Flux
% 5-HT Response
5-HT2A
5-HT2C
5-HT2B
Log [ligand M]
C
HTR Counts (30 Min)
****
***
25N-NBI (10) Dose (mg/kg)
D
5-HT2A 25N-NBI (10)
Gq Dissociation
β-Arr Recruitment
% 5-HT NET BRET
log [ligand M]
E
25N-NBI (10) TM6 Mutants
5-HT2A wild-type
F339L6.51
F340L6.52
N343A6.55
25N-NBI (10) TM5 and EL2 Mutants
L229AEL2.52
S239A5.43
S242A5.46
25N-NBI (10) TM3 Mutants
W151F3.28
W151L3.28
I152A3.29
S159A3.33
S159C3.33
25N-NBI (10) TM7 Mutants
L362F7.35
L362A7.35
V366A7.39
Y370F7.43
Y370W7.43
Gq dissociation (NET BRET)
log [25N-NBI (10)] M

**Supplementary Figure 5. Induced fit docking for 25CN-NBOH and target 25N compounds.** Induced fit docking at the human active state 5-$HT_{2A}$ receptor displays ligand binding mode and interactions with key orthosteric residues. (HB = H-bond, Ion = ionic interaction, PI = pi interaction). (A) Induced fit docking pose of 25CN-NBOH overlaid with the experimental binding mode for 25CN-NBOH (PDB: 6WHA). (B) Induced fit docking pose of 25N-NBOH (3). (C) Induced fit docking pose of 25N-NBOMe (4). (D) Induced fit docking pose of 25N-NBI (10). € Induced fit docking pose of 25N-N1-Nap (16). (F) Induced fit docking pose of 25N-NBPh (17). (G) Induced fit docking pose of 25N-NB-2-OH-3-Me (18). (H) Induced fit docking pose of 25N-NB-2,5-DiMeO (20).

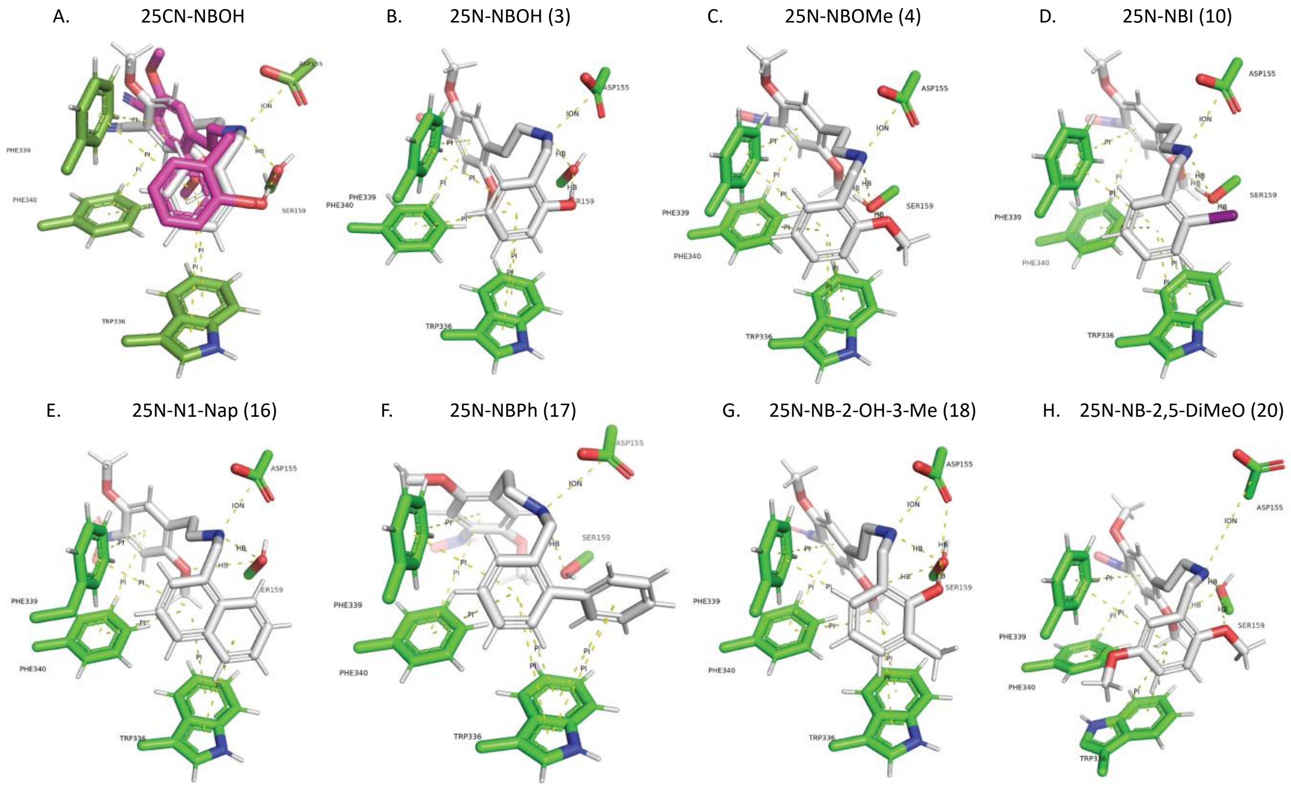

**Supplementary Figure 6. Extensive validation of β-arrestin bias SAR, 5-$HT_{2A}$R selectivity, and 5-$HT_{2A}$R mutagenesis (A)** Examination of 2-substituted 25N analogs measuring 5-$HT_{2A}$ receptor Gq dissociation (red) and β-arrestin2 (blue) recruitment indicating reduced Gq dissociation efficacy. Data represent the mean and SEM from 3 independent experiments performed at 37°C with 60 minute incubation. **(B)** 5-$HT_2$ selectivity comparing 5-$HT_{2A}$ receptor (green) to 5-$HT_{2B}$ receptor (red) and 5-$HT_{2C}$ receptor (purple) measuring Gq dissociation. Data represent the mean and SEM from 3 independent experiments performed at 37°C with 60 minute incubation. **(C)** Comparison of β-arrestin2 recruitment at 5-$HT_{2A}$ receptor (blue) to 5-$HT_{2B}$ receptor (orange) and 5-$HT_{2C}$ receptor (purple) for 25N-N1-Nap (16) and 25N-NBPh (17). Data represent the mean and SEM from 3 independent experiments performed at 37°C with 60 minute incubation. **(D)** Comparison of 5-$HT_{2A}$ activities measuring β-arrestin2 (blue), Gq (red) and G11 (pink) comparing 25N-NBOMe (4) to 25N-N1-Nap (16) and 25N-NBPh (17). Data represent the mean and SEM from 3 independent experiments performed at 37°C with 60 minute incubation. **(E)** 5-$HT_{2A}$-Gq/11-mediated calcium flux activity comparing 25N-NBOMe (4) to 25N-N1-Nap (16)and 25N-NBPh (17). Calcium flux traces comparing activity of 5-HT (purple) and vehicle (black) to 25N-N1-Nap (16) (red) and 25N-NBPh (17) (blue) at 1 µM concentration over 10 minutes. Calcium flux is expressed as relative fluorescence units RFU and represent mean and SEM from three independent experiments. **(F)** Comparison of activities across the 5-HT receptor GPCRome measuring G protein dissociation comparing 5-HT to to 25N-N1-Nap (16) (red) and 25N-NBPh (17). Data represent the mean and SEM from 3 independent experiments performed at 37°C with 60 minute incubation. **(G)** Comparison of Gq-partial agonist and assessment of partial antagonist activity measuring 5-$HT_{2A}$-Gq dissociation by BRET for 25N-N1-Nap (16, $IC_{50}$ = 15.6 nM) and 25N-NBPh (17, $IC_{50}$ = 68.6 nM). Data represent the mean and SEM from 3 independent experiments performed at 37°C with 60 minute incubation. **(H)** 5-$HT_{2A}$ receptor Gq dissociation (red) and β-arrestin2 (blue) recruitment kinetics comparing 25N-NBOMe (4) to 25N-N1-Nap (16) and 25N-NBPh (17). Data represent the mean and SEM from three independent experiments performed at 37°C at indicated time points. **(I)** Comparison of net BRET ratios relative to baseline of 5-$HT_{2A}$ receptor wild-type (green) to W336$^{6.48}$L (red) and W336$^{6.48}$Y (blue). Data represent the mean and SEM from three independent experiments performed at 37°C with 60 minute incubation.

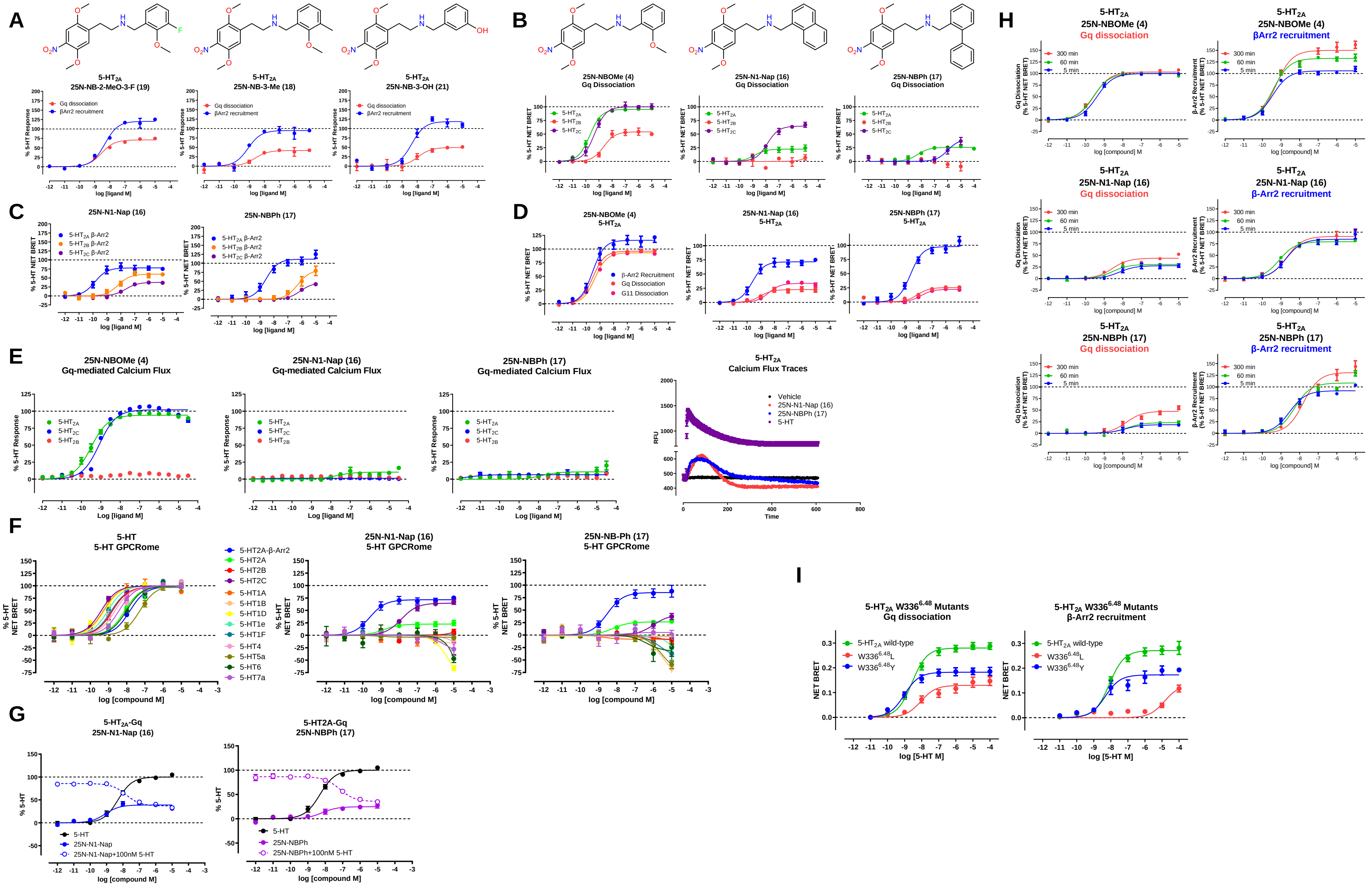
A
B
C
D
E
F
G
H
I
5-HT2A
25N-NB-2-MeO-3-F (19)
25N-NB-3-Me (18)
25N-NB-3-OH (21)
Gq dissociation
βArr2 recruitment
% 5-HT Response
log [ligand M]
25N-NBOMe (4)
Gq Dissociation
25N-N1-Nap (16)
25N-NBPh (17)
% 5-HT NET BRET
β-Arr2 Recruitment
G11 Dissociation
Gq-mediated Calcium Flux
Calcium Flux Traces
Vehicle
5-HT
RFU
Time
5-HT GPCRome
25N-NB-Ph (17)
5-HT2A-β-Arr2
5-HT2A
5-HT2B
5-HT2C
5-HT1A
5-HT1B
5-HT1D
5-HT1e
5-HT1F
5-HT4
5-HT5a
5-HT6
5-HT7a
log [compound M]
5-HT2A-Gq
25N-N1-Nap
25N-NBPh
25N-N1-Nap+100nM 5-HT
25N-NBPh+100nM 5-HT
% 5-HT
300 min
60 min
5 min
Gq Dissociation (% 5-HT NET BRET)
β-Arr2 Recruitment (% 5-HT NET BRET)
log [compound] M
5-HT2A W336 6.48 Mutants
β-Arr2 recruitment
5-HT2A wild-type
W336 6.48 L
W336 6.48 Y
NET BRET
log [5-HT M]

**Supplementary Figure 7. Time Series Heat Map for 25CN-NBOH MD Simulation.** Key interactions over the course of the simulation include D155$^{3.32}$, S159$^{3.36}$, F339$^{6.51}$, F340$^{6.52}$, and W336$^{6.48}$.

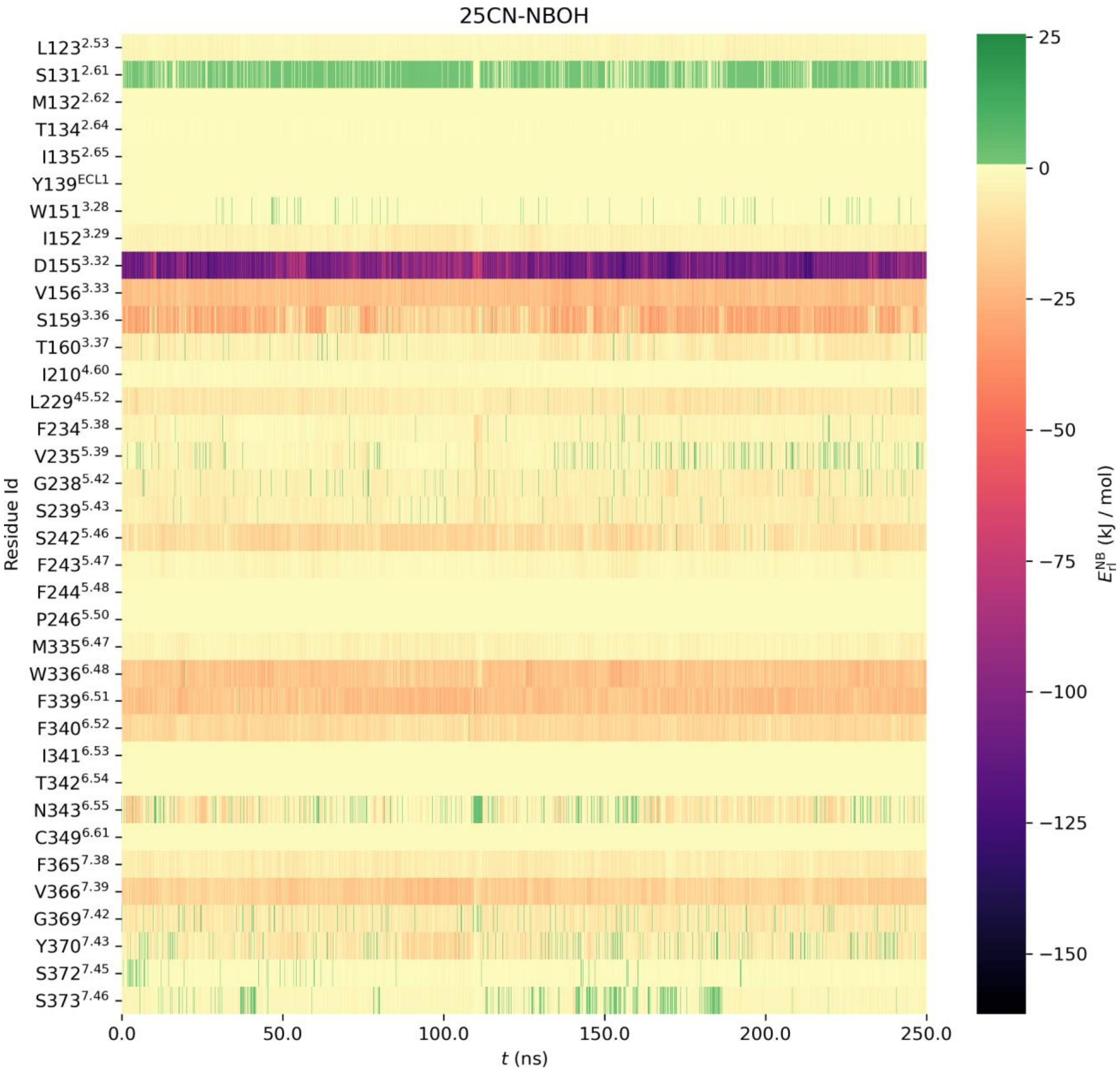

**Supplementary Figure 8. Time Series Heat Map for 25N-N1-Nap (16) MD Simulation.** Key interactions over the course of the simulation include D155$^{3.32}$, S159$^{3.36}$, F339$^{6.51}$, F340$^{6.52}$, and W336$^{6.48}$.

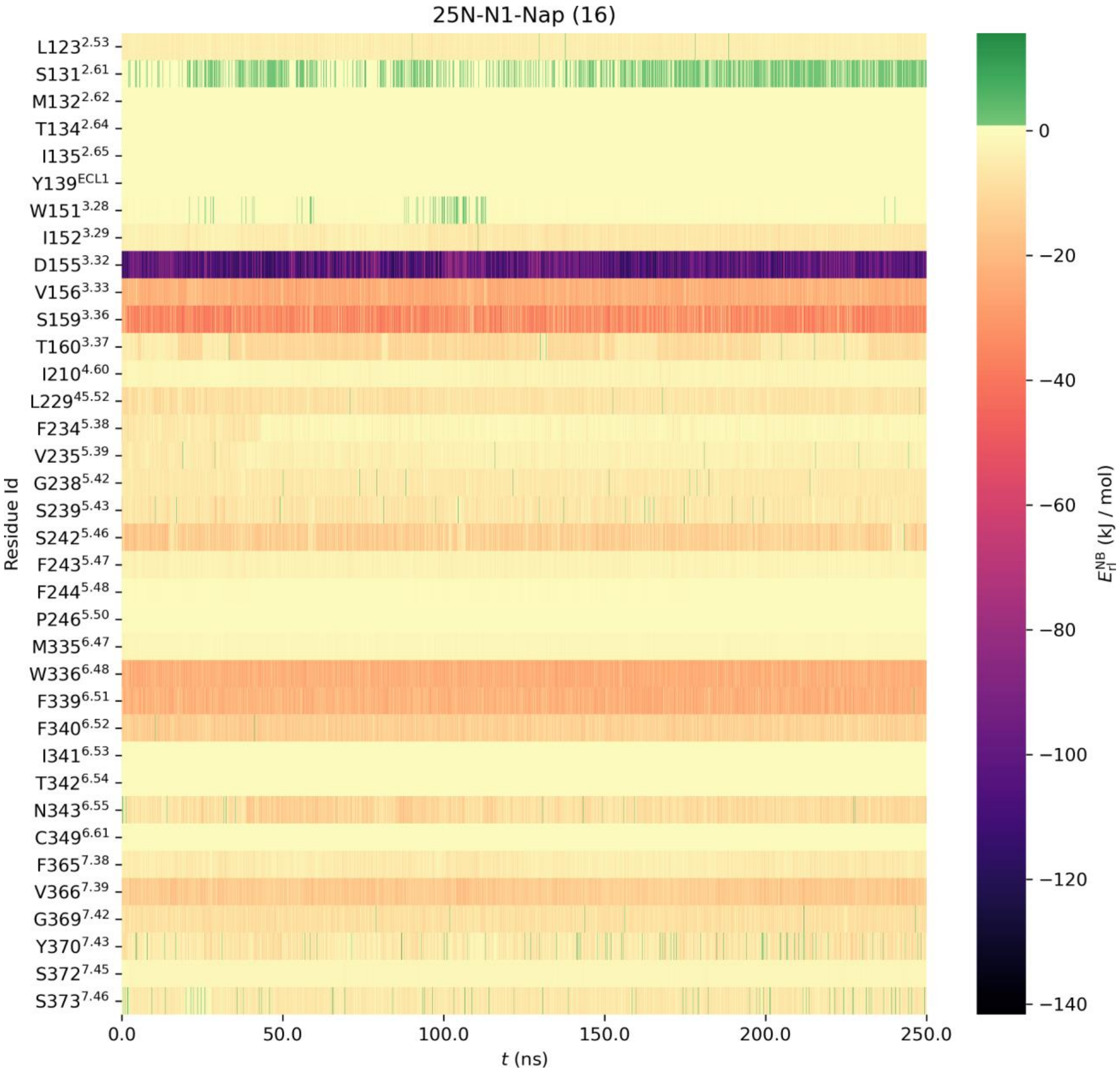

**Supplementary Figure 9. Mouse 5-$HT_{2A}$ activity and additional β-arrestin biased agonists.** **(A)** Mouse 5-$HT_{2A}$ receptor Gq dissociation (red) and β-arrestin2 (blue) recruitment comparing 25N-N1-Nap (16) (*left*) and 25N-NBPh (17) (*right*). Data represent mean and SEM from three independent experiments. **(B)** 25O-N1-Nap (26) (*left*) and 2C2-N1-Nap (29) (*right*) 5-$HT_{2A}$ receptor Gq dissociation (red) and β-arrestin2 (blue) activities. Data represent the mean and SEM from three independent experiments. **(C)** Effect of 25O-N1-Nap (28) (*left*) and 2C2-N1-Nap (29) (*right*) on the head-twitch response (HTR). **(D)** Pretreatment with 25O-N1-Nap (28) (*left*) and 2C2-N1-Nap (29) (*right*) blocks the HTR induced by 1 mg/kg (±)-DOI (25O-N1-Nap: $W_{4,11.42}$ = 80.05, $p$<0.0001; 2C2-N1-Nap: $F_{4,19}$ = 8.93, $p$=0.0003). **$p$<0.01, ****$p$<0.0001, significant difference between groups (Tukey's test or Dunnett's T3 test). HTR counts from individual male C57BL/6J mice as well as group means are shown.

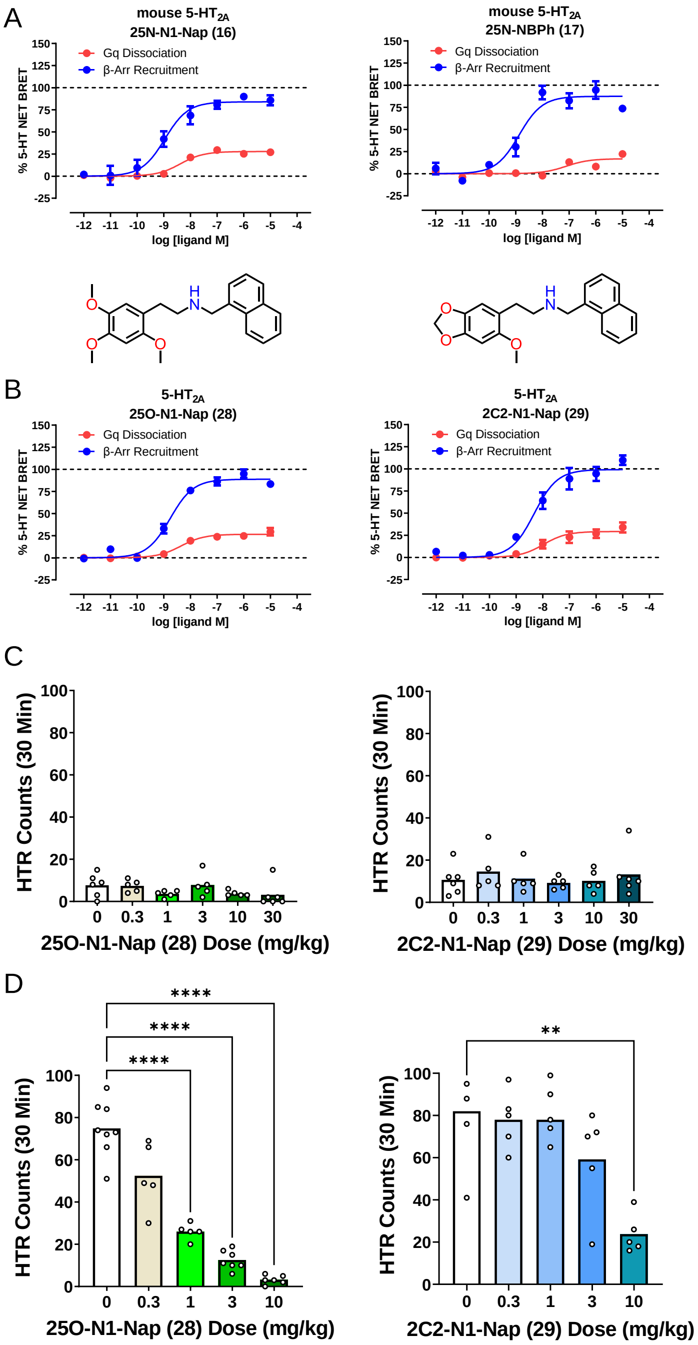

**Supplementary Figure 10. Correlation graphs and prediction ligand activities. (A)** Scatter plot showing the relationship between 5-$HT_{2A}$ receptor efficacy ($E_{MAX}$ values from calcium flux assays) and head-twitch response (HTR) magnitude (maximum counts per minute induced by the compound) for 15 members of the 25N series. Spearman's rank correlation coefficient $R_s$ is shown. The regression line generated by fitting the data using non-linear regression is included as a visual aid. **(B)** Scatter plot and linear regression showing the correlation between 5-$HT_{2A}$ receptor binding affinity ($K_i$ vs. [$^3$H]-ketanserin) and potencies in the HTR assay ($ED_{50}$) for eleven 25N derivatives. Pearson's correlation coefficient *R* is shown. **(C)** Scatter plot and linear regression showing the correlation between 5-$HT_{2A}$ receptor Gq activation potencies ($EC_{50}$) and potencies in the HTR assay ($ED_{50}$) for eight 25N derivatives. **(D)** Scatter plot and linear regression showing the correlation between 5-$HT_{2A}$ receptor Gq activation potencies ($EC_{50}$) and potencies in the HTR assay ($ED_{50}$) for 24 phenethylamine psychedelics. Pearson's correlation coefficient *R* is shown. **(E)** Scatter plot and linear regression showing the correlation between 5-$HT_{2A}$ receptor β-arrestin2 recruitment potencies ($EC_{50}$) and potencies in the HTR assay ($ED_{50}$) for eight 25N derivatives. **(F)** Scatter plot and linear regression showing the correlation between 5-$HT_{2A}$ receptor β-arrestin2 recruitment potencies ($EC_{50}$) and potencies in the HTR assay ($ED_{50}$) for 24 phenethylamine psychedelics. **(G-L)** 5-$HT_{2A}$ receptor Gq dissociation and β-arrestin2 recruitment activation profiles for the synthesized prediction compounds 2C2-NBOMe (31), 25O-NBOMe (32), 25O-NBcP (33), 25D-N1-Nap (26), 25O-NB-3-I (34), and 25O-NBPh-10'-OH (35). Data represent the mean and SEM from 3 independent experiments performed at 37°C with 60 minute incubation.

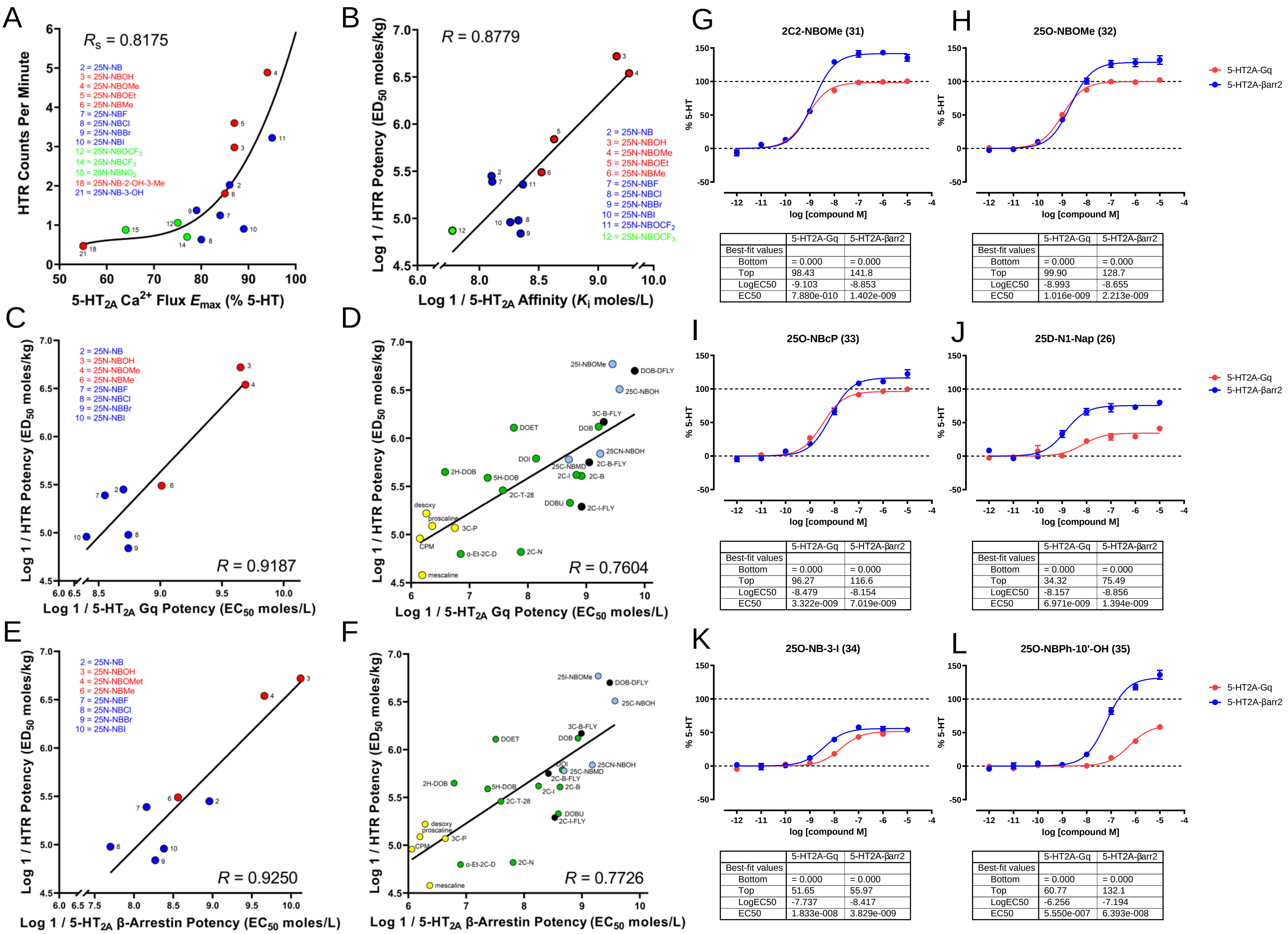

A
$R_S$ = 0.8175
2 = 25N-NB
3 = 25N-NBOH
4 = 25N-NBOMe
5 = 25N-NBOEt
6 = 25N-NBMe
7 = 25N-NBF
8 = 25N-NBCl
9 = 25N-NBBr
10 = 25N-NBI
12 = 25N-NBOCF$_3$
14 = 25N-NBCF$_3$
15 = 25N-NBNO$_2$
18 = 25N-NB-2-OH-3-Me
21 = 25N-NB-3-OH
HTR Counts Per Minute
5-HT$_{2A}$ Ca$^{2+}$ Flux $E_{max}$ (% 5-HT)
B
$R$ = 0.8779
2 = 25N-NB
3 = 25N-NBOH
4 = 25N-NBOMe
5 = 25N-NBOEt
6 = 25N-NBMe
7 = 25N-NBF
8 = 25N-NBCl
9 = 25N-NBBr
10 = 25N-NBI
11 = 25N-NBOCF$_2$
12 = 25N-NBOCF$_3$
Log 1 / HTR Potency (ED$_{50}$ moles/kg)
Log 1 / 5-HT$_{2A}$ Affinity ($K_i$ moles/L)
C
2 = 25N-NB
3 = 25N-NBOH
4 = 25N-NBOMe
6 = 25N-NBMe
7 = 25N-NBF
8 = 25N-NBCl
9 = 25N-NBBr
10 = 25N-NBI
$R$ = 0.9187
Log 1 / HTR Potency (ED$_{50}$ moles/kg)
Log 1 / 5-HT$_{2A}$ Gq Potency (EC$_{50}$ moles/L)
D
25I-NBOMe
DOB-DFLY
25C-NBOH
3C-B-FLY
DOB
DOET
25CN-NBOH
DOI
25C-NBMD
2C-B-FLY
2C-I
2C-B
2H-DOB
5H-DOB
2C-T-28
DOBU
2C-I-FLY
desoxy
proscaline
3C-P
CPM
α-Et-2C-D
2C-N
mescaline
$R$ = 0.7604
Log 1 / HTR Potency (ED$_{50}$ moles/kg)
Log 1 / 5-HT$_{2A}$ Gq Potency (EC$_{50}$ moles/L)
E
2 = 25N-NB
3 = 25N-NBOH
4 = 25N-NBOMet
6 = 25N-NBMe
7 = 25N-NBF
8 = 25N-NBCl
9 = 25N-NBBr
10 = 25N-NBI
$R$ = 0.9250
Log 1 / HTR Potency (ED$_{50}$ moles/kg)
Log 1 / 5-HT$_{2A}$ β-Arrestin Potency (EC$_{50}$ moles/L)
F
25I-NBOMe
DOB-DFLY
25C-NBOH
3C-B-FLY
DOB
DOET
DOI
25CN-NBOH
25C-NBMD
2C-B-FLY
2H-DOB
5H-DOB
2C-I
2C-B
2C-T-28
DOBU
2C-I-FLY
desoxy
proscaline
3C-P
CPM
α-Et-2C-D
2C-N
mescaline
$R$ = 0.7726
Log 1 / HTR Potency (ED$_{50}$ moles/kg)
Log 1 / 5-HT$_{2A}$ β-Arrestin Potency (EC$_{50}$ moles/L)
G
2C2-NBOMe (31)
% 5-HT
log [compound M]
| | 5-HT2A-Gq | 5-HT2A-βarr2 |
| Best-fit values | | |
| Bottom | = 0.000 | = 0.000 |
| Top | 98.43 | 141.8 |
| LogEC50 | -9.103 | -8.853 |
| EC50 | 7.880e-010 | 1.402e-009 |
H
25O-NBOMe (32)
% 5-HT
log [compound M]
| | 5-HT2A-Gq | 5-HT2A-βarr2 |
| Best-fit values | | |
| Bottom | = 0.000 | = 0.000 |
| Top | 99.90 | 128.7 |
| LogEC50 | -8.993 | -8.655 |
| EC50 | 1.016e-009 | 2.213e-009 |
5-HT2A-Gq
5-HT2A-βarr2
I
25O-NBcP (33)
% 5-HT
log [compound M]
| | 5-HT2A-Gq | 5-HT2A-βarr2 |
| Best-fit values | | |
| Bottom | = 0.000 | = 0.000 |
| Top | 96.27 | 116.6 |
| LogEC50 | -8.479 | -8.154 |
| EC50 | 3.322e-009 | 7.019e-009 |
J
25D-N1-Nap (26)
% 5-HT
log [compound M]
| | 5-HT2A-Gq | 5-HT2A-βarr2 |
| Best-fit values | | |
| Bottom | = 0.000 | = 0.000 |
| Top | 34.32 | 75.49 |
| LogEC50 | -8.157 | -8.856 |
| EC50 | 6.971e-009 | 1.394e-009 |
5-HT2A-Gq
5-HT2A-βarr2
K
25O-NB-3-I (34)
% 5-HT
log [compound M]
| | 5-HT2A-Gq | 5-HT2A-βarr2 |
| Best-fit values | | |
| Bottom | = 0.000 | = 0.000 |
| Top | 51.65 | 55.97 |
| LogEC50 | -7.737 | -8.417 |
| EC50 | 1.833e-008 | 3.829e-009 |
L
25O-NBPh-10'-OH (35)
% 5-HT
log [compound M]
| | 5-HT2A-Gq | 5-HT2A-βarr2 |
| Best-fit values | | |
| Bottom | = 0.000 | = 0.000 |
| Top | 60.77 | 132.1 |
| LogEC50 | -6.256 | -7.194 |
| EC50 | 5.550e-007 | 6.393e-008 |
5-HT2A-Gq
5-HT2A-βarr2

**Supplementary Figure 11. Structures of Examined Phenethylamine Psychedelics**

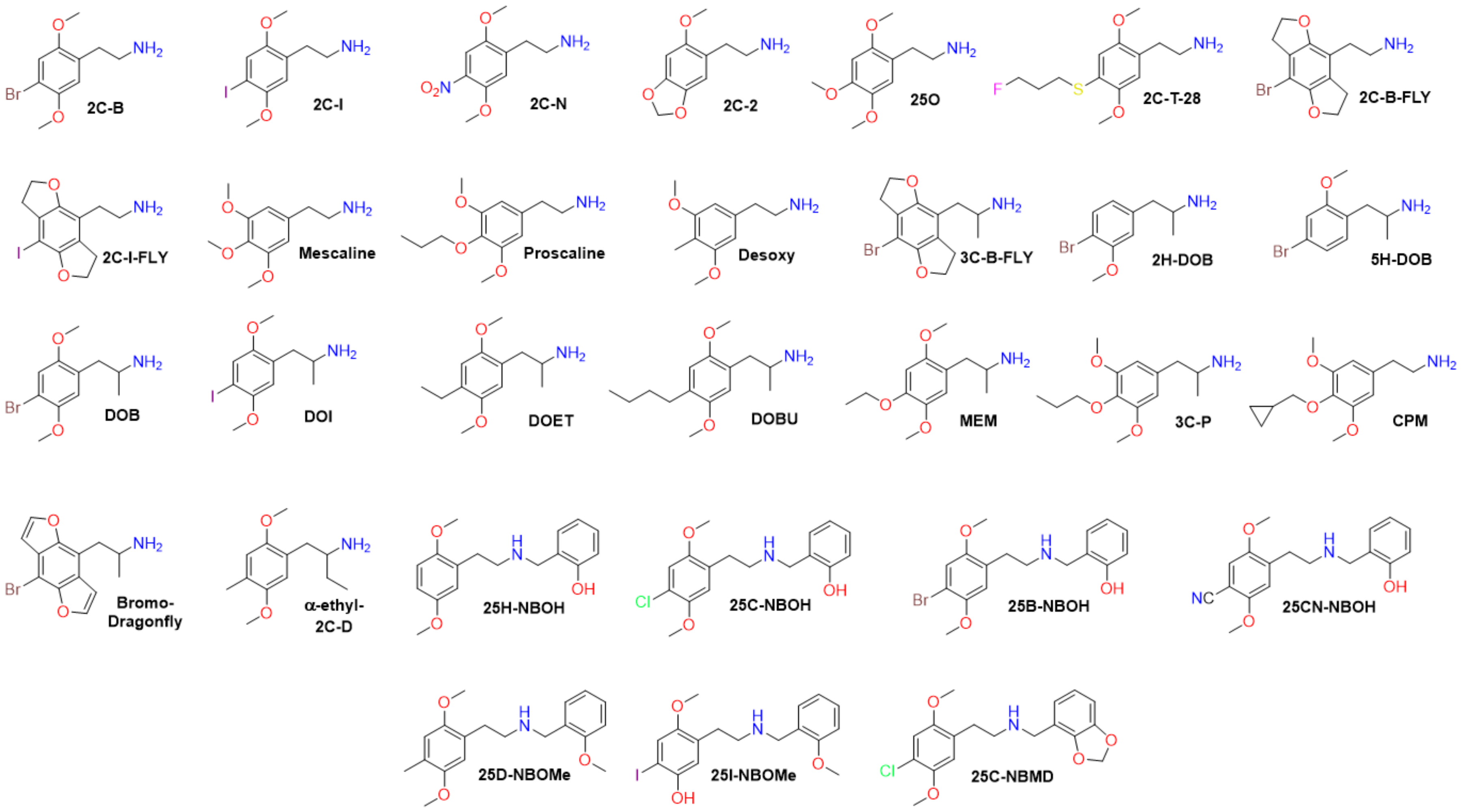

### Supplementary Table 1. 5-$HT_2$ Receptor Assay Parameters

Table of system, time, and temperature assay parameters for compounds tested in this study. Drug incubation times were varied in experiments in Supplementary Figure 1.

| **Parameter** | **G Protein Dissociation BRET** | **β-arrestin2 Recruitment BRET** |
|---|---|---|
| Transducer Pathway | Gq protein | β-Arrestin 2 |
| Cell Line | HEK 293T | HEK 293T |
| Data Collection Time Point (min) | 60 and various | 60 and various |
| Temperature (°C) | 37 | 37 |
| Reference Ligand for Emax | 5-HT | 5-HT |
| Reference Ligand for bias | 5-HT | 5-HT |
| Measured Process | Dissociation | Recruitment |
| Measured Molecule 1 | Gq-Rluc subunit | 5-$HT_2$-Rluc Receptor |
| Measured Molecule 2 | $GFP^2$-Gγ1 subunit | Venus-β-arrestin2 |
| Co-expressed Molecule | β1 | None |
| Signal Detection Technique | BRET | BRET |

**Supplementary Table 2. Naming scheme for the *N*-benzyl portion of target ligands.**

| Compound | *N*-Benzyl Ring Position | | | |
|---|---|---|---|---|
| | **2'-** | **3'-** | **4'-** | **5'-** |
| **25N-NB (2)** | H | H | H | H |
| **25N-NBOH (3)** | OH | H | H | H |
| **25N-NBOMe (4)** | $OCH_3$ | H | H | H |
| **25N-NBOEt (5)** | $OCH_2CH_3$ | H | H | H |
| **25N-NBMe (6)** | $CH_3$ | H | H | H |
| **25N-NBF (7)** | F | H | H | H |
| **25N-NBCl (8)** | Cl | H | H | H |
| **25N-NBBr (9)** | Br | H | H | H |
| **25N-NBI (10)** | I | H | H | H |
| **25N-NBOCF$_2$H (11)** | $OCF_2H$ | H | H | H |
| **25N-NBOCF$_3$ (12)** | $OCF_3$ | H | H | H |
| **25N-NBMDF$_2$ (13)** | $OCF_2O$ | | H | H |
| **25N-NBCF$_3$ (14)** | $CF_3$ | H | H | H |
| **25N-NBNO$_2$ (15)** | $NO_2$ | H | H | H |
| **25N-N1-Nap (16)** | $(CH)_4$ | | H | H |
| **25N-NBPh (17)** | Ph | H | H | H |
| **25N-NB-2-OH-3-Me (18)** | OH | $CH_3$ | H | H |
| **25N-NB-2-MeO-3-F (19)** | $OCH_3$ | F | H | H |

| | | | | |
|---|---|---|---|---|
| **25N-NB-2,5-DiMeO (20)** | $OCH_3$ | H | H | $OCH_3$ |
| **25N-NB-3-OH (21)** | H | OH | H | H |
| **25N-NB-3-Me (22)** | H | $CH_3$ | H | H |
| **25N-NB-4-Me (23)** | H | H | $CH_3$ | H |
| **25N-NB-3-F (24)** | H | F | H | H |
| **25N-NB-4-F (25)** | H | H | F | H |
| **25D-N1-Nap (26)** | $(CH_2)_4$ | H | H | H |
| **25D-NBPh (27)** | Ph | H | H | H |
| **25O-N1-Nap (28)** | $(CH)_4$ | | H | H |
| **2C2-N1-Nap (29)** | $(CH)_4$ | | H | H |
| **Ψ-DOB-N1-Nap (30)** | $(CH)_4$ | | H | H |
| **2C2-NBOMe (31)** | $OCH_3$ | H | H | H |
| **25O-NBOMe (32)** | $OCH_3$ | H | H | H |
| **25O-NBcP (33)** | $CH(CH_2)_2$ | H | H | H |
| **25O-NB-3-I (34)** | H | I | H | H |
| **25O-NBPh-10'-OH (35)** | 4-HO-Ph | H | H | H |

**Supplementary Table 3. Hammett σ Constants from the literature [Hansch et al. 1973, 1991].** Hansch, C., Leo, A. and Taft, R.W., 1991. A survey of Hammett substituent constants and resonance and field parameters. *Chemical Reviews*, *91*(2), pp.165-195. Hansch, C., Leo, A., Unger, S.H., Kim, K.H., Nikaitani, D. and Lien, E.J., 1973. Aromatic substituent constants for structure-activity correlations. *Journal of Medicinal Chemistry*, *16*(11), pp.1207-1216. Color code: Red (Hammett σ range = -0.50-0.00), blue (Hammett σ range = 0.01-0.30), green (Hammett σ range = 0.31-1.00).

| Compound | Functional Group | Hammett σ constant to $C_{5'}$ |
|---|---|---|
| 25N-NB-2-OH-3-Me (18) | 2-OH-3-Me- | -0.44* |
| 25N-NBOH (3) | 2-OH | -0.37 |
| 25N-NBOMe (4) | 2-OMe | -0.27 |
| 25N-NBOEt (5) | 2-OEt | -0.24 |
| 25N-NBMe (6) | 2-Me | -0.17 |
| 25N-NBPh (17) | 2-Ph | -0.01 |
| 25N-NB-3-Me (22) | 3-Me | -0.07 |
| 25N-NB (2) | 2-H | 0.00 |
| 25N-NBF (7) | 2-F | 0.06 |
| 25N-NBI (10) | 2-I | 0.18 |
| 25N-NBOCF$_2$H (11) | 2-OCF$_2$H | 0.18 |
| 25N-NBCl (8) | 2-Cl | 0.23 |
| 25N-NBBr (9) | 2-Br | 0.23 |
| 25N-N1-Nap (16) | 2,3-(C$_4$H$_4$) | 0.04 |
| 25NB-2-MeO-3-F (19) | 2-MeO-3-F | 0.07 |
| 25N-NB-3-OH (21) | 3-OH | 0.12 |
| 25N-NB-3-F (24) | 3-F | 0.34 |
| 25N-NBOCF$_3$ (12) | 2-OCF$_3$ | 0.35 |
| 25N-NBMDF$_2$ (13) | 2,3-MDF$_2$ | 0.36 |
| 25N-NBCF$_3$ (14) | 2-CF$_3$ | 0.54 |
| 25N-NBNO$_2$ (15) | 2-NO$_2$ | 0.78 |

*Additive value from corresponding $\sigma_m$ and $\sigma_p$ values. Color code: Red = strong electron donating (Hammett σ range: -0.5 to -0.10, Blue = Neutral electronic range = -0.09 to 0.30, Green = strong electron withdrawing = Hammett σ range: 0.31 to 1.00.

**Supplementary Table 4. Calculated Hirshfeld Charges for protonated substituted *N*-benzylamine groups.** Order based on highest to lowest negative charge around $C_{5'}$ carbon. Color code: Red (Hammett σ range = -0.50-0.00), blue (Hammett σ range = 0.01-0.30), green (Hammett σ range = 0.31-1.00). See F.L. Hirshfeld, *Theoretica chimica acta* 44, 129-138 (1977).

| Compound | $C_{1'}$ | $C_{2'}$ | $C_{3'}$ | $C_{4'}$ | $C_{5'}$ | $C_{6'}$ |
|---|---|---|---|---|---|---|
| **NB (2b)** | -0.02595 | -0.03538 | -0.01392 | -0.00226 | -0.0139 | -0.03547 |
| **NBOH (3b)** | -0.03682 | 0.06355 | -0.04431 | -0.00235 | -0.02444 | -0.02295 |
| **NBOMe (4b)** | -0.03767 | 0.06272 | -0.04656 | -0.0057 | -0.02684 | -0.02475 |
| **NBOEt (5b)** | -0.03876 | 0.06212 | -0.04683 | -0.00624 | -0.02772 | -0.02595 |
| **NBMe (6b)** | -0.03172 | 0.00711 | -0.02309 | -0.0044 | -0.0212 | -0.03588 |
| **NBF (7b)** | -0.03535 | 0.0822 | -0.02632 | 0.00429 | -0.01595 | -0.02042 |
| **NBCl (8b)** | -0.02719 | 0.02398 | -0.01848 | 0.00372 | -0.01224 | -0.02245 |
| **NBBr (9b)** | -0.0343 | -0.01277 | -0.03002 | -0.00157 | -0.01527 | -0.02763 |
| **NBI (10b)** | -0.03145 | -0.03839 | -0.02894 | -0.00369 | -0.01572 | -0.02808 |
| **NBOCF$_2$H (11b)** | -0.03293 | 0.06314 | -0.04114 | 0.001 | -0.01887 | -0.02022 |
| **NBOCF$_3$ (12b)** | -0.03063 | 0.05923 | -0.03605 | 0.0037 | -0.01456 | -0.02023 |
| **NBCF$_3$ (14b)** | -0.0158 | -0.03269 | -0.01037 | 0.00398 | -0.00167 | -0.02342 |
| **NBNO$_2$ (15b)** | -0.00343 | 0.02686 | -0.00146 | 0.00275 | 0.00144 | -0.04535 |
| **N1-Nap (16b)** | -0.02464 | -0.00428 | 0.01353 | -0.00566 | -0.02505 | -0.02773 |
| **NB-2-OH-3-Me (18b)** | -0.03864 | 0.05792 | -0.00397 | -0.01281 | -0.02755 | -0.03011 |
| **NB-2-OMe-3-F (19b)** | -0.03263 | 0.049 | 0.08264 | -0.01998 | -0.01724 | -0.03131 |
| **NB-3-OH (21b)** | -0.02277 | -0.05655 | 0.08567 | -0.03577 | -0.01337 | -0.05222 |

**Supplementary Table 5. Pearson Rank Correlation Statistics for Electrostatic Structure Activity Relationship Studies.**

| Correlation | N | Pearson's r | P value (Two-Tailed) |
|---|---|---|---|
| Hammett σ to $C_{5'}$ vs. 5-$HT_{2A}$ $pK_i$ | 17 | -0.8980 | <0.0001 |
| Hirshfeld partial charge vs. 5-$HT_{2A}$ $pK_i$ | 17 | -0.8980 | <0.0001 |
| $H_{5'}$ $^{1}H$ NMR chemical shift vs. 5-$HT_{2A}$ $pK_i$ | 17 | -0.8266 | <0.0001 |
| $C_{5'}$ $^{13}C$ NMR chemical shift vs. 5-$HT_{2A}$ $pK_i$ | 17 | -0.8920 | <0.0001 |
| Hammett σ to $C_{5'}$ vs. 5-$HT_{2A}$ $Ca^{2+}$ Flux $pEC_{50}$ | 19 | -0.8558 | <0.0001 |
| Hammett σ to $C_{5'}$ vs. 5-$HT_{2A}$ Gq BRET | 16 | -0.9072 | <0.0001 |
| Hammett σ to $C_{5'}$ vs. 5-$HT_{2A}$ Arrestin BRET | 16 | -0.8736 | <0.0001 |

**Supplementary Table 6. Linear Regression Correlation Statistics for Electrostatic Structure Activity Relationship Studies.**

| Correlation | N | $R^2$ | F Statistic | P value (Two-Tailed) | Line Equation |
|---|---|---|---|---|---|
| Hammett σ to $C_{5'}$ vs. 5-$HT_{2A}$ $pK_i$ | 17 | 0.8064 | $F_{(1,15)} = 62.49$ | <0.0001 | Y = -63.70X + 7.257 |
| Hirshfeld partial charge vs. 5-$HT_{2A}$ $pK_i$ | 17 | 0.8064 | $F_{(1,15)} = 62.49$ | <0.0001 | Y = -63.70x + 7.257 |
| $H_5$' $^1$H NMR chemical shift vs. 5-$HT_{2A}$ $pK_i$ | 17 | 0.6832 | $F_{(1,15)} = 32.35$ | <0.0001 | Y = -1.65X + 20.42 |
| $C_5$' $^{13}$C NMR chemical shift vs. 5-$HT_{2A}$ $pK_i$ | 17 | 0.7957 | $F_{(1,15)} = 58.42$ | <0.0001 | Y = -0.1250X + 24.06 |
| Hammett σ to $C_{5'}$ vs. $Ca^{2+}$ Flux $pEC_{50}$ | 19 | 0.7323 | $F_{(1,17)} = 46.51$ | <0.0001 | Y = -1.217X + 8.788 |
| Hammett σ to $C_{5'}$ vs. Gq BRET | 16 | 0.8230 | $F_{(1,14)} = 65.07$ | <0.0001 | Y = -1.955X + 8.752 |
| Hammett σ to $C_{5'}$ vs. Arrestin BRET | 16 | 0.7631 | $F_{(1,14)} = 45.10$ | <0.0001 | Y = -2.230X + 8.735 |

**Supplementary Table 7. 5-$HT_2$ Gq-mediated $Ca^{2+}$ Flux Data.** Data presented as mean ± SEM from three biological replicates. NA = no activity; NC = not calculated.

| | **5-$HT_{2A}$** | | **5-$HT_{2B}$** | | **5-$HT_{2C}$** | |
|---|---|---|---|---|---|---|
| | **$EC_{50}$, nM ($pEC_{50}$ ± SEM)** | **Emax % 5-HT** | **$EC_{50}$, nM ($pEC_{50}$ ± SEM)** | **Emax % 5-HT** | **$EC_{50}$, nM ($pEC_{50}$ ± SEM)** | **Emax % 5-HT** |
| **5-HT** | 0.42 (9.37 ± 0.03) | 100 | 1.10 (8.96 ± 0.03) | 100 | 0.22 (9.66 ± 0.02) | 100 |
| **25N (1)** | 4.56 (8.31 ± 0.03) | 100 ± 1 | 50.3 (7.30 ± 0.09) | 72 ± 3 | 60.1 (7.22 ± 0.07) | 69 ± 2 |
| **25N-NB (2)** | 1.27 (8.90 ± 0.04) | 86 ± 1 | NA | <10 | 26.2 (7.58 ± 0.05) | 75 ± 1 |
| **25N-NBOH (3)** | 0.32 (9.50 ± 0.06) | 87 ± 2 | NA | <10 | 1.78 (8.75 ± 0.04) | 100 ± 2 |
| **25N-NBOMe (4)** | 0.32 (9.50 ± 0.03) | 94 ± 1 | NA | <10 | 0.84 (9.07 ± 0.03) | 102 ± 1 |
| **25N-NBOEt (5)** | 0.72 (9.14 ± 0.08) | 87 ± 2 | NA | <10 | 0.88 (9.06 ± 0.04) | 99 ± 1 |
| **25N-NBMe (6)** | 1.19 (8.92 ± 0.04) | 85 ± 1 | NA | <10 | 33.2 (7.48 ± 0.03) | 89 ± 2 |
| **25N-NBF (7)** | 1.56 (8.81 ± 0.04) | 84 ± 1 | NA | <10 | 28.8 (7.54 ± 0.05) | 82 ± 2 |
| **25N-NBCl (8)** | 2.03 (8.69 ± 0.07) | 80 ± 2 | NA | <10 | 89.3 (7.05 ± 0.05) | 81 ± 2 |
| **25N-NBBr (9)** | 2.36 (8.63 ± 0.08) | 79 ± 2 | NA | <10 | 144 (6.84 ± 0.06) | 78 ± 2 |
| **25N-NBI (10)** | 4.57 (8.34 ± 0.05) | 89 ± 1 | NA | <10 | 682 (6.17 ± 0.08) | 45 ± 2 |
| **25N-NBO$CF_2$H (11)** | 1.92 (8.72 ± 0.04) | 95 ± 1 | NA | <10 | 3.99 (8.40 ± 0.03) | 101 ± 1 |
| **25N-NBO$CF_3$ (12)** | 8.09 (8.09 ± 0.09) | 75 ± 2 | NA | <10 | 58.3 (7.23 ± 0.04) | 89 ± 1 |
| **25N-NBMD$F_2$ (13)** | 3.02 (8.52 ± 0.05) | 86 ± 2 | NA | <10 | 431 (6.36 ± 0.20) | 49 ± 4 |
| **25N-NB$CF_3$ (14)** | 12.3 (7.91 ± 0.04) | 77 ± 1 | NA | <10 | 271 (6.57 ± 0.22) | 20 ± 2 |
| **25N-NB$NO_2$ (15)** | 15.6 (7.81 ± 0.07) | 64 ± 2 | NA | <10 | 583 (6.23 ± 0.08) | 36 ± 2 |
| **25N-N1-Nap (16)** | NA | <10 | NA | <10 | NA | <10 |
| **25N-NBPh (17)** | NA | <10 | NA | <10 | NA | <10 |
| **25N-NB-2-OH-3-Me (18)** | 1.58 (8.80 ± 0.13) | 55 ± 2 | NA | <10 | NA | <10 |
| **25N-NB-2-MeO-3-F (19)** | 2.35 (8.63 ± 0.05) | 83 ± 1 | NA | <10 | 74.5 (7.13 ± 0.08) | 77 ± 2 |
| **25N-NB-2,5-DiMeO (20)** | 133 (6.88 ± 0.04) | 72 ± 1 | NA | <10 | NA | <10 |

| | | | | | | |
|---|---|---|---|---|---|---|
| **25N-NB-3-OH (21)** | 3.32<br>(8.48 ± 0.09) | 55 ± 2 | NA | <10 | 49.1<br>(7.31 ± 0.19) | 53 ± 4 |
| **25N-NB-3-Me (22)** | 2.27<br>(8.64 ± 0.09) | 54 ± 1 | NA | <10 | 14.3<br>(7.84 ± 0.30) | 15 ± 2 |
| **25N-NB-4-Me (23)** | 38.3<br>(7.42 ± 0.17) | 42 ± 3 | NA | <10 | 844<br>(6.07 ± 0.15) | 14 ± 1 |
| **25N-NB-3-F (24)** | 2.25<br>(8.65 ± 0.06) | 85 ± 2 | NA | <10 | 149<br>(6.83 ± 0.04) | 54 ± 1 |
| **25N-NB-4-F (25)** | 19.7<br>(7.71 ± 0.14) | 48 ± 2 | NA | <10 | 413<br>(6.38 ± 0.15) | 21 ± 2 |

**Supplementary Table 8. Affinity Constants for Select Compounds at 5-HT Receptors.** Compounds were assayed as described by NIMH PDSP. Data presented as mean ± SEM (N = 3-6 separate experiments) except where single value without SEM then N = 1. <5 indicates compounds were inactive in a primary screening at 10 µM. ND = not determined.

| **Compound** | **5-HT Receptors ($pK_i$ ± SEM)** | | | | | | | | | | |
|---|---|---|---|---|---|---|---|---|---|---|---|
| | $5\text{-HT}_{1A}$ | $5\text{-HT}_{1B}$ | $5\text{-HT}_{1D}$ | $5\text{-HT}_{1e}$ | $5\text{-HT}_{2A}$ | $5\text{-HT}_{2B}$ | $5\text{-HT}_{2C}$ | $5\text{-HT}_{3}$ | $5\text{-HT}_{5a}$ | $5\text{-HT}_{6}$ | $5\text{-HT}_{7}$ |
| **2C-N (1)** | 5.84 ± 0.18 | <5 | 6.08 ± 0.09 | 6.17 ± 0.13 | 7.14 ± 0.02 | 6.91 ± 0.03 | 6.79 ± 0.18 | <5 | <5 | 6.60 ± 0.18 | <5 |
| **25N-NB (2)** | 5.72 ± 0.09 | <5 | <5 | <5 | 8.13 ± 0.10 | 7.84 ± 0.08 | 7.49 ± 0.04 | <5 | <5 | 6.50 ± 0.06 | <5 |
| **25N-NBOH (3)** | 5.80 ± 0.06 | <5 | <5 | <5 | 9.18 ± 0.06 | 8.31 ± 0.11 | 8.10 ± 0.03 | 5.38 ± 0.23 | 5.73 ± 0.05 | 6.72 ± 0.17 | <5 |
| **25N-NBOMe (4)** | 5.73 ± 0.15 | <5 | 5.25 ± 0.06 | <5 | 9.26 ± 0.15 | 8.35 ± 0.08 | 8.16 ± 0.07 | <5 | <5 | 7.26 ± 0.13 | <5 |
| **25N-NBOEt (5)** | 6.23 ± 0.11 | <5 | <5 | <5 | 8.65 ± 0.07 | 8.82 ± 0.22 | 8.39 ± 0.07 | <5 | <5 | 7.22 ± 0.05 | <5 |
| **25N-NBMe (6)** | 5.80 ± 0.15 | 5.53 ± 0.09 | <5 | <5 | 8.56 ± 0.09 | 7.34 ± 0.11 | 7.29 ± 0.12 | <5 | <5 | 6.48 ± 0.07 | <5 |
| **25N-NBF (7)** | 5.66 ± 0.08 | <5 | <5 | <5 | 8.11 ± 0.06 | 7.47 ± 0.06 | 7.28 ± 0.01 | 5.41 ± 0.03 | <5 | 6.29 ± 0.06 | <5 |
| **25N-NBCl (8)** | 5.83 ± 0.05 | <5 | <5 | <5 | 8.37 ± 0.12 | 7.33 ± 0.16 | 7.26 ± 0.11* | <5 | 5.42 ± 0.01 | 6.34 ± 0.11 | <5 |
| **25N-NBBr (9)** | 5.86 ± 0.06 | <5 | <5 | <5 | 8.35 ± 0.06 | 7.32 ± 0.17 | 7.38 ± 0.03 | <5 | <5 | 6.43 ± 0.14 | <5 |
| **25N-NBI (10)** | 6.00 ± 0.06 | <5 | <5 | <5 | 8.27 ± 0.08 | 7.70 ± 0.17 | 7.36 ± 0.02 | <5 | <5 | 6.42 ± 0.04 | <5 |
| **25N-NBOCF$_2$H (11)** | 5.93 ± 0.09 | <5 | <5 | <5 | 8.45 ± 0.15 | 8.49 ± 0.13 | 8.11 ± 0.11 | <5 | <5 | 6.59 ± 0.05 | <5 |
| **25N-NBOCF$_3$ (12)** | 5.95 ± 0.08 | <5 | <5 | <5 | 7.84 ± 0.14 | 7.86 ± 0.12 | 7.33 ± 0.08 | <5 | 5.80 ± 0.10 | 6.12 ± 0.09 | <5 |
| **25N-NBMDF$_2$ (13)** | 6.00 | <5 | <5 | <5 | 8.92 | 7.33 | 6.67 | <5 | ND | ND | ND |

| | | | | | | | | | | | |
|---|---|---|---|---|---|---|---|---|---|---|---|
| **25N-NB$CF_3$ (14)** | 5.97 ± 0.05 | <5 | <5 | <5 | 7.70 ± 0.05 | 7.23 ± 0.22 | 6.99 ± 0.08 | <5 | <5 | 6.39 ± 0.02 | <5 |
| **25N-NB$NO_2$ (15)** | 5.71 ± 0.15 | <5 | <5 | <5 | 7.09 ± 0.04 | 6.72 ± 0.05 | 6.55 ± 0.06 | <5 | <5 | <5 | <5 |
| **25N-N1-Nap (16)** | 6.62 ± 0.02 | <5 | 6.06 ± 0.06 | <5 | 8.94 ± 0.14 | 8.93 ± 0.27 | 8.37 ± 0.06 | <5 | 5.96 ± 0.16 | 7.10 ± 0.11 | 6.73 ± 0.10 |
| **25N-NBPh (17)** | 5.75 | <5 | 5.92 | <5 | 9.48 | 5.85 | 6.43 | <5 | ND | ND | ND |
| **25N-NB-2-OH-3-Me (18)** | 6.73 ± 0.10 | <5 | 5.89 ± 0.08 | <5 | 9.32 ± 0.09 | 9.11 ± 0.14 | 8.54 ± 0.02 | <5 | 6.30 ± 0.08 | 6.68 ± 0.02 | 6.06 ± 0.04 |
| **25N-NB-2-MeO-3-F (19)** | 6.00 ± 0.12 | <5 | <5 | <5 | 8.13 ± 0.05 | 7.60 ± 0.17 | 7.39 ± 0.04 | <5 | 5.54 ± 0.02 | <5 | <5 |
| **25N-NB-2,5-DiMeO (20)** | 6.29 ± 0.07 | <5 | <5 | <5 | 6.64 ± 0.05 | 6.87 ± 0.12 | 6.84 ± 0.04 | <5 | <5 | <5 | <5 |
| **25N-NB-3-OH (21)** | 5.73 ± 0.10 | <5 | <5 | <5 | 7.84 ± 0.08 | 7.45 ± 0.09 | 7.08 ± 0.05 | <5 | <5 | 7.16 ± 0.10 | <5 |
| **25N-NB-3-Me (22)** | 5.75 | <5 | 6.03 | <5 | 9.68 | 8.54 | 7.40 | <5 | ND | ND | ND |
| **25N-NB-4-Me (23)** | 5.58 | <5 | <5 | <5 | 8.26 | 7.28 | 6.20 | <5 | ND | ND | ND |
| **25N-NB-3-F (24)** | <5 | <5 | <5 | <5 | 8.52 | 7.37 | 6.65 | <5 | ND | ND | ND |
| **25N-NB-4-F (25)** | <5 | <5 | <5 | <5 | 8.21 | 7.07 | 6.09 | <5 | ND | ND | ND |

**Supplementary Table 9. Off-Target Competitive Radioligand Binding Affinity (pKi) Values for 25N Series.** Compounds were assayed as described by NIMH PDSP. Data presented as N = 1. <5 indicates compounds were inactive in a primary screening at 10 μM.

| Receptors | 25N (1) | 25N-NB (2) | 25N-NBOH (3) | 25N-NBOMe (4) | 25N-NBOEt (5) | 25N-NBMe (6) | 25N-NBF (7) | 25N-NBCl (8) | 25N-NBBr (9) | 25N-NBI (10) | 25N-NBOCF$_2$H (11) | 25N-NBOCF$_3$ (12) | 25N-NBNO$_2$ (13) |
|---|---|---|---|---|---|---|---|---|---|---|---|---|---|
| **Alpha1A** | <5.00 | <5.00 | <5.00 | <5.00 | <5.00 | <5.00 | <5.00 | <5.00 | <5.00 | <5.00 | <5.00 | <5.00 | <5.00 |
| **Alpha1B** | <5.00 | <5.00 | <5.00 | <5.00 | <5.00 | <5.00 | <5.00 | <5.00 | <5.00 | 6.06 | <5.00 | <5.00 | <5.00 |
| **Alpha1D** | <5.00 | <5.00 | <5.00 | <5.00 | 5.81 | <5.00 | <5.00 | <5.00 | <5.00 | <5.00 | 5.27 | <5.00 | <5.00 |
| **Alpha2A** | 6.62 | 5.74 | 6.04 | 6.30 | 5.92 | 6.19 | 6.27 | 6.14 | 6.17 | 5.57 | 5.90 | 6.05 | <5.00 |
| **Alpha2B** | 5.65 | 5.71 | 5.85 | 5.96 | 5.57 | 5.72 | 5.71 | 5.93 | 5.92 | 5.58 | 5.75 | 5.86 | <5.00 |
| **Alpha2C** | 6.05 | 6.20 | 6.39 | 6.16 | 6.38 | 5.87 | 6.37 | 6.13 | 6.29 | 6.36 | 6.46 | 6.45 | 5.77 |
| **Beta1** | <5.00 | <5.00 | 5.81 | <5.00 | <5.00 | <5.00 | <5.00 | <5.00 | <5.00 | <5.00 | <5.00 | <5.00 | <5.00 |
| **Beta2** | <5.00 | <5.00 | 6.18 | 5.15 | <5.00 | <5.00 | <5.00 | <5.00 | <5.00 | <5.00 | <5.00 | <5.00 | <5.00 |
| **Beta3** | <5.00 | <5.00 | <5.00 | <5.00 | <5.00 | <5.00 | <5.00 | <5.00 | <5.00 | <5.00 | <5.00 | <5.00 | <5.00 |
| **BZP** | <5.00 | <5.00 | <5.00 | <5.00 | <5.00 | <5.00 | <5.00 | <5.00 | <5.00 | <5.00 | <5.00 | <5.00 | <5.00 |
| **D1** | <5.00 | <5.00 | <5.00 | <5.00 | <5.00 | <5.00 | <5.00 | <5.00 | <5.00 | <5.00 | <5.00 | <5.00 | <5.00 |
| **D2** | <5.00 | <5.00 | <5.00 | 5.17 | <5.00 | <5.00 | <5.00 | 5.19 | <5.00 | <5.00 | <5.00 | <5.00 | <5.00 |
| **D3** | <5.00 | 5.90 | 5.86 | 5.66 | 5.61 | 5.65 | 5.84 | 5.88 | 6.06 | 6.18 | 6.08 | 6.00 | <5.00 |
| **D4** | <5.00 | 5.96 | 5.54 | <5.00 | <5.00 | 5.76 | 5.69 | 6.03 | 5.88 | 6.13 | 5.77 | 5.64 | <5.00 |
| **D5** | <5.00 | <5.00 | <5.00 | <5.00 | <5.00 | <5.00 | <5.00 | <5.00 | <5.00 | <5.00 | <5.00 | <5.00 | <5.00 |
| **DAT** | <5.00 | <5.00 | 5.80 | <5.00 | <5.00 | <5.00 | 5.06 | 5.93 | 5.28 | <5.00 | 5.36 | 5.57 | <5.00 |
| **DOR** | <5.00 | <5.00 | <5.00 | <5.00 | <5.00 | <5.00 | <5.00 | <5.00 | <5.00 | <5.00 | <5.00 | <5.00 | <5.00 |
| **GABAA** | <5.00 | <5.00 | <5.00 | <5.00 | <5.00 | <5.00 | <5.00 | <5.00 | <5.00 | <5.00 | <5.00 | <5.00 | <5.00 |
| **H1** | <5.00 | 6.65 | 5.65 | 7.04 | 6.47 | 6.41 | 5.86 | 6.47 | 6.60 | 6.55 | 6.07 | 6.74 | <5.00 |
| **H2** | <5.00 | 6.66 | <5.00 | 5.97 | 6.02 | 6.17 | <5.00 | 5.93 | 6.32 | 6.33 | ND | 6.00 | 6.19 |
| **H3** | 5.26 | <5.00 | <5.00 | <5.00 | <5.00 | <5.00 | <5.00 | <5.00 | <5.00 | <5.00 | <5.00 | <5.00 | <5.00 |
| **H4** | <5.00 | <5.00 | <5.00 | <5.00 | <5.00 | <5.00 | <5.00 | <5.00 | <5.00 | <5.00 | <5.00 | <5.00 | <5.00 |
| **KOR** | <5.00 | <5.00 | 5.32 | 5.55 | 5.33 | 5.71 | <5.00 | 6.12 | 5.51 | 5.79 | 5.47 | 5.47 | <5.00 |
| **M1** | <5.00 | <5.00 | <5.00 | <5.00 | <5.00 | <5.00 | <5.00 | <5.00 | <5.00 | <5.00 | <5.00 | <5.00 | <5.00 |
| **M2** | <5.00 | <5.00 | <5.00 | <5.00 | <5.00 | <5.00 | <5.00 | <5.00 | <5.00 | <5.00 | <5.00 | <5.00 | <5.00 |
| **M3** | 5.00 | <5.00 | <5.00 | <5.00 | <5.00 | <5.00 | <5.00 | <5.00 | <5.00 | <5.00 | <5.00 | <5.00 | <5.00 |
| **M4** | <5.00 | <5.00 | <5.00 | <5.00 | <5.00 | <5.00 | <5.00 | <5.00 | 5.00 | <5.00 | <5.00 | <5.00 | <5.00 |
| **M5** | <5.00 | <5.00 | <5.00 | <5.00 | <5.00 | <5.00 | <5.00 | <5.00 | 5.00 | <5.00 | <5.00 | <5.00 | <5.00 |
| **MOR** | <5.00 | 5.29 | <5.00 | <5.00 | 5.95 | <5.00 | <5.00 | <5.00 | 5.81 | 5.86 | 5.72 | 5.67 | 5.25 |
| **NET** | <5.00 | <5.00 | <5.00 | <5.00 | <5.00 | <5.00 | <5.00 | <5.00 | 5.49 | <5.00 | <5.00 | <5.00 | <5.00 |
| **SERT** | <5.00 | <5.00 | <5.00 | 5.85 | <5.00 | <5.00 | <5.00 | <5.00 | 5.00 | <5.00 | <5.00 | <5.00 | <5.00 |
| **Sigma 1** | <5.00 | 7.37 | 6.65 | 6.27 | 5.87 | 6.84 | 6.72 | 6.81 | 6.48 | 6.25 | 5.89 | 5.44 | 6.32 |

| | | | | | | | | | | | | | |
|---|---|---|---|---|---|---|---|---|---|---|---|---|---|
| **Sigma 2** | <5.00 | 6.95 | 6.76 | 7.24 | 7.09 | 7.55 | 7.33 | 7.55 | 7.51 | 7.26 | 7.26 | 7.18 | 6.83 |

**Supplementary Table 10. Off-Target Competitive Radioligand Binding Affinity (pKi) Values for 25N Series.** Compounds were assayed as described by NIMH PDSP. Data presented as N = 1. <5 indicates compounds were inactive in a primary screening at 10 μM.

| Receptors | 25N-N1-Nap (16) | 25N-NBPh (17) | 25N-NB-2-OH-3-Me (18) | 25N-NB-2-MeO-3-F (19) | 25N-NB-2,5-DiMeO (20) | 25N-NB-3-OH (21) | 25N-NB-3-Me (22) | 25N-NB-4-Me (23) | 25N-NB-3-F (24) | 25N-NB-4-F (25) |
|---|---|---|---|---|---|---|---|---|---|---|
| **Alpha1A** | 6.32 | 5.82 | <5.00 | <5.00 | <5.00 | <5.00 | <5.00 | <5.00 | <5.00 | <5.00 |
| **Alpha1B** | 6.35 | 5.89 | <5.00 | <5.00 | <5.00 | <5.00 | <5.00 | <5.00 | <5.00 | <5.00 |
| **Alpha1D** | 6.19 | <5.00 | <5.00 | <5.00 | <5.00 | <5.00 | <5.00 | <5.00 | <5.00 | <5.00 |
| **Alpha2A** | 6.35 | 6.36 | 5.24 | 5.90 | 5.99 | 5.71 | 6.80 | 5.98 | 6.11 | 6.03 |
| **Alpha2B** | 7.00 | <5.00 | <5.00 | 5.83 | 5.63 | <5.00 | 6.14 | 5.63 | 5.50 | 5.66 |
| **Alpha2C** | 7.00 | 5.86 | 6.04 | 5.95 | 5.84 | 6.15 | 5.77 | 5.37 | 5.45 | <5.00 |
| **Beta1** | <5.00 | <5.00 | 6.04 | <5.00 | <5.00 | <5.00 | <5.00 | <5.00 | <5.00 | <5.00 |
| **Beta2** | 5.82 | <5.00 | <5.00 | <5.00 | <5.00 | <5.00 | <5.00 | <5.00 | <5.00 | <5.00 |
| **Beta3** | 6.04 | <5.00 | <5.00 | <5.00 | <5.00 | <5.00 | <5.00 | <5.00 | <5.00 | <5.00 |
| **BZP** | <5.00 | <5.00 | <5.00 | <5.00 | <5.00 | <5.00 | <5.00 | <5.00 | <5.00 | <5.00 |
| **D1** | 6.21 | <5.00 | <5.00 | <5.00 | <5.00 | <5.00 | <5.00 | <5.00 | <5.00 | <5.00 |
| **D2** | <5.00 | <5.00 | <5.00 | <5.00 | <5.00 | <5.00 | <5.00 | <5.00 | <5.00 | <5.00 |
| **D3** | 7.07 | 5.98 | 5.72 | <5.00 | 5.95 | <5.00 | 5.88 | 6.61 | 5.77 | 5.95 |
| **D4** | 6.25 | 5.87 | 5.86 | 5.82 | 5.71 | <5.00 | 6.16 | 6.19 | <5.00 | <5.00 |
| **D5** | <5.00 | <5.00 | <5.00 | <5.00 | <5.00 | <5.00 | <5.00 | <5.00 | <5.00 | <5.00 |
| **DAT** | <5.00 | 5.63 | <5.00 | <5.00 | <5.00 | <5.00 | 5.48 | 5.66 | 5.46 | 5.76 |
| **DOR** | <5.00 | <5.00 | <5.00 | <5.00 | <5.00 | <5.00 | <5.00 | <5.00 | <5.00 | <5.00 |
| **GABAA** | <5.00 | <5.00 | <5.00 | <5.00 | <5.00 | <5.00 | <5.00 | <5.00 | <5.00 | <5.00 |
| **H1** | 5.80 | 6.13 | 5.17 | 6.82 | <5,.00 | 6.22 | 6.15 | <5.00 | 5.97 | <5.00 |
| **H2** | <5.00 | 6.97 | 6.23 | <5.00 | 6.39 | 6.34 | 6.93 | 6.87 | 6.42 | 6.72 |
| **H3** | <5.00 | <5.00 | <5.00 | <5.00 | <5.00 | <5.00 | <5.00 | <5.00 | <5.00 | <5.00 |
| **H4** | <5.00 | <5.00 | <5.00 | <5.00 | <5.00 | <5.00 | <5.00 | <5.00 | <5.00 | <5.00 |
| **KOR** | 5.55 | 6.45 | 5.53 | 5.98 | 6.13 | 6.10 | 5.51 | 5.90 | 5.05 | <5.00 |
| **M1** | <5.00 | <5.00 | <5.00 | <5.00 | 5.42 | <5.00 | <5.00 | <5.00 | <5.00 | <5.00 |
| **M2** | <5.00 | <5.00 | <5.00 | <5.00 | <5.00 | <5.00 | <5.00 | <5.00 | <5.00 | <5.00 |
| **M3** | 5.11 | <5.00 | <5.00 | <5.00 | 5.56 | <5.00 | <5.00 | <5.00 | <5.00 | <5.00 |
| **M4** | 6.05 | <5.00 | <5.00 | <5.00 | <5.00 | <5.00 | <5.00 | <5.00 | <5.00 | <5.00 |
| **M5** | <5.00 | <5.00 | <5.00 | <5.00 | <5.00 | <5.00 | <5.00 | <5.00 | <5.00 | <5.00 |
| **MOR** | 5.70 | 6.10 | 5.25 | 6.03 | 6.07 | 5.68 | <5.00 | 5.51 | <5.00 | <5.00 |
| **NET** | <5.00 | 6.19 | <5.00 | <5.00 | <5.00 | 5.56 | <5.00 | <5.00 | <5.00 | <5.00 |
| **SERT** | <5.00 | 5.21 | <5.00 | <5.00 | <5.00 | 5.39 | 5.31 | 5.40 | 5.39 | 5.31 |

| | | | | | | | | | | |
|---|---|---|---|---|---|---|---|---|---|---|
| **Sigma 1** | 5.93 | 6.30 | 5.89 | 6.42 | 6.38 | 6.83 | 7.62 | 7.74 | 7.82 | 8.10 |

**Supplementary Table 11. 5-$HT_{2A}$ Gq and β-Arrestin2 Bioluminescence Resonance Energy Transfer (BRET) Assay Functional Data for Phenethylamine Psychedelics.** Data presented as mean ± SEM from three biological replicates.

| Compound | 5-$HT_{2A}$ | | | |
|---|---|---|---|---|
| | Gq Dissociation (BRET) | | β-Arrestin2 Association (BRET) | |
| | pEC50 ± SEM | $E_{MAX}$ ± SEM (%5-HT) | pEC50 ± SEM | $E_{MAX}$ ± SEM (%5-HT) |
| **2C-B** | 8.92 ± 0.03 | 100.8 ± 0.9 | 8.62 ± 0.04 | 84.0 ± 1.0 |
| **2C-I** | 8.83 ± 0.05 | 93.4 ± 1.4 | 8.25 ± 0.07 | 74.3 ± 1.6 |
| **25N (1)** | 7.88 ± 0.04 | 87.9 ± 1.4 | 7.81 ± 0.08 | 83.8 ± 2.5 |
| **2C-2** | 6.83 ± 0.02 | 102.0 ± 0.9 | 6.23 ± 0.10 | 107.3 ± 4.6 |
| **2C-O** | 6.71 ± 0.04 | 99.5 ± 1.6 | 6.17 ± 0.08 | 96.2 ± 3.1 |
| **2C-T-28** | 7.57 ± 0.17 | 95.3 ± 5.0 | 7.60 ± 0.09 | 87.7 ± 2.4 |
| **2C-B-FLY** | 9.05 ± 0.03 | 102.0 ± 0.7 | 8.42 ± 0.06 | 80.1 ± 1.5 |
| **2C-I-FLY** | 8.92 ± 0.04 | 100.1 ± 1.2 | 8.53 ± 0.07 | 73.5 ± 1.4 |
| **Mescaline** | 6.19 ± 0.07 | 74.3 ± 2.3 | 6.37 ± 0.16 | 33.0 ± 2.1 |
| **Proscaline** | 6.36 ± 0.06 | 96.9 ± 2.4 | 6.20 ± 0.05 | 92.7 ± 2.0 |
| **Desoxy** | 6.26 ± 0.06 | 98.2 ± 2.3 | 6.29 ± 0.09 | 82.9 ± 3.1 |
| **3C-B-FLY** | 9.30 ± 0.03 | 105.0 ± 1.0 | 8.99 ± 0.05 | 89.0 ± 1.3 |
| **2H-DOB** | 6.58 ± 0.07 | 95.8 ± 2.4 | 6.79 ± 0.06 | 93.3 ± 1.9 |
| **5H-DOB** | 7.31 ± 0.06 | 102.1 ± 2.1 | 7.37 ± 0.09 | 87.9 ± 2.6 |
| **DOB** | 9.21 ± 0.03 | 104.9 ± 1.0 | 8.93 ± 0.05 | 101.0 ± 1.3 |
| **DOI** | 8.14 ± 0.12 | 108.2 ± 4.2 | 8.66 ± 0.08 | 88.8 ± 1.9 |
| **DOET** | 7.76 ± 0.03 | 102.3 ± 1.1 | 7.51 ± 0.06 | 98.1 ± 1.8 |
| **DOBU** | 8.72 ± 0.04 | 103.5 ± 1.3 | 8.59 ± 0.05 | 98.0 ± 1.3 |
| **MEM** | 6.53 ± 0.06 | 101.5 ± 2.3 | 6.79 ± 0.11 | 104.7 ± 4.5 |
| **3C-P** | 6.75 ± 0.06 | 96.4 ± 2.2 | 6.64 ± 0.13 | 87.5 ± 4.3 |
| **CPM** | 6.15 ± 0.06 | 95.5 ± 2.4 | 6.06 ± 0.11 | 82.5 ± 3.8 |
| **Bromo-Dragonfly** | 9.83 ± 0.06 | 104.7 ± 1.6 | 9.48 ± 0.06 | 97.6 ± 1.6 |
| **α-Ethyl-2C-D** | 6.85 ± 0.05 | 89.1 ± 1.6 | 6.90 ± 0.05 | 83.0 ± 1.5 |
| **25H-NBOH** | 8.17 ± 0.07 | 97.8 ± 2.0 | 8.32 ± 0.08 | 118.2 ± 2.8 |
| **25C-NBOH** | 9.57 ± 0.04 | 100.6 ± 0.9 | 9.57 ± 0.04 | 114.4 ± 1.3 |
| **25B-NBOH** | 9.58 ± 0.05 | 100.5 ± 1.2 | 9.54 ± 0.08 | 112.5 ± 2.4 |
| **25CN-NBOH** | 9.24 ± 0.06 | 97.0 ± 1.6 | 9.18 ± 0.04 | 112.4 ± 1.3 |
| **25D-NBOMe** | 9.65 ±0.05 | 93.0 ± 1.3 | 9.43 ± 0.09 | 126.0 ± 3.4 |
| **25I-NBOMe** | 9.45 ± 0.04 | 99.8 ± 1.2 | 9.28 ± 0.09 | 137.2 ± 3.7 |
| **25C-NBMD** | 8.70 ± 0.07 | 85.0 ± 1.6 | 8.69 ± 0.15 | 121.1 ± 5.0 |

**Supplementary Table 12. 5-HT$_{2B}$ and 5-HT$_{2C}$ Gq Bioluminescence Resonance Energy Transfer (BRET) Assay Functional Data for Phenethylamine Psychedelics.** Data presented as mean ± SEM from three biological replicates.

| Compound | 5-HT$_{2B}$ Gq Dissociation (BRET) | | 5-HT$_{2C}$ Gq Dissociation (BRET) | |
|---|---|---|---|---|
| | pEC50 ± SEM | $E_{MAX}$ ± SEM (%5-HT) | pEC50 ± SEM | $E_{MAX}$ ± SEM (%5-HT) |
| **2C-B** | 7.90 ± 0.04 | 97.4 ± 1.4 | 9.20 ± 0.05 | 97.8 ± 1.5 |
| **2C-I** | 7.72 ± 0.07 | 101.0 ± 2.1 | 9.34 ± 0.05 | 106.9 ± 1.5 |
| **25N (1)** | 7.13 ± 0.08 | 87.7 ± 2.8 | 7.90 ± 0.09 | 78.4 ± 2.5 |
| **2C-2** | 6.69 ± 0.06 | 103.8 ± 2.4 | 8.01 ± 0.08 | 109.3 ± 2.7 |
| **2C-O** | 6.57 ± 0.09 | 103.9 ± 3.5 | 7.30 ± 0.11 | 99.6 ± 3.6 |
| **2C-T-28** | 6.78 ± 0.13 | 100.5 ± 4.9 | 9.23 ± 0.15 | 105.4 ± 4.4 |
| **2C-B-FLY** | 9.14 ± 0.05 | 108.4 ± 1.4 | 9.91 ± 0.04 | 103.6 ± 1.1 |
| **2C-I-FLY** | 8.92 ± 0.06 | 109.9 ± 2.0 | 9.73 ± 0.06 | 99.3 ± 1.5 |
| **Mescaline** | 5.96 ± 0.10 | 91.1 ± 4.1 | 6.50 ± 0.12 | 108.5 ± 5.0 |
| **Proscaline** | 6.18 ± 0.09 | 83.3 ± 3.1 | 7.42 ± 0.08 | 100.5 ± 2.7 |
| **Desoxy** | 6.66 ± 0.16 | 100.4 ± 6.1 | 7.98 ± 0.19 | 117.8 ± 7.3 |
| **3C-B-FLY** | 9.37 ± 0.06 | 108.8 ± 1.7 | 9.90 ± 0.04 | 107.2 ± 1.2 |
| **2H-DOB** | 7.05 ± 0.10 | 105.1 ± 3.7 | 7.73 ± 0.10 | 113.9 ± 3.5 |
| **5H-DOB** | 7.29 ± 0.11 | 103.5 ± 3.9 | 8.28 ± 0.08 | 111.2 ± 2.7 |
| **DOB** | 8.55 ± 0.05 | 100.4 ± 1.4 | 9.30 ± 0.04 | 104.5 ± 1.3 |
| **DOI** | 7.90 ± 0.09 | 103.2 ± 2.8 | 9.08 ± 0.11 | 114.3 ± 3.4 |
| **DOET** | 6.96 ± 0.05 | 108.1 ± 2.1 | 7.77 ± 0.04 | 101.9 ± 1.3 |
| **DOBU** | 7.08 ± 0.08 | 69.7 ± 2.1 | 9.04 ± 0.04 | 100.1 ± 1.3 |
| **MEM** | 6.36 ± 0.10 | 95.7 ± 3.7 | 7.07 ± 0.12 | 97.7 ± 4.0 |
| **3C-P** | 6.77 ± 0.15 | 70.1 ± 3.8 | 7.06 ± 0.11 | 111.2 ± 4.5 |
| **CPM** | 5.57 ± 0.16 | 105.3 ± 7.9 | 6.86 ± 0.09 | 111.0 ± 3.7 |
| **Bromo-Dragonfly** | 9.85 ± 0.08 | 105.0 ± 2.1 | 10.21 ± 0.06 | 104.0 ± 1.6 |
| **α-Ethyl-2C-D** | 6.23 ± 0.08 | 85.3 ± 2.7 | 6.92 ± 0.06 | 92.6 ± 2.1 |
| **25H-NBOH** | 6.10 ± 0.15 | 46.9 ± 2.9 | 8.12 ± 0.07 | 106.7 ± 2.2 |
| **25C-NBOH** | 7.66 ± 0.17 | 55.5 ± 3.0 | 9.67 ± 0.04 | 105.5 ± 1.1 |
| **25B-NBOH** | 7.45 ± 0.18 | 55.3 ± 3.2 | 9.96 ± 0.04 | 104.8 ± 1.0 |
| **25CN-NBOH** | 6.84 ± 0.14 | 56.9 ± 2.9 | 8.32 ± 0.07 | 113.9 ± 2.5 |
| **25I-NBOMe** | 8.47 ± 0.10 | 78.6 ± 2.6 | 10.01 ± 0.09 | 123.8 ± 3.1 |
| **25C-NBMD** | 6.36 ± 0.32 | 33.6 ± 4.2 | 8.83 ± 0.07 | 117.1 ± 2.3 |

**Supplementary Table 13. 5-HT$_{2A}$ Gq and β-Arrestin2 Bioluminescence Resonance Energy Transfer (BRET) Assay Functional Data for 25N Series.** Data presented as mean ± SEM from three biological replicates.

| Compound | 5-HT$_{2A}$ | | | |
|---|---|---|---|---|
| | Gq Dissociation (BRET) | | β-Arrestin2 Association (BRET) | |
| | pEC50 ± SEM | $E_{MAX}$ ± SEM (%5-HT) | pEC50 ± SEM | $E_{MAX}$ ± SEM (%5-HT) |
| **25N-NB (2)** | 8.70 ± 0.06 | 79.5 ± 1.5 | 8.96 ± 0.11 | 115.1 ± 4.1 |
| **25N-NBOH (3)** | 9.65 ± 0.03 | 99.7 ± 0.8 | 10.12 ± 0.19 | 106.7 ± 5.8 |
| **25N-NBOMe (4)** | 9.69 ± 0.04 | 95.2 ± 1.2 | 9.66 ± 0.09 | 136.5 ± 3.6 |
| **25N-NBMe (6)** | 9.01 ± 0.05 | 82.2 ± 1.3 | 8.56 ± 0.22 | 83.2 ± 5.9 |
| **25N-NBF (7)** | 8.55 ± 0.06 | 80.1 ± 1.6 | 8.16 ± 0.24 | 89.5 ± 7.5 |
| **25N-NBCl (8)** | 8.74 ± 0.06 | 79.0 ± 1.6 | 7.70 ± 0.21 | 100.8 ± 7.7 |
| **25N-NBBr (9)** | 8.74 ± 0.05 | 75.7 ± 1.2 | 8.27 ± 0.29 | 77.2 ± 8.0 |
| **25N-NBI (10)** | 8.40 ± 0.04 | 73.9 ± 1.0 | 8.38 ± 0.12 | 106.4 ± 4.1 |
| **25N-NB$CF_3$ (14)** | 7.62 ± 0.09 | 76.9 ± 2.6 | 7.37 ± 0.13 | 104.4 ± 5.2 |
| **25N-NB$NO_2$ (15)** | 7.26 ± 0.07 | 54.0 ± 1.6 | 7.40 ± 0.08 | 88.7 ± 2.7 |
| **25N-N1-Nap (16)** | 8.84 ± 0.29 | 22.5 ± 2.1 | 9.39 ± 0.08 | 78.2 ± 1.8 |
| **25N-NBPh (17)** | 8.26 ± 0.18 | 25.7 ± 1.4 | 8.39 ± 0.08 | 109.4 ± 2.8 |
| **25N-NB-2-OH-3-Me (18)** | 9.49 ± 0.11 | 50.8 ± 1.6 | 9.43 ± 0.10 | 130.7 ± 4.0 |
| **25N-NB-2-MeO-3-F (19)** | 8.56 ± 0.08 | 71.1 ± 1.9 | 8.33 ± 0.08 | 120.4 ± 3.3 |
| **25N-NB-3-OH (21)** | 7.94 ± 0.23 | 49.8 ± 4.3 | 8.36 ± 0.13 | 118.6 ± 5.2 |
| **25N-NB-3-Me (22)** | 8.64 ± 0.25 | 41.7 ± 3.4 | 9.08 ± 0.14 | 94.7 ± 4.2 |
| **25D-N-1-Nap (26)** | 8.16 ± 0.24 | 34.3 ± 2.9 | 8.86 ± 0.10 | 75.5 ± 2.4 |
| **25D-NBPh (27)** | 6.09 ± 0.16 | 23.2 ± 4.8 | 7.64 ± 0.12 | 107.0 ± 4.7 |
| **25O-N1-Nap (28)** | 8.26 ± 0.14 | 24.3 ± 1.1 | 8.79 ± 0.08 | 74.9 ± 1.9 |
| **2C2-N1-Nap (29)** | 7.81 ± 0.20 | 26.4 ± 1.9 | 8.23 ± 0.09 | 80.1 ± 2.4 |
| **Ψ-DOB-N1-Nap (30)** | 7.50 ± 0.08 | 36.7 ± 1.1 | 8.31 ± 0.08 | 98.3 ± 3.3 |
| **2C2-NBOMe (31)** | 9.10 ± 0.03 | 98.4 ± 1.0 | 8.85 ± 0.05 | 141.8 ± 2.1 |
| **25O-NBOMe (32)** | 8.99 ± 0.03 | 99.9 ± 0.8 | 8.66 ± 0.06 | 128.7 ± 2.3 |
| **25O-NBcP (33)** | 8.48 ± 0.04 | 96.3 ± 1.4 | 8.15 ± 0.07 | 116.6 ± 2.7 |
| **25O-NB-3-I (34)** | 7.74 ± 0.07 | 51.7 ± 1.3 | 8.42 ± 0.10 | 56.0 ± 1.8 |
| **25O-NBPh-10'-OH (35)** | 6.26 ± 0.06 | 60.8 ± 1.9 | 7.19 ± 0.06 | 132.1 ± 3.3 |

**Supplementary Table 14. 5-$HT_{2B}$ and 5-$HT_{2C}$ Gq Bioluminescence Resonance Energy Transfer (BRET) Assay Functional Data for 25N Series.** Data presented as mean ± SEM from three biological replicates. NA = no activity; NC = not calculated.

| Compound | 5-$HT_{2B}$ | | 5-$HT_{2C}$ | |
|---|---|---|---|---|
| | Gq Dissociation (BRET) | | Gq Dissociation (BRET) | |
| | pEC50 ± SEM | $E_{MAX}$ ± SEM (%5-HT) | pEC50 ± SEM | $E_{MAX}$ ± SEM (%5-HT) |
| **25N-NB (2)** | 7.42 ± 0.19 | 32.6 ± 2.3 | 8.07 ± 0.07 | 89.4 ± 2.3 |
| **25N-NBOMe (4)** | 8.63 ± 0.14 | 54.1 ± 2.5 | 9.34 ± 0.08 | 100.0 ± 2.5 |
| **25N-NBMe (6)** | 8.02 ± 0.44 | 22.2 ± 3.2 | 7.94 ± 0.07 | 85.9 ± 1.9 |
| **25N-NBF (7)** | 7.12 ± 0.27 | 24.8 ± 3.2 | 7.85 ± 0.07 | 83.1 ± 2.1 |
| **25N-NBCl (8)** | 6.69 ± 0.25 | 27.0 ± 3.0 | 7.67 ± 0.10 | 82.5 ± 2.8 |
| **25N-NBBr (9)** | 6.61 ± 0.28 | 17.0 ± 2.1 | 7.51 ± 0.08 | 81.0 ± 2.3 |
| **25N-NBI (10)** | 7.53 ± 0.31 | 21.8 ± 2.3 | 7.06 ± 0.07 | 71.5 ± 2.2 |
| **25N-$NBNO_2$ (15)** | NA | NA | 6.83 ± 0.17 | 61.3 ± 4.4 |
| **25N-N1-Nap (16)** | NA | NA | 7.88 ± 0.09 | 64.6 ± 2.1 |
| **25N-NBPh (17)** | NA | NA | <6.00 | NC |
| **25N-NB-2-OH-3-Me (18)** | NA | NA | 8.67 ± 0.17 | 45.8 ± 2.4 |
| **25O-N1-Nap (28)** | NA | NA | 7.90 ± 0.09 | 64.4 ± 2.0 |
| **2C2-N1-Nap (29)** | NA | NA | 7.77 ± 0.07 | 90.5 ± 2.3 |
| **psi-DOB-N1-Nap (30)** | NA | NA | 7.19 ± 0.07 | 80.3 ± 2.4 |

**Supplementary Table 15. Effect of Test Compounds on the Head-Twitch Response (HTR) in Mice.** *N.D.* not determined.

| **Table 15. Effect of 2C-N derivatives on the head-twitch response (HTR) in mice.** | | | |
|---|---|---|---|
| **Compound** | **HTR $ED_{50}$** | | **Maximum magnitude of the HTR (counts per minute)** |
| | **mg/kg (95% CI)** | **μmol/kg (95% CI)** | |
| **25N-NB (2)** | 1.25 (0.96-1.62) | 3.53 (2.71-4.60) | 2.027 |
| **25N-NBOH (3)** | 0.07 (0.04-0.11) | 0.19 (0.12-0.29) | 2.980 |
| **25N-NBOMe (4)** | 0.11 (0.08-0.16) | 0.29 (0.20-0.43) | 4.890 |
| **25N-NBOEt (5)** | 0.57 (0.45-0.73) | 1.44 (1.13-1.83) | 3.600 |
| **25N-NBMe (6)** | 1.19 (0.49-2.91) | 3.24 (1.32-7.94) | 1.807 |
| **25N-NBF (7)** | 1.52 (0.98-2.35) | 4.10 (2.65-6.33) | 1.251 |
| **25N-NBCl (8)** | 4.04 (1.59-10.3) | 10.4 (4.11-26.5) | 0.633 |
| **25N-NBBr (9)** | 6.31 (2.90-13.7) | 14.6 (6.71-31.8) | 1.378 |
| **25N-NBI (10)** | 5.23 (2.18-12.54) | 10.9 (4.55-26.2) | 0.907 |
| **25N-NBOCF$_2$H (11)** | 1.81 (1.23-2.67) | 4.32 (2.93-6.38) | 3.222 |
| **25N-NBOCF$_3$ (12)** | 5.90 (2.97-11.7) | 13.5 (6.81-26.8) | 1.060 |
| **25N-NBCF$_3$ (14)** | Inactive up to 30[1] | *N.D.* | 0.700 |
| **25N-NBNO$_2$ (15)** | Inactive up to 100 | *N.D.* | 0.880 |
| **25N-N1-Nap (16)** | Inactive up to 30 | *N.D.* | 0.561 |
| **25N-NBPh (17)** | Inactive up to 100 | *N.D.* | 0.692 |
| **25N-NB-2-HO-3-Me (18)** | Inactive up to 10 | *N.D.* | 0.467 |
| **25N-NB-3-HO (21)** | Inactive up to 30 | *N.D.* | 0.473 |

*N.D.*, not determined.
[1]Based on the absence of significant post-hoc pairwise difference between drug and vehicle control.

### Supplementary Table 16. Summary of the Head-Twitch Response (HTR) Data for 25N Analogs in Mice.

| Compound | ANOVA | Dose (mg/kg) | *N* | Mean | SEM | Post-Hoc Result[1] |
|---|---|---|---|---|---|---|
| **25N-NB (2)** | $F_{5,26} = 84.56$, $p < 0.0001$ | 0 | 6 | 6.7 | 1.1 | |
| | | 0.3 | 5 | 11.8 | 0.6 | |
| | | 1 | 5 | 32.0 | 2.1 | **** |
| | | 3 | 6 | 42.8 | 2.3 | **** |
| | | 10 | 5 | 60.8 | 2.7 | **** |
| | | 30 | 5 | 35.0 | 3.4 | **** |
| **25N-NBOH (3)** | $W_{5,9.70} = 40.11$, $p < 0.0001$ | 0 | 5 | 10.2 | 1.2 | |
| | | 0.01 | 4 | 15.0 | 2.5 | |
| | | 0.03 | 5 | 33.0 | 5.1 | |
| | | 0.1 | 5 | 62.2 | 5.1 | ** |
| | | 0.3 | 5 | 89.4 | 6.8 | ** |
| | | 1 | 5 | 89.4 | 15.6 | |
| **25N-NBOMe (4)** | $W_{5,19.46} = 84.56$, $p < 0.0001$ | 0 | 10 | 9.4 | 1.4 | |
| | | 0.03 | 10 | 40.5 | 7.3 | * |
| | | 0.1 | 11 | 70.7 | 7.3 | **** |
| | | 0.3 | 11 | 122.9 | 8.2 | **** |
| | | 1 | 10 | 146.7 | 12.7 | **** |
| | | 3 | 6 | 143.5 | 21.0 | * |
| **25N-NBOEt (5)** | $F_{5,25} = 61.29$, $p < 0.0001$ | 0 | 6 | 8.2 | 1.9 | |
| | | 0.1 | 5 | 18.2 | 3.8 | |
| | | 0.3 | 5 | 35.8 | 3.2 | ** |
| | | 1 | 5 | 74.4 | 4.3 | **** |
| | | 3 | 5 | 108.0 | 9.2 | **** |
| | | 10 | 5 | 77.8 | 5.5 | **** |
| **25N-NBMe (6)** | $F_{5,23} = 8.34$, $p = 0.0001$ | 0 | 5 | 6.8 | 1.7 | |
| | | 0.3 | 4 | 15.8 | 3.1 | |
| | | 1 | 5 | 31.4 | 8.4 | |
| | | 3 | 5 | 35.4 | 6.4 | * |
| | | 10 | 5 | 54.2 | 6.0 | *** |
| | | 30 | 5 | 51.0 | 8.9 | *** |
| **25N-NBF (7)** | $W_{5,32.56} = 40.02$, $p < 0.0001$ | 0 | 17 | 8.3 | 0.7 | |
| | | 0.3 | 15 | 13.1 | 1.9 | |
| | | 1 | 15 | 19.3 | 1.8 | *** |
| | | 3 | 15 | 30.1 | 2.8 | **** |
| | | 10 | 15 | 37.5 | 2.4 | **** |
| | | 30 | 10 | 37.5 | 5.6 | ** |
| **25N-NBCl (8)** | $F_{4,21} = 5.14$, $p = 0.0048$ | 0 | 6 | 8.2 | 2.4 | |
| | | 3 | 5 | 12.2 | 1.5 | |
| | | 10 | 5 | 18.6 | 2.2 | * |

| | | | | | | |
|---|---|---|---|---|---|---|
| | | 30 | 5 | 19.0 | 1.8 | ** |
| | | 100 | 5 | 15.6 | 2.0 | |
| **25N-NBBr (9)** | $F_{5,30} = 9.53$, $p < 0.0001$ | 0 | 7 | 8.9 | 0.8 | |
| | | 1 | 5 | 8.2 | 0.9 | |
| | | 3 | 6 | 15.7 | 2.4 | |
| | | 10 | 6 | 31.0 | 4.5 | * |
| | | 30 | 6 | 35.2 | 7.1 | ** |
| | | 100 | 6 | 41.3 | 7.1 | *** |
| **25N-NBI (10)** | $F_{5,24} = 14.48$, $p < 0.0001$ | 0 | 6 | 9.0 | 0.9 | |
| | | 0.3 | 4 | 10.5 | 3.0 | |
| | | 1 | 5 | 11.2 | 2.3 | |
| | | 3 | 5 | 14.4 | 2.5 | |
| | | 10 | 5 | 23.8 | 2.1 | *** |
| | | 30 | 5 | 27.2 | 1.4 | **** |
| **25N-NBOCF$_2$ (11)** | $F_{5,26} = 38.09$, $p < 0.0001$ | 0 | 6 | 7.1 | 1.1 | |
| | | 0.3 | 5 | 15.0 | 1.7 | |
| | | 1 | 5 | 44.4 | 4.2 | ** |
| | | 3 | 5 | 57.4 | 8.0 | **** |
| | | 10 | 6 | 96.7 | 8.5 | **** |
| | | 30 | 5 | 92.2 | 8.6 | **** |
| **25N-NBOCF$_3$ (12)** | $F_{6,53} = 3.49$, $p = 0.0055$ | 0 | 13 | 8.2 | 1.1 | |
| | | 0.3 | 5 | 8.8 | 0.5 | |
| | | 1 | 5 | 14.6 | 2.3 | |
| | | 3 | 10 | 14.4 | 1.8 | |
| | | 10 | 10 | 24.0 | 4.5 | |
| | | 30 | 10 | 31.8 | 8.7 | ** |
| | | 100 | 6 | 22.0 | 5.9 | |
| **25N-NBCF$_3$ (14)** | $F_{5,23} = 2.72$, $p = 0.0451$ | 0 | 5 | 8.0 | 2.0 | |
| | | 0.3 | 5 | 9.6 | 1.0 | |
| | | 1 | 6 | 10.7 | 2.0 | |
| | | 3 | 4 | 10.5 | 4.3 | |
| | | 10 | 5 | 16.4 | 1.2 | |
| | | 30 | 4 | 21.0 | 6.2 | |
| **25N-NBNO$_2$ (15)** | $F_{5,24} = 1.17$, $p = 0.3528$ | 0 | 5 | 14.0 | 4.1 | |
| | | 1 | 5 | 16.8 | 2.4 | |
| | | 3 | 5 | 19.4 | 4.2 | |
| | | 10 | 5 | 26.4 | 4.6 | |
| | | 30 | 5 | 15.0 | 3.2 | |
| | | 100 | 5 | 21.2 | 6.0 | |
| **25N-N1-Nap (16)** | $F_{5,26} = 1.39$, $p = 0.2618$ | 0 | 6 | 14.5 | 3.3 | |
| | | 0.3 | 5 | 8.4 | 2.4 | |
| | | 1 | 5 | 9.0 | 0.9 | |

| | | | | | |
|---|---|---|---|---|---|
| | | 3 | 5 | 7.6 | 1.4 |
| | | 10 | 6 | 16.8 | 5.2 |
| | | 30 | 5 | 12.8 | 2.6 |
| **25N-NBPh (17)** | $F_{4,18} = 0.89$, $p = 0.4903$ | 0 | 5 | 10.0 | 2.1 |
| | | 3 | 4 | 20.8 | 9.7 |
| | | 10 | 4 | 17.3 | 5.6 |
| | | 30 | 6 | 17.7 | 3.5 |
| | | 100 | 4 | 10.8 | 1.4 |
| **25N-NB-2-OH-3-Me (18)** | $F_{5,25} = 0.55$, $p = 0.7377$ | 0 | 6 | 13.7 | 3.2 |
| | | 0.1 | 5 | 9.6 | 2.3 |
| | | 0.3 | 5 | 9.8 | 2.0 |
| | | 1 | 5 | 9.6 | 1.9 |
| | | 3 | 5 | 13.4 | 4.6 |
| | | 10 | 5 | 14.0 | 2.9 |
| **25N-NB-3-OH (21)** | $F_{5,25} = 2.41$, $p = 0.0645$ | 0 | 6 | 8.5 | 1.2 |
| | | 0.3 | 5 | 9.6 | 0.8 |
| | | 1 | 5 | 8.8 | 1.1 |
| | | 3 | 5 | 9.2 | 1.5 |
| | | 10 | 5 | 12.8 | 1.9 |
| | | 30 | 5 | 14.2 | 2.3 |

[1]Post-hoc pairwise comparisons were performed using Tukey's test (for ANOVAs) or Dunnett's T3 multiple comparisons test (for Welsh ANOVAs). *$p < 0.05$, **$p < 0.01$, ***$p < 0.001$, ****$p < 0.0001$, significant difference vs. vehicle control group.

**Supplementary Table 17. Head-Twitch Response (HTR) Data for Selected Phenethylamine Psychedelics.**

| Compound | ANOVA | Dose (mg/kg) | *N* | Mean | SEM | Post-hoc result[1] |
|---|---|---|---|---|---|---|
| **Desoxy** | $F_{4,21} = 26.31$, $p < 0.0001$ | 0 | 6 | 6.2 | 0.7 | |
| | | 0.3 | 5 | 15.4 | 2.5 | |
| | | 1 | 5 | 35.6 | 6.1 | |
| | | 3 | 5 | 107.6 | 10.1 | **** |
| | | 10 | 5 | 54.6 | 13.9 | ** |
| **Cyclopropylmescaline** | $F_{4,20} = 8.33$, $p = 0.0004$ | 0 | 5 | 8.0 | 1.9 | |
| | | 1.5 | 5 | 10.6 | 1.0 | |
| | | 3 | 5 | 25.2 | 2.5 | |
| | | 6 | 5 | 44.6 | 7.0 | ** |
| | | 12 | 5 | 35.4 | 9.4 | * |
| **25C-NBMD** | $F_{4,19} = 13.09$, $p < 0.0001$ | 0 | 5 | 4.2 | 1.3 | |
| | | 0.3 | 4 | 10.3 | 1.7 | |
| | | 1 | 5 | 31.0 | 4.9 | ** |
| | | 3 | 5 | 39.8 | 5.8 | **** |
| | | 10 | 5 | 28.6 | 4.2 | ** |
| **25N** | $W_{5,12.18} = 26.34$, $p < 0.0001$ | 0 | 6 | 11.3 | 2.2 | |
| | | 1.5 | 6 | 18.5 | 1.0 | |
| | | 3 | 6 | 28.3 | 2.2 | ** |
| | | 6 | 5 | 42.4 | 4.9 | ** |
| | | 12 | 7 | 57.4 | 3.6 | **** |
| | | 24 | 5 | 32.2 | 7.7 | |
| **25C-NBOH** | $F_{4,20} = 26.25$, $p < 0.0001$ | 0 | 5 | 10.6 | 1.5 | |
| | | 0.03 | 5 | 22.0 | 4.2 | |
| | | 0.3 | 5 | 84.6 | 9.7 | **** |
| | | 1 | 5 | 94.8 | 8.1 | **** |
| | | 3 | 5 | 69.2 | 9.6 | *** |
| **(±)-DOI** | $W_{4,8.34} = 51.87$, $p < 0.0001$ | 0 | 5 | 6.2 | 1.2 | |
| | | 0.1 | 5 | 37.4 | 7.5 | * |
| | | 0.3 | 5 | 80.6 | 6.2 | *** |
| | | 1 | 5 | 117.0 | 12.0 | ** |
| | | 3 | 5 | 63.6 | 10.0 | * |

[1]Post-hoc pairwise comparisons were performed using Tukey's test (for ANOVAs) or Dunnett's T3 multiple comparisons test (for Welsh ANOVAs). *$p < 0.05$, **$p < 0.01$, ***$p < 0.001$, ****$p < 0.0001$, significant difference vs. vehicle control group.

**Supplementary Table 18. Sources of Head-Twitch Response (HTR) Data for Phenethylamine Psychedelics.**

| Drug | HTR Potency (-pED$_{50}$ in moles/kg) | HTR Magnitude (counts per minute) | Reference |
|---|---|---|---|
| **Bromo-Dragonfly** | 6.70 | 4.150 | (Halberstadt et al. 2019b) |
| **2C-B** | 5.61 | 2.640 | (Halberstadt et al. 2019b) |
| **2C-B-FLY** | 5.75 | 2.507 | (Halberstadt et al. 2019b) |
| **3C-B-FLY** | 6.17 | 5.340 | (Halberstadt et al. 2019b) |
| **2C-I** | 5.62 | 3.127 | (Halberstadt and Geyer 2014) |
| **2C-I-FLY** | 5.29 | 2.427 | (Halberstadt et al. 2019b) |
| **25N** | 4.82 | 1.913 | Supplementary Table 17 |
| **25C-NBMD** | 5.78 | 1.327 | Supplementary Table 17 |
| **25C-NBOH** | 6.51 | 3.160 | Supplementary Table 17 |
| **25CN-NBOH** | 5.84 | 3.450 | (Halberstadt et al. 2019c) |
| **3C-P** | 5.07 | 1.900 | (Halberstadt et al. 2019a) |
| **2C-T-28** | 5.46 | 1.400 | (Halberstadt et al. 2023) |
| **Cyclopropylmescaline** | 4.96 | 1.487 | Supplementary Table 17 |
| **Desoxy** | 5.22 | 3.587 | Supplementary Table 17 |
| **DOB** | 6.12 | 3.960 | (Halberstadt et al. 2019b) |
| **DOBU** | 5.33 | 3.727 | (Halberstadt et al. 2020) |
| **DOET** | 6.11 | 4.067 | (Halberstadt et al. 2020) |
| **(±)-DOI** | 5.79 | 3.900 | Supplementary Table 17 |
| **α-Ethyl-2C-D (Ariadne)** | 4.80 | 0.867 | (Halberstadt et al. 2020) |
| **5H-DOB** | 5.59 | 1.378 | (Marcher-Rorsted et al. 2020) |
| **2H-DOB** | 5.65 | 1.427 | (Marcher-Rorsted et al. 2020) |
| **25I-NBOMe** | 6.77 | 3.420 | (Halberstadt and Geyer 2014) |
| **Mescaline** | 4.58 | 2.050 | (Halberstadt et al. 2019a) |
| **Proscaline** | 5.09 | 2.940 | (Halberstadt et al. 2019a) |

### Supplementary Table 19. Head-twitch response (HTR) Data Used to Test Whether Activity can be Predicted Based on 5-HT$_{2A}$ Gq Emax.

| Compound | ANOVA | Dose (mg/kg) | *N* | Mean | SEM | Post-hoc result[1] | $ED_{50}$ mg/kg (95% CI) |
|---|---|---|---|---|---|---|---|
| **25O-NBcP (33)** | $F_{5,25}$ = 20.57, $p < 0.0001$ | 0 | 6 | 8.8 | 2.2 | | 2.1 (1.5-2.9) |
| | | 1 | 5 | 29.3 | 2.0 | | |
| | | 3 | 5 | 59.0 | 7.8 | *** | |
| | | 10 | 5 | 92.2 | 6.4 | **** | |
| | | 30 | 5 | 53.8 | 12.6 | *** | |
| | | 100 | 5 | 20.6 | 5.1 | | |
| **2C2-NBOMe (31)** | $W_{6,12.67}$ = 96.28, $p < 0.0001$ | 0 | 7 | 8.4 | 1.4 | | 0.25 (0.18-0.35) |
| | | 0.03 | 5 | 10.2 | 1.3 | | |
| | | 0.1 | 5 | 23.2 | 5.0 | | |
| | | 0.3 | 6 | 53.0 | 5.2 | ** | |
| | | 1 | 5 | 89.2 | 3.0 | **** | |
| | | 3 | 7 | 74.4 | 10.0 | ** | |
| | | 10 | 4 | 44.0 | 9.5 | | |
| **25O-NBOMe (32)** | $F_{5,26}$ = 37.01, $p < 0.0001$ | 0 | 6 | 4.7 | 1.1 | | 0.70 (0.52-0.93) |
| | | 0.1 | 5 | 12.6 | 2.1 | | |
| | | 0.3 | 5 | 24.8 | 3.1 | | |
| | | 1 | 5 | 59.4 | 5.5 | **** | |
| | | 3 | 6 | 96.0 | 4.8 | **** | |
| | | 10 | 5 | 60.4 | 12.8 | **** | |
| **25O-NBPh-10'-OH (35)** | $F_{4,27}$ = 2.58, $p = 0.0601$ | 0 | 7 | 12.4 | 2.4 | | |
| | | 1 | 6 | 10.5 | 2.2 | | |
| | | 3 | 6 | 24.3 | 6.5 | | |
| | | 10 | 7 | 20.7 | 5.3 | | |
| | | 30 | 6 | 26.5 | 4.0 | | |
| **25O-NB-3-I (34)** | $F_{5,25}$ = 0.26, $p = 0.9288$ | 0 | 5 | 12.6 | 2.9 | | |
| | | 0.3 | 5 | 9.2 | 1.6 | | |
| | | 1 | 5 | 12.4 | 2.0 | | |
| | | 3 | 5 | 12.4 | 4.4 | | |
| | | 10 | 6 | 11.2 | 2.8 | | |
| | | 30 | 5 | 13.4 | 2.5 | | |
| **25D-N1-Nap (26)** | $W_{3,10.5}$ = 0.85, $p < 0.4942$ | 0 | 6 | 9.5 | 1.4 | | |
| | | 3 | 6 | 11.7 | 3.5 | | |
| | | 10 | 6 | 13.0 | 1.7 | | |
| | | 30 | 6 | 13.0 | 3.5 | | |

[1]Post-hoc pairwise comparisons were performed using Tukey's test (for ANOVAs) or Dunnett's T3 multiple comparisons test (for Welsh ANOVAs). *$p < 0.05$, **$p < 0.01$, ***$p < 0.001$, ****$p < 0.0001$, significant difference vs. vehicle control group.

**Supplementary Table 20. Summary of Induced-Fit Docking Results for Selection of 25N Compounds Docked Against Agonist Receptor Structure 6WHA.** Total number of poses and shown for major cluster, and interaction counts (over all poses in the cluster) between ligands and key binding site residues. (Some additional interactions noted.) Color coding: Red, 90% or more of poses include feature; orange at least 50%; yellow, at least 25%

| Compound | Num Poses | Moiety | Phe339 p-p | Phe340 p-p | Trp336 p-p | Ser159 H-bond |
|---|---|---|---|---|---|---|
| 25CN-NBOH (1) | 27 | cation | | | | 26 |
| | | 25CN | 7 | 20 | | |
| | | NBOH | 26 | 27 | 27 | 26 |
| 25N-NBOH (3) | 25 | cation | | | | 18 |
| | | 25N | 10 | 22 | | 6 |
| | | NBOH | 23 | 24 | 25 | 22 |
| 25N-NBOMe (4) | 22 | cation | | | | 4 |
| | | 25N | 20 | 16 | | 8 |
| | | NBOMe | 21 | 21 | 22 | 10 |
| 25N-NBI (10) | 12 | cation | | | | 5 |
| | | 25N | 8 | 8 | | 4 |
| | | NBI | 12 | 11 | 11 | 3 |
| 25N-N1-Nap (16) | 15 | cation | | | | 10 |
| | | 25N | 11 | 8 | | 7 |
| | | Naph | 15 | 13 | 15 | |
| 25N-NBPh (17) | 8 | cation | | | | 1 |
| | | 25N | 2 | 2 | | |
| | | NB | 8 | 8 | 7 | |
| | | phenyl | | | 6 | |
| 25N-NB-2-OH-3-Me (18) | 20 | cation | | | | 9 |
| | | 25N | 10 | 13 | | 3 |
| | | NB-2-OH-3-Me | 19 | 17 | 20 | 17 |
| 25N-NB-2,5-DiMeO (20) | 7 | cation | | | | 4 |
| | | 25N | 6 | 5 | | 4 |
| | | NB-2,5-DiMeO | 5 | 5 | 7 | 1 |

**Supplementary Table 21. Averaged W336$^{6.48}$ $\chi^2$ angle for 25CN-NBOH and 25N-N1-Nap (16) MD Simulations.**

| Trajectory | W366$^{6.48}$ $\chi^2$ angle | | |
|---|---|---|---|
| | **Peak Position** | **Peak Range** | **Width ½ Max** |
| 25CN-NBOH | 80° | 19-125° | 29 |
| 25N-N1-Nap (16) | 112° | 29-155° | 25 |

**Supplementary Table 22. W336$^{6.48}$ $\chi^2$ angles of published 5-$HT_{2A}$ structures.**

| **PDB Entry** (Ligand) | | **6WHA** (25CN-NBOH) | **6WH4** (Methiothepin) | **6WGT** (LSD) | **6A93** (Risperidone) | **6A94** (Zotepine) |
|---|---|---|---|---|---|---|
| | **Chain** | | | | | |
| W336$^{6.48}$ $\chi^2$ | A | 71.8 | 110.3 | 119.1 | 119.5 | 106.9 |
| | B | - | 107.8 | 105.4 | 119.7 | 104.9 |
| | C | - | 95.9 | 90.5 | | |
| Circular Mean | | 71.8 | 104.7 | 105.0 | 119.6 | 105.9 |
| Circular Stdev | | - | 0.4 | 0.4 | 0.7 | 0.4 |

**Supplementary Table 23. Parameters in common between 25CN-NBOH and 25N-N1-Nap (16) MD simulations**

| III. Parmeter | value |
|---|---|
| MD iterator | leap-frog |
| Constraints (h-bonds) | LINCS[24] |
| LINCS iter | 1 |
| LINCS order | 4 |
| Water Constraints | SETTLE[25] |
| Coulomb Type | PME[26] |
| PME order | 4 |
| Fourier Spacing | 0.16 |
| Coulomb Cutoff (Å) | 12 |
| vdW Cutoff | 12 |
| Neighbor list cutoff | 12 |
| Neighbor search type | grid |
| Neighbor list update (fs) | 5 |
| dt (fs) | 1.25 |
| Temperature (K) | 323 |
| Pressure (bar) | 1 |
| P-couple type | semi-isotropic |
| Compressibility ($bar^{-1}$) | 4.50E-05 |
| Dispersion correction | Energy and Pressure |

**Supplementary Table 24. Parameters that vary between 25CN-NBOH and 25N-N1-Nap (16) MD simulations**

| Ligand (Set) | Phase | Simulation | *k* (kJ / mol / nm) | Duration (ns) | Thermostat | $\tau_T$ (ps) | Barostat | $\tau_P$ (ps) |
|---|---|---|---|---|---|---|---|---|
| 25CN-NBOH | Minimization | Steepest Descent | N/A | N/A | N/A | N/A | N/A | N/A |
| | Equilibration | NVT | 1000 | 1 | V-rescale | 0.1 | N/A | N/A |
| | | NPT | 1000 | 10 | Nosé-Hoover | 0.5 | Berendsen | 5 |
| | Production | NPT | N/A | 250 | Nosé-Hoover | 0.5 | Parrinello-Rahman | 2 |
| 25N-N1-Nap (16) | Minimization | Steepest Descent | 1000 | N/A | N/A | N/A | N/A | N/A |
| | | Steepest Descent | N/A | N/A | N/A | N/A | N/A | N/A |
| | Equilibration | NVT | 1000 | 1 | V-rescale | 0.1 | N/A | N/A |
| | | NPT | 1000 | 10 | Nosé-Hoover | 0.5 | Berendsen | 5 |
| | Production | NPT | N/A | 250 | Nosé-Hoover | 0.5 | Parrinello-Rahman | 2 |

**Synthesis Schemes**

**Scheme 1. Synthetic Scheme Used for 25N Series (1-25).**

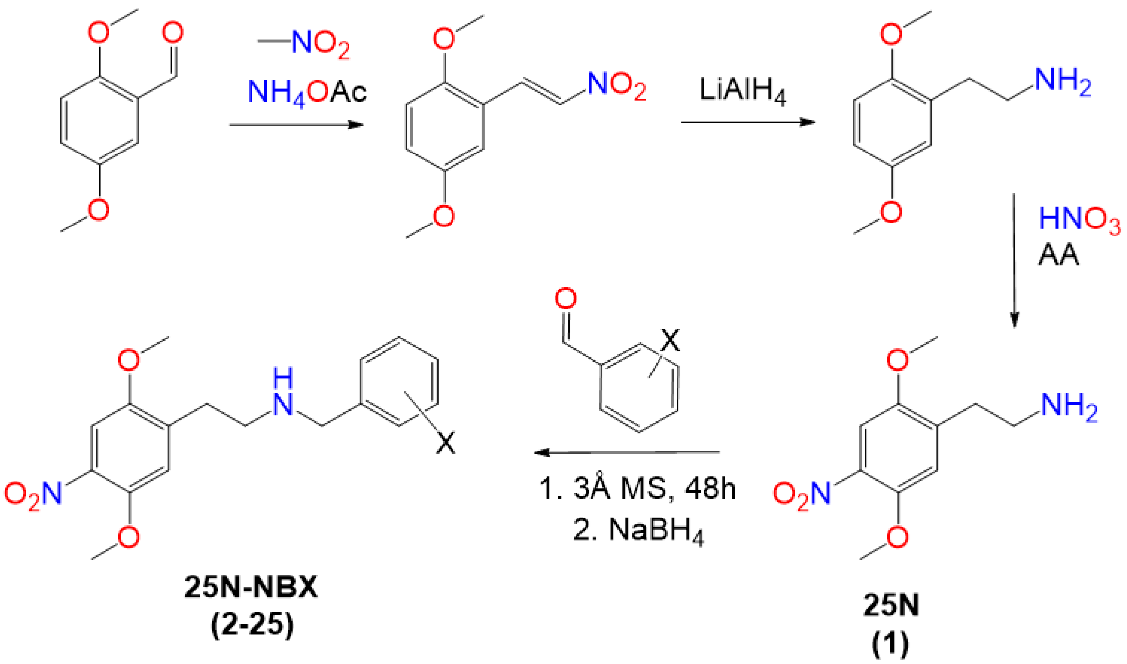

**Scheme 2. Synthetic Scheme Used for 25D, 25D-N1-Nap (26) and 25D-NBPh (27).**

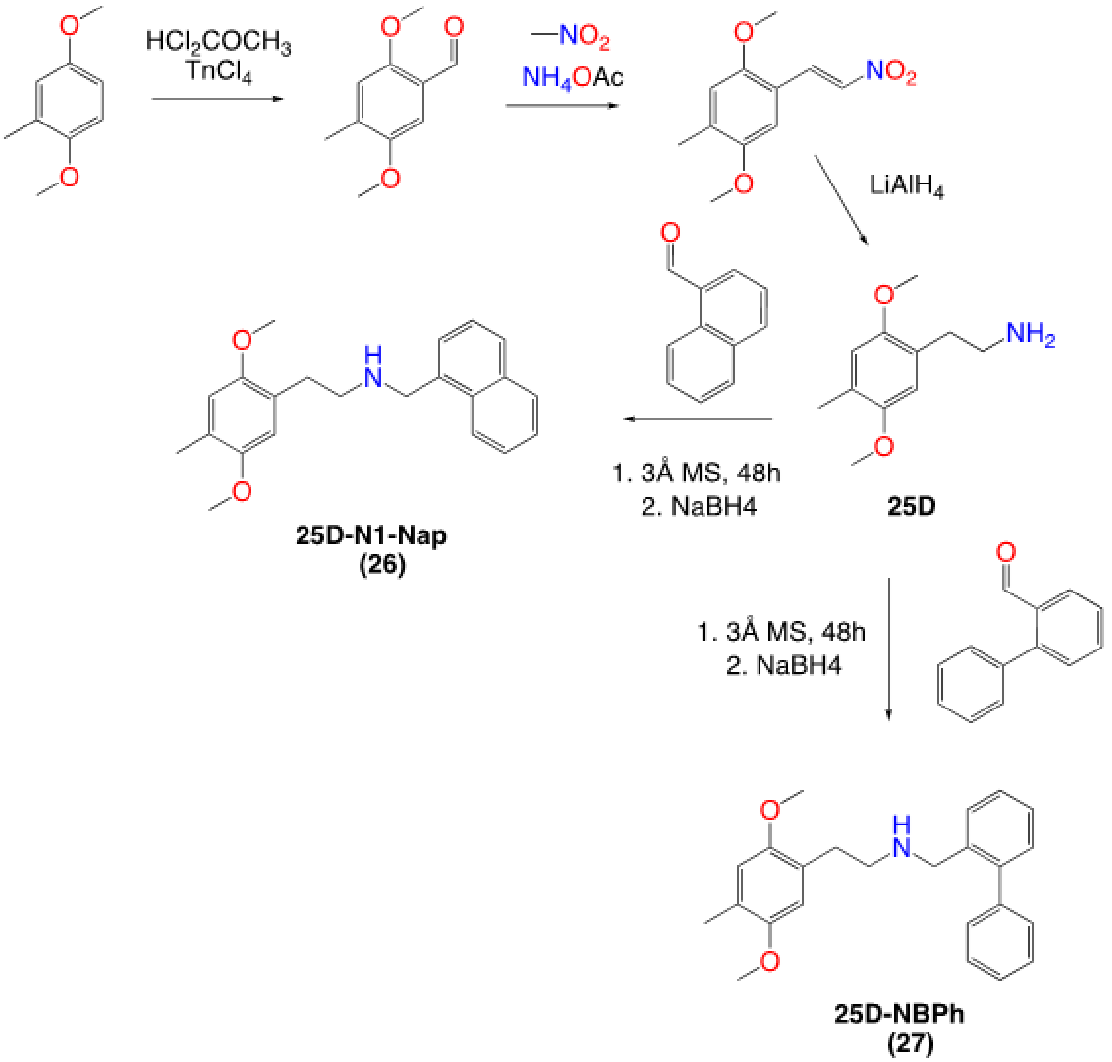

**Scheme 3. Synthetic Scheme Used for 25O and 25O-N1-Nap (28).**

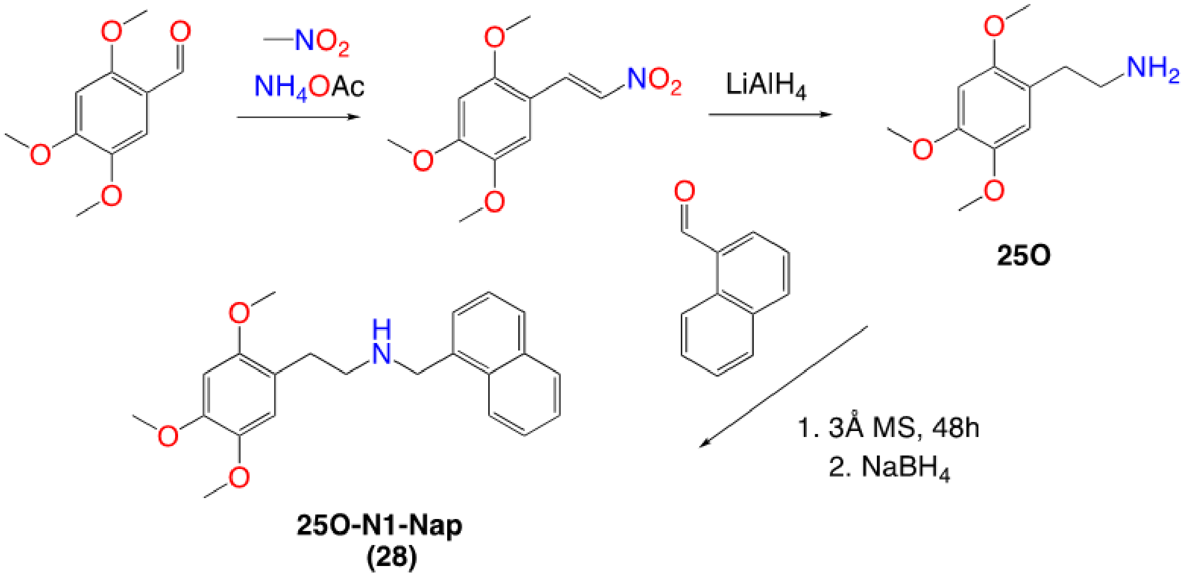

**Scheme 4.** **Synthetic Scheme Used for 2C-2 and 2C2-N1-Nap (29).**

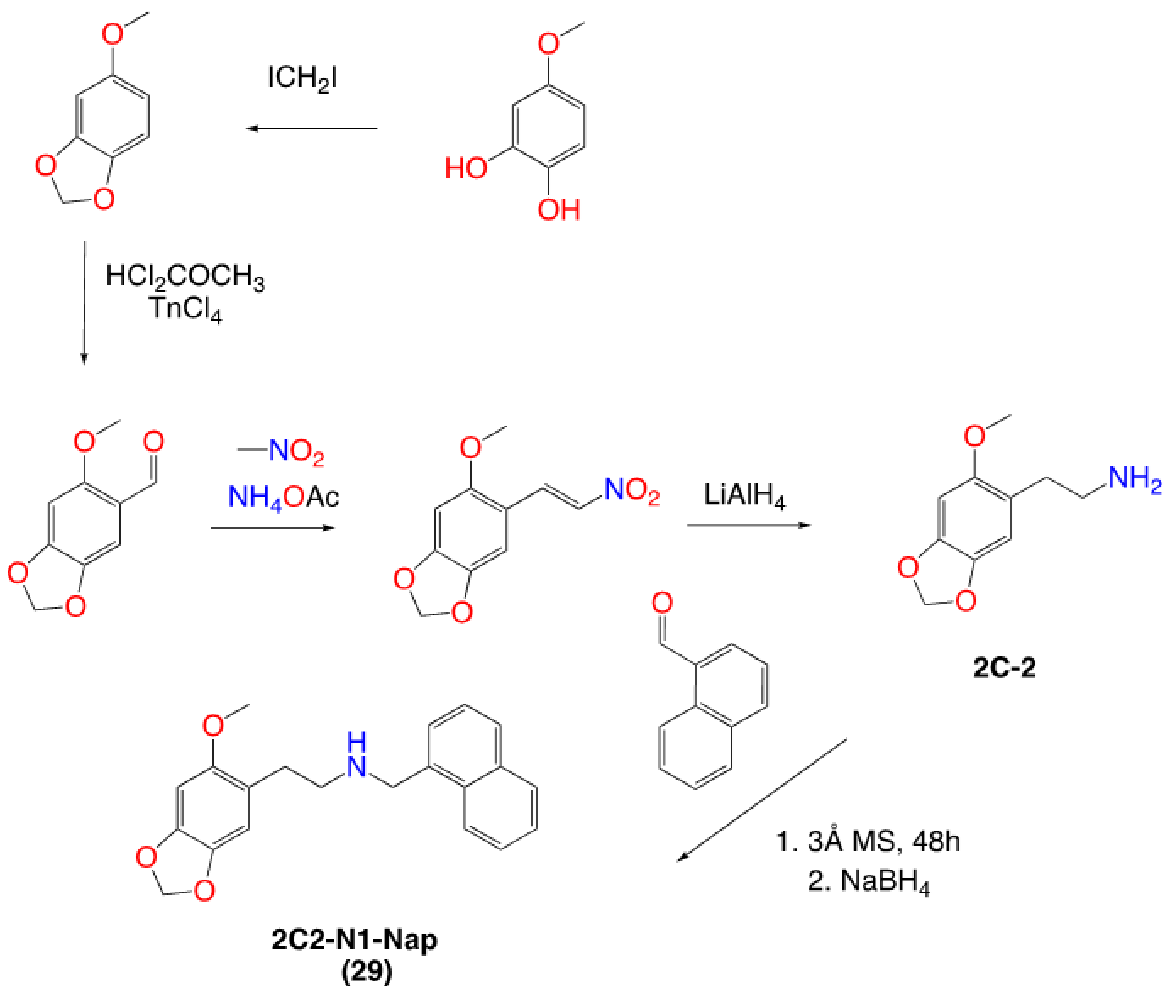

### Supplementary Methods

#### Instrumentation:

**Nuclear Magnetic Resonance.** $^{1}H$ and $^{13}C$ NMR spectra data were obtained on a Bruker Avance III with PA BBO 400S1 BBF-H-D-05 Z plus probe (Bruker Corporation, Billerica, MA, USA). Samples were prepared at a concentration of ~20 mg/mL in anhydrous DMSO-$d_6$. Chemical shifts are reported in parts per million (ppm) against the solvent signal (DMSO-$d_6$ $^{1}H$ = 2.50 ppm, $^{13}C$ = 39.52 ppm, fluorotrichloromethane ($CFCl_3$), $^{19}F$ = 0.00). Assignments based on 1D relative chemical shift positions, $^{1}H$ chemical shift multiplicities, PENDANT $^{13}C$ experiments, and 2-D homo (COSY) and heteronuclear (HMQC, HSQC, HMBC) experiments. For the 25N compounds blank spectra, side by side experiments were run on the anhydrous d6-DMSO solvent (which still contains some water) to rule out the presence of hydrates (no evidence of hydrates was observed). Full NMR chemical shift assignments for $^{1}H$, $^{13}C$, and $^{19}F$ signals are presented in table format in the supplementary information document.

**Atmospheric Solids Analysis Probe Mass Spectroscopy (ASAP-MS).** Low resolution mass spectra for reaction monitoring were obtained on an Advion Expression$^{s}$ CMS Spectrometer with a quadrupole mass analyzer. Samples were ionized via Atmospheric Solids Analysis source using an Atmospheric Pressure Chemical Ionization (APCI) attachment. Data was processed in Advion Data Express software. Measurement parameters were as follows: Capillary Temperature = 150 °C, Capillary Voltage = 120 V, Source Gas Temperature = 200 °C, and APCI corona discharge = 5 µA.

**High resolution mass spectral analysis (HRMS)**. HRMS data were obtained on a Thermo Orbitrap Exactive Mass Spectrometer with an Orbitrap mass analyzer. The instrument was calibrated using electrospray ionization with Pierce™ LTQ ESI Positive Ion Calibration Solution from ThermoFisher Scientific. Samples were introduced into the instrument and ionized via an Atmospheric Solids Analysis Probe (ASAP). Parameters: Spray voltage- 3.50 V; Capillary temperature-275 ℃; Capillary voltage-25.00 V; Tube lens voltage- 65.00 V; Skimmer voltage-14.00 V; Heater temperature-100 ℃. Data was analyzed in the Thermo Xcalibur Qual Browser software and identity was confirmed if <5 ppm error.

**High Performance Liquid Chromatography (HPLC)**

HPLC analyses were performed on an Agilent 1260 Infinity system that includes a 1260 quaternary pump VL, a 1260 ALS autosampler, a 1260 Thermostatted Column Compartment, and a DAD Multiple Wavelength Detector (Agilent Technologies, Santa Clara, CA, USA). The detection wavelengths were set at 220, 230, 254, and 280 nm. Separation was achieved using a Zorbax Eclipse XDB-C18 analytical column (5 µm, 4.6 x 150 mm) from Agilent (Agilent Technologies, Santa Clara, CA, USA). Mobile phase A consisted of 10 mM aqueous ammonium formate buffer titrated to pH 4.5 and mobile phase B consisted of acetonitrile. The injection volume of samples was 10 µL, flow rate was 1.0 mL/min, and the column temperature was set at 25ºC. Samples were prepared by weighing analyte into a vial and making a 1 mg/mL solution in 1:1 A:B. All samples were injected in duplicate with a wash in between each run. Run time was 10 minutes with a mobile phase ratio (isocratic) of 1:1 for A:B. Chromatograms were analyzed using the Agilent ChemStation Software (Agilent Technologies, Santa Clara, CA, USA).

**Elemental Analysis**

Elemental analysis (C, H, N) was run on select compounds by Galbraith Laboratories, Inc. (Knoxville, TN).

#### Syntheses:

**1,4-dimethoxy-2-[(1E)-2-nitroethenyl]benzene**

0.0722 mol (12.0 g) 2,5-dimethoxybenzaldehyde was dissolved in 25 mL nitromethane containing 0.0157 mol (1.21 g) ammonium acetate. The reaction was heated at 80 °C on a water bath for 6 hours. After which the solution rapidly set to a solid orange cake. This was allowed to sit at room temperature overnight, the solids were then collected by gravity filtration, dissolved in 50 mL boiling isopropanol and allowed to sit at room temperature for several days. The deep orange crystals were collected by vacuum filtration and dried to give 0.0529 mol (11.6 g) 1,4-dimethoxy-2-[(1E)-2-nitroethenyl]benzene as orange crystalline needles. An additional 0.4 g was obtained as a second crop (after recrystallization from 10 mL 200 proof ethanol). Total of 12.0 g (79.5 % yield).

**2-(2,5-dimethoxyphenyl)ethan-1-amine (2C-H)**

0.0554 mol (11.6 g) of 1,4-dimethoxy-2-[(1E)-2-nitroethenyl]benzene was dissolved in 150 mL anhydrous THF and added slowly over 1 hour dropwise to stirred suspension of 0.166 mol (6.3 g) $LiAlH_4$ in 100 mL anhydrous THF on an ice-water bath under argon. After addition the grey solution was allowed to recover to room temperature at which point it was placed on a mild reflux. Progress was monitored by TLC and MS-ASAP. After 3 hours the reaction was finished. The reaction was placed on an ice-water bath and the excess hydride was quenched by the slow (~30 minutes) dropwise addition of $H_2O$:THF (3:1). A few mL of an aqueous KOH solution was added and the solution was then diluted with 200 mL ethyl acetate and inorganics removed by gravity filtration. The solids were washed heavily with ethyl acetate (~200 mL). The resulting ethyl acetate solution was extracted with aqueous 1N HCl (3 x 150 mL). The pooled aqueous solutions were then made basic with the addition of KOH pellets. The resulting cloudy solution was then extracted with ethyl acetate (3 x 100 mL), each extraction washed with 10 mL brine and then pooled and dried with anhydrous $Na_2SO_4$. The solvent was removed under vacuum to give an amber oil. This crude freebase was immediately distilled using a Kugelrohr (170-200 °C) to give 5.2 g of 2C-H as a colorless oil (51.8 % yield). This oil set to a white solid upon storage at -20 °C under argon.

**2-(2,5-dimethoxy-4-nitrophenyl)ethan-1-amine (25N, 2C-N) (1)**

2C-N (1) was synthesized using a modification of the method described by Shulgin and Shulgin [Shulgin and Shulgin. 1991]. 0.027589 mol (5.0 g) 2,5-dimethoxyphenethylamine freebase was dissolved in 50 mL glacial acetic acid and placed on an ice bath while vigorously stirring. 16.5 mL of 70% nitric acid was added dropwise over several minutes. The initially clear solution turned yellow upon addition of the nitric acid. Stirring on ice was continued and after 12 minutes a spatula was used to scratch the side of the flask resulting in the precipitation of a small amount of yellow crystals. The solution then set to a yellow crystalline mass over 1 minute. This was stirred for an additional 20 minutes, at which point 75 mL of diethyl ether ($Et_2O$) was slowly added. The resulting light-yellow crystals were collected onto Whatman paper by vacuum filtration, washed with additional $Et_2O$ and dried at room temperature to give 6.66 g of fluffy canary yellow crystals. An additional 0.64 g of material (sparkling darker yellow crystals) was collected as a slower precipitate from the combined filtrate and washes. Total yield, 7.3 g (91.43% yield) of 2C-N (1) nitrate. This material was dissolved in water, basified with excess KOH pellets and extracted with ethyl acetate (3 x 75 mL), pooled, washed with brine, dried over anhydrous $Na_2SO_4$ and concentrated under vacuum to give 2C-N (1) freebase as a yellow-orange waxy solid that set to single solid mass in near quantitative yield from the nitrate salt. The HCl salt was prepared by dissolving the freebase in 20 mL ethanol (200 proof), which was titrated to an acidic pH (pH <3) with concentrated HCl while stirring. The solvent was then evaporated under warm air flow. Additional EtOH was added and evaporation repeated until all excess water and acid was gone (~4 x 10 mL volumes of EtOH). This resulted in light yellow powder which was washed with $Et_2O$ (10 mL) and dried with gentle heating. The resulting solids were recrystallized by dissolving in ~5 mL boiling EtOH followed by the addition of ~20 mL $Et_2O$ and storing at room temperature (~1 hour) followed by -20 °C overnight. The resulting crystals were then washed with $Et_2O$ (2 x 10 mL) followed by ethyl acetate (5 mL). This was repeated for a total of three crystallizations to give light yellow crystalline solids of 2C-N (25N) (1) HCl that were then dried in a vacuum desiccator for ~48 hours, mp: 201.0-202.3 °C (Lit: 193-195 °C [Shulgin and Shulgin 1991]. HRMS: Observed: 227.1015 (100), Theoretical: $C_{10}H_{15}N_2O_4$: 227.1026, Δppm: -4.84.

***N*-benzyl-2-(2,5-dimethoxy-4-nitrophenyl)ethan-1-amine(25N-NB) (2)**

0.00088 mol (200 mg) 2C-N (1) freebase and 0.001056 mol (112 mg) benzaldehyde were dissolved in 10 mL dry (3Å molecular sieves) were dissolved in 10 mL of methanol and 2 mL anhydrous THF containing ~1 g 3Å molecular sieves. The reaction was sealed under argon and protected from light and left for 4 days. After which, the reaction was placed on an ice-water bath and under argon flow, at which point 0.0044 mol (166 mg) $NaBH_4$ was added in small portions over ~10 minutes with vigorous mixing. The reaction was mixed occasionally for an additional hour at which point it was removed from the ice-water bath and active argon flow (but sealed under argon) and left to sit (with occasional mixing) for 4 hours. The reaction was then quenched by slow addition of the solution to 300 mL 2N aqueous HCl solution. The solution was washed with ethyl acetate (2 x 60 mL). Organic washes were pooled and extracted twice with 2N aqueous HCl (3 x 60 mL). The acidic aqueous phases were pooled, made basic with KOH pellets and extracted with ethyl acetate (3 x 60 mL). Organic extracts were washed with brine (10 mL), pooled, dried over anhydrous magnesium sulfate and evaporated under vacuum to give a yellow oil. This crude freebase was purified via flash column chromatography on silica gel with hexanes:ethyl acetate (3:2) containing 1% triethylamine. The ethyl acetate was slowly increased to 50%. Pure fractions were identified using MS-ASAP and TLC and combined to give 140 mg (50.4 % yield) of a light-yellow oil. A straight to base work up was later observed to give substantially higher yields on *N*-benzyl-phenethylamines and its likely product is lost in the organic washes with the acid base workup. The HCl salt was prepared as described for 2C-N (1) to give a beige-tan crystalline powder (mp: 220.0-221.0 °C). HRMS: Observed: 317.1480 (100), Theoretical: $C_{17}H_{21}N_2O_4$: 317.1496 (100), Δppm: -0.22.

**2-(((2,5-dimethoxy-4-nitrophenethyl)amino)methyl)phenol (25N-NBOH) (3)**

Prepared as described for 25N-NB (2) using 0.00088 mol (200 mg) 2C-N (1) and 0.00132mol (185 μL) 2-hydroxybenzaldehyde to give an amber oil (purified by column chromatography). The HCl salt was prepared as described for 2C-N (1) HCl to give 203.3 mg (62.6% yield) light yellow crystalline solids (mp: 205-206.7 °C). HRMS: Observed: 333.1443 (100), Theoretical: $C_{17}H_{21}N_2O_5$: 333.1445, Δppm: -0.60. Elemental Analysis: Calc: C, 55.36; H, 5.74; N, 7.6. Found: C, 54.92; H, 5.81; N, 7.38

**2-(2,5-dimethoxy-4-nitrophenyl)-*N*-(2-methoxybenzyl)ethan-1-amine (25N-NBOMe) (4)**

Prepared as described for 25N-NB (2) using 0.0088 mol (2 g) 2C-N (1) and 0.01056mol (1.44 g) 2-methoxybenzaldehyde to give 1.95 g (64% yield) of a dark yellow oil (after flash column chromatography). The HCl salt was prepared as described for 2C-N (1) HCl to give large sparkling transparent yellow needles (mp: 166.4-167.8 °C). HRMS: Observed 347.1599 (100), Theoretical: $C_{18}H_{23}N_2O_5$: 347.1602, Δppm: -0.86. Elemental analysis: Calc: C, 56.47; N, 6.06; N, 7.32. Found: C, 56.17; H, 5.87; N, 7.43.

**2-(2,5-dimethoxy-4-nitrophenyl)-*N*-(2-ethoxybenzyl)ethan-1-amine (25N-NBOEt) (5)**

Prepared as described for 25N-NB (2) using 0.00088 mol (200 mg) 2C-N (1) and 0.00132mol (185 μL) 2-ethoxybenzaldehyde to give a yellow solid which was purified by crystallization from ethyl acetate and hexanes at 0 °C to give 185 mg (58.4% yield) of yellow crystals. Of note an additional 39 mg of the HCl salt was recovered from the initial organic washes. Later in the project it was found that a straight to base workup as described for 25O-N1-Nap (28) improved recoveries. The HCl salt was prepared as described for 2C-N (1) HCl to give yellow needles (mp: 202.5-203.5 °C). HRMS: Observed: 361.1759 (100), Theoretical: $C_{19}H_{25}N_2O_5$: 361.1758, Δppm: 0.28.

**2-(2,5-dimethoxy-4-nitrophenyl)-*N*-(2-methylbenzyl)ethan-1-amine (25N-NBMe) (6)**

Prepared as described for 25N-NB (2) using 0.00088 mol (200 mg) 2C-N (1) and 0.001056 mol (122 μL) o-tolualdehyde to give a yellow oil which was purified by crystallization (2X) from ethyl acetate and hexanes at 0 °C to give 170 mg (58.4 % yield) of transparent yellow crystalline clusters. The HCl salt was prepared as described for 2C-N (1) HCl to give a fluffy beige yellow crystalline solid (mp: 205-205.5 °C). HRMS: Observed 331.1648 (100), Theoretical: $C_{18}H_{23}N_2O_4$: 331.1652, Δppm: -1.21. Elemental Analysis: Calc: C, 58.93 6.32; N, 6.99. Found: C, 58.62; H, 6.26; N: 6.90.

**2-(2,5-dimethoxy-4-nitrophenyl)-N-(2-fluorobenzyl)ethan-1-amine (25N-NBF) (7)**

Prepared as described for 25N-NB (2) using 0.00088 mol (200 mg) 2C-N (1) and 0.00132 mol (139.1 μL) 2-fluorobenzaldehyde to give a light yellow oil (after flash column chromatography). The HCl salt was prepared as described for 2C-N (1) HCl to give 126 mg (38.7% yield) of light-yellow crystalline solids (mp: 186.7-187.0 °C). HRMS: Observed: 335.1395 (100), Theoretical: $C_{17}H_{19}FN_2O_4$+H:

335.1402, Δppm: -2.09. Elemental Analysis: $C_{17}H_{20}ClFN_2O_4$-0.2$H_2O$. Calc: C, 54.53; H, 5.49; N, 7.48. Found: C, 54.97; H, 5.32; N, 7.06.

***N*-(2-chlorobenzyl)-2-(2,5-dimethoxy-4-nitrophenyl)ethan-1-amine (25N-NBCl) (8)**

Prepared as described for 25N-NB (2) using 0.00088 mol (200 mg) 2C-N (1) and 0.001056 mol (119 μL) 2-chlorobenzaldehyde to give a solid which was purified by crystallization (2X) from ethyl acetate and hexanes at 0 C to give 185 mg (59.9% yield) of yellow crystalline solids. The HCl salt was prepared as described for 2C-N (1) HCl to give a fluffy bright yellow crystalline solid (mp: 194-195.6 °C). HRMS: Observed: 351.1100 (100), 352.113 (20), 353.1071 (30), Theoretical: $C_{17}H_{20}ClN_2O_4$: 351.1106, Δppm: -1.71. Elemental Analysis: $C_{17}H_{20}Cl_2N_2O_4$, Calc: C, 52.73; N, 5.21; N, 7.23. Found: C, 52.78; H, 5.19; N, 7.0.

***N*-(2-bromobenzyl)-2-(2,5-dimethoxy-4-nitrophenyl)ethan-1-amine (25N-NBBr) (9)**

Prepared as described for 25N-NB (2) using 0.00088 mol (200 mg) 2C-N (1) and 0.001056 mol (123 μL) 2-bromobenzaldehyde to give 166 mg (47.7 % yield) of a transparent yellow oil (after flash column chromatography). The HCl salt was prepared as described for 2C-N (1) HCl to give fluffy bright yellow crystalline solids (mp: 210.6-211.0 °C). HRMS: Observed: 397.0573 (100), 395.0594 (100), Theoretical: $C_{17}H_{20}BrN_2O_4$: 397.0580 (100), Δppm: -1.76, 395.0601 (100), Δppm: -1.77.

**2-(2,5-dimethoxy-4-nitrophenyl)-*N*-(2-iodobenzyl)ethan-1-amine (25N-NBI) (10)**

Prepared as described for 25N-NB (2) using 0.00088 mol (200 mg) 2C-N (1) and 0.001056 mol (245 mg) 2-iodobenzaldehyde to give 110 mg (28.3 % yield) of an orange oil which solidified upon storage at 0 °C. The HCl salt was prepared as described for 2C-N (1) HCl to give a fluffy light yellow crystalline solid (mp: 226.7-228.0 °C). An additional 120 mg of HCl salt as an orange crystalline solid was obtained from the evaporated organic washes, from the work up and purified by recrystallization three times (mp: 227.5-228.7 °C with decomposition). 56.6% combined yield. HRMS: Observed: 443.0458 (100), Theoretical: $C_{17}H_{20}IN_2O_4$: 443.0462, Δppm: -0.902.

***N*-(2-(difluoromethoxy)benzyl)-2-(2,5-dimethoxy-4-nitrophenyl)ethan-1-amine (25N-NBOCF$_2$H) (11)**

Prepared as described for 25N-NB (2) using 0.00088 mol (200 mg) 2C-N (1) and 0.00088 mol (151 mg) 2-dlfluoromethoxy-benzaldehyde to give a golden oil. The HCl salt was prepared as for 2C-N (1) HCl to give 65 mg (17.6% yield) canary yellow solids (mp: 197.5-199.0 °C). HRMS: Observed: 383.1408 (100), Theoretical $C_{18}H_{21}F_2N_2O_5$: 383.1413, Δppm: -1.31. Elemental Analysis: $C_{18}H_{21}ClF_2N_2O_5$-0.18$H_2O$, Calc: C, 51.22; H, 5.10; N, 6.63. Found: C, 50.85; H, 5.18; N, 6.41.

**2-(2,5-dimethoxy-4-nitrophenyl)-*N*-(2-(trifluoromethoxy)benzyl)ethan-1-amine (25N-NBOCF$_3$) (12)**

Prepared as described for 25N-NB (2) using 0.00176 mol (400 mg) 2C-N (1) and 0.002 mol (285 μL) 2-(trifluoromethoxy)-benzaldehyde to give a yellow oil. The HCl salt was prepared as described for 2C-N (1) HCl to give 300 mg (39.0% yield) yellow crystalline solids (mp: 163.0-164.3 °C). HRMS: Observed: 401.1314 (100), Theoretical: $C_{18}H_{20}F_3N_2O_5$: 401.1319, Δppm: -1.25.

***N*-((2,2-difluorobenzo[d][1,3]dioxol-4-yl)methyl)-2-(2,5-dimethoxy-4-nitrophenyl)ethan-1-amine (25N-NBMDF$_2$) (13)**

Prepared as described for 25N-NB (2) using 0.00088 mol (200 mg) 2C-N (1) and 0.00132 mol (208 mg) 2,2-difluoro-1,3-benzodioxole-4-carboxaldehyde to give a yellow oil (after flash column chromatography) that was crystallized (ethyl acetate:hexanes) to a yellow crystalline solid. The HCl salt was prepared as described for 2C-N (1) HCl to give 87 mg (24.9% yield) of beige crystalline solids (mp: 208.4-209.8 °C). HRMS: Observed: 397.1198 (100), Theoretical: $C_{18}H_{19}F_2N_2O_6$, 397.1206 (100), Δppm: -2.01.

**2-(2,5-dimethoxy-4-nitrophenyl)-*N*-(2-(trifluoromethyl)benzyl)ethan-1-amine (25N-NBCF$_3$) (14)**

Prepared as described for 25N-NB (2) using 0.00088 mol (200 mg) 2C-N (1) and 0.00132 mol (177 μL) 2-(trifluoromethyl)benzaldehyde to give a transparent yellow oil (after flash column chromatography). The HCl salt was prepared as described for 2C-N (1) HCl to give 166 mg (44.8% yield) of fluffy yellow needles (mp: 156.7-159.1 °C). HRMS: Observed: 385.1366 (100) Theoretical: $C_{18}H_{20}F_3N_2O_4$, 385.1370 (100), Δppm: -1.03.

**2-(2,5-dimethoxy-4-nitrophenyl)-*N*-(2-nitrobenzyl)ethan-1-amine 25N-NBNO$_2$) (15)**

Prepared as described for 25N-NB (2) using 0.00088 mol (200 mg) 2C-N (1) and 0.00158 mol (239 mg) 2-nitrobenzaldehyde to give a transparent golden oil which set to an opaque gold solid. The solids were purified by crystallization from ethyl acetate diluted with hexanes to give 195 mg (61.3% yield) of bright yellow salt-granule-like crystals. The HCl salt was prepared as described for 2C-N (1) HCl to give a yellow crystalline powder (mp: 200.5-201.7 °C). HRMS: Observed: 362.1346 (100), Theoretical: $C_{17}H_{20}N_3O_6$, 362.1347 (100), Δppm: -0.27.

**2-(2,5-dimethoxy-4-nitrophenyl)-*N*-(naphthalen-1-ylmethyl)ethan-1-amine (25N-N1-Nap) (16)**

Prepared as described for 25N-NB (2) using 0.00088 mol (200 mg) 2C-N (1) and 0.00132 mol (179 μL) 1-napthaldehyde to give a yellow oil which was purified by crystallization (2X) from ethyl acetate and hexanes at 0 °C to give 103 mg of yellow needles (32.0% yield). An additional 200 mg of a water soluble solid was recovered from the organic washes which was the HCl salt of the product. Total % yield: 83.0%. The HCl salt was prepared as described for 2C-N (1) HCl to give fluffy beige crystalline solid mp: 191.8-192.3 °C. HRMS: Observed: 367.1653 (100), Theoretical: $C_{21}H_{23}N_2O_4$, 367.1652 (100), Δppm: 0.27.

***N*-([1,1'-biphenyl]-2-ylmethyl)-2-(2,5-dimethoxy-4-nitrophenyl)ethan-1-amine amine (25N-NBPh) (17)**

Prepared as described for 25N-NB (2) using 0.00088 mol (200 mg) 2C-N (1) and 0.00132 mol (240.5 mg) biphenyl-2-carboxylate to give a yellow oil (after flash column chromatography). The HCl salt was prepared as described for 2C-N (1) HCl to give 104.3 mg (27.6% yield) of light yellow fluffy crystalline needles mp: 181.5-182.5 °C). HRMS: Observed: 393.1807 (100), Theoretical: $C_{23}H_{25}N_2O_4$: 393.1809 (100), Δppm: -0.51.

**2-(((2,5-dimethoxy-4-nitrophenethyl)amino)methyl)-6-methylphenol (25N-NB-2-OH-3-Me) (18)**

Prepared as described for 25N-NB (2) using 0.00088 mol (200 mg) 2C-N (1) and 0.00132 mol (179.7 mg) 2-hydroxy-3-methylbenzaldehyde to give a yellow oil. The HCl salt was prepared as for 2C-N (1) HCl to give 180 mg (53.4 % yield) transparent neon yellow crystalline solids (mp: 177.6-179.5 °C). HRMS: Observed, 347.1588, Theoretical: $C_{18}H_{23}N_2O_5$, 347.1601, Δppm: -3.744.

**2-(2,5-dimethoxy-4-nitrophenyl)-N-(3-fluoro-2-methoxybenzyl)ethan-1-amine (25N-NB-2-MeO-3-F) (19)**

Prepared as described for 25N-NB (2) using 0.00088 mol (200 mg) 2C-N (1) and 0.00088 mol (136 mg) 2-methoxy-3-dluorobenzaldehyde to give a golden oil. HCl salt prepared as for 2C-N (1) HCl to give 130 mg (36.9% yield) transparent yellow needles (mp: 155.0-156.7 °C). HRMS: Observed, 365.1500, Theoretical: $C_{18}H_{22}FN_2O_5$, 365.1507, Δppm: -1.91.

**3-(((2,5-dimethoxy-4-nitrophenethyl)amino)methyl)phenol (25N-NB-3-OH) (21)**

Prepared as described for 25N-NB (2) using 0.00088 mol (200 mg) 2C-N (1) and 0.001056 mol (129 mg) 3-hydroxybenzaldehyde to give a yellow oil which solidified to a yellow solid upon sitting. This was purified by crystallization (dissolve in boiling 10 mL ethyl acetate and 2 mL ethanol followed by dilution with 20 mL hexanes and storing at 0 °C) to give 195 mg transparent tan needles (66.8 % yield). The HCl salt prepared as described for 2C-N (1) HCl to give transparent orange flat edged rectangular crystals (mp: 194.0-195.5 °C). HRMS: Observed: 333.1443 (100), Theoretical: $C_{17}H_{21}N_2O_5$, 333.1445 (100), Δppm: -0.60.

**2-(2,5-dimethoxy-4-nitrophenyl)-*N*-(3-methylbenzyl)ethan-1-amine (25N-NB-3-Me) (22)**

Prepared as described for 25N-NB (2) using 0.000663 mol (150 mg) 2C-N (1) and 0.001326 mol (156 μL) p-tolualdehyde to give a yellow-orange solid. This was crystalized (2X) from ethyl acetate and hexanes stored at 0 °C to give 160 mg (73.1 % yield) of crystalline orange needle clusters. The HCl salt prepared as described for 2C-N (1) HCl to give a beige-yellow crystalline powder (mp: 194.4-195.5 °C). HRMS: Observed: 331.1645 (100), Theoretical: $C_{18}H_{23}N_2O_4$, 331.1652 (100), Δppm: -2.11.

**2-(2,5-dimethoxy-4-nitrophenyl)-N-(4-methylbenzyl)ethan-1-amine (25N-NB-4-Me) (23)**

Prepared as described for 25N-NB (2) using 0.000663 mol (150 mg) 2C-N (1) and 0.001326 mol (156 μL) m-tolualdehyde to give 115 mg (52.5 % yield) of a yellow oil (after flash column chromatography). The HCl salt was prepared as described for 2C-N (1) HCl to give fluffy slightly-yellow crystalline powder (mp: 182.0-184.0 °C). HRMS: Observed: 331.1645 (100), Theoretical: $C_{18}H_{23}N_2O_4$, 331.1652 (100), Δppm: -2.11.

**2-(2,5-dimethoxy-4-nitrophenyl)-*N*-(3-fluorobenzyl)ethan-1-amine (25N-NB-3-F) (24)**

Prepared as described for 25N-NB (2) using 0.000663 mol (150 mg) 2C-N (1) and 0.001326 mol (140.7 μL) 3-fluorobenzaldehyde to give 115.4 mg (52.0% yield) of a yellow oil (after flash column chromatography). The HCl salt was prepared as described for 2C-N (1) HCl to give a light-yellow crystalline needles (mp: 216.2-217.5 °C). HRMS: Observed: 335.1393 (100), Theoretical: $C_{17}H_{20}FN_2O_4$, 335.1402 (100), Δppm: -2.68.

**2-(2,5-dimethoxy-4-nitrophenyl)-*N*-(4-fluorobenzyl)ethan-1-amine (25N-NB-4-F) (25)**

Prepared as described for 25N-NB (2) using 0.000663 mol (150 mg) 2C-N (1) and 0.001326 mol (156 μL) 4-fluorobenzaldehyde to give small circular orange crystalline clusters. The HCl salt was prepared as described for 2C-N (1) HCl to give 106 mg (43.0% yield) of orange-brown fluffy crystalline needles (mp: 180.0-181.0 °C). HRMS: Observed: 335.1396 (100), Theoretical: $C_{17}H_{20}FN_2O_4$, 335.1402 (100), Δppm: -1.79.

**1,4-dimethoxy-2-methyl-5-(2-nitroethenyl)benzene**

18.5 g 2,5-dimethoxy-4-methylbenzaldehyde was dissolved in 50 mL nitromethane containing 2.0 g anhydrous ammonium acetate and 20 drops of 1,2-diaminocyclohexane. The solution was heated on a hot water bath (80 °C) for 2 hours and then left to sit for 4 days. At this point light orange crystals had formed. 30 mL of methanol was added and the solids were collected by gravity filtration. The collected crystals were washed twice with a small amount of methanol and then dried in an oven (~70 °C) to give 11 g of 1,4-dimethoxy-2-methyl-5-(2-nitroethenyl)benzene as light-yellow crystals.

**2-(2,5-dimethoxy-4-methylphenyl)ethan-1-amine (2C-D, 25D)**

(0.0493 mol) 11 g of the 1,4-dimethoxy-2-methyl-5-(2-nitroethenyl)benzene were dissolved in 60 mL dry (4Å MS) THF and added dropwise over ~20 minutes to a stirred suspension of (0.1478 mol) 5.62 g $LiAlH_4$ dissolved in THF on an ice bath while under nitrogen flow. Following the addition, the reaction was kept on ice for 20 min, at which point it was removed and placed under reflux. Reflux was maintained for 48 hours at which point the reaction was placed back on an ice bath and quenched by the cautious addition of ice chips and dilute KOH solution. 400 mL of ethyl acetate were then added and the suspension gravity filtered. The collected solids were washed extensively with additional ethyl acetate. The pooled organic was extracted with 0.5 N aqueous HCl (3 x ~200 mL). The pooled aqueous extracts were made basic with KOH pellets and extracted with ethyl acetate (3 x 150 mL). The pooled ethyl acetate extracts were washed with brine, dried over anhydrous $Na_2SO_4$ and evaporated under vacuum to give an amber oil which spontaneously formed a white solid (3.5 g, 36.4% yield). HCl salt was made as described for 25N (1) to give white crystalline solids.

**2-(2,5-dimethoxy-4-methylphenyl)-*N*-(2-methoxybenzyl)ethan-1-amine (25D-NBOMe)**

Prepared as described for 25N-NB (2) using 0.00107 mol (250 mg) 2C-D HCl and 0.00107 mol (147 mg) 2-methoxybenzaldehyde with the addition of 0.25 mL TEA (to convert the 2C-D HCl to the freebase) to give an amber oil (after flash column chromatography). The HCl salt prepared as described for 2C-N (1) HCl to give 203.3 mg (62.6% yield). The HCl salt was prepared as described for 2C-N (1) HCl as fluffy transparent needle clusters (mp: 168.6-168.8 °C). HRMS: Observed: 316.1904 (100), Theoretical: $C_{19}H_{26}NO_3$, 316.1904 (100), Δppm: 0.00.

**2-(2,5-dimethoxy-4-methylphenyl)-*N*-(naphthalen-1-ylmethyl)ethan-1-amine (25D-N1-Nap) (26)**

Prepared as described for 25N-NB (2) using 0.000866 mol (200 mg) 2C-N (1) and 0.00184 mol (287 mg) 1-napthaldehyde to give 460 mg of a yellow oil (which contained material from reduced 1-napthaldehyde) which was purified by crystallization (2X) from ethyl acetate and hexanes at 0 °C to give 230 mg (74.2% yield) of a fluffy white flake/scale crystalline solid (mp: 187.6-188.4 °C). HRMS: Observed: 336.1949 (100), Theoretical: $C_{22}H_{26}NO_2$, 336.1958 (100)Δppm: 2.67.

***N*-([1,1'-biphenyl]-2-ylmethyl)-2-(2,5-dimethoxy-4-methylphenyl)ethan-1-amine (25D-NBPh) (27)**
Prepared as described for 25N-NBPh (17) using 0.000866 mol (230 mg) 2C-D and 0.00184 mol (288 mL) biphenyl-2-carboxylate to give a light-yellow oil. Note: A straight to base workup was used as the HCl salt was soluble in organic solvent. The HCl salt was prepared as described for 2C-N (1) HCl to give 247.4 mg (71.8% yield) of a white crystalline powder (mp: 185.2-186.3 °C). HRMS: Observed: 362.2104 (100), Theoretical: $C_{24}H_{28}NO_2$, 362.2115 (100), Δppm: 3.04.

**1,2,4-trimethoxy-5-(2-nitrovinyl) benzene**
To a dry, argon flushed round-bottom flask was added anhydrous $NH_4Oac$ (1.17 g, 15.18 mmol) and nitromethane (42 mL). Cyclohexylamine (200 µL, 173 mg) and 2,4,5-trimethoxy benzaldehyde (10.02 g, 51.07 mmol) were added to the flask and the reaction was sealed with parafilm and wrapped in aluminum foil. The reaction was allowed to sit at ambient temperature for five days. An orangish-yellow precipitate formed during the reaction. The suspension was concentrated under vacuum and trace nitromethane was removed with the aid of repeated evaporations with added EtOH. The crude residue was washed in cold EtOH (200 proof) and then suspended in 15 ml of boiling EtOH followed by cooling at -20 °C overnight. The resulting orange crystalline solids were collected by gravity filtration. The product was recrystallized further by dissolving in a minimum volume of hot methanol and layering with $Et_2O$ and was combined with other crops to afford an orangish-yellow solid (9.39 g, 76.3% yield), (mp: 129.4-131.2 °C). Lit range: 127-130 °C [Shulgin and Shulgin. 1991] HRMS: Observed: 240.0861, Theoretical: $C_{11}H_{14}NO_5$: 240.0866, Δppm: 2.08.

**2-(2,4,5-trimethoxyphenyl)ethan-1-amine (25O)**
A dry, three-neck round-bottom flask with a teflon-coated stir bar was charged with dry (3Å molecular sieves) THF (135 mL) and the flask was cooled to 0 ℃. With vigorous stirring, lithium aluminum hydride (1.30 g, 34.3 mmol) was added in one portion as the flask was kept under active argon flow. The flask was fitted with a dry addition funnel and a solution of $AlCl_3$ (1.52 g, 11.40 mmol) in dry (3Å molecular sieves) THF (135 mL) was added dropwise at 0 ℃ over 15 min. When complete, a solution of 1,2,4-trimethoxy-5-[(1E)-2-nitroethenyl]benzene (4.0 g, 16.7 mmol) in dry (3Å molecular sieves) THF (135 mL) was added to the addition funnel and the solution was added dropwise over 1 hour at 0 ℃. The reaction was then stirred for 2 hours at rt and then quenched by slow dropwise addition of cold 1:1 THF: $H_2O$ (~50 mL) while cooled to 0 ℃. The reaction was basified with the addition of KOH solution to fully suspend all the inorganics and the resulting suspension was transferred to a separatory funnel. The aqueous suspension was extracted with EtOAc (3 x 100 mL) and the combined organics were washed with brine, dried over anhydrous $Na_2SO_4$, and concentrated under vacuum to afford a light amber oil. The next day, the oil was purified by short path distillation using a Kugelrohr for 11.5 hours at 170-175 °C (~0.3 mmHg) to obtain white waxy solids. (1.95 g, 55.2% yield), (mp: 103.7-105.0 °C). The HCl salt was prepared as described for 2C-N (1) HCl to give white, flaky particles, (mp: 191.3-192.5 °C). Lit: 187-188 °C [Shulgin and Shulgin. 1991] HRMS: Observed: 212.1278 (100), Theoretical: $C_{11}H_{18}NO_3$, 212.1281 (100), Δppm: 1.41.

**5-methoxy-6-(2-nitrovinyl) benzo[d] [1,3] dioxole**
0.0278 mol (5.0 g) 6-methoxy-2H-1,3-benzodioxole-5-carbaldehyde (prepared using a procedure comparable to Shulgin and Shulgin. 1991) was dissolved in 50 mL nitromethane and 0.5 g ammonium acetate was added followed by ~8 drops of cyclohexylamine from a glass pipette. The solution was sealed under argon and heated for 5 hours in an 80 °C water bath and then left to sit at room temperature overnight. The next day dark orange crystals had precipitated, and the reaction was placed at -20 °C for 24 hours. The crystalline solids were collected by decanting and washing sparingly with ethanol (200 proof), followed by drying with gentle heating under argon flow. The resulting solids were boiled in 100 mL methanol with grinding in an attempt to crystalize but were only partially soluble. The suspension was placed in the freezer for 24 hours, at which point the solids were collected by gravity filtration, washed twice with 10 mL ethanol (200 proof) and dried to give 4.3 g 5-methoxy-6-(2-nitrovinyl) benzo[d] [1,3] dioxole as orange-red solids (69.5% yield). The product was brought to the next step without further purification or characterization.

**2-(6-methoxybenzo[d][1,3]dioxol-5-yl)ethan-1-amine (2C-2)**
0.01927 mol (4.3 g) 5-methoxy-6-(2-nitrovinyl) benzo[d] [1,3] dioxole was dissolved in 60 mL anhydrous THF and added dropwise over 30 minutes to a stirred suspension of 0.0578 mol (2.19 g) $LiAlH_4$ in 100 mL anhydrous THF which was kept under argon and on an ice-water bath. After the addition, the reaction was allowed to recover to room temperature and then placed under reflux. Reflux was maintained for ~3 hours at which point TLC showed complete conversion. The workup was performed as described for 2C-H to give an amber oil of the crude product after evaporation of the ethyl acetate solvent. The crude freebase was purified via flash column chromatography (silica gel) starting with ethyl acetate:hexanes (4:1) containing 0.4% triethylamine and increasing to 10% ethanol in ethyl acetate (0.4% triethylamine). Desired fractions were pooled using MS (ASAP) and TLC to give 2.59 g of a beige waxy solid (68.9% yield). The HCl salt was prepared as described for 2C-N (1) HCl (with two additional crystallizations, 5-total, from MeOH: $Et_2O$) to give 2C-2 HCl as an off-white crystalline powder (mp: 227.8-229.2 °C). HRMS: Observed: 196.0967, Theoretical: $C_{10}H_{14}NO_3$, 196.0968, Δppm: 0.51.
**5-bromo-1,3-dimethoxy-2-(2-nitroprop-1-en-1-yl) benzene**
To a dry, argon flushed round-bottom flask was added anhydrous $NH_4OAc$ (503 mg, 6.52 mmol) and nitroethane (20 mL). Cyclohexylamine (100 µL, 86.7 mg, 0.87 mmol) and 4-bromo-2,6-dimethoxy benzaldehyde (5.0 g, 20.4 mmol) was added to the flask and the reaction was sealed and wrapped in aluminum foil. The reaction was allowed to sit at ambient temperature for ten days. A yellow precipitate formed during this time and the RBF was then allowed to stand at -20 °C overnight. The solid was filtered and washed with 95% EtOH (2 x 5 mL) to give 3.39 g of β-(4-bromo-2,6-dimethoxy)-α-methyl-nitrostyrene as yellow needle-like crystals. The filtrates were returned to -20 °C overnight to afford a second crop of the product as yellow crystalline solids (2.71 g, total mass was 6.1 g, ~quantitative yield) (mp: 110.2-111.4 °C). The product was brought to the next step without further purification or characterization.
**1-(4-bromo-2,6-dimethoxyphenyl)propan-2-amine (psi-DOB)**
A dry, three-neck round-bottom flask with a stir bar was charged with anhydrous THF (75 mL) and the flask was cooled to 0℃. With vigorous stirring, lithium aluminum hydride (1.50 g, 39.5 mmol) was added in one portion and the flask was flushed with argon. The flask was fitted with a dry addition funnel and a solution of $AlCl_3$ (1.76 g, 13.1 mmol) in dry THF (75 mL) was added dropwise at 0℃, over 15 min. When complete, a solution of β-(4-bromo-2,6-dimethoxy)-α-methyl nitrostyrene (6.0 g, 19.8 mmol) in dry THF (100 mL) was added to the addition funnel and the solution was added dropwise over 80 min at 0 °C. The reaction was then stirred for 2.5 hours at room temperature and then quenched by dropwise addition of cold 1:1 THF: $H_2O$ (~50 mL) at 0℃. The reaction was diluted into $H_2O$ (~500 mL) and basified with the addition of KOH. The resulting suspension was transferred to a separatory funnel and EtOAc (200 mL) was added. The mixture was extracted and then the aqueous layer was extracted further with EtOAc (2 x 100 mL). The combined organics were washed with brine, dried over anhydrous $Na_2SO_4$, and concentrated to afford a white solid (5.45 g, quantitative). The amine was converted to the hydrochloride salt by dissolving in absolute EtOH and adding a stoichiometric equivalent of concentrated HCl. Solvent and excess HCl were removed with a stream of warm air and the salt was washed with $Et_2O$ (3 x ~10 mL). The salt was crystallized three times from EtOH: $Et_2O$ as described previously. The freebase was directly converted to the hydrochloride salt as described for 25N (1) HCl to obtain white particles, (mp: 245.9-246.7 °C). HRMS: Observed: 276.0412 (90), 274.0431 (100), Theoretical: $C_{11}H_{17}BrNO_2$, 276.0417 (100), Δppm: -1.81, 274.0437 (100), Δppm: -2.18.
***N*-(naphthalen-1-ylmethyl)-2-(2,4,5-trimethoxyphenyl)ethan-1-amine (25O-N1-Nap) (28)**
0.00095 mol (200 mg) 25O freebase and 0.0014 mol (192.9 µL) 1-naphthaldehyde were dissolved in 15 mL dry (3Å molecular sieves) methanol, 4 mL dry (3Å molecular sieves) THF containing ~1.5g 3Å molecular sieves. The reaction was sealed under argon and protected from light for ~4 days. In general reductions were done after a minimum of 48 hours. The reaction was next placed on an ice-water bath and argon flow, at which point 0.00264 mol (100 mg) $NaBH_4$ was added in small portions over 10 minutes with vigorous mixing. After addition, the ice-bath was removed, and the reaction was continued

for an additional two hours at room temperature. The reaction was quenched by slow addition to a dilute KOH solution and then extracted with ethyl acetate (3 x 60 mL). Organic extracts were washed with brine (10 mL), pooled, dried over anhydrous $Na_2SO_4$ and evaporated under vacuum to give a yellow oil. This crude freebase was purified via flash column chromatography on silica gel with hexanes: ethyl acetate (1:4) containing 0.5% triethylamine. The ethyl acetate was slowly increased to 100%. Pure fractions were identified using MS-ASAP and TLC and combined to give 180 mg (54.1% yield) of a yellow oil. This was converted to the HCl salt by dissolving the freebase in 200 proof EtOH (~30 mL) and adding a stoichiometric equivalent of concentrated HCl solution. Solvent and excess HCl and water were removed via repeat evaporations of ethanol under a stream of warm air flow. The resulting solids were washed with $Et_2O$ (3 x ~10 mL) and the salt crystalized by dissolving in a minimum volume of hot EtOH followed by the addition of $Et_2O$ to yield, on standing at -20 °C overnight and repeated 3 times gave a white, flaky crystalline solids (280 mg, 75.9%) (mp: 178.4-179.6 °C). HRMS: Observed: 352.1907 (100), Theoretical: $C_{22}H_{26}NO_3$, 352.1907, Δppm: 0.00.

**2-(6-methoxybenzo[d][1,3]dioxol-5-yl)-*N*-(naphthalen-1-ylmethyl)ethan-1-amine (2C2-N1-Nap) (29)**

Prepared as described for 25N-NB (2) using 0.000866 mol (200 mg) 2C-2 HCl and 0.00184 mol (287 mg) 1-napthaldehyde to give 350 mg of an amber oil (which contained material from reduced 1-napthaldehyde). The HCl salt was prepared as described for 2C-N (1) HCl to give 270.6 mg (84.0% yield) of white crystalline powder (mp: 190.8-191.1 °C). HRMS: Observed: 336.1582 (100), Theoretical: $C_{21}H_{22}NO_3$, 336.1594 (100), Δppm: 3.57.

**1-(4-bromo-2,6-dimethoxyphenyl)-N-(naphthalen-1-ylmethyl)propan-2-amine (Psi-DOB-N1-Nap) (30)**

Prepared as described for 25O-N1-NAP (28), using 0.0013 mol (400 mg) psi-DOB HCl salt and 0.0019 mol (262.6 µL) 1-naphthaldehyde to give 490 mg of psi-DOB-N1-Nap HCl as a white powder (84.2 % yield). The procedure was the same except that 0.00258 mol (362.26 □L) of TEA was added to convert the starting amine HCl salt to the freebase and column chromatography was not carried out for purification. The HCl salt was prepared as described for 2C-N (1) HCl to give a white crystalline powder (mp: 207.9-209.2 °C). HRMS: Observed: 414.1059 (100), Theoretical: $C_{22}H_{25}BrNO_2$, 416.1043 (100), Δppm: -1.68, 414.1063, Δppm: 0.97.

**2-(6-methoxybenzo[d][1,3]dioxol-5-yl)-*N*-(2-methoxybenzyl)ethan-1-amine (2C2-NBOMe) (31)**

Prepared as described for 25N-NB (2) using 0.000768 mol (150 mg) 2C-2 and 0.00122 mol (167 mg) 2-methoxybenzaldehyde to give after flash column chromatography 110 mg (45.4% yield) of a colorless oil. The HCl salt was prepared as described for 2C-N (1) HCl to give white crystalline solids, mp: 147.1-148.1 °C.

***N*-(2-methoxybenzyl)-2-(2,4,5-trimethoxyphenyl)ethan-1-amine (25O-NBOMe) (32)**

Prepared as described for 25O-N1-Nap (28) using 0.000807 mol (200 mg) 25O HCl and 0.00125 mol (170 mg) 2-methoxybenzaldehyde to give 240 mg of the product as the HCl salt (80.8% yield). The workup was performed as described for that of 25O-N1-Nap (28) except that column chromatography was not carried out for purification and (~2M equivalent, 0.226 mL) TEA was added to the salt at the beginning of the reaction. The HCl salt was prepared as described for 2C-N (1) HCl to give an off-white crystalline powder (mp: 156.5-157.7 °C). HRMS: Observed: 332.1855, Theoretical: $C_{19}H_{26}NO_4$, 332.1856, Δppm: 0.30.

***N*-(2-cyclopropylbenzyl)-2-(2,4,5-trimethoxyphenyl)ethan-1-amine (25O-NBcP) (33)**

Prepared as described for 25N-NB (2) using 0.0012 mol (300 mg) 25O HCl and 0.00182 mol (266 mg) 2-cyclopropylbenzaldehyde to give an amber oil. The procedure was performed as described for 25O-N1-Nap (28) except that column chromatography was not carried out for purification and (~2M equivalents) TEA was added to the salt at the beginning of the reaction. The HCl salt was prepared as described for 2C-N (1) HCl to give 370 mg (80.9% yield) fluffy, white, crystalline powder (mp: 180.1-181.4°C). HRMS: Observed: 342.2047, Theoretical: $C_{21}H_{28}NO_3$, 342.2064, Δppm: 4.97.

***N*-(3-iodobenzyl)-2-(2,4,5-trimethoxyphenyl)ethan-1-amine (25O-NB-3-I) (34)**

Prepared as described for 25O-N1-Nap (28) using 0.00121 mol (300 mg) 25O HCl and 0.00147 mol (340 mg) 3-Iodobenzaldehyde to give 440 mg HCl salt (78.4% yield). The procedure was performed as described for 25O-N1-Nap (28) except that column chromatography was not carried out for purification and (~2M equivalent, 0.340 mL) TEA was added to the salt at the beginning of the reaction. The HCl salt was made as described for 2C-N (1) HCl to give fluffy white crystalline solids (mp: 155.5-156.1°C). HRMS: Observed: 428.0703, Theoretical: $C_{18}H_{23}INO_3$, 428.0717, Δppm: 3.27.

**2'-(((2,4,5-trimethoxyphenethyl)amino)methyl)-[1,1'-biphenyl]-4-ol (25O-NBPh-10'-OH) (35)**

Prepared as described for 25O-N1-Nap (28) using 0.00097 mol (240 mg) 25O HCl salt and 0.00116 mol (230 mg) 4'-hydroxy-biphenyl-3-carbaldehyde to give 128.9 mg of the HCl salt (32.3% yield). The procedure was performed as described for 25O-N1-Nap (28) except that column chromatography was not carried out for purification, $NH_4OH$ was used for base extraction and (~2M equivalent, 0.27 mL) TEA was added to the salt at the beginning of the reaction. The HCl salt was prepared as described for 2C-N (1) HCl to give fluffy white crystalline solids (mp: 259.1-260°C). HRMS: Observed: 394.1995, Theoretical: $C_{24}H_{28}NO_4$, 394.2013, Δppm: 4.57.

### NMR Numbering Schemes

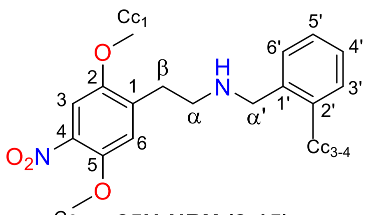

**25N-NBX (2-15)**

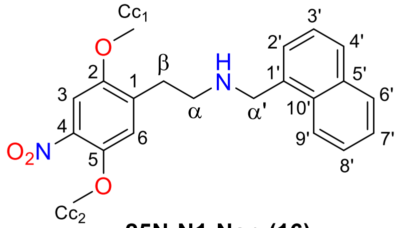

**25N-N1-Nap (16)**

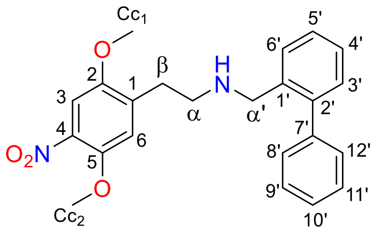

**25N-NBPh (17)**

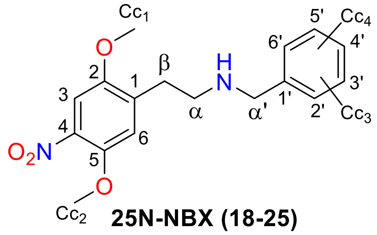

**25N-NBX (18-25)**

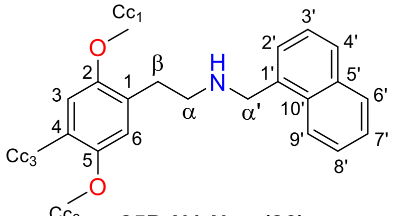

**25D-N1-Nap (26)**

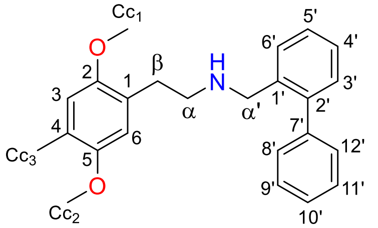

**25D-NBPh (27)**

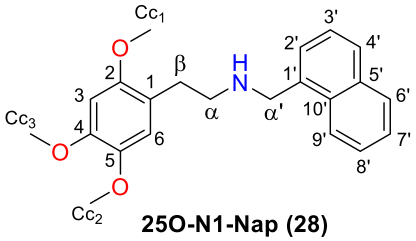

**25O-N1-Nap (28)**

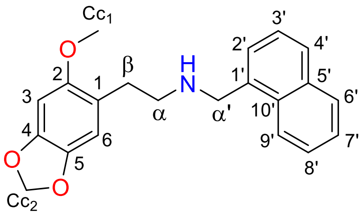

**2C2-N1-Nap (29)**

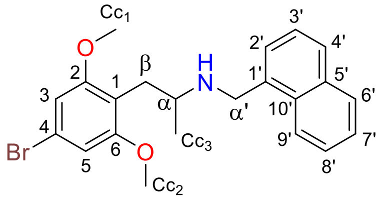

**ψ-DOB-N1-Nap (30)**

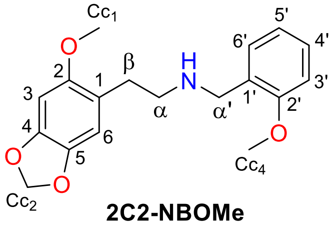

**2C2-NBOMe**

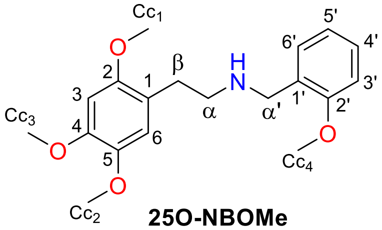

**25O-NBOMe**

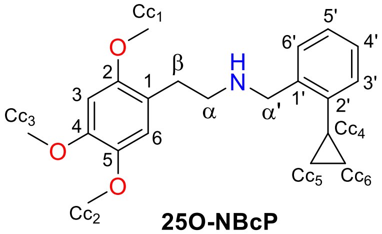

**25O-NBcP**

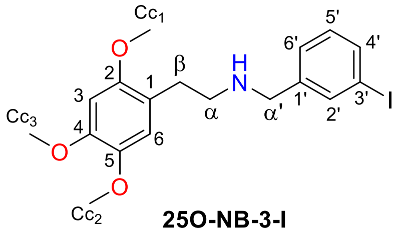

**25O-NB-3-I**

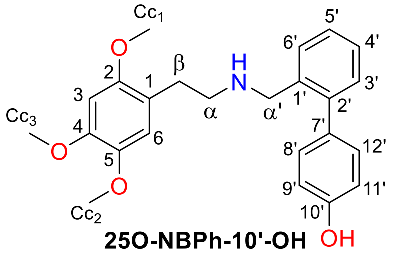

**25O-NBPh-10'-OH**

**$^1$H Chemical Shift Assignments Table 1 (compounds 2-6)**

Compounds dissolved at 20 mg/ml in d6-DMSO.

| Proton | δ (ppm) | | | | |
|---|---|---|---|---|---|
| | **25N-NB (2)** | **25N-NBOH (3)** | **25N-NBOMe (4)** | **25N-NBOEt (5)** | **25N-NBMe (6)** |
| $H_1$ | - | - | - | - | - |
| $H_2$ | - | - | - | - | - |
| $H_3$ | 7.49 s(1H) | 7.50 s(1H) | 7.50 s(1H) *overlap with $H_{6'}$ | 7.49 s(1H) | 7.51 s(1H) |
| $H_4$ | - | - | - | - | - |
| $H_5$ | - | - | - | - | - |
| $H_6$ | 7.33 s(1H) | 7.28 s(1H) | 7.30 s(1H) | 7.29 s(1H) | 7.33 s(1H) |
| α | 3.16-3.04 m(2H) *overlap with β | 3.19-3.09 m(2H) | 3.18-3.04m(2H) *overlap with β | 3.20-3.12 m(2H) | 3.28-3.18 m(2H) |
| β | 3.16-3.04 m(2H) *overlap with α | 3.08-3.01 m(2H) | 3.18-3.04 m(2H) *overlap with α | 3.12-3.05 m(2H) | 3.17-3.08 m(2H) |
| $α_1$ | 4.16 s(2H) | 4.10 s(2H) | 4.13 s(2H) | 4.13 s(2H) | 4.16 s(2H) |
| $H_{1'}$ | - | - | - | - | - |
| $H_{2'}$ | 7.62-7.56 m(1H) | - | - | - | - |
| $H_{3'}$ | 7.46-7.37 m(1H) *overlap with $H_{4'}$ | 6.97 dd($J$ = 8.2, 1.1 Hz, 1H) | 7.09 dd ($J$ = 8.3, 1.1 Hz, 1H) | 7.07 dd($J$ = 8.4, 1.0 Hz, 1H) | 7.30-7.24 m(1H) |
| $H_{4'}$ | 7.46-7.37 m(1H) Overlap with $H_{3','5}$ | 7.23 ddd($J$ = 8.2, 7.4, 1.7 Hz, 1H) | 7.41 ddd($J$ = 8.3, 7.5, 1.7 Hz, 1H) | 7.39 ddd($J$ = 8.3, 7.4, 1.7 Hz, 1H) | 7.30-7.24 m(1H) |
| $H_{5'}$ | 7.46-7.37 m(1H) *overlap with $H_{4'}$ | 6.84 td($J$ = 7.5, 1.2 Hz, 1H) | 7.00 td($J$ = 7.4, 1.0 Hz, 1H) | 6.99 td($J$ = 7.5, 1.1 Hz, 1H) | 7.30-7.24 m(1H) |
| $H_{6'}$ | 7.62-7.56 m(1H) | 7.40 dd($J$ = 7.6, 1.7 Hz, 1H) | 7.50 dd($J$ = 7.7, 1.7 Hz) *overlap with $H_3$ | 7.49 dd($J$ = 7.0, 1.8 Hz, 1H) | 7.59-7.53 m(1H) |
| $H_{c1}$ | 3.80 s($OCH_3$) | 3.80 s($OCH_3$) | 3.80 s($OCH_3$) | 3.81 s($OCH_3$) | 3.81 s($OCH_3$) |
| $H_{c2}$ | 3.88 s($OCH_3$) | 3.89 s($OCH_3$) | 3.89 s($OCH_3$) | 3.89 s($OCH_3$) | 3.90 s($OCH_3$) |
| $H_{c3}$ | - | - | 3.83 s($OCH_3$) | 4.09 q($J$ = 7.0 Hz) | 2.40 s($CH_3$) |
| $H_{C4}$ | - | - | - | 1.36 t($J$ = 6.9 Hz) | - |
| $NH^+$ | 9.60 s(2 $NH^+$) | 9.11 s(2 $NH^+$) | 9.27 s(2 $NH^+$) | 9.23 s(2 $NH^+$) | 9.44 s(2 $NH^+$) |
| OH | - | 10.30 s(1H) | - | - | - |

$^1$H NMR chemical shifts. Compounds dissolved at 20 mg/ml in d6-DMSO.

**$^1$H Chemical Shift Assignments Table 2 (compounds 7-11)**
Compounds dissolved at 20 mg/ml in d6-DMSO.

| Proton | δ (ppm) | | | | |
|---|---|---|---|---|---|
| | **25N-NBF (7)** | **25N-NBCl (8)** | **25N-NBBr (9)** | **25N-NBI (10)** | **25N-NBOCF$_2$H (11)** |
| $H_1$ | - | - | - | - | - |
| $H_2$ | - | - | - | - | - |
| $H_3$ | 7.50 s(1H) | 7.51 s(1H) | 7.51 s(1H) | 7.52 s(1H) *overlap with $H_{5'}$ | 7.50 s(1H) *overlap with $H_{4'}$ |
| $H_4$ | - | - | - | - | - |
| $H_5$ | - | - | - | - | - |
| $H_6$ | 7.31 s(1H) | 7.32 s(1H) | 7.33 s(1H) | 7.33 s(1H) | 7.30 s(1H) *overlap with $H_{5'}$ and $C_{c3}$ |
| α | 3.24-3.15 m (2H) | 3.29-3.10 m(2H) | 3.28-3.21 m(2H) | 3.25 dd(*J* = 9.6, 6.1 Hz, 2H) | 3.23-3.14 m(2H) |
| β | 3.13-3.04 m(2H) | 3.17-3.07 m(2H) | 3.12 dd(J = 9.3, 6.1 Hz, 2H) | 3.12 dd(*J* = 9.3, 6.1 Hz, 2H) | 3.09 dd (*J* = 9.1, 5.6 Hz, 2H) |
| $α_1$ | 4.22 s(2H) | 4.30 s(2H) | 4.29 s(2H) | 4.25 s(2H) | 4.20 s(2H) |
| $H_{1'}$ | - | - | - | - | - |
| $H_{2'}$ | - | - | - | - | - |
| $H_{3'}$ | 7.36-7.25 m(1H) *overlap with H5’ | 7.60-7.53 m(1H) | 7.72 dd(*J* = 8.0, 1.1 Hz, 1H) | 7.96 dd(*J* = 7.9, 1.2 Hz) | 7.27 dd(*J* = 8.3, 1.3 Hz, 1H) |
| $H_{4'}$ | 7.54-7.44 m(1H) | 7.49-7.42 m(1H) | 7.37 td(*J* = 7.7, 1.7 Hz, 1H) | 7.16 td(*J* = 7.7, 1.6 Hz, 1H) | 7.50 td(*J* = 7.7, 1.7 Hz, 1H) *overlap with $H_3$ |
| $H_{5'}$ | 7.36-7.25 m(1H) *overlap with $H_{3'}$ | 7.49-7.42 m(1H) | 7.48 td(*J* = 7.6, 1.1 Hz, 1H) | 7.50 td(*J* = 7.7, 1.1 Hz, 1H) *overlap with $H_3$ | 7.33 td(*J* = 7.7, 1.7 Hz, 1H) *overlap with $H_6$ |
| $H_{6'}$ | 7.74 td(*J* = 7.6, 1.7 Hz, 1H) | 7.84-7.76 m(1H) | 7.80 dd(*J* = 7.7, 1.7 Hz, 1H) | 7.76 dd(*J* = 7.8, 1.6 Hz, 1H) | 7.74 dd(*J* = 7.7, 1.7 Hz, 1H) |
| $H_{c1}$ | 3.81 s($OCH_3$) | 3.82s($OCH_3$) | 3.82 s($OCH_3$) | 3.82 s($OCH_3$) | 3.81 s($OCH_3$) |
| $H_{c2}$ | 3.89 s($OCH_3$) | 3.89 s($OCH_3$) | 3.90 s($OCH_3$) | 3.90 s($OCH_3$) | 3.89 s($OCH_3$) |
| $H_{c3}$ | - | - | - | - | 7.29 t(*J* = 73.60 Hz, 1H) |
| $NH^+$ | 9.59 s(2 $NH^+$) | 9.65 s(2 $NH^+$) | 9.68 s(2 $NH^+$) | 9.66 s(2 $NH^+$) | 9.54 s(2 $NH^+$) |

**$^1$H Chemical Shift Assignments Table 3 (compounds 12-15)**
Compounds dissolved at 20 mg/ml in d6-DMSO.

| Proton | δ (ppm) | | | |
|---|---|---|---|---|
| | **25N-NBOCF$_3$ (12)** | **25N-NBMDF$_2$ (13)** | **25N-NBCF$_3$ (14)** | **25N-NBNO$_2$ (15)** |
| $H_1$ | - | - | - | - |
| $H_2$ | - | - | - | - |
| $H_3$ | 7.51 s(1H) *overlap with $H_{5'}$ | 7.51 s(1H) | 7.51 s(1H) | 7.52 s(1H) |
| $H_4$ | - | - | - | - |
| $H_5$ | - | - | - | - |
| $H_6$ | 7.32 s(1H) | 7.32 s(1H) *overlap with $H_{5'}$ | 7.33 s(1H) | 7.33 s(1H) |
| α | 3.27-3.18 m(2H) | 3.29-3.20 m(2H) | 3.30-3.24 m(2H) | 3.29 dd(*J* = 9.5, 6.2 Hz, 2H) |
| β | 3.10 dd(*J* = 9.2, 9.2 Hz, 2H) | 3.15-3.01 m(2H) | 3.12 dd(*J* = 9.4, 6.2 Hz, 2H) | 3.13 dd(*J* = 9.2, 6.4 Hz, 2H) |
| $α_1$ | 4.24 s(2H) | 4.25 s(2H) | 4.33 s(2H) | 4.49 s(2H) |
| $H_{1'}$ | - | - | - | - |
| $H_{2'}$ | - | - | - | - |
| $H_{3'}$ | 7.48-7.43 m(1H) | - | 7.84 d(*J* = 7.9 Hz) | 8.22 dd(*J* = 8.2, 1.3 Hz, 1H) |
| $H_{4'}$ | 7.57 td(*J* = 7.7, 1.7 Hz, 1H) | 7.47 dd(*J* = 8.1, 0.9 Hz, 1H) | 7.65 t(*J* = 7.8 Hz) | 7.73 ddd(*J* = 8.7, 7.3, 1.6 Hz, 1H) |
| $H_{5'}$ | 7.52-7.48 m(1H) *overlap with and $H_3$ | 7.29 t(*J* = 8.1 Hz, 1H) *overlap with $H_6$ | 7.80 t(*J* = 7.8 Hz) | 7.87 td(*J* = 7.5, 1.3 Hz, 1H) |
| $H_{6'}$ | 7.91 dd(*J* = 7.7, 1.7 Hz, 1H) | 7.56 d(*J* = 7.4 Hz, 1H) | 8.04 d(*J* = 7.8 Hz, 1H) | 7.93 dd(*J* =7.8, 1.6 Hz, 1H) |
| $H_{c1}$ | 3.81 s($OCH_3$) | 3.79 s($OCH_3$) | 3.82 s($OCH_3$) | 3.84 s($OCH_3$) |
| $H_{c2}$ | 3.89 s($OCH_3$) | 3.89 s($OCH_3$) | 3.90 s($OCH_3$) | 3.90 s($OCH_3$) |
| $NH^+$ | 9.71 s(2 $NH^+$) | 9.72 s(2 $NH^+$) | 9.83 s(2 $NH^+$) | 9.66 s(2 $NH^+$) |

**$^1$H Chemical Shift Assignments Table 4 (compounds 16 and 17)**
Compounds dissolved at 20 mg/ml in d6-DMSO.

| Proton | δ (ppm) | |
|---|---|---|
| | **25N-N1-Nap (16)** | **25N-NBPh (17)** |
| $H_1$ | - | - |
| $H_2$ | - | - |
| $H_3$ | 7.51 s(1H) | 7.50 s(1H) *overlap with $H_{3'}$, $H_{4'}$, $H_{9'}$,$H_{10'}$, $H_{11'}$ |
| $H_4$ | - | - |
| $H_5$ | - | - |
| $H_6$ | 7.31 s(1H) | 7.21 s(1H) |
| α | 3.33-3.23 m(2H) | 3.10-2.80 m(2H) |
| β | 3.19-3.06 m(2H) | 3.10-2.80 m(2H) |
| $α_1$ | 4.68 s(2H) | 4.11 s(2H) |
| $H_{1'}$ | - | - |
| $H_{2'}$ | 7.83 d(*J* = 6.4 Hz, 1H) | - |
| $H_{3'}$ | 7.58 dd(*J* = 9.0, 7.2 Hz, 1H) | 7.54-7.41 m(1H) *overlap $H_3$, $H_{4'}$, $H_{9'}$,$H_{10'}$, $H_{11'}$ |
| $H_{4'}$ | 8.02 d(*J* = 8.4 Hz, 1H) | 7.54-7.41 m(1H) *overlap $H_{3'}$, $H_3$, $H_{9'}$,$H_{10'}$, $H_{11'}$ |
| $H_{5'}$ | - | 7.35-7.30 m(1H) |
| $H_{6'}$ | 8.02 d(*J* = 8.4 Hz, 1H) | 7.98-7.82 m(1H) |
| $H_{7'}$ | 7.61 ddd(*J* = 8.3, 6.8, 1.5 Hz, 1H) | - |
| $H_{8'}$ | 7.65 ddd(*J* = 8.4, 6.9, 1.6 Hz, 1H) | 7.41-7.36 m(1H) |
| $H_{9'}$ | 8.25 d(*J* = 8.4 Hz, 1H) | 7.54-7.41 m(1H) *overlap with $H_{3'}$, $H_3$, $H_{4'}$, $H_{10'}$, |
| $H_{10'}$ | - | 7.54-7.41 m(1H) *overlap with $H_{3'}$, $H_3$, $H_{4'}$, $H_{9'}$, $H_{11'}$ |
| $H_{11'}$ | - | 7.54-7.41 m(1H) *overlap with $H_{3'}$, $H_3$, $H_{4'}$,$H_{10'}$, |
| $H_{12'}$ | - | 7.41-7.36 m(1H) |

| | | |
|---|---|---|
| $H_{c1}$ | 3.78 s($OCH_3$) | 3.73 ($OCH_3$) |
| $H_{c2}$ | 3.88 s($OCH_3$) | 3.84 ($OCH_3$) |
| $NH^+$ | 9.57 s(2 $NH^+$) | 9.63 s(2 $NH^+$) |

**$^{1}H$ Chemical Shift Assignments Table 5 (compounds 18-21)**
Compounds dissolved at 20 mg/ml in d6-DMSO.

| Proton | δ (ppm) | | | |
|---|---|---|---|---|
| | **25N-NB-2-HO-3-Me (18)** | **25N-NB-2-MeO-3-F (19)** | **25N-NB-2,5-DiMeO (20)** | **25N-NB-3-OH (21)** |
| $H_1$ | - | - | - | - |
| $H_2$ | - | - | - | - |
| $H_3$ | 7.50 s(1H) | 7.50 s(1H) | 7.50 s(1H) | 7.50 s(1H) |
| $H_4$ | - | - | - | - |
| $H_5$ | - | - | - | - |
| $H_6$ | 7.29 s(1H) | 7.31 s(1H) | 7.30 s(1H) | 7.29 s(1H) |
| α | 3.14 s(2H) | 3.16 s(2H) | 3.12 s(2H) | 3.28-3.0 m(2H) *overlap with β |
| β | 3.07 dd(*J* = 8.7, 5.4 Hz, 2H) | 3.08 dd(*J* = 8.8, 5.3 Hz, 2H) | 3.07 dd(*J* = 8.6, 5.2 Hz, 2H) | 3.28-3.0 m(2H) *overlap with α |
| $α_1$ | 4.15 s(2H) | 4.19 s(2H) | 4.11 s(2H) | 4.06 s(2H) |
| $H_{1'}$ | - | - | - | - |
| $H_{2'}$ | - | - | - | 6.95 t(J = 2.0 Hz, 1H) |
| $H_{3'}$ | - | - | 7.00 t(*J* = 9.0 Hz, 1H) | - |
| $H_{4'}$ | 7.15 ddd(*J* = 7.6, 1.8, 0.9 Hz, 1H) | 7.35 ddd(*J* = 12.0, 8.3, 1.5 Hz, 1H) | 6.96 dd(*J* = 9.0, 3.0 Hz, 1H) | 6.82 ddd(*J* = 8.2, 2.5, 1.0 Hz, 1H) |
| $H_{5'}$ | 6.82 t(*J* = 7.5 Hz, 1H) | 7.17 td(*J* = 8.0, 5.1 Hz, 1H) | - | 7.21 t(*J* = 7.8 Hz, 1H) |
| $H_{6'}$ | 7.27 dd(*J* = 7.6, 1.5 Hz, 1H) | 7.45 dt(*J* = 7.8, 1.2 Hz) | 7.20 d(*J* = 3.0 Hz, 1H) | 6.98 dm(*J* = 7.6 Hz, 1H) |
| $H_{c1}$ | 3.81 s($OCH_3$) | 3.81 s($OCH_3$) | 3.80 s($OCH_3$) | 3.78 s($OCH_3$) |
| $H_{c2}$ | 3.89 s($OCH_3$) | 3.89 s($OCH_3$) | 3.89 s($OCH_3$) | 3.89 s($OCH_3$) |
| $H_{c3}$ | 9.08 s(OH) | 3.94 s($OCH_3$) | 3.78 s($OCH_3$) | 9.72 s(OH) |
| $Hc_4$ | 2.21 s($CH_3$) | - | 3.73 s($OCH_3$) | - |
| $NH^+$ | 9.16 s(2 $NH^+$) | 9.48 s(2 $NH^+$) | 9.28 s(2 $NH^+$) | 9.43 s(2 $NH^+$) |

**$^1$H Chemical Shift Assignments Table 6 (compounds 22-25)**
Compounds dissolved at 20 mg/ml in d6-DMSO.

| Proton | δ (ppm) | | | |
|---|---|---|---|---|
| | **25N-NB-3-Me (22)** | **25N-NB-4-Me (23)** | **25N-NB-3-F (24)** | **25N-NB-4-F (25)** |
| $H_1$ | - | - | - | - |
| $H_2$ | - | - | - | - |
| $H_3$ | 7.50 s(1H) | 7.49 s(1H) | 7.55-7.44 m(1H) *overlap with $C_{2'}$, $C_{5'}$ | 7.50 s(1H) |
| $H_4$ | - | - | - | - |
| $H_5$ | - | - | - | - |
| $H_6$ | 7.32 s(1H) | 7.30 s(1H) | 7.30 s(1H) | 7.30 s(1H) *overlap with $H_{3'}$,$H_{5'}$ |
| α | 3.19-3.00 m(2H) *overlap with β | 3.17-2.99 m(2H) *overlap with β | 3.36-2.88 m(2H) *overlap with β | 3.20-3.00 m(2H) *overlap with β |
| β | 3.19-3.00 m(2H) *overlap with α | 3.17-2.99 m(2H) *overlap with α | 3.36-2.88 m(2H) *overlap with α | 3.20-3.00 m(2H) *overlap with α |
| $\alpha_1$ | 4.09 s(2H) | 4.08 s(2H) | 4.19 s(2H) | 4.16 s(2H) |
| $H_{1'}$ | - | - | - | - |
| $H_{2'}$ | 7.39 bs(1H) *Overlap with $H_{6'}$ | 7.45 d(*J* = 8.0 Hz, 1H) | 7.55-7.44 m(1H) *overlap with $H_3$ and $H_{5'}$ | 7.64 ddm(*J* = 8.6, 5.5 Hz, 1H) |
| $H_{3'}$ | - | 7.23 d(*J* = 7.9 Hz, 1H) | - | 7.28 tm(*J* = 8.4 Hz, 1H) *overlap with $H_6$ |
| $H_{4'}$ | 7.22 d(*J* = 7.3 Hz, 1H) | - | 7.46 t(*J* = 8.6 Hz, 1H) | - |
| $H_{5'}$ | 7.32 tm(*J* = 7.8 Hz, 1H) *overlap with $H_6$ | 7.23 d(*J* = 7.9 Hz, 1H) | 7.55-7.44 m(1H) *overlap with $H_{2'}$ and $H_3$ | 7.28 tm(*J* = 8.4 Hz, 1H) *overlap with $H_6$ |
| $H_{6'}$ | 7.37 d(*J* = 5.8 Hz, 1H) *overlap with $H_{2'}$ | 7.45 d(*J* = 8.0 Hz, 1H) | 7.41 d(*J* = 7.5 Hz, 1H) | 7.64 ddm(*J* = 8.6, 5.5 Hz, 1H) |
| $H_{c1}$ | 3.80 s($OCH_3$) | 3.79 s($OCH_3$) | 3.80 s($OCH_3$) | 3.81 s($OCH_3$) |
| $H_{c2}$ | 3.89 s($OCH_3$) | 3.85 s($OCH_3$) | 3.88 s($OCH_3$) | 3.89 s($OCH_3$) |
| $H_{c3}$ | 2.32 s($CH_3$) | 2.31 s($CH_3$) | - | - |
| $NH^+$ | 9.48 m(2 $NH^+$) | 9.45 s(2 $NH^+$) | 9.51 s(2 $NH^+$) | 9.49 s(2 $NH^+$) |

**$^1$H Chemical Shift Assignments Table 7 (compounds 26-30)**
Compounds dissolved at 20 mg/ml in d6-DMSO.

| Proton | δ (ppm) | | | | |
|---|---|---|---|---|---|
| | **25D-N1-Nap (26)** | **25D-NBPh (27)** | **25O-N1-Nap (28)** | **2C2-N1-Nap (29)** | **Ψ-DOB-N1-Nap (30)** |
| $H_1$ | - | - | - | - | - |
| $H_2$ | - | - | | | - |
| $H_3$ | 6.83 s(1H) | 6.68 s(1H) | 6.69 s(1H) | 6.79 s(1H) | 6.88 s(1H) *overlap with $H_5$ |
| $H_4$ | - | - | - | - | - |
| $H_5$ | - | - | - | - | 6.88 s(1H) *overlap with $H_3$ |
| $H_6$ | 6.80 s(1H) | 6.78 s(1H) | 6.83 s(1H) | 6.81 s(1H) | - |
| α | 3.26-3.17 m(2H) | 2.94-2.87 m(2H) | 3.24-3.15 m(2H) | 3.21-3.11 m(2H) | 3.55-3.41 m(2H) |
| β | 3.05-2.95 m(2H) | 2.83-2.76 m(2H) | 3.00-2.90 m(2H) | 2.99-2.90 m(2H) | 3.07 dd(*J* = 12.7, 4.2 Hz, 1H) 2.96 dd(*J* = 12.7, 10.1 Hz, 1H) |
| $α_1$ | 4.67 s(1H) | 4.10 s(2H) | 4.66 s(2H) | 4.65 s(2H) | 4.79-4.61 m(1H) |
| $H_{1'}$ | - | - | - | - | - |
| $H_{2'}$ | 7.81 dd(*J* = 7.1, 0.8 Hz, 1H) | - | 7.81 dd(*J* = 7.0, 2.7 Hz, 1H) | 7.81 dd(*J* = 7.0, 2.0 Hz, 1H) | 7.84 d(*J* = 7.1 Hz, 1H) |
| $H_{3'}$ | 7.57 dd (*J* = 8.2, 7.2 Hz, 1H) *overlap with $H_{7'}$ | 7.52-7.45 m(1H) *overlap with $H_{4'}$, $H_{9'}$, $H_{11'}$ | 7.58 t(*J* = 7.9 Hz, 1H) *overlap with $H_{7'}$ | 7.57 t (*J* = 7.8 Hz, 1H) *overlap with $H_{7'}$ | 7.59 t(*J* = 8.0 Hz, 1H) *overlap with $H_{7'}$ |
| $H_{4'}$ | 8.01 d(*J* = 8.3 Hz, 1H) | 7.52-7.45 m(1H) *overlap with $H_{3'}$, $H_{9'}$, $H_{11'}$ | 8.02 d(*J* = 8.1 Hz, 1H) *overlap with $H_{6'}$ | 8.01 d(*J* = 8.1 Hz, 1H) *overlap with $H_{6'}$ | 8.02-7.99 m(1H) |
| $H_{5'}$ | - | 7.34-7.30 m(1H) | - | - | - |
| $H_{6'}$ | 8.01 d(*J* = 8.3 Hz, 1H) | 7.89-7.84 m(1H) | 8.02 d(*J* = 8.1 Hz, 1H) *overlap with $H_{4'}$ | 8.01 d(*J* = 8.1 Hz, 1H) *overlap with $H_{4'}$ | 8.05-8.02 m(1H) |
| $H_{7'}$ | 7.60 ddd(*J* = 8.8, 8.5, 1.4 Hz, 1H) *overlap with $H_{3'}$ and $H_{7'}$ | - | 7.63-7.57 m(1H) *overlap with $H_{3'}$ and $H_{8'}$ | 7.62-7.58 m(1H) overlap with $H_{3'}$ and $H_{8'}$ | 7.64-7.58 m(1H) overlap with $H_{3'}$ and $H_{8'}$ |
| $H_{8'}$ | 7.65 ddd(*J* = 8.3, 6.5, 1.6 Hz, 1H) *overlap with $H_{7'}$ | 7.40-7.36 m(1H) *overlap with $H_{12'}$ | 7.68-7.63 m(1H) *overlap with $H_{7'}$ | 7.65 ddd(*J* = 8.2, 6.9, 1.1 Hz, 1H) *overlap with $H_{7'}$ | 7.67 ddd(*J* = 8.2, 6.8, 1.3 Hz, 1H) *overlap with $H_{7'}$ |
| $H_{9'}$ | 8.24 d(*J* = 2.1 Hz, 1H) | 7.52-7.45 m(1H) *overlap with $H_{3'}$, $H_{4'}$, $H_{11'}$ | 8.25 d(*J* = 8.3 Hz, 1H) | 8.25 d(*J* = 8.2 Hz, 1H) | 8.22 d(*J* = 8.3 Hz, 1H |
| $H_{10'}$ | - | 7.45-7.42 m(1H) | - | - | - |
| $H_{11'}$ | - | 7.52-7.45 m(1H) *overlap with $H_{3'}$, $H_{4'}$, $H_{9'}$ | - | - | - |
| $H_{12'}$ | - | 7.40-7.36 m(1H) *overlap with $H_{12'}$ | - | - | - |
| $H_{c1}$ | 3.71 s($OCH_3$) | 3.66 s($OCH_3$) | - | 3.71 s(2H) | - |
| $H_{c2}$ | 3.74 s($OCH_3$) | 3.70 s($OCH_3$) | - | 5.95 s(2H) | - |
| $H_{c3}$ | 2.13 s($CH_3$) | 2.11 s($CH_3$) | - | - | - |

| | | | | | |
|---|---|---|---|---|---|
| $NH^+$ | 9.45 bs($NH_2^+$) | 9.45 bs($NH_2^+$) | 9.43 s($NH_2^+$) | 9.47 s($NH_2^+$) | 9.45 s($NH_2^+$)<br>9.32 s($NH_2^+$) |

**$^1$H Chemical Shift Assignments Table 8 (prediction compounds)**
Compounds dissolved at 20 mg/ml in d6-DMSO.

| Proton | δ (ppm) | | | | |
|---|---|---|---|---|---|
| | **2C2-NBOMe (31)** | **25O-NBOMe (32)** | **25O-NBcP (33)** | **25O-NB-3-I (34)** | **25O-NBPh-10'-OH (35)** |
| $H_1$ | - | - | - | - | - |
| $H_2$ | - | - | - | - | - |
| $H_3$ | 6.78, s(1H) *overlap with $H_6$ | 6.68, s(1H) | 6.69, s(1H) | 6.68, s(1H) | 6.65, s(1H) |
| $H_4$ | - | - | - | - | - |
| $H_5$ | - | - | - | - | - |
| $H_6$ | 6.79, s(1H) *overlap with $H_3$ | 6.80, s(1H) | 6.83, s(1H) | 6.80, s(1H) | 6.72, s(1H) |
| $H_\alpha$ | 3.03-2.95, m(2H) | 3.05-2.96, m(2H) | 3.18-3.08, m(2H) | 3.05-2.96, m(2H) | 2.93-2.85, m(2H) |
| $H_\beta$ | 2.93-2.84, m(2H) | 2.92-2.85, m(2H) | 2.99-2.89, m(2H) | 2.92-2.85, m(2H) | 2.81-2.73, m(2H) |
| $H_{\alpha'}$ | 4.10, s(2H) | 4.11, s(1H) | 4.34, s(2H) | 4.11, s(1H) | 4.10, s(1H) |
| $H_{1'}$ | - | - | - | - | - |
| $H_{2'}$ | - | 7.97, s(1H) | - | 7.97, s(1H) | - |
| $H_{3'}$ | 7.08, d (*J* = 8.3 Hz, 1H) | - | 7.08, d (*J* = 7.5 Hz, 1H) | - | 7.31-7.25, m(1H) |
| $H_{4'}$ | 7.44-7.35, tm (*J* = 7.9 Hz, 1H) | 7.80-7.75, dm (*J* = 8.0 Hz, 1H) | 7.34-7.28, tm (*J* = 7.4 Hz, 1H) | 7.80-7.75, dm (*J* = 8.0 Hz, 1H) | 7.46-7.39, m(1H) *overlap with $H_{5'}$ |
| $H_{5'}$ | 6.99, t (*J* = 7.5 Hz, 1H) | 7.24, t (*J* = 7.8 Hz, 1H) | 7.28-7.22, tm (*J* = 7.4 Hz, 1H) | 7.24, t (*J* = 7.8 Hz, 1H) | 7.46-7.39, m(1H) *overlap with $H_{4'}$ |
| $H_{6'}$ | 7.51-7.45, dm (*J* = 7.3 Hz, 1H) | 7.61-7.56, dm (*J* = 7.7 Hz, 1H) | 7.61-7.48, m(1H) | 7.61-7.56, dm (*J* = 7.7 Hz, 1H) | 7.85-7.78, m(1H) |
| $H_{7'}$ | - | - | - | - | - |
| $H_{8'}$ | - | - | - | - | 7.20-7.14, dm *J* = 8.5 Hz, 1H *overlap with $H_{12'}$ |
| $H_{9'}$ | - | - | - | - | 6.89-6.84, dm *J* = 8.5 Hz, 1H *overlap with $H_{11'}$ |
| $H_{10'}$ | - | | | | |

| | | | | | |
|---|---|---|---|---|---|
| $H_{11'}$ | - | - | - | - | 6.89-6.84, dm (*J* = 8.5 Hz, 1H) *overlap with $H_{9'}$ |
| $H_{12'}$ | - | - | - | - | 7.20-7.14, dm *J* = 8.5 Hz, 1H *overlap with $H_{8'}$ |
| $H_{c1}$ | 3.71, s($OCH_3$) | 3.76, s($OCH_3$) *overlap with $H_{Cc3}$ | 3.77, s($OCH_3$) *overlap with $H_{Cc3}$ | 3.76, s($OCH_3$) *overlap with $H_{Cc3}$ | 3.72, s($OCH_3$) |
| $H_{c2}$ | 5.94, s(2H) | 3.70, s($OCH_3$) | 3.70, s($OCH_3$) | 3.70, s($OCH_3$) | 3.67, s($OCH_3$) |
| $H_{c3}$ | 3.83, s($OCH_3$) | 3.77, s($OCH_3$) *overlap with $H_{Cc1}$ | 3.78, s($OCH_3$) *overlap with $H_{Cc1}$ | 3.77, s($OCH_3$) *overlap with $H_{Cc1}$ | 3.76, s($OCH_3$) |
| $H_{c4}$ | - | - | 2.20-2.02, m(1H) | - | - |
| $H_{c5}$ | - | - | 0.98-0.91, m(1H)<br><br>0.69-0.62, m(1H) | - | - |
| $H_{c6}$ | - | - | 0.98-0.91, m(1H)<br><br>0.69-0.62, m(1H) | - | - |
| $NH_2^+$ | 9.18, bs($1NH^+$) *overlap with 9.12<br><br>9.12, bs($1NH^+$) *overlap with 9.1 | 9.38, bs($NH_2^+$) | 9.28, bs($NH_2^+$) | 9.38, bs($NH_2^+$) | 9.40, bs($NH_2^+$) |
| OH | - | - | - | - | 9.70, s(OH) |

$^{13}$**C Chemical Shift Assignments Table 1 (compounds 2-6)**. Compounds dissolved at 20 mg/ml in d6-DMSO.

| Carbon | δ (ppm) | | | | |
|---|---|---|---|---|---|
| | **25N-NB (2)** | **25N-NBOH (3)** | **25N-NBOMe (4)** | **25N-NBOEt (5)** | **25N-NBMe (6)** |
| $C_1$ | 132.55 | 132.54 | 132.52 | 132.45 | 132.62 |
| $C_2$ | 150.38 | 150.37 | 150.34 | 150.34 | 150.37 |
| $C_3$ | 107.34 | 107.36 | 107.31 | 107.31 | 107.35 |
| $C_4$ | 137.66 | 137.65 | 137.64 | 137.69 | 137.65 |
| $C_5$ | 146.20 | 146.22 | 146.18 | 146.16 | 146.20 |
| $C_6$ | 116.76 | 116.73 | 116.69 | 116.66 | 116.65 |
| α | 45.22 | 45.06 | 45.38 | 45.41 | 45.90 |
| β | 26.56 | 26.54 | 26.45 | 26.48 | 26.47 |
| $α_1$ | 49.83 | 45.39 | 44.84 | 44.82 | 47.29 |
| $C_{1'}$ | 132.01 | 118.05 | 119.68 | 119.75 | 130.52 |
| $C_{2'}$ | 130.14 | 156.08 | 157.48 | 156.78 | 137.37 |
| $C_{3'}$ | 128.62 | 115.42 | 111.10 | 111.87 | 130.50 |
| $C_{4'}$ | 128.90 | 130.47 | 130.78 | 130.73 | 128.95 |
| $C_{5'}$ | 128.63 | 119.07 | 120.38 | 120.25 | 126.06 |
| $C_{6'}$ | 130.13 | 131.66 | 131.46 | 131.39 | 130.50 |
| $C_{c1}$ | 56.34 | 56.37 | 56.35 | 56.33 | 56.38 |
| $C_{c2}$ | 57.02 | 57.03 | 57.02 | 57.03 | 57.04 |
| $C_{c3}$ | - | - | 55.61 | 63.60 | 19.14 |
| $Cc_4$ | - | - | - | 14.47 | - |

**$^{13}$C Chemical Shift Assignments Table 2 (compounds 7-11)**. Compounds dissolved at 20 mg/ml in d6-DMSO.

| Carbon | δ (ppm) | | | | |
|---|---|---|---|---|---|
| | **25N-NBF (7)** | **25N-NBCl (8)** | **25N-NBBr (9)** | **25N-NBI (10)** | **25N-NBOCF$_2$H (11)** |
| $C_1$ | 132.47* | 132.43 | 132.42 | 132.45 | 132.44 |
| $C_2$ | 150.38 | 150.37 | 150.36 | 150.36 | 150.39 |
| $C_3$ | 107.34 | 107.34 | 107.34 | 107.36 | 107.36 |
| $C_4$ | 137.68 | 137.68 | 137.67 | 137.67 | 137.70 |
| $C_5$ | 146.18 | 146.19 | 146.19 | 146.22 | 146.23 |
| $C_6$ | 116.76 | 116.71 | 116.68 | 116.67 | 116.52 |
| α | 45.63 | 45.84 | 45.84 | 45.87 | 45.67 |
| β | 26.52 | 26.50 | 26.52 | 26.60 | 26.52 |
| $α_1$ | 43.03<br>42.99 | 46.98 | 49.53 | 54.36 | 44.05 |
| $C_{1'}$ | 119.20<br>119.05 | 129.90 | 131.60 | 134.86 | 122.82 |
| $C_{2'}$ | 161.79<br>159.34 | 133.51 | 124.07 | 101.28 | 149.54 |
| $C_{3'}$ | 115.71<br>115.50 | 129.61 | 132.91 | 139.53 | 117.91 |
| $C_{4'}$ | 132.47 | 130.86 | 130.99 | 130.82 | 130.93 |
| $C_{5'}$ | 124.73<br>124.70 | 127.57 | 128.12 | 128.69 | 125.30 |
| $C_{6'}$ | 131.52<br>131.44 | 131.90 | 131.68 | 130.60 | 132.04 |
| $C_{c1}$ | 56.37 | 56.38 | 56.38 | 56.42 | 56.38 |
| $C_{c2}$ | 57.02 | 57.04 | 57.05 | 57.07 | 57.04 |
| $C_{c3}$ | - | - | - | - | 116.52 (triplet, $J$ = 26 Hz) |
| * Coalesced with $C_{6'}$ only able to see via HMBC cross coupling with $C_3$ and $C_6$ $^1$H shifts. | | | | | |

$^{13}$**C Chemical Shift Assignments Table 3 (compounds 12-15)**. Compounds dissolved at 20 mg/ml in d6-DMSO.

| Carbon | δ (ppm) | | | |
|---|---|---|---|---|
| | **25N-NBOCF$_3$ (12)** | **25N-NBMDF$_2$ (13)** | **25N-NBCF$_3$ (14)** | **25N-NBNO$_2$ (15)** |
| $C_1$ | 132.40 | 132.37 | 132.41 | 132.44 |
| $C_2$ | 150.35 | 150.41 | 150.35 | 150.35 |
| $C_3$ | 107.31 | 107.36 | 107.32 | 107.31 |
| $C_4$ | 137.71 | 137.72 | 137.67 | 137.66 |
| $C_5$ | 146.16 | 146.21 | 146.19 | 146.19 |
| $C_6$ | 116.16 | 116.75 | 116.62 | 116.65 |
| α | 45.79 | 45.78 | 46.17 | 46.19 |
| β | 26.43 | 26.52 | 26.44 | 26.53 |
| $α_1$ | 43.70 | 42.94 | 46.35 | 47.26 |
| $C_{1'}$ | 124.56 | 115.08 | 130.41 | 127.30 |
| $C_{2'}$ | 146.69 | 141.75 | 127.45 (q, *J* = 29.3 Hz) | 148.34 |
| $C_{3'}$ | 120.36 | 142.63 | 126.27 126.21 | 125.27 |
| $C_{4'}$ | 130.97 | 110.79 | 132.97 | 130.70 |
| $C_{5'}$ | 127.49 | 124.62 | 129.45 | 134.33 |
| $C_{6'}$ | 131.89 | 126.25 | 130.41 | 133.41 |
| $C_{c1}$ | 56.32 | 56.38 | 56.37 | 56.39 |
| $C_{c2}$ | 57.02 | 57.02 | 57.05 | 57.03 |
| $C_{c3}$ | 120.00 (q, *J* = 256 Hz) | 131.04 (t, *J* = 254.5 Hz) | 124.00 (q, *J* = 28.3 Hz) | - |

**$^{13}$C Chemical Shift Assignments Table 4 (compounds 16 and 17)**. Compounds dissolved at 20 mg/ml in d6-DMSO.

| Carbon | δ (ppm) | |
|---|---|---|
| | **25N-N1-Nap (16)** | **25N-NBPh (17)** |
| $C_1$ | 132.61 | 132.39 |
| $C_2$ | 150.36 | 150.28 |
| $C_3$ | 107.34 | 107.27 |
| $C_4$ | 137.65 | 137.62 |
| $C_5$ | 146.18 | 146.18 |
| $C_6$ | 116.66 | 116.51 |
| α | 46.06 | 45.55 |
| β | 26.53 | 26.35 |
| $α_1$ | 46.74 | 47.20 |
| $C_{1'}$ | 128.09 | 129.53* |
| $C_{2'}$ | 129.03 | 142.06 |
| $C_{3'}$ | 125.32 | 127.91 |
| $C_{4'}$ | 129.53 | 128.76 |
| $C_{5'}$ | 133.25 | 130.37 |
| $C_{6'}$ | 128.64 | 129.43 |
| $C_{7'}$ | 126.24 | 139.55 |
| $C_{8'}$ | 126.77 | 129.30 |
| $C_{9'}$ | 123.69 | 128.60 |
| $C_{10'}$ | 131.06 | 127.68* |
| $C_{11'}$ | - | 128.60 |
| $C_{12'}$ | - | 129.30 |
| $C_{c1}$ | 56.34 | 56.31 |
| $C_{c2}$ | 57.02 | 57.04 |
| $C_{c3}$ | - | - |
| $C_{C4}$ | | - |
| * Chemical shifts assigned to 1',10' appear very close so assignment is tentative. | | |

**$^{13}$C Chemical Shift Assignments Table 5 (compounds 19-21)**. Compounds dissolved at 20 mg/ml in d6-DMSO.

| Carbon | δ (ppm) | | | |
|---|---|---|---|---|
| | **25N-NB-2-OH-3-Me (18)** | **25N-NB-2-MeO-3-F (19)** | **25N-NB-2,5-DiMeO (20)** | **25N-NB-3-OH (21)** |
| $C_1$ | 132.53 | 132.49 | 132.50 | 132.53 |
| $C_2$ | 150.35 | 150.37 | 150.36 | 150.39 |
| $C_3$ | 107.33 | 107.34 | 107.33 | 107.35 |
| $C_4$ | 137.65 | 137.66 | 137.65 | 137.67 |
| $C_5$ | 146.17 | 146.21 | 146.19 | 146.19 |
| $C_6$ | 116.71 | 116.71 | 116.71 | 116.90 |
| α | 45.39 | 45.63 | 45.36 | 45.21 |
| β | 26.55 | 26.49 | 26.47 | 26.55 |
| $α_1$ | 45.51 | 44.07<br>44.04 | 44.67 | 49.84 |
| $C_{1'}$ | 119.32 | 126.48<br>126.45 | 120.45 | 133.14 |
| $C_{2'}$ | 153.75 | 145.71<br>145.60 | 151.42 | 116.74 |
| $C_{3'}$ | 125.69 | 155.71<br>153.27 | 112.11 | 157.57 |
| $C_{4'}$ | 131.74 | 118.10<br>117.91 | 115.06 | 115.84 |
| $C_{5'}$ | 119.68 | 123.99<br>123.91 | 152.88 | 129.67 |
| $C_{6'}$ | 129.33 | 126.91<br>126.88 | 117.28 | 120.41 |
| $C_{c1}$ | 56.34 | 56.38 | 56.35 | 56.34 |
| $C_{c2}$ | 57.01 | 57.03 | 57.02 | 57.02 |
| $C_{c3}$ | - | 61.67 | 56.00 | - |
| $C_{c4}$ | 16.67 | - | 55.58 | - |

**$^{13}$C Chemical Shift Assignments Table 6 (compounds 22-25)**. Compounds dissolved at 20 mg/ml in d6-DMSO.

| Carbon | δ (ppm) | | | |
|---|---|---|---|---|
| | **25N-NB-3-Me (22)** | **25N-NB-4-Me (23)** | **25N-NB-3-F (24)** | **25N-NB-4-F (25)** |
| $C_1$ | 132.46 | 132.56 | 132.44 | 132.40 |
| $C_2$ | 150.38 | 150.41 | 150.43 | 150.38 |
| $C_3$ | 107.33 | 107.35 | 107.38 | 107.34 |
| $C_4$ | 137.70 | 137.70 | 137.73 | 137.71 |
| $C_5$ | 146.15 | 146.22 | 146.22 | 146.16 |
| $C_6$ | 116.76 | 116.81 | 116.83<br>*overlap with $C_{2'}$ | 116.78 |
| α | 45.24 | 45.13 | 45.38 | 45.17 |
| β | 26.56 | 26.59 | 26.64 | 26.58 |
| $α_1$ | 49.85 | 49.64 | 49.21 | 49.02 |
| $C_{1'}$ | 131.84 | 128.88 | 134.66<br>134.58 | 128.24 |
| $C_{2'}$ | 127.10 | 130.12 | 117.07<br>116.83<br>*overlap with $C_6$ | 132.57<br>132.48 |
| $C_{3'}$ | 137.81 | 129.18 | 163.19<br>160.76 | 115.58<br>115.36 |
| $C_{4'}$ | 129.48 | 138.39 | 115.96<br>115.76 | 163.59<br>161.15 |
| $C_{5'}$ | 128.55 | 129.18 | 130.80<br>130.72 | 115.58<br>115.36 |
| $C_{6'}$ | 130.60 | 130.12 | 126.33<br>126.30 | 132.57<br>132.48 |
| $C_{c1}$ | 56.34 | 56.37 | 56.38 | 56.34 |
| $C_{c2}$ | 57.01 | 57.06 | 57.06 | 57.02 |
| $C_{c3}$ | 20.91 | 20.81 | - | - |

$^{13}$**C Chemical Shift Assignments Table 7 (compounds 26-30)**. Compounds dissolved at 20 mg/ml in d6-DMSO.

| Carbon | δ (ppm) | | | | |
|---|---|---|---|---|---|
| | **25D-N1-Nap (26)** | **25D-NBPh (27)** | **25O-N1-Nap (28)** | **2C2-N1-Nap (29)** | **Ψ-DOB-N1-Nap (30)** |
| $C_1$ | 122.70 | 124.93 | 115.95 | 116.85 | 120.98 |
| $C_2$ | 150.71 | 151.05 | 151.43 | 152.21 | 158.67 |
| $C_3$ | 112.84 | 112.72 | 98.48 | 95.26 | 107.64 |
| $C_4$ | 124.97 | 122.42 | 148.60 | 146.76 | 111.75 |
| $C_5$ | 151.12 | 150.61 | 142.55 | 140.44 | 107.64 |
| $C_6$ | 114.05 | 113.95 | 115.03 | 109.70 | 158.67 |
| α | 46.94 | 47.07 | 47.02 | 46.97 | 53.35 |
| β | 26.44 | 26.27 | 25.91 | 26.16 | 25.97 |
| $\alpha_1$ | 46.69 | 46.38 | 46.64 | 46.62 | 44.35 |
| $C_{1'}$ | 128.21 | 129.60 | 128.20 | 128.18 | 128.37 |
| $C_{2'}$ | 129.08 | 142.11 | 129.05 | 129.01 | 128.78 |
| $C_{3'}$ | 125.37 | 127.89 | 125.33 | 125.32 | 125.32 |
| $C_{4'}$ | 129.57 | 128.74 | 129.53 | 129.50 | 129.43 |
| $C_{5'}$ | 133.30 | 130.33 | 133.27 | 133.26 | 133.28 |
| $C_{6'}$ | 128.68 | 129.49 | 128.65 | 128.64 | 128.70 |
| $C_{7'}$ | 126.29 | 139.55 | 126.25 | 126.24 | 126.24 |
| $C_{8'}$ | 126.82 | 129.27 | 126.77 | 126.76 | 126.81 |
| $C_{9'}$ | 123.76 | 128.57 | 123.74 | 123.74 | 123.59 |
| $C_{10'}$ | 131.13 | 127.66 | 131.10 | 131.09 | 131.05 |
| $C_{11'}$ | - | 128.57 | - | - | - |
| $C_{12'}$ | - | 129.27 | - | - | - |
| $C_{c1}$ | 55.73 | 55.73 | 56.15 | 100.95 | 56.20 |
| $C_{c2}$ | 55.91 | 55.84 | 56.39 | 56.37 | 56.20 |
| $C_{c3}$ | 16.02 | 16.00 | 55.88 | - | 15.71 |

$^{13}$**C Chemical Shift Assignments Table 8**. Compounds dissolved at 20 mg/ml in d6-DMSO.

| Carbon | δ (ppm) | | | | |
|---|---|---|---|---|---|
| | **2C2-NBOMe (31)** | **25O-NBOMe (32)** | **25O-NBcP (33)** | **25O-NB-3-I (34)** | **25O-NBPh-10'-OH (35)** |
| $C_1$ | 116.74<br>116.72 | 115.88 | 115.87 | 115.77 | 115.69 |
| $C_2$ | 152.22 | 151.40 | 151.45 | 151.43 | 151.34 |
| $C_3$ | 95.25 | 98.47 | 98.49 | 98.46 | 98.39 |
| $C_4$ | 146.78 | 148.59 | 148.64 | 148.64 | 148.57 |
| $C_5$ | 140.43 | 142.53 | 142.56 | 142.53 | 142.48 |
| $C_6$ | 109.76 | 115.04 | 115.09 | 115.07 | 114.88 |
| $C_\alpha$ | 46.25<br>46.22 | 46.28 | 46.94<br>*overlap with $C_{\alpha'}$ | 46.33 | 46.44 |
| $C_\beta$ | 26.08 | 25.81 | 25.93 | 25.96 | 25.74 |
| $C_{\alpha'}$ | 44.79<br>44.74 | 44.74 | 46.94<br>*overlap with $C_\alpha$ | 48.91 | 47.09 |
| $C_{1'}$ | 119.70 | 119.71 | 131.70 | 134.59 | 129.64 |
| $C_{2'}$ | 157.49 | 157.49 | 141.95 | 138.55 | 142.24 |
| $C_{3'}$ | 111.08 | 111.07 | 126.08 | 94.90 | 130.37<br>*overlap with $C_{8'}$ and $C_{12'}$ |
| $C_{4'}$ | 130.76 | 130.73 | 129.02 | 137.52 | 128.60 |
| $C_{5'}$ | 120.36 | 120.35 | 125.93 | 130.68 | 127.26 |
| $C_{6'}$ | 131.47 | 131.47 | 129.92 | 129.55 | 129.38 |
| $C_{7'}$ | - | - | - | - | 129.98 |
| $C_{8'}$ | - | - | - | - | 130.37<br>*overlap with $C_{3'}$ and $C_{12'}$ |
| $C_{9'}$ | - | - | - | - | 115.31<br>*overlap with $C_{11'}$ |
| $C_{10'}$ | - | - | - | - | 157.04 |
| $C_{11'}$ | - | - | - | - | 115.31<br>*overlap with $C_{9'}$ |
| $C_{12'}$ | - | - | - | - | 130.37<br>*overlap with $C_{3'}$ and $C_{8'}$ |
| $Cc_1$ | - | 56.14 | 56.19 | 56.17 | 56.10 |
| $Cc_2$ | - | 56.38 | 56.44 | 56.40 | 56.37 |
| $Cc_3$ | - | 55.89 | 55.91 | 55.89 | 55.87 |
| $Cc_4$ | - | 55.59 | 12.52 | - | - |
| $Cc_5$ | - | - | 7.39<br>*overlap with $Cc_6$ | - | - |
| $Cc_6$ | - | - | 7.39<br>*overlap with $Cc_5$ | - | - |

**$^{19}$F NMR Chemical Shift Assignments Table 1.** Compounds dissolved at 20 mg/ml in d6-DMSO. Reference $CFCl_3$ (0.00 ppm).

| Fluorine | δ (ppm) | | | | | | | |
|---|---|---|---|---|---|---|---|---|
| | **25N-NBF (7)** | **25N-NBOCF$_2$H (11)** | **25N-NBOCF$_3$ (11)** | **25N-NBMDF$_2$ (13)** | **25N-NBCF$_3$ (14)** | **25N-NB-2-MeO-3-F (19)** | **25N-NB-3-F (24)** | **25N-NB-4-F (25)** |
| | -115.54 | -81.26 | -55.80 | -48.13 | -57.19 | -129.72 | -112.17 | -112.45 |

**[1]H NMR Spectra for Synthesized *N*-benzyl-phenethylamines.**

**Spectra 1. [1]H NMR of 25N-NB (2) HCl.**

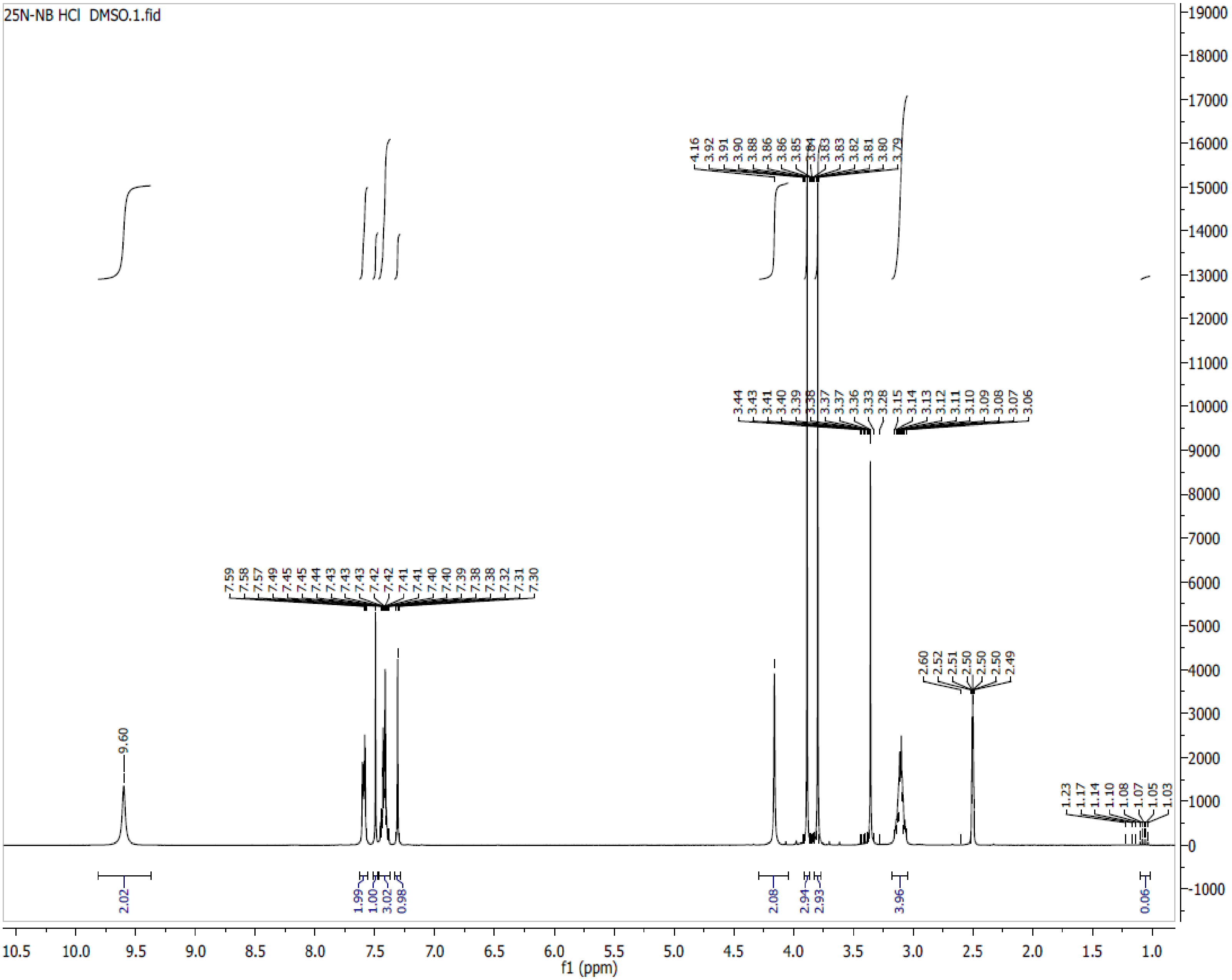

**Spectra 2. $^1$H NMR of 25N-NBOH (3) HCl**

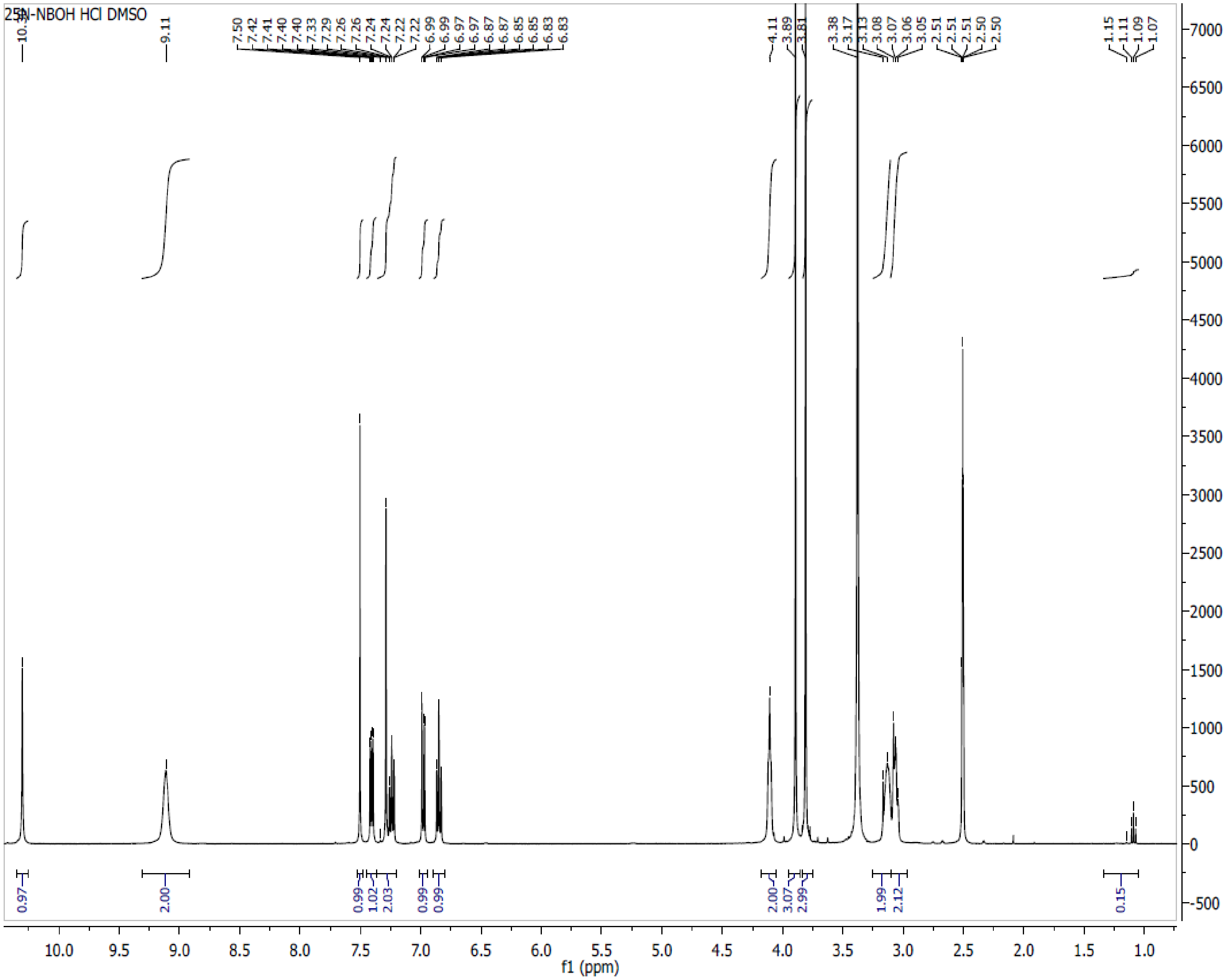

**Spectra 3. $^1$H NMR of 25N-NBOMe (4) HCl**

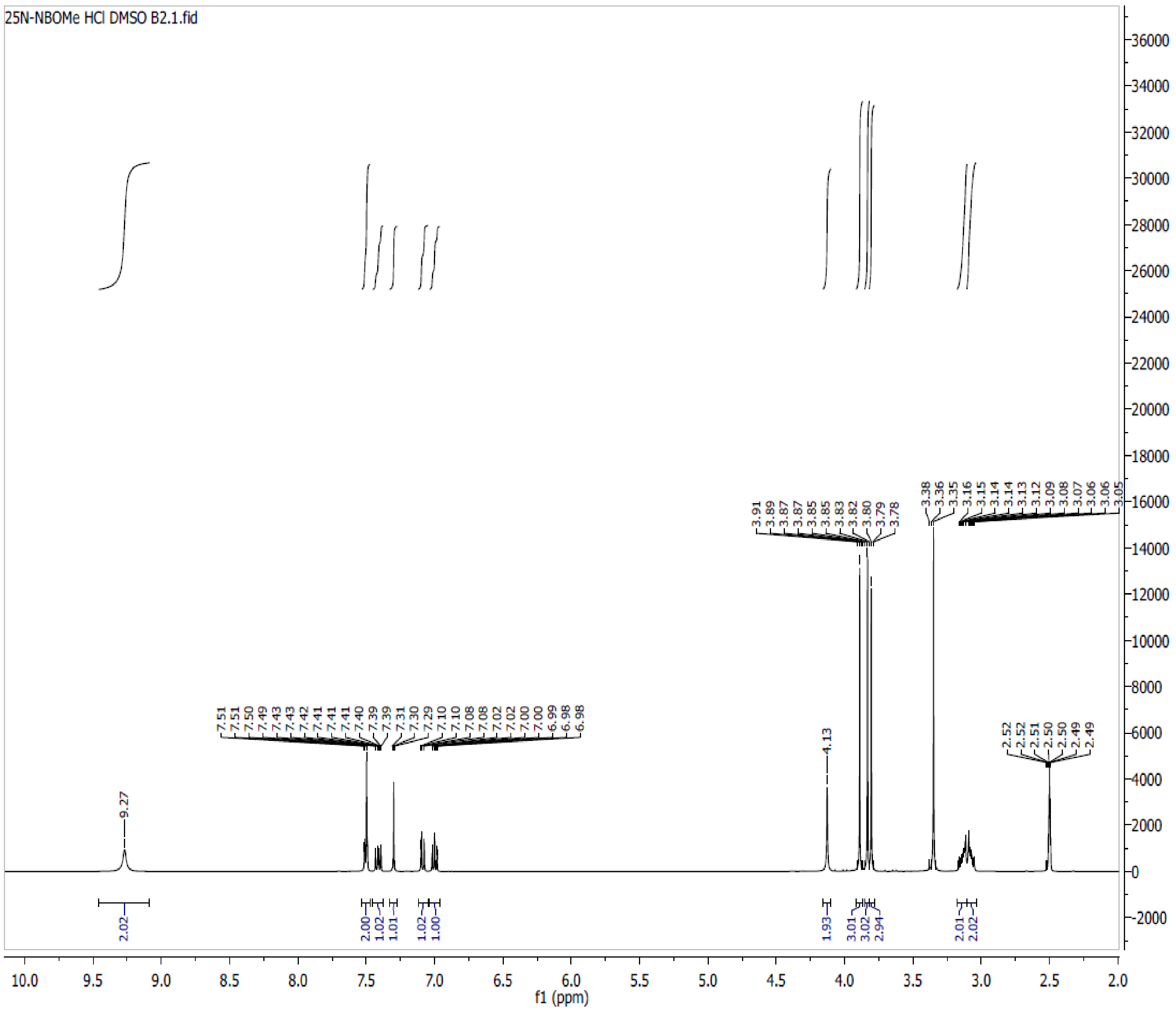

**Spectra 4. $^{1}$H NMR of 25N-NBOEt (5) HCl**

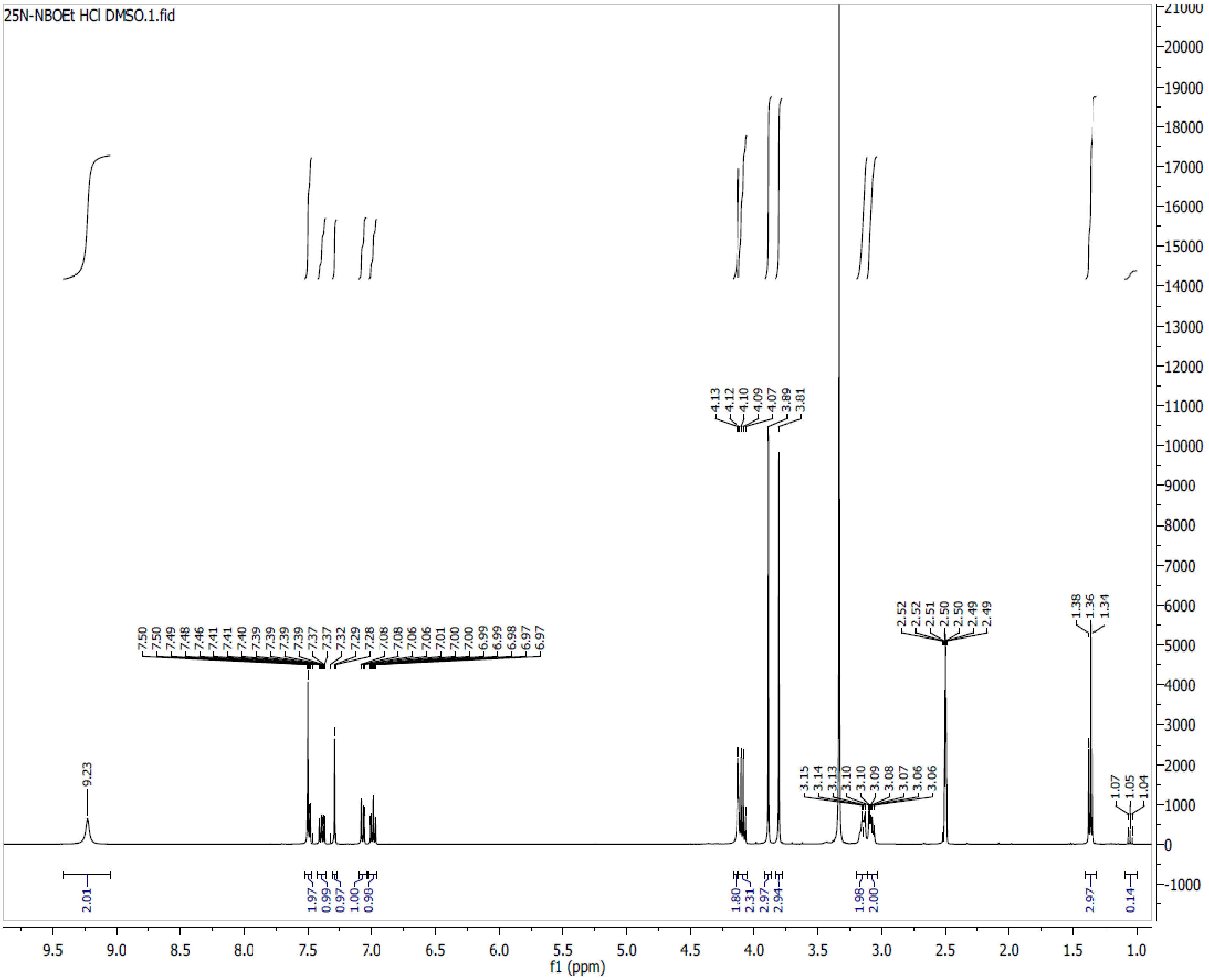

**Spectra 5. $^{1}$H NMR of 25N-NBMe (6) HCl**

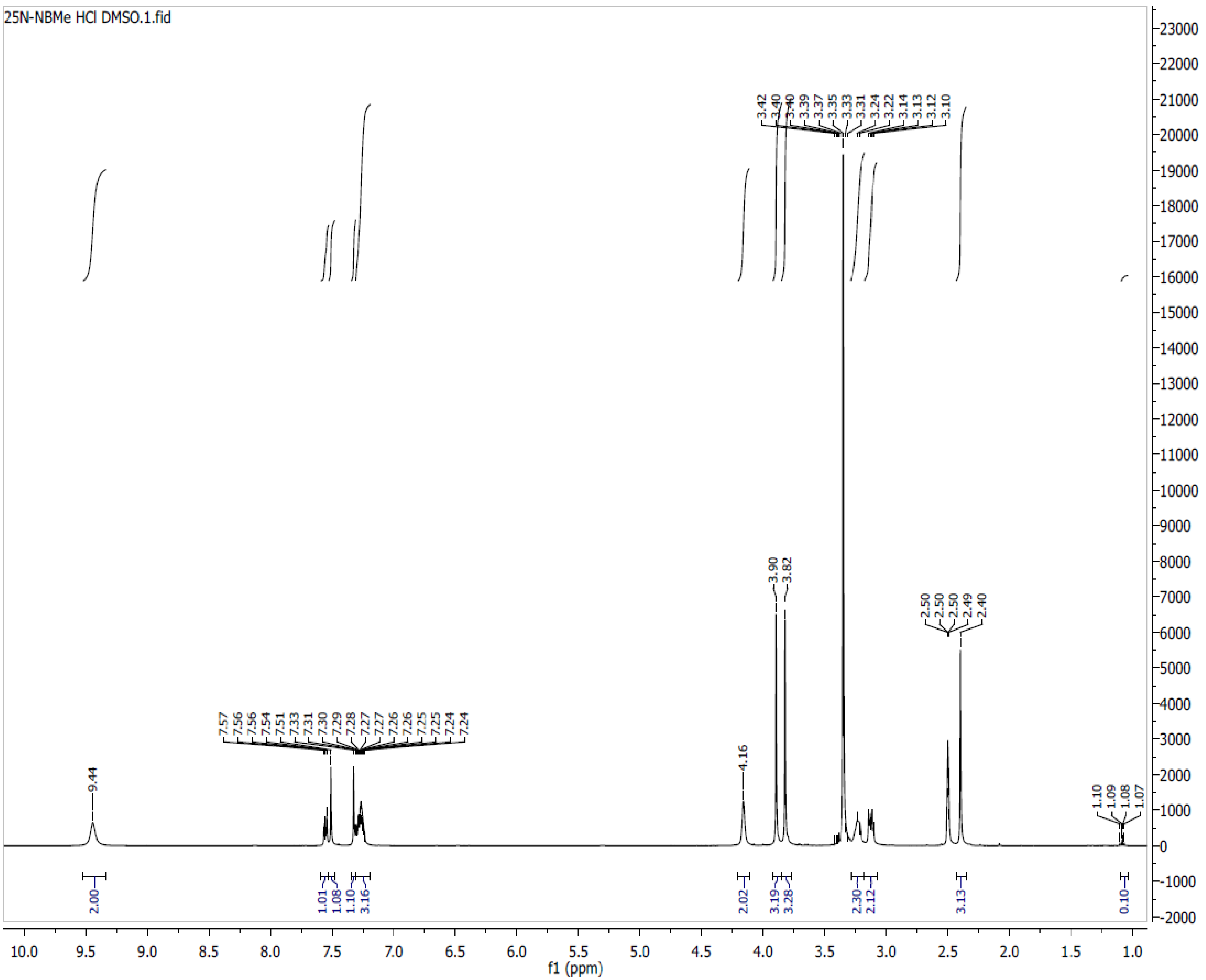

**Spectra 6. $^{1}$H NMR of 25N-NBF (7) HCl**

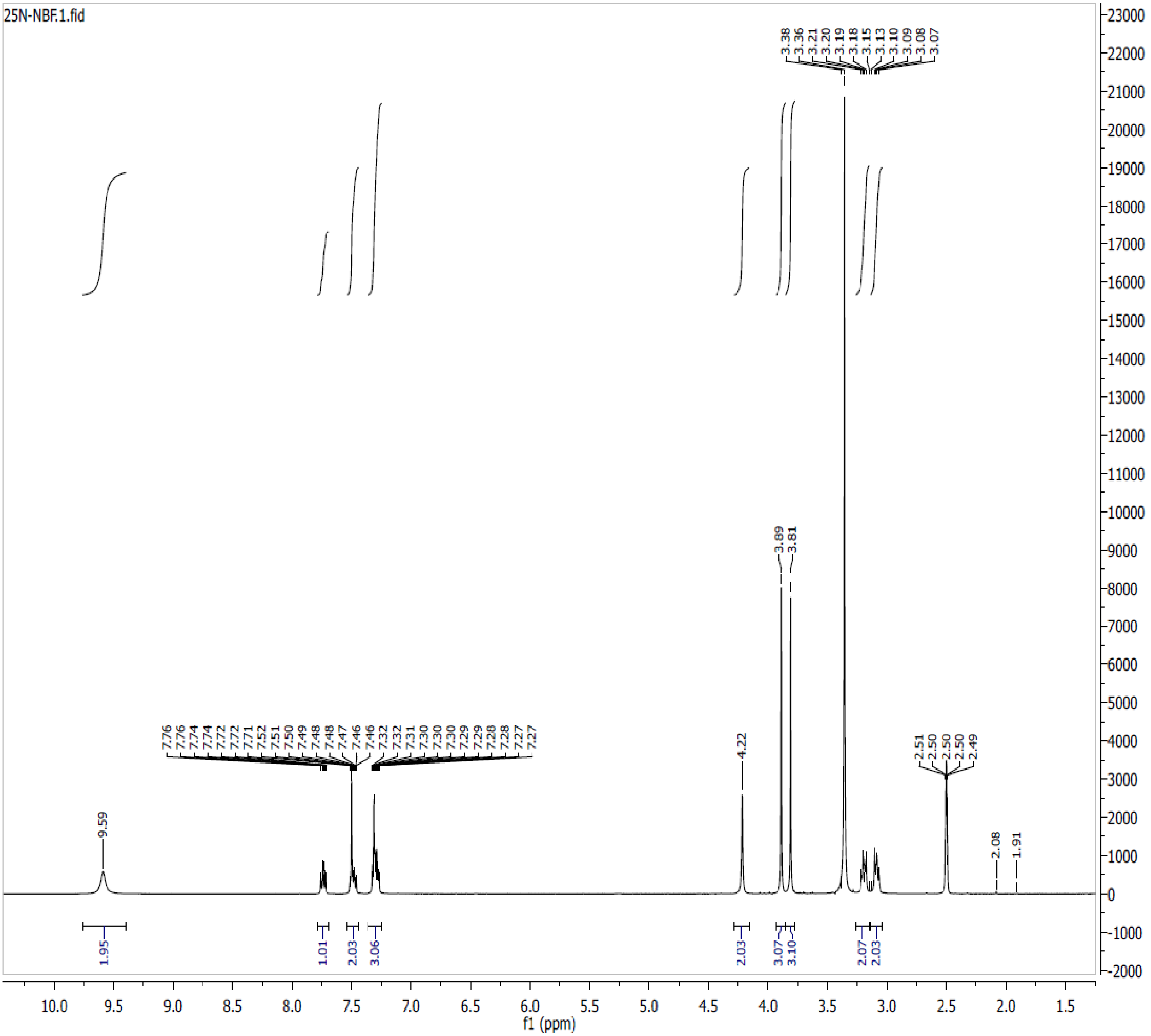

**Spectra 7. $^{1}$H NMR of 25N-NBCl (8) HCl**

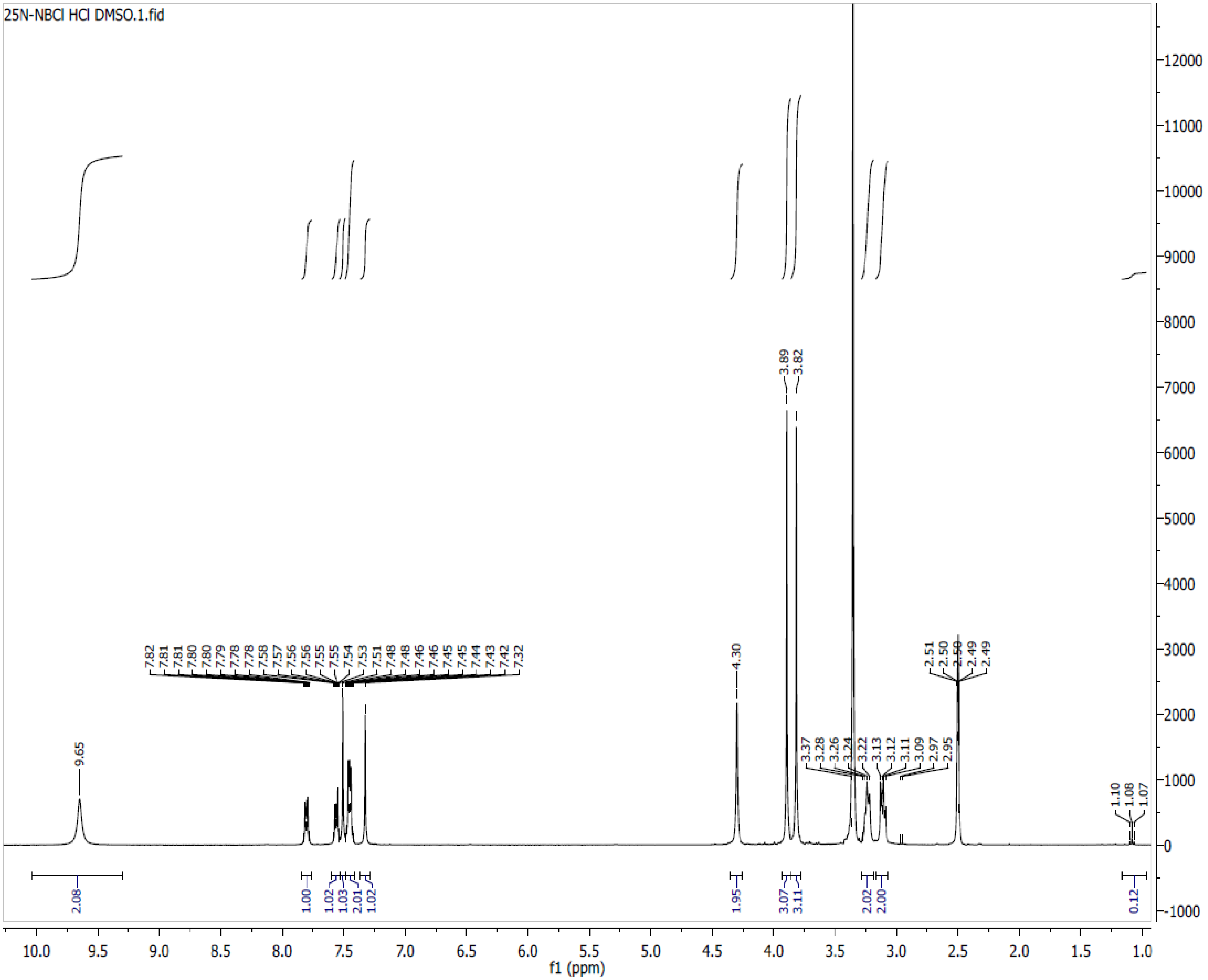

**Spectra 8. $^{1}$H NMR of 25N-NBBr (9) HCl**

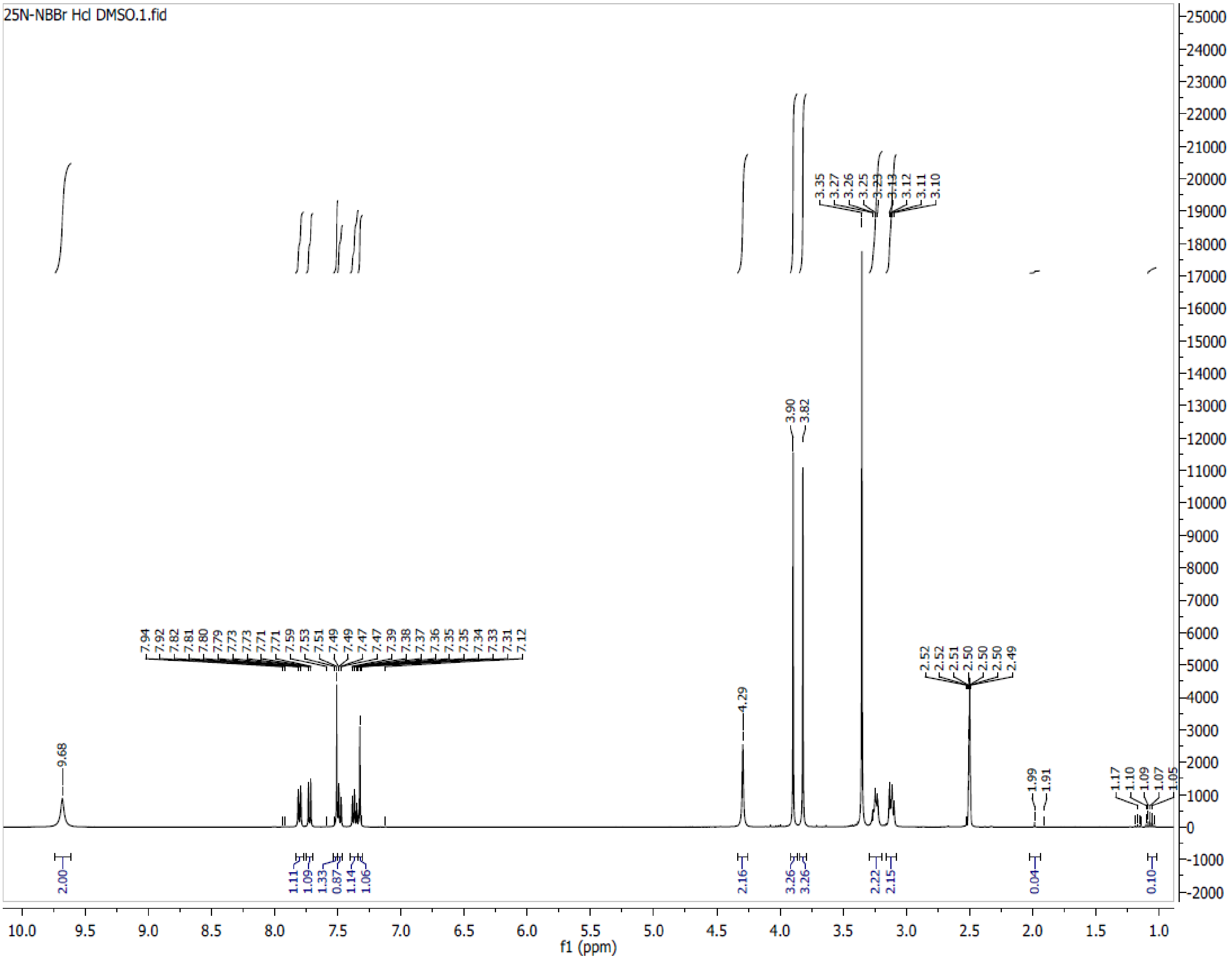

**Spectra 9. $^{1}$H NMR of 25N-NBI (10) HCl**

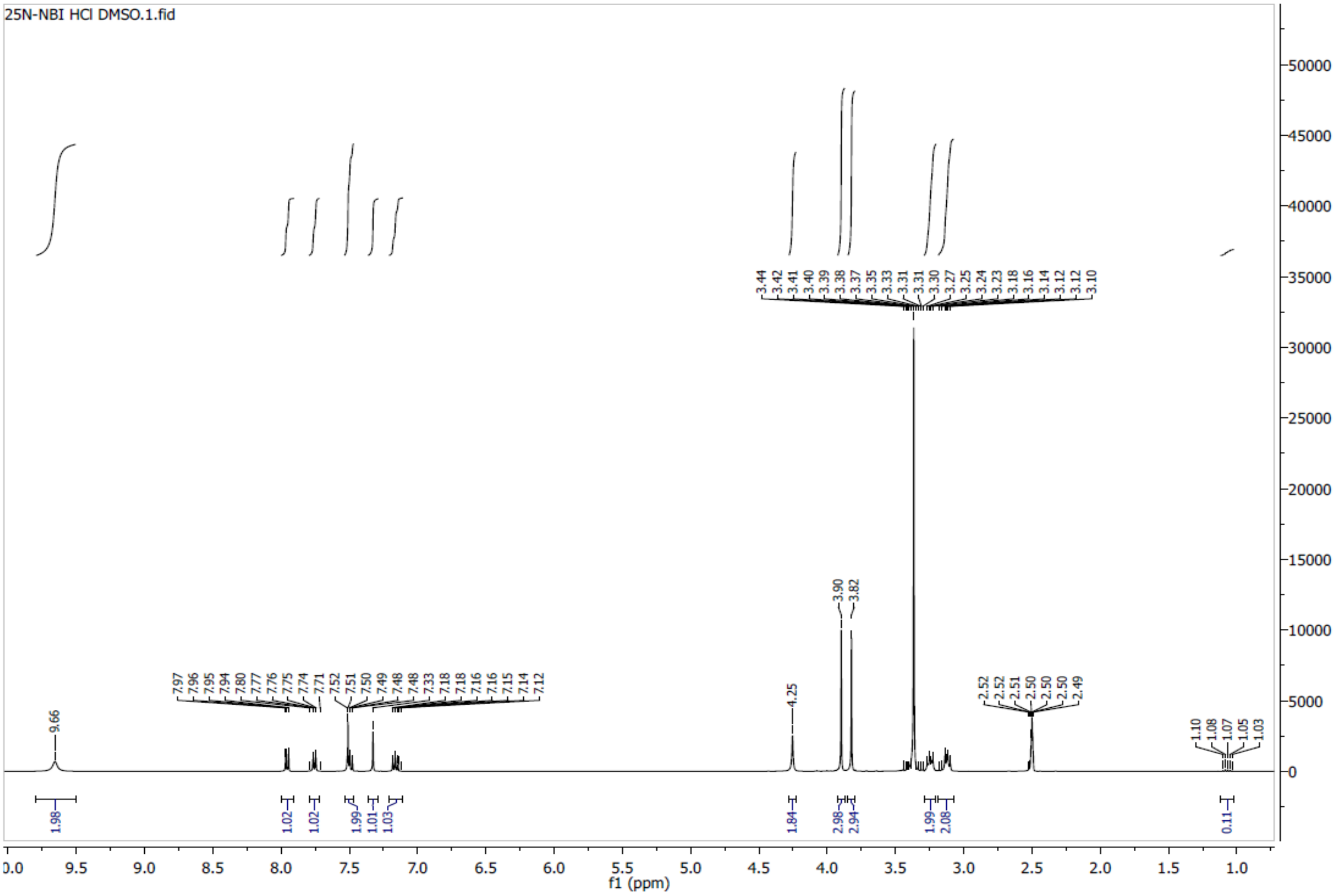

**Spectra 10**. $^{1}H$ NMR of 25N-NBOCF$_2$H (11) HCl

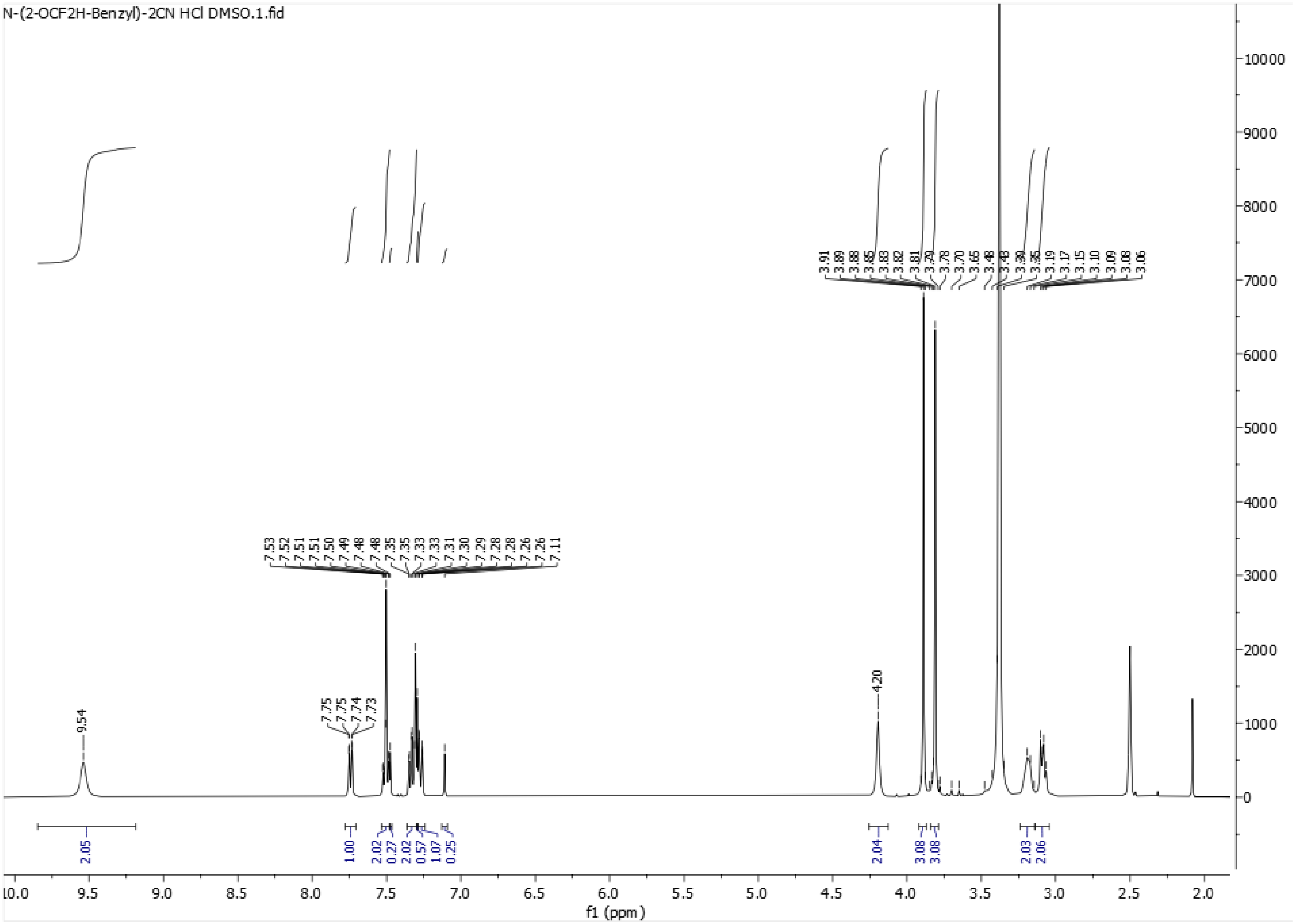

**Spectra 11.** $^1$**H NMR of 25N-NBOCF**$_3$ **(12) HCl**

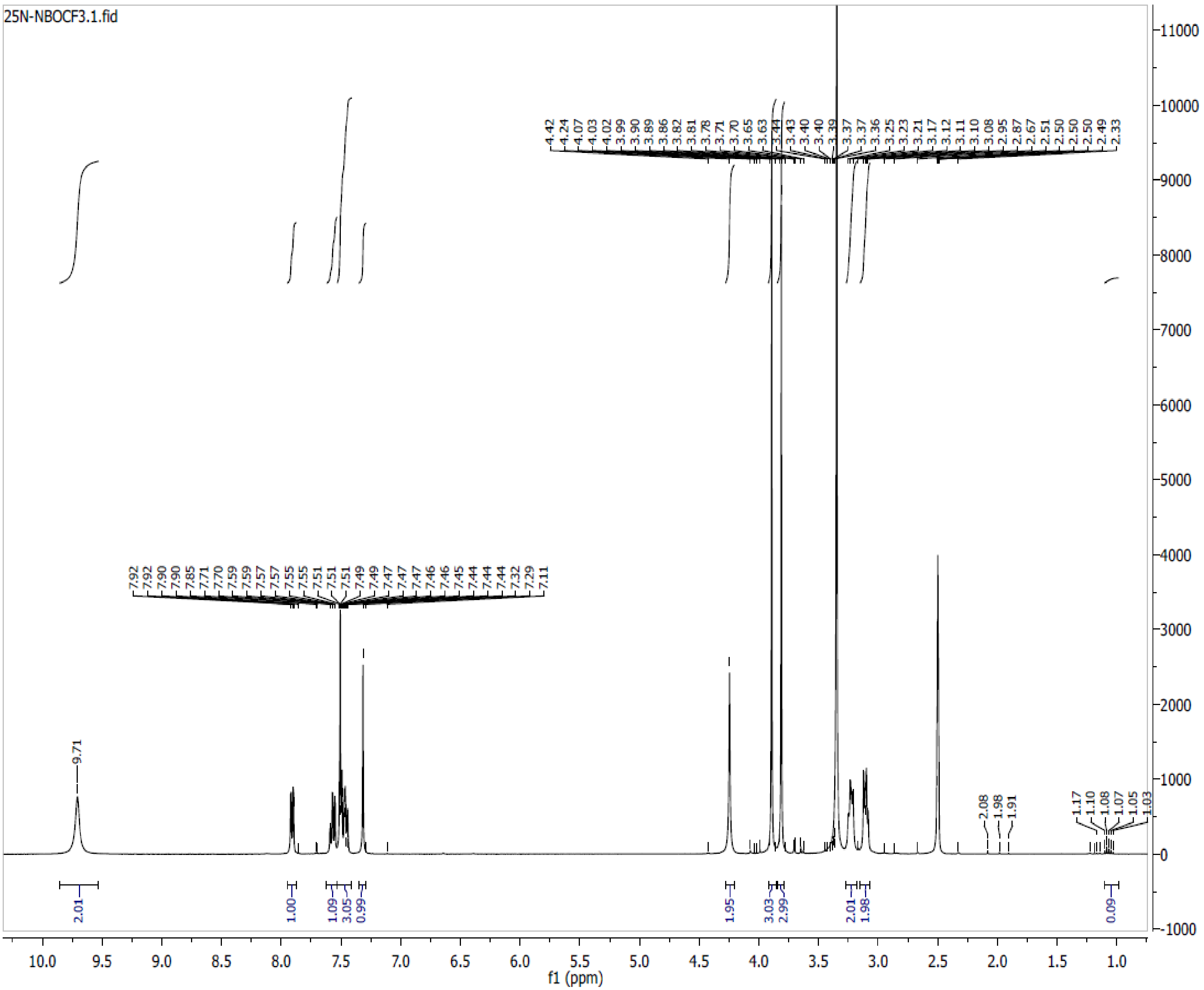

**Spectra 12. $^{1}$H NMR of 25N-NBMDF$_2$ (13) HCl**

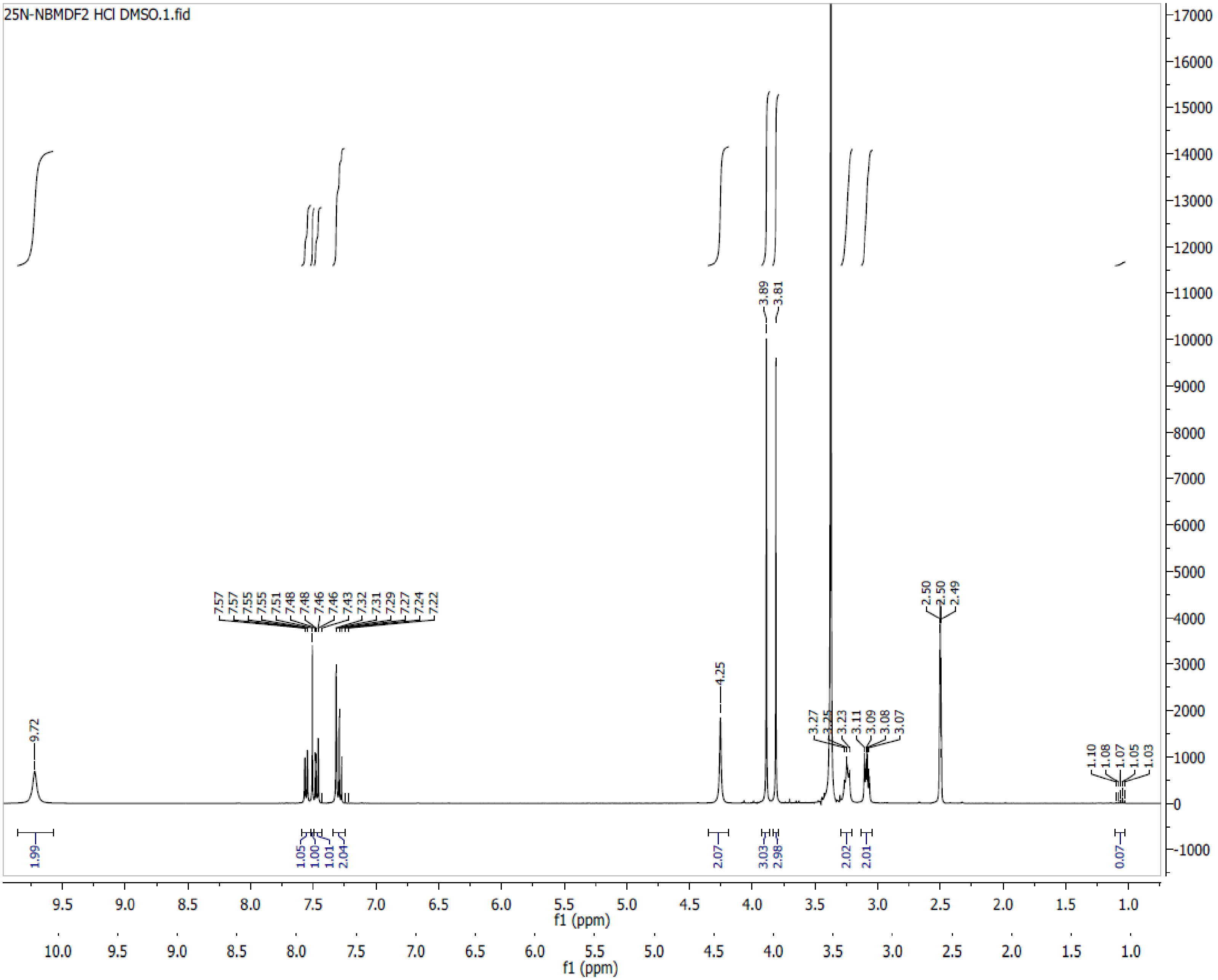

**Spectra 13. $^{1}$H NMR of 25N-NBCF$_3$ (14) HCl**

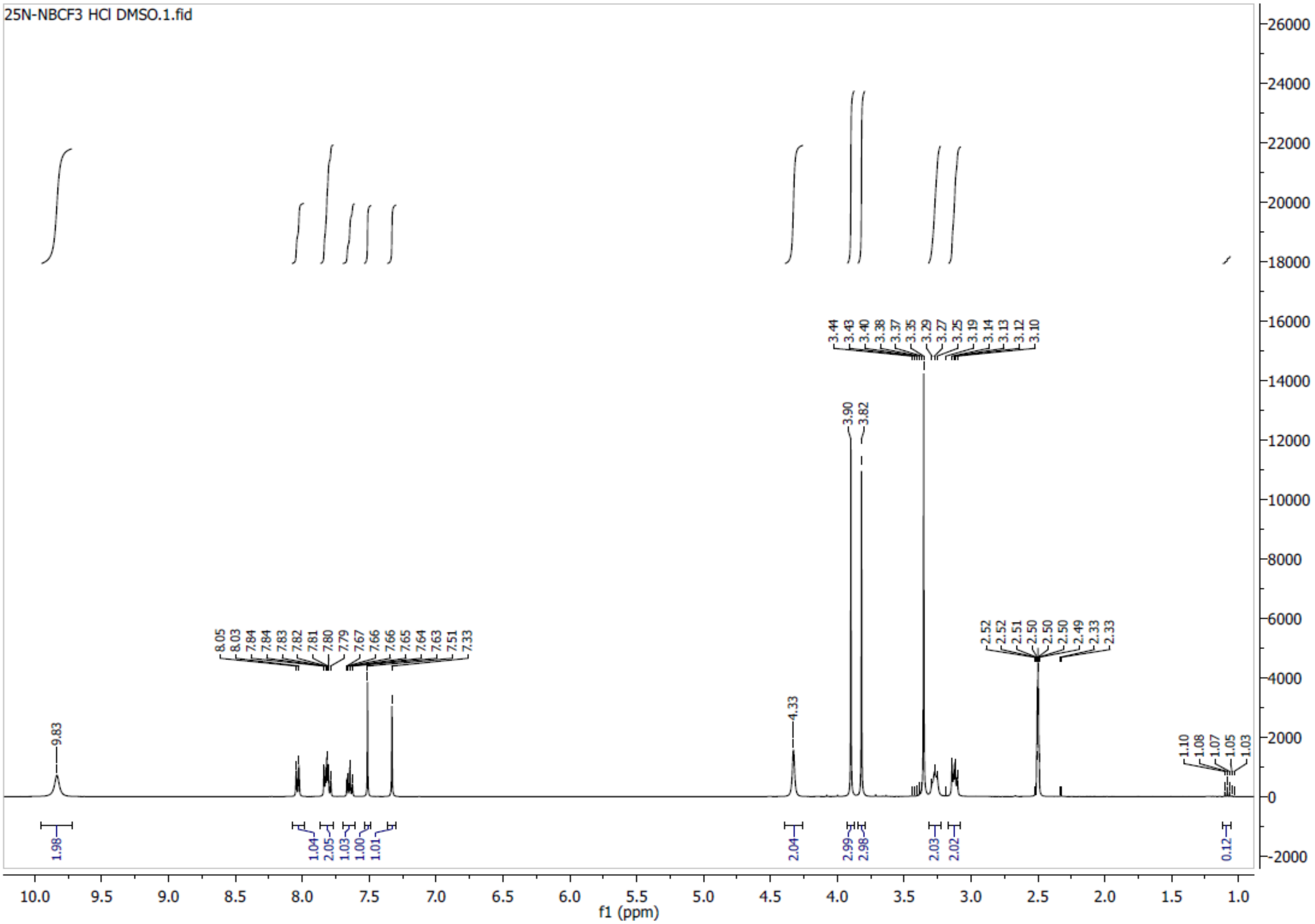

**Spectra 14. $^1$H NMR of 25N-NBNO$_2$ (15) HCl**

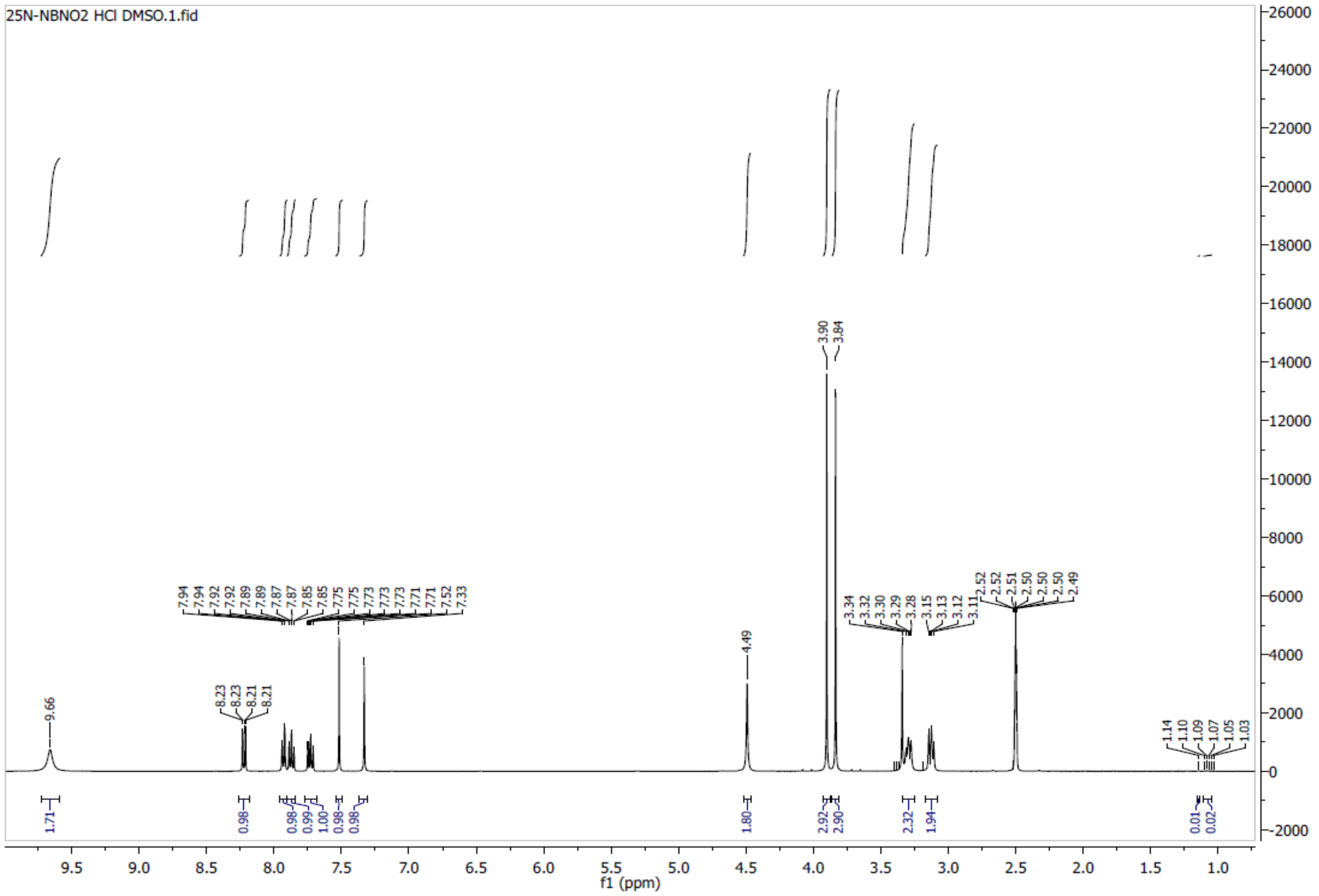

**Spectra 15**. **$^{1}$H NMR of 25N-N1-Nap (16) HCl**

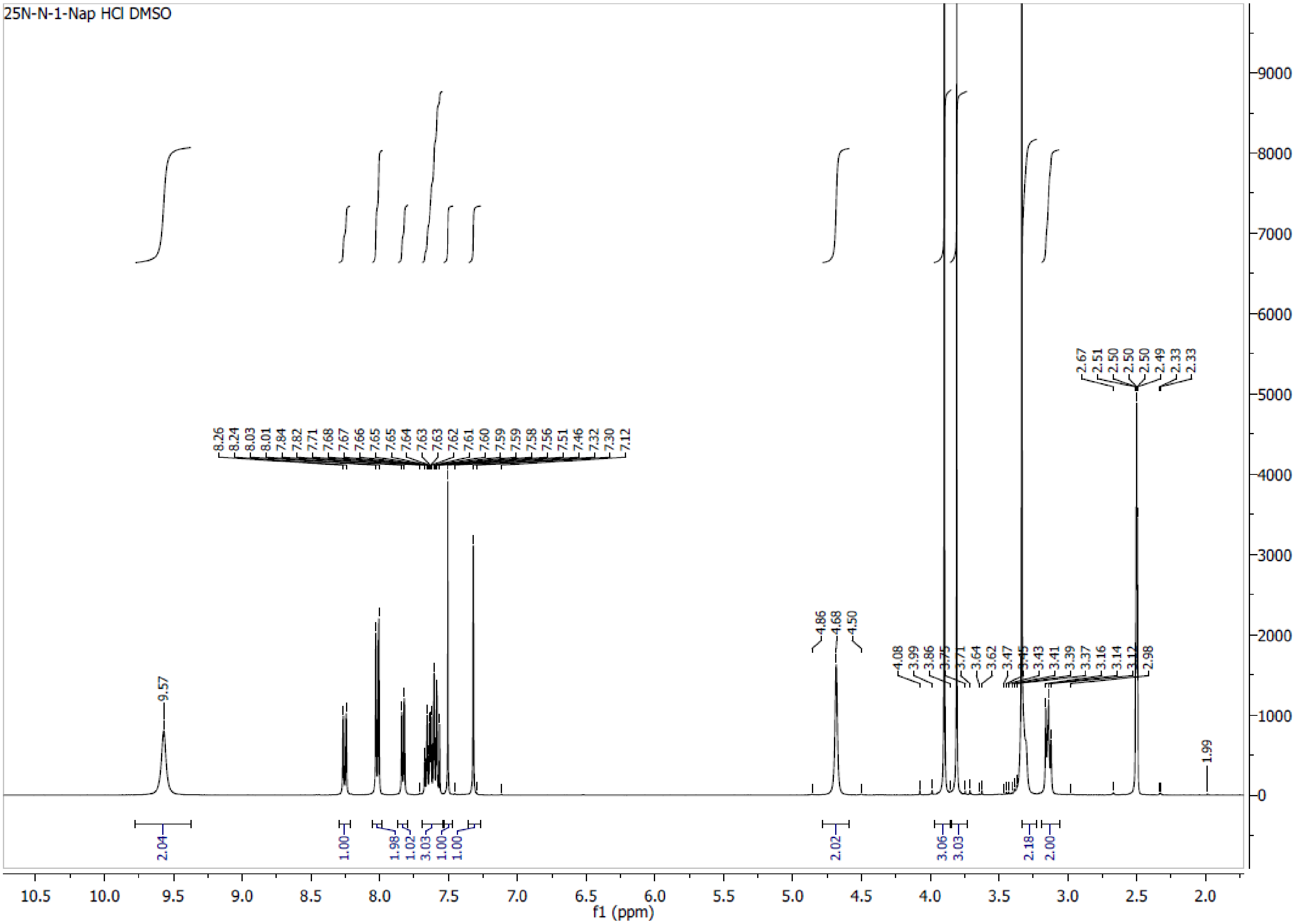

**Spectra 16. $^1$H NMR of 25N-NBPh (17) HCl**

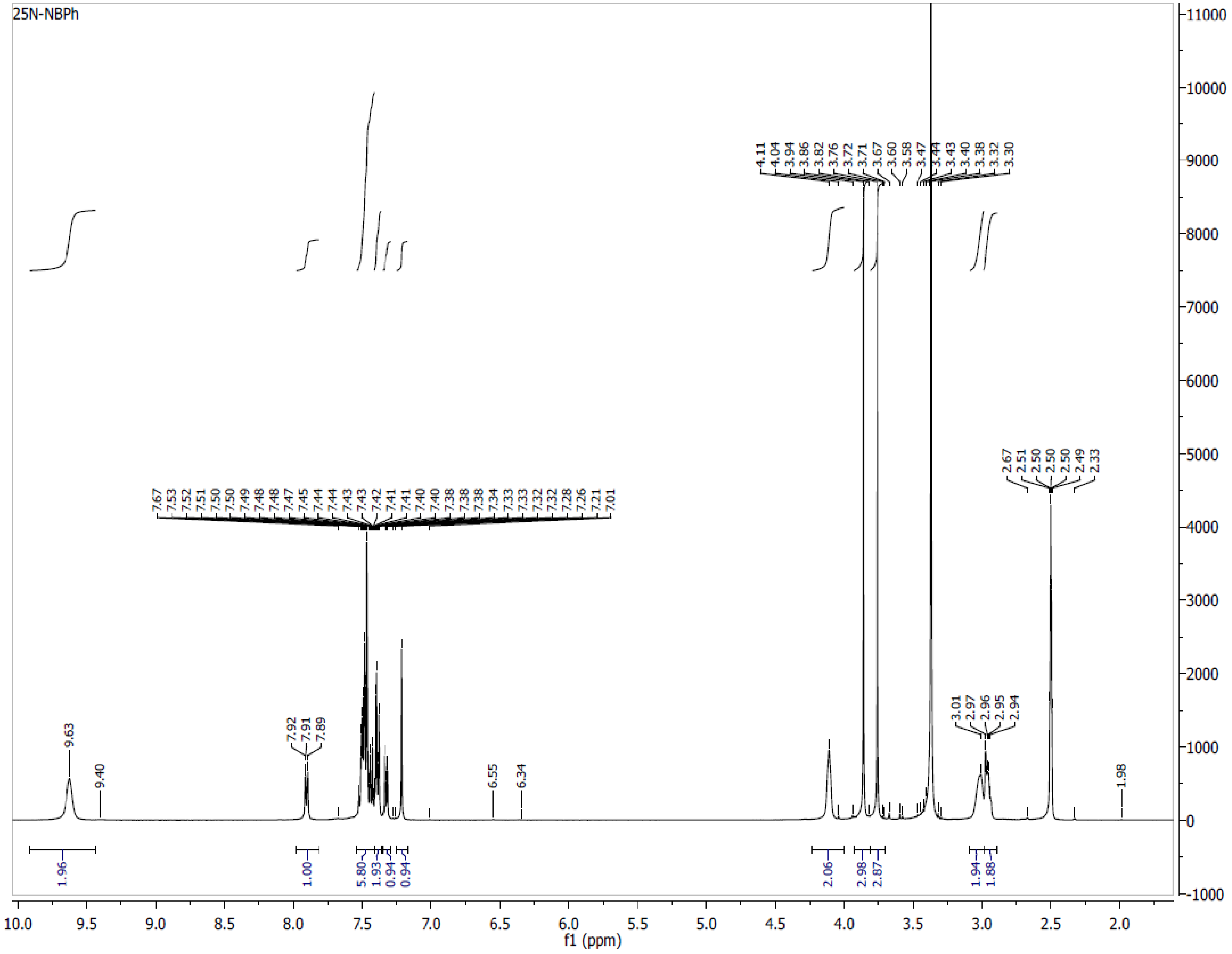

**Spectra 17. $^{1}$H NMR of 25N-NB-2-OH-3-Me (18) HCl**

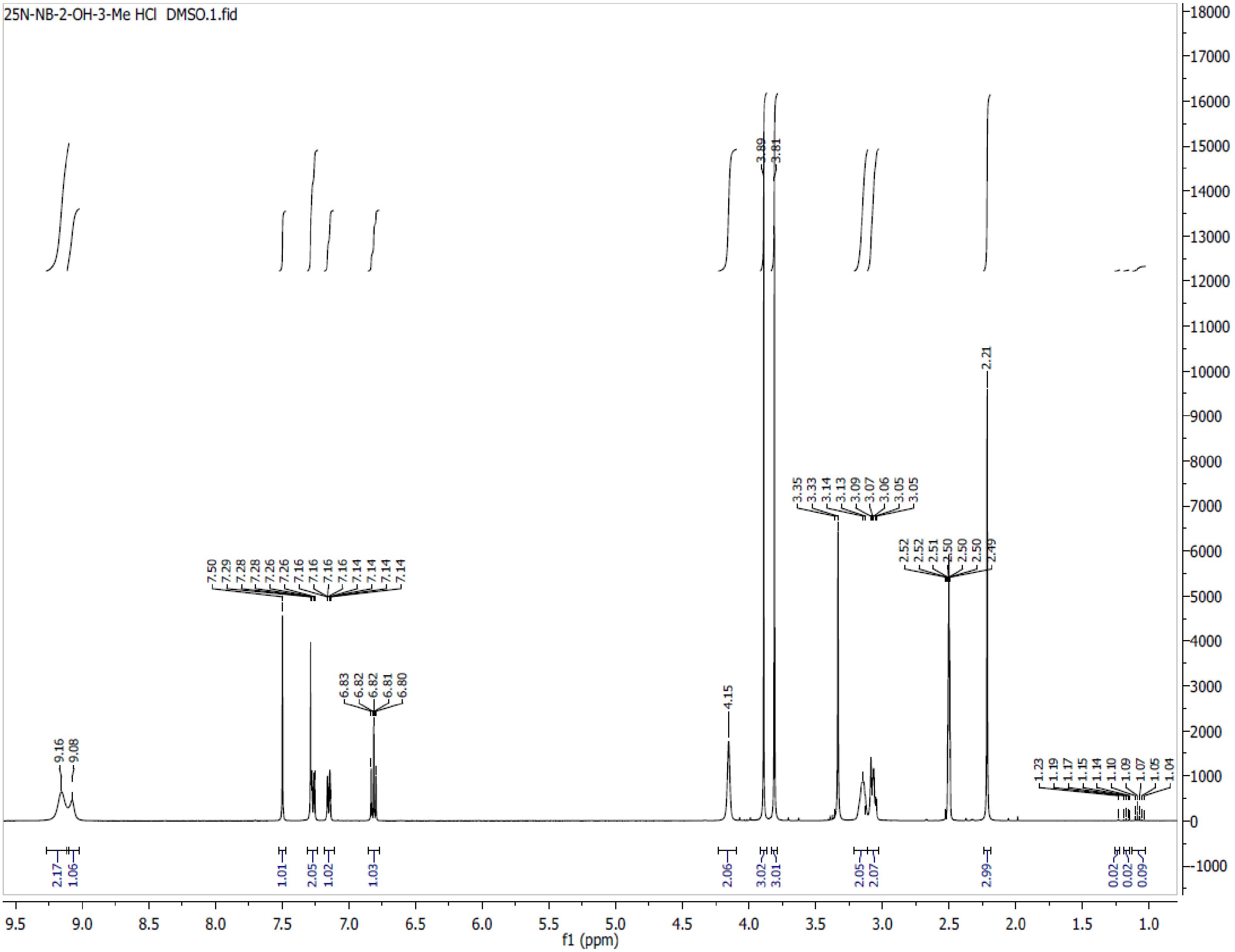

**Spectra 18. $^{1}$H NMR of 25N-NB-2-MeO-3-F (19) HCl**

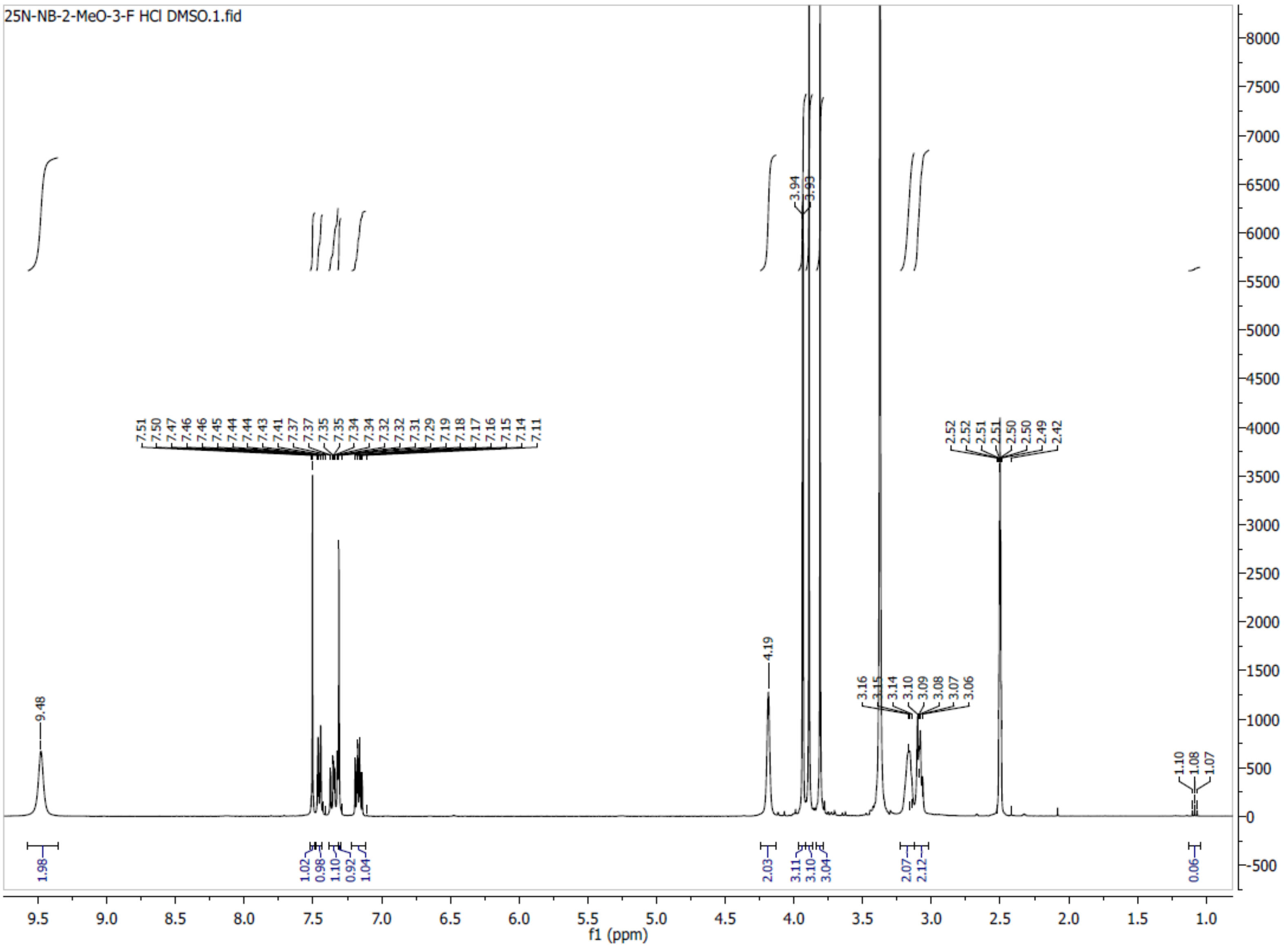

**Spectra 19**. **$^{1}$H NMR of 25N-NB-2,5-DiMeO (20) HCl**

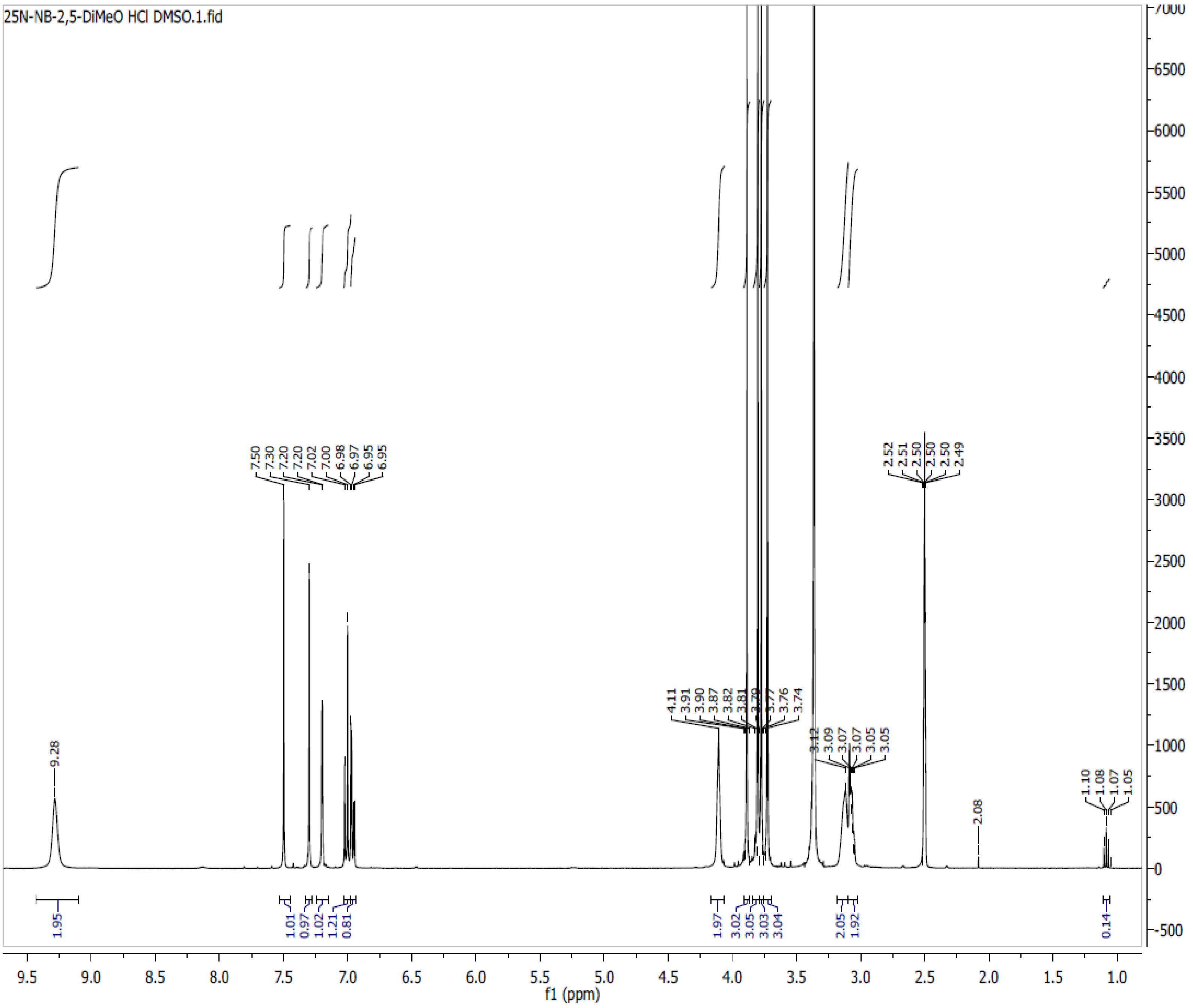

**Spectra 20. ¹H NMR of 25N-NB-3-OH (21) HCl**

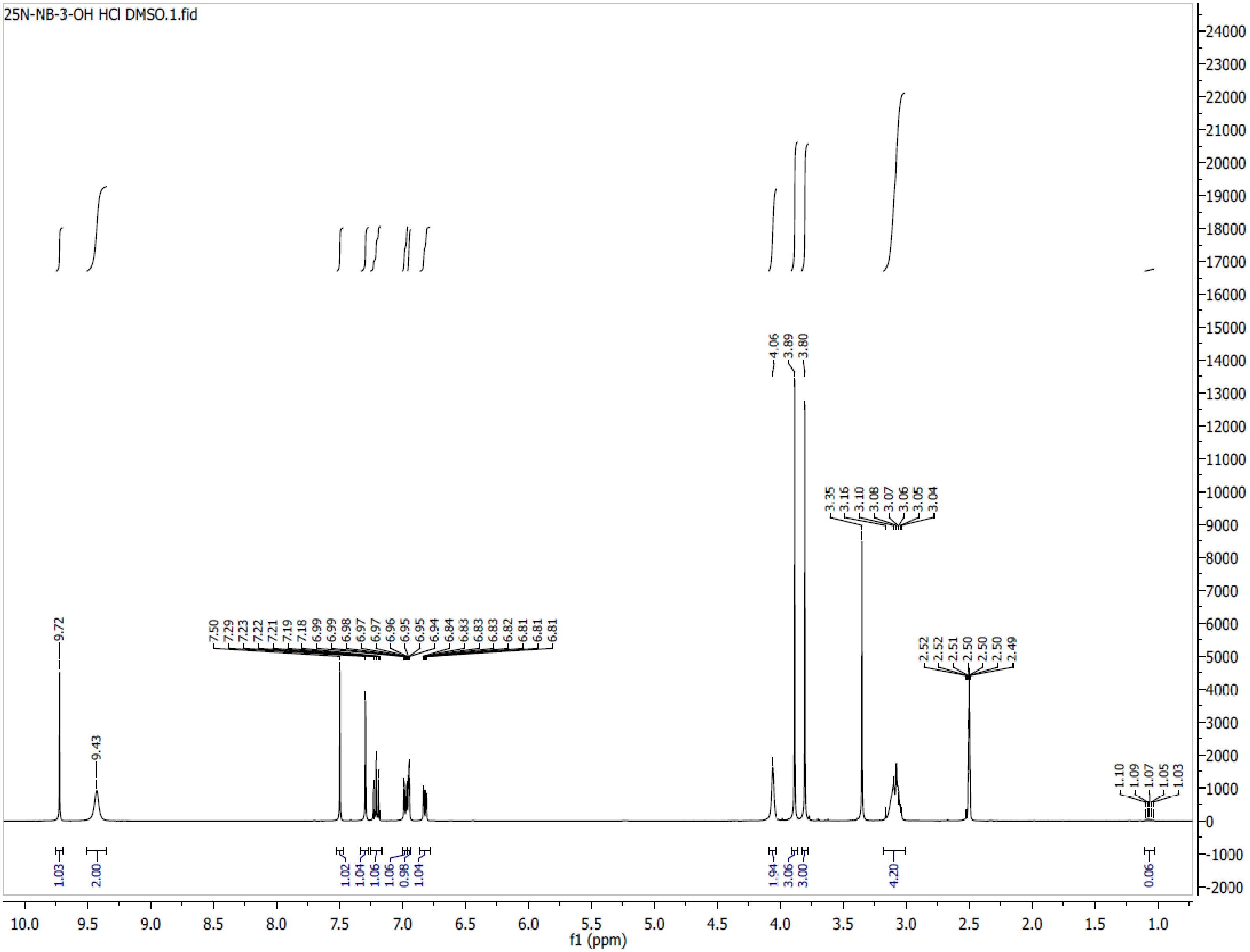

**Spectra 21. $^{1}$H NMR of 25N-NB-3-Me (22) HCl**

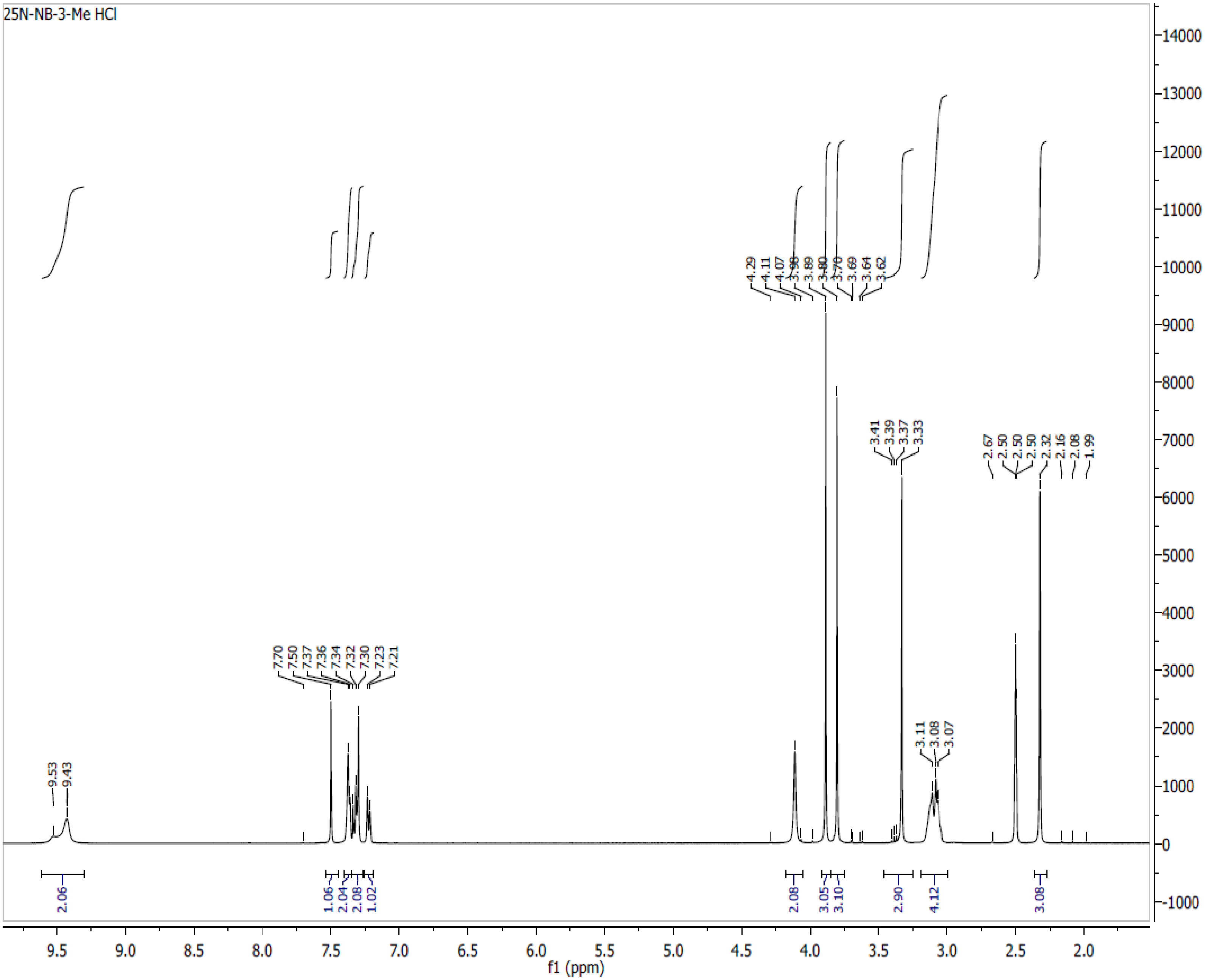

**Spectra 22. $^{1}$H NMR of 25N-NB-4-Me (23) HCl**

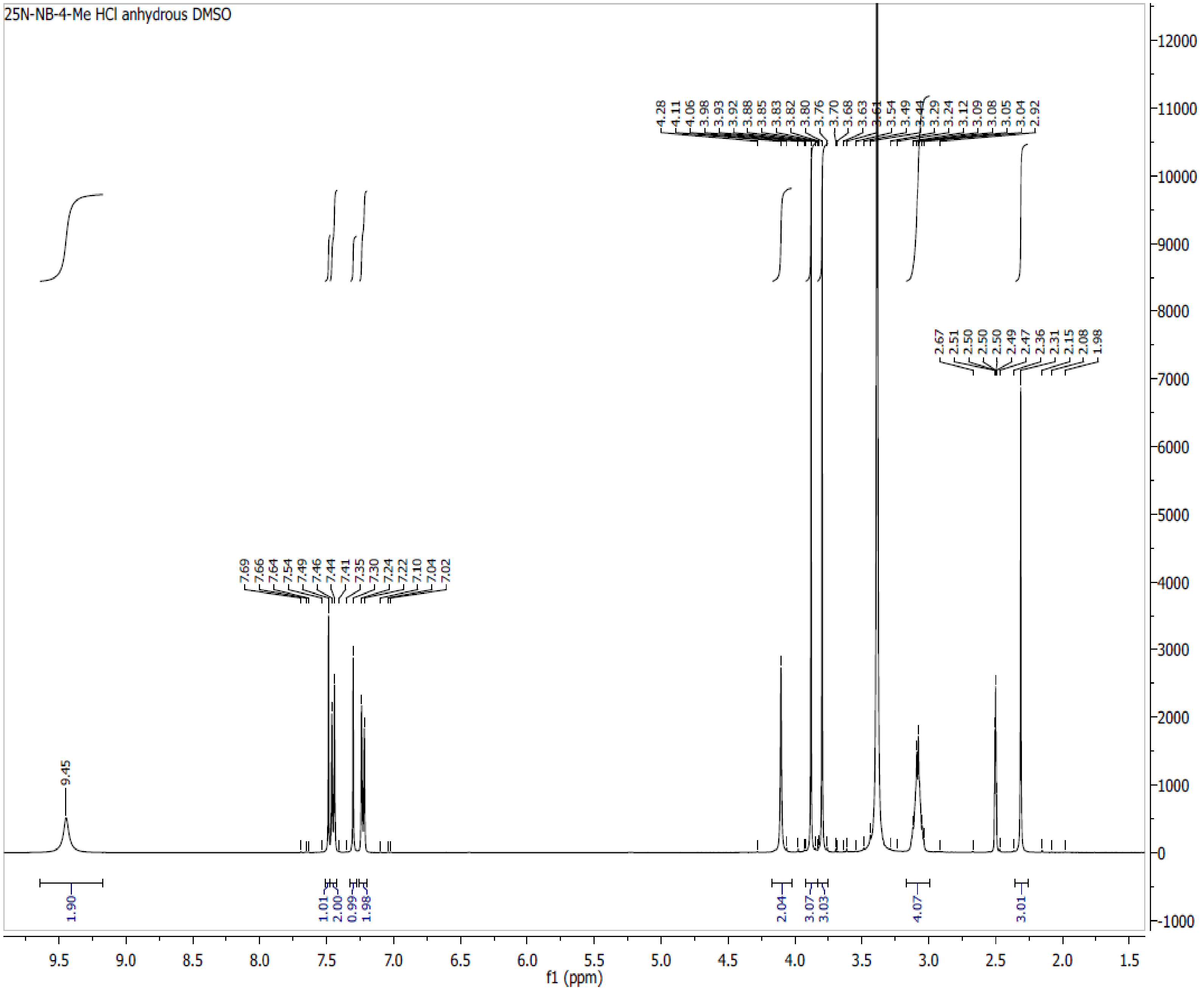

**Spectra 23. $^1$H NMR of 25N-NB-3-F (24) HCl**

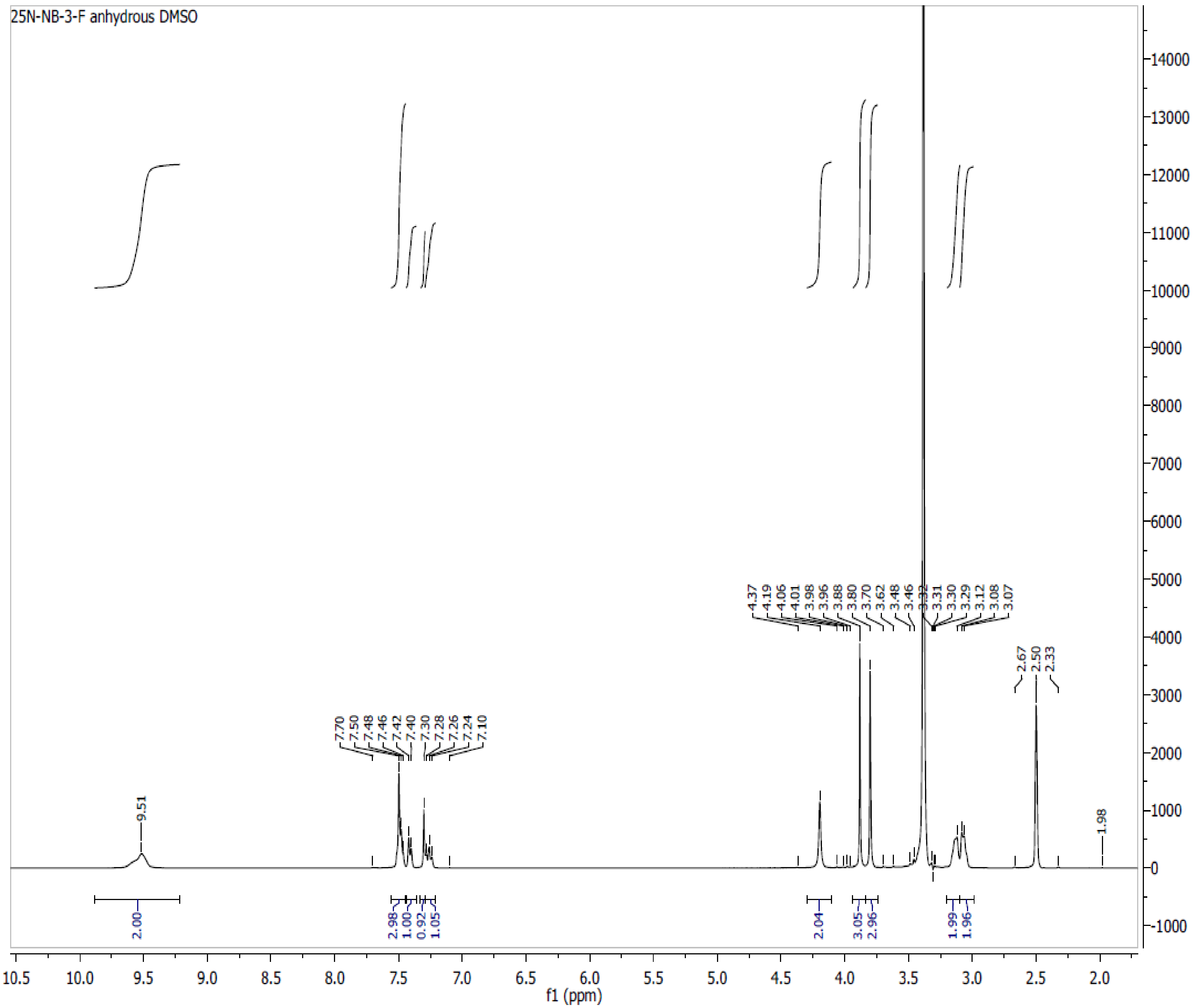

**Spectra 24. $^{1}$H NMR of 25N-NB-4-F (25) HCl**

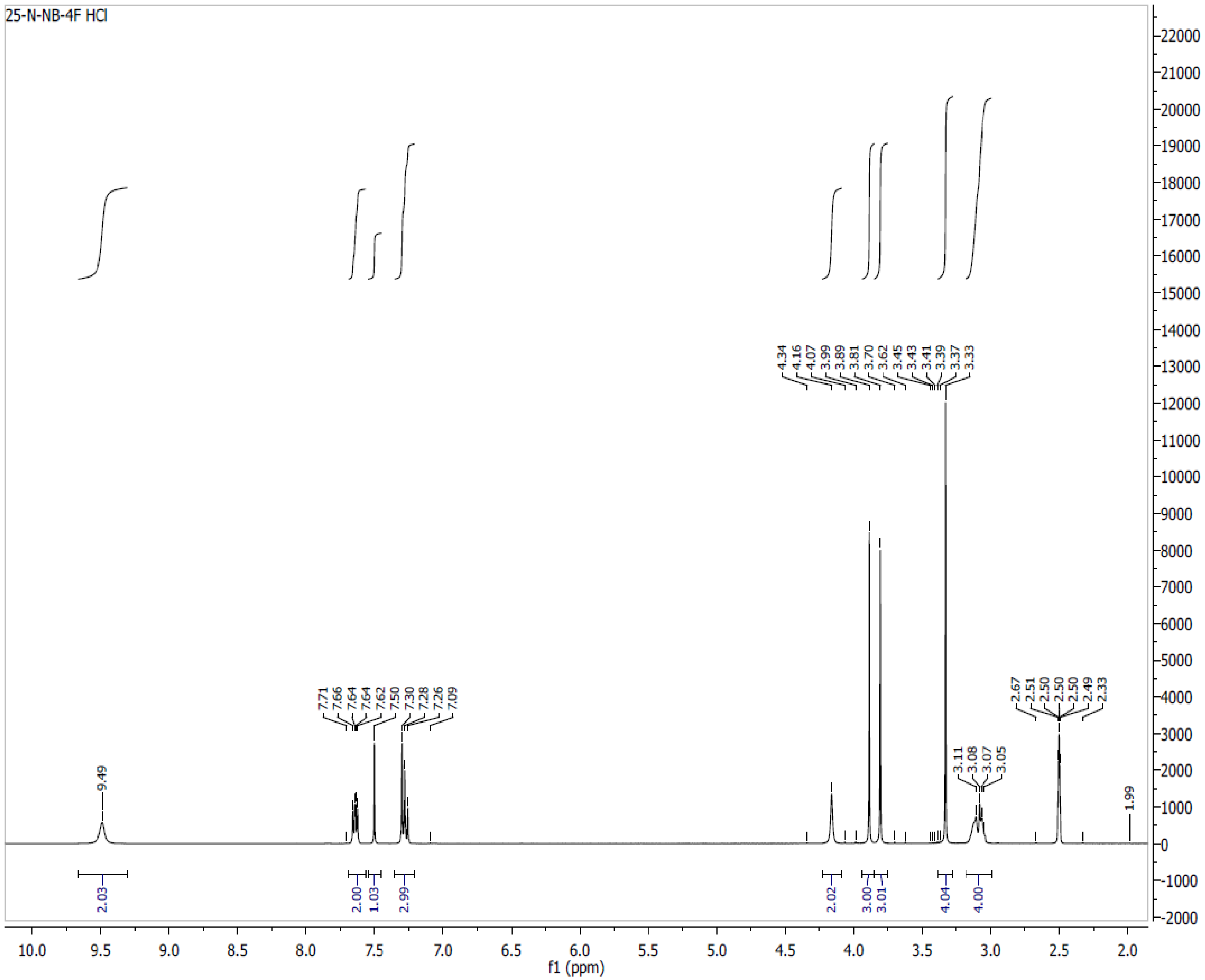

**Spectra 25**. **$^{1}$H NMR of 25D-N1-Nap (26) HCl**

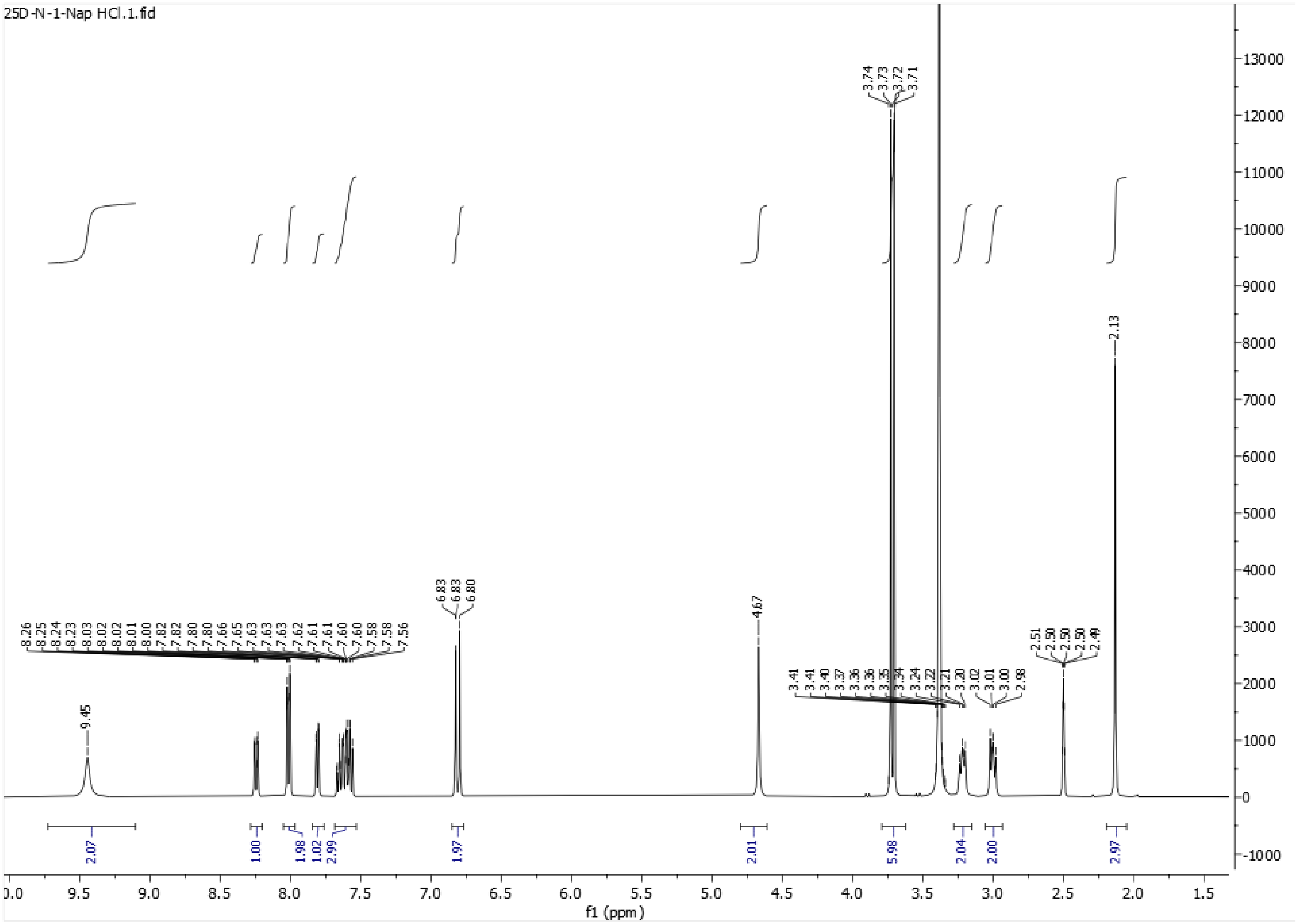

### Spectra 26. $^{1}$H NMR of 25D-NBPh (27) HCl

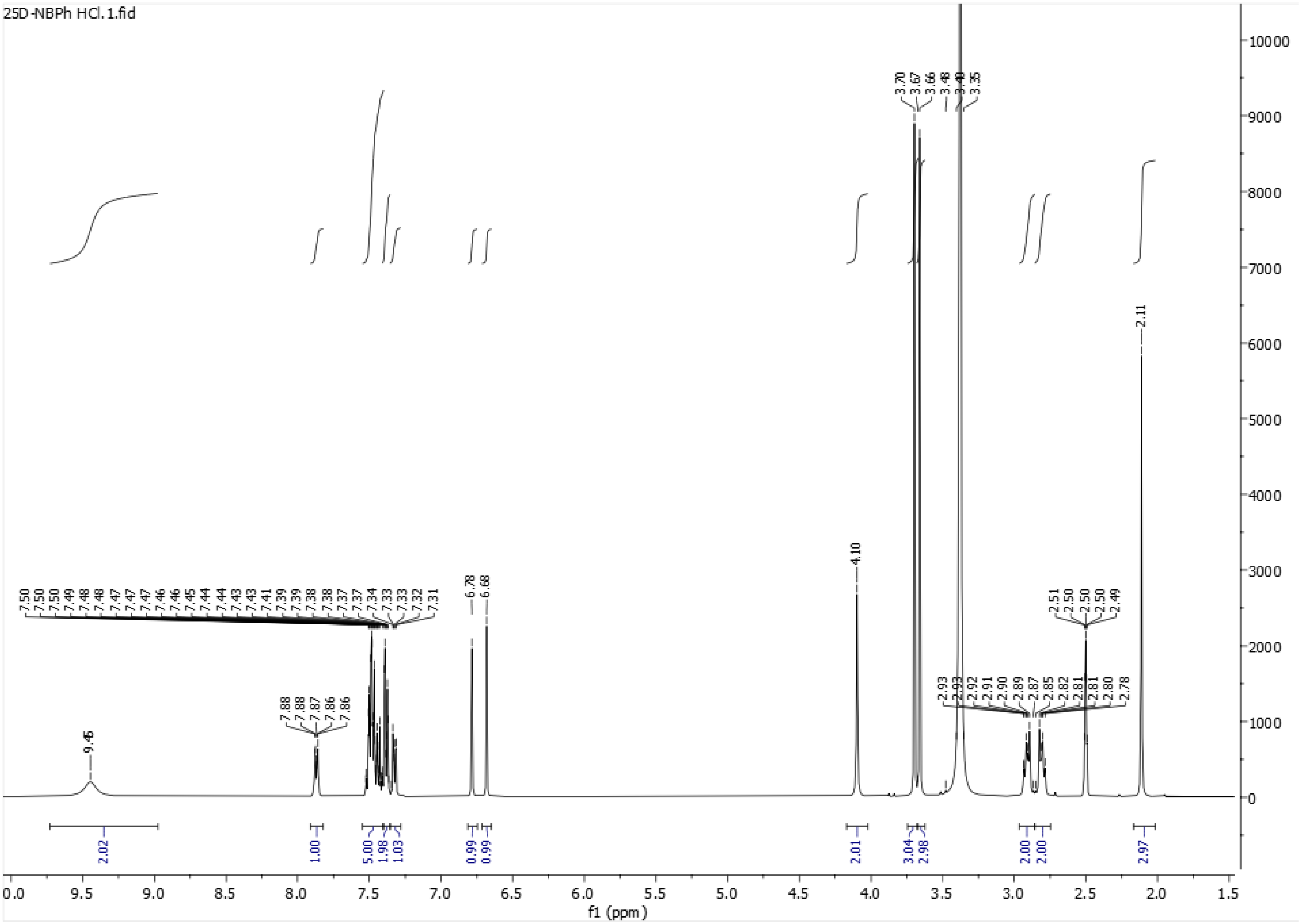

**Spectra 27. $^{1}$H NMR of 25O-N1-Nap (28) HCl**

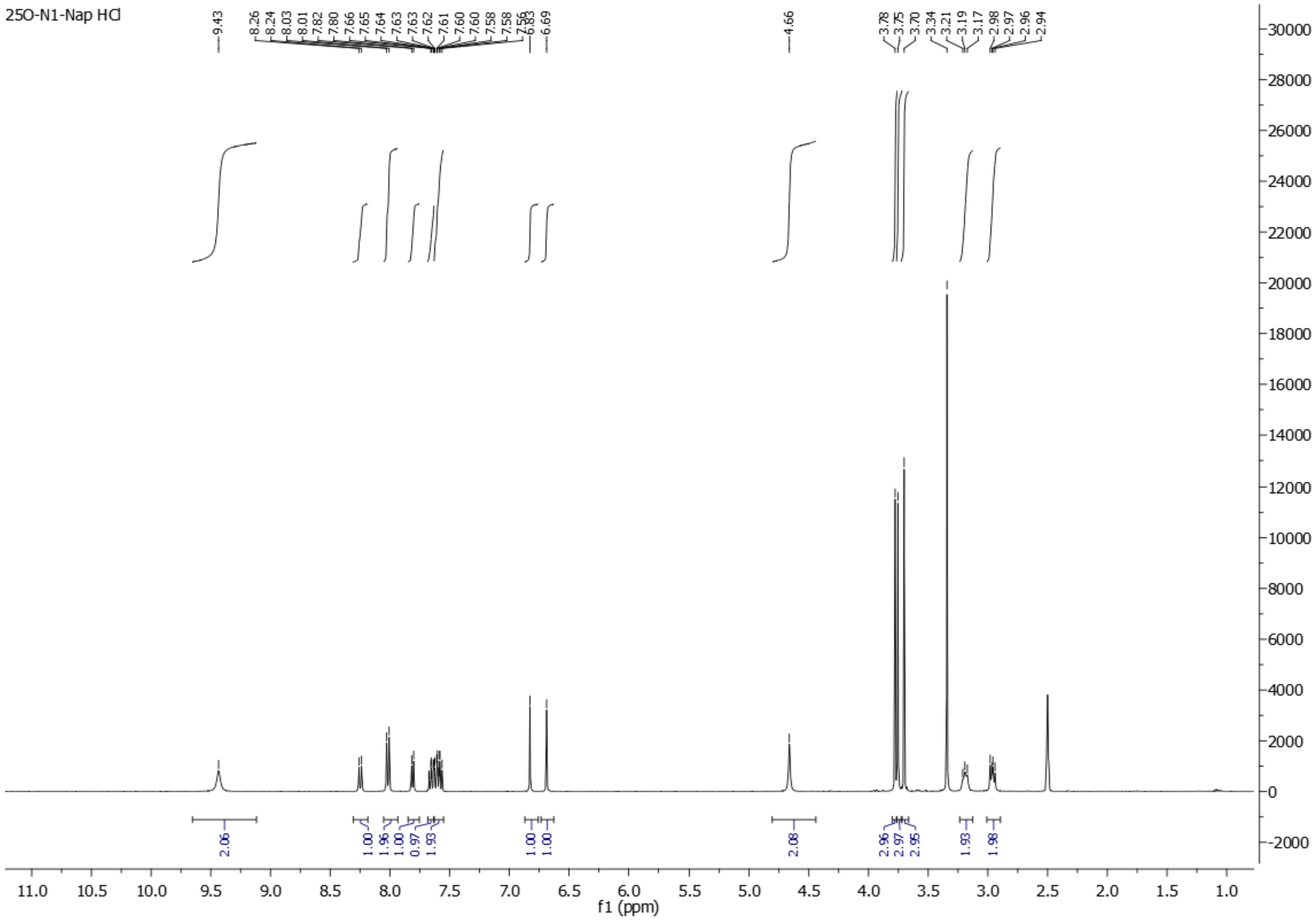

### Spectra 28. $^{1}$H NMR spectrum of 2C2-N1-Nap (29) HCl

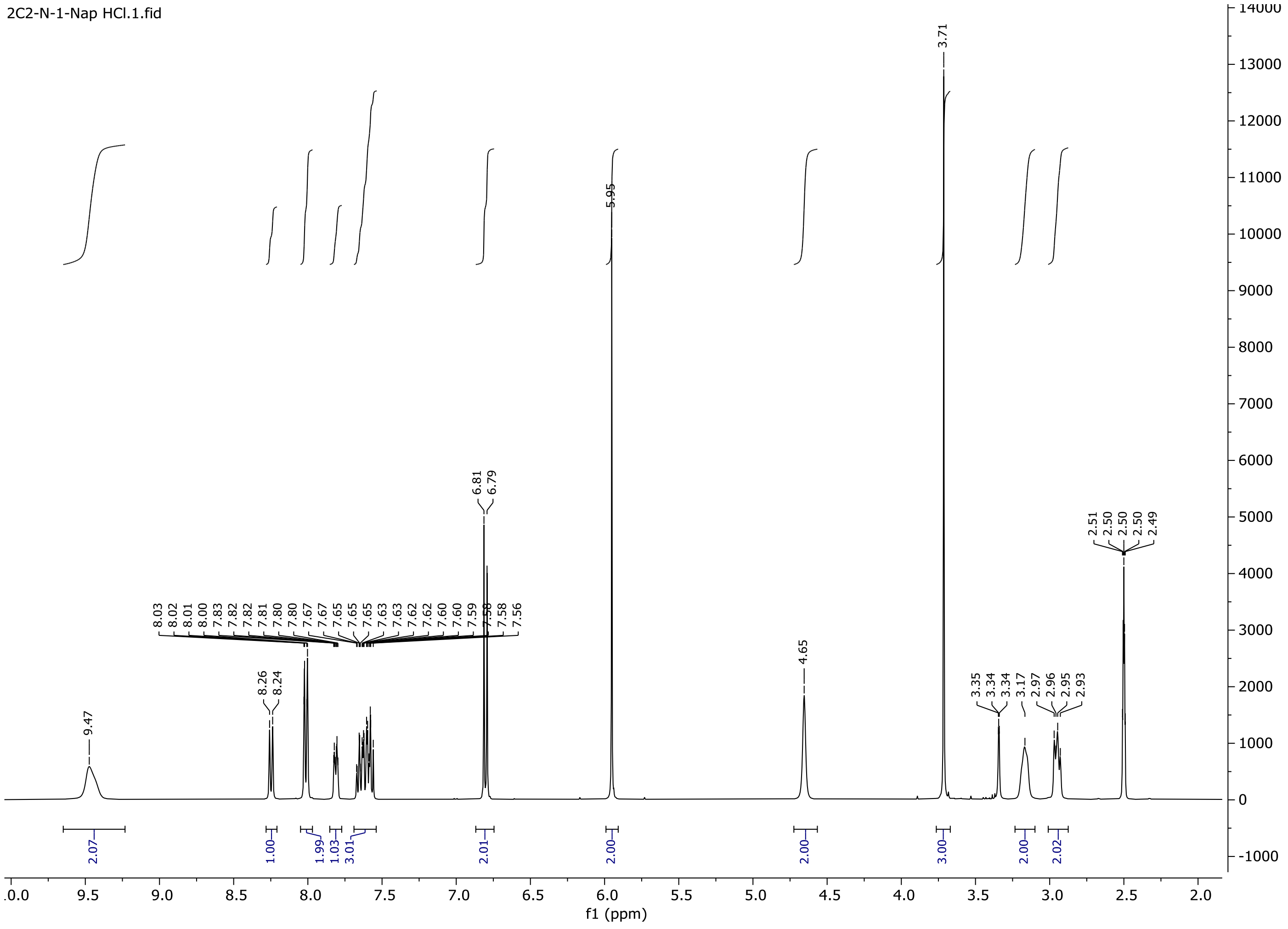

**Spectra 29.** $^{1}$H NMR spectrum of **Ψ-DOB-N1-Nap (30) HCl**

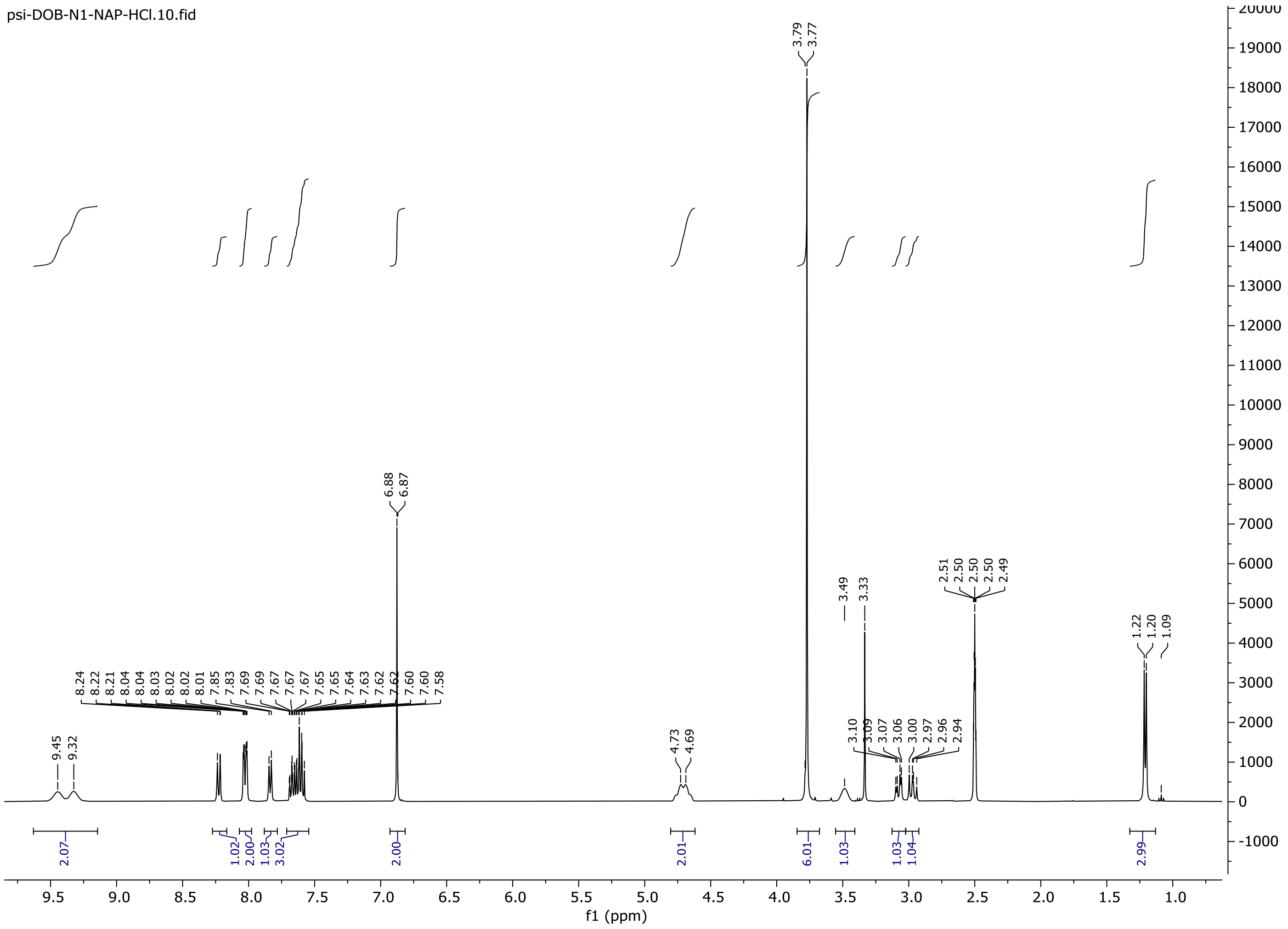

### Spectra 30. $^{1}$H NMR of 2C2-NBOMe (31) HCl

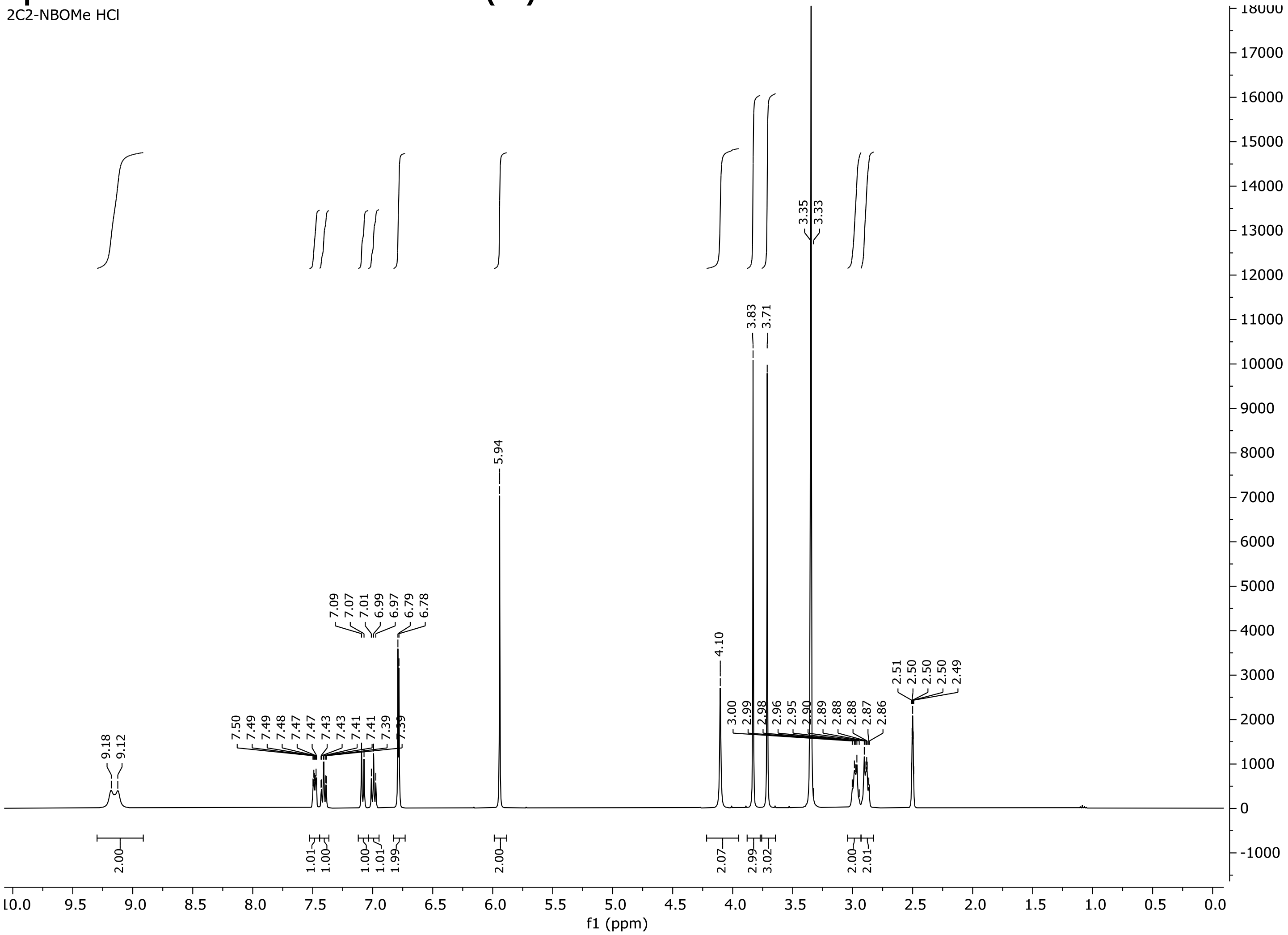

### Spectra 31. $^1$H NMR of 25O-NBOMe (32) HCl

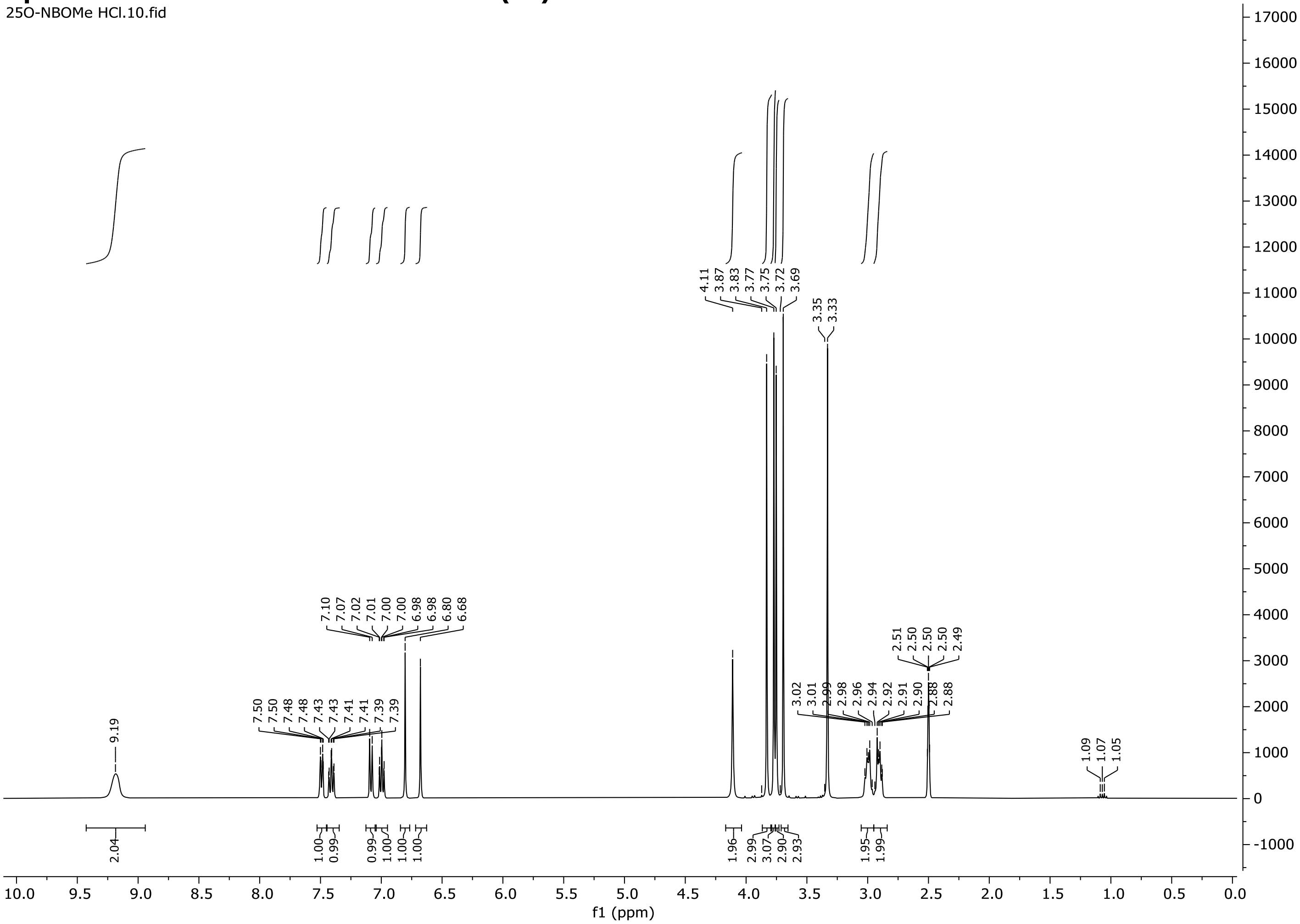

### Spectra 32. $^{1}$H NMR of 25O-NBcP (33) HCl

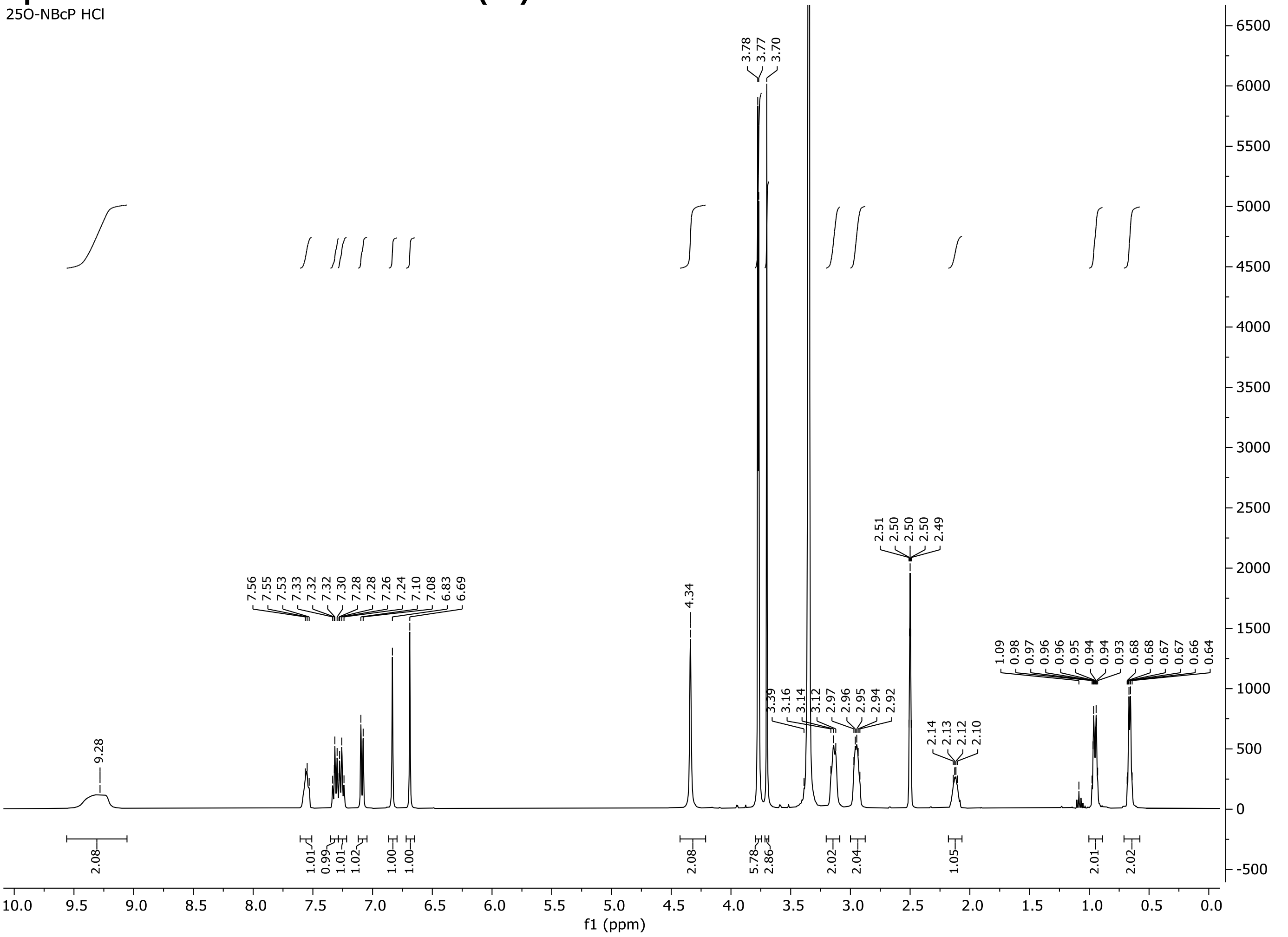

### Spectra 33. $^{1}$H NMR of 25O-NB-3-I (34) HCl

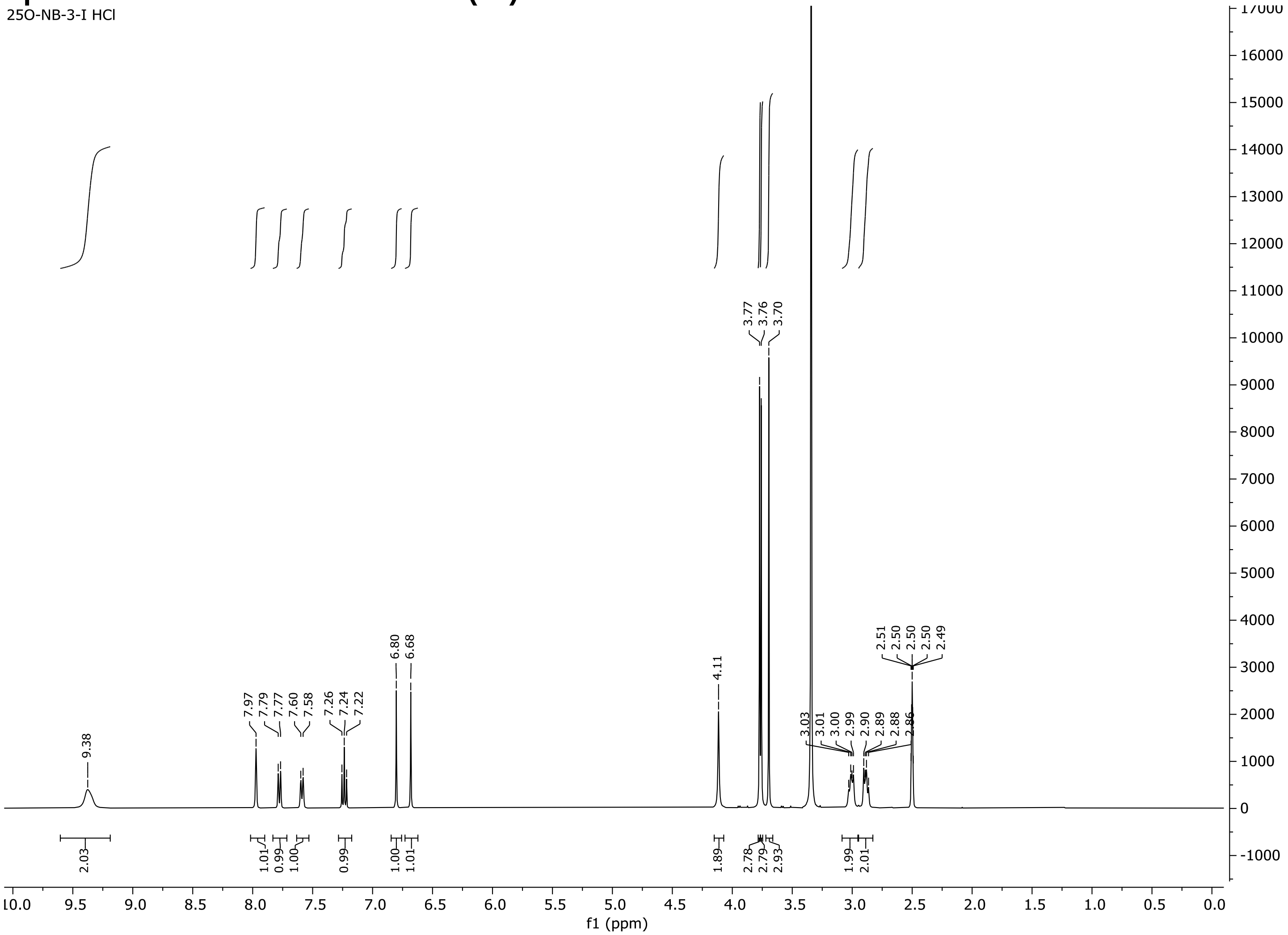

### Spectra 34. $^1$H NMR of 25O-NBPh-10'-OH (35) HCl

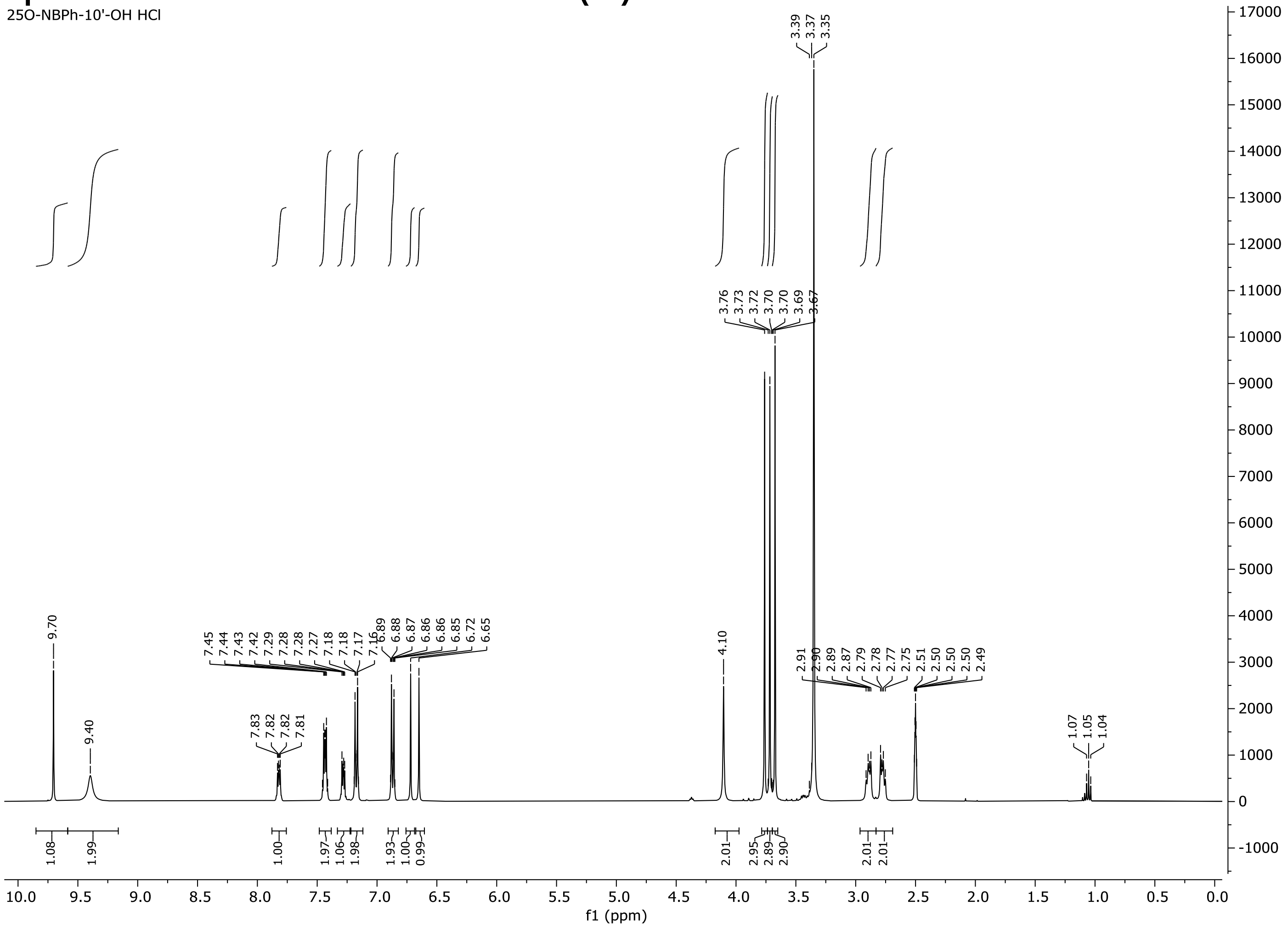

### $^{13}$C NMR Spectra of Synthesized N-Benzyl Phenethylamines

#### Spectra 35. $^{13}$C NMR of 25N-NB (2) HCl

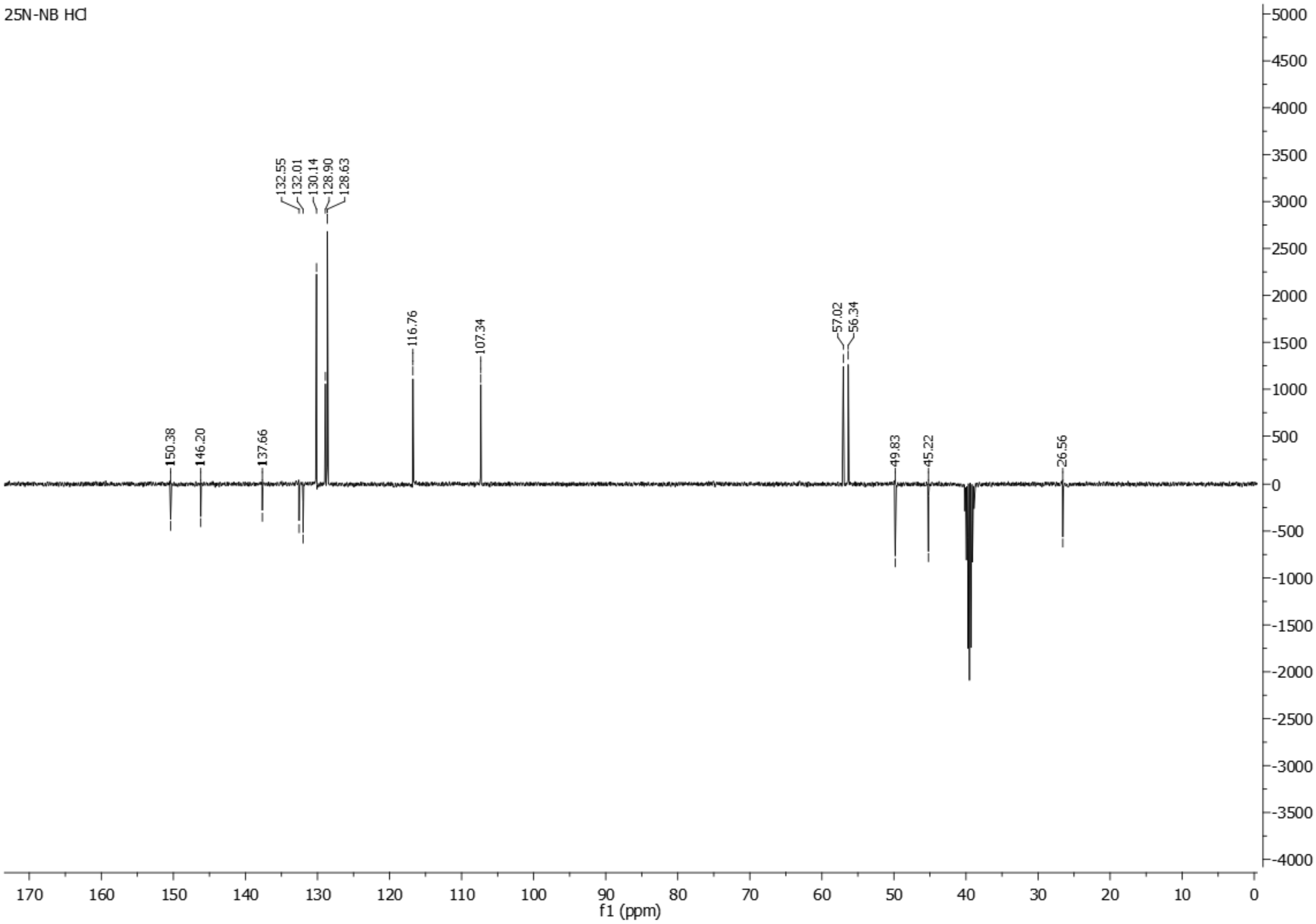

**Spectra 36. $^{13}C$ NMR of 25N-NBOH (3) HCl**

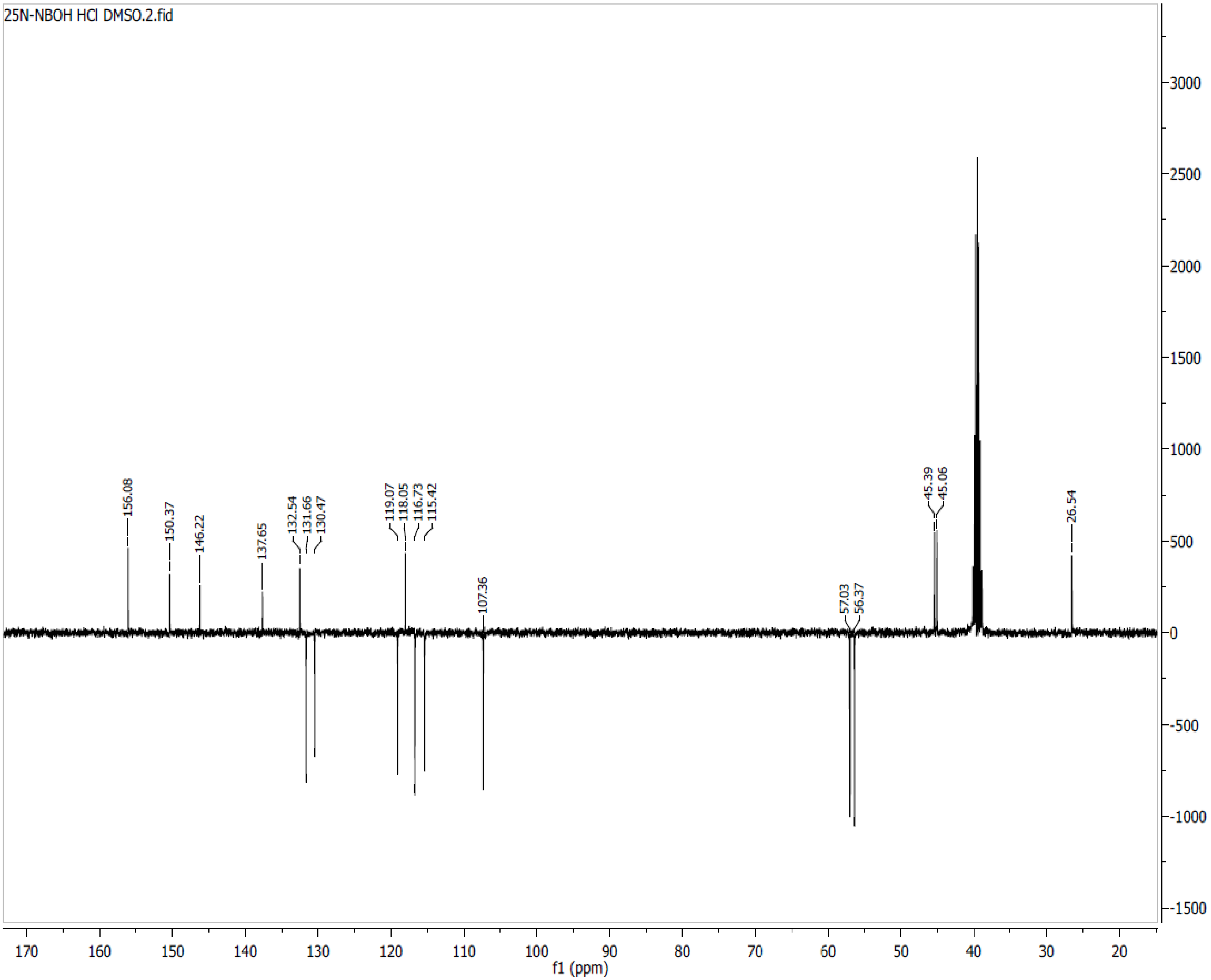

**Spectra 37. $^{13}$C NMR of 25N-NBOMe (4) HCl**

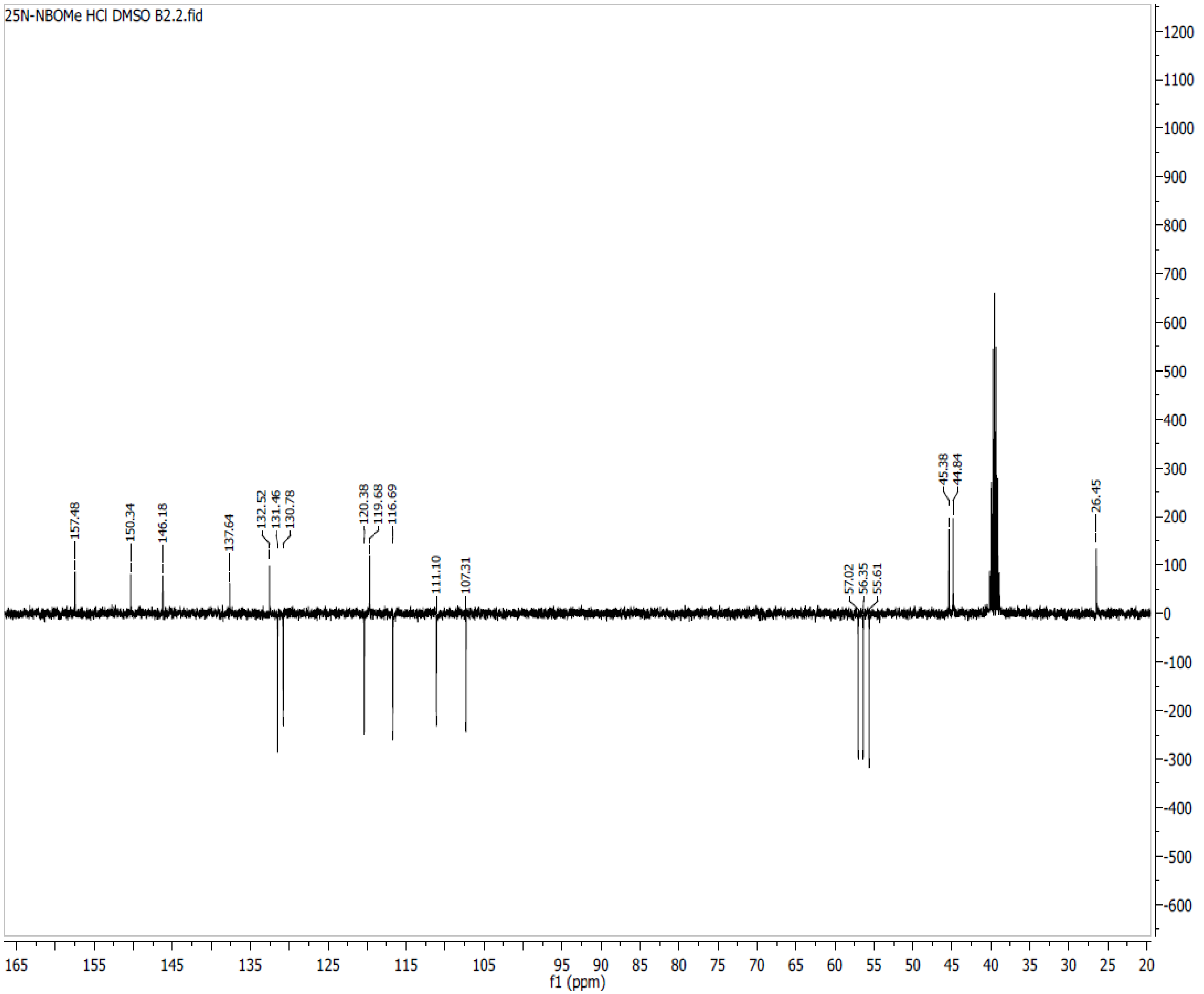

**Spectra 38. $^{13}C$ NMR of 25N-NBOEt (5) HCl**

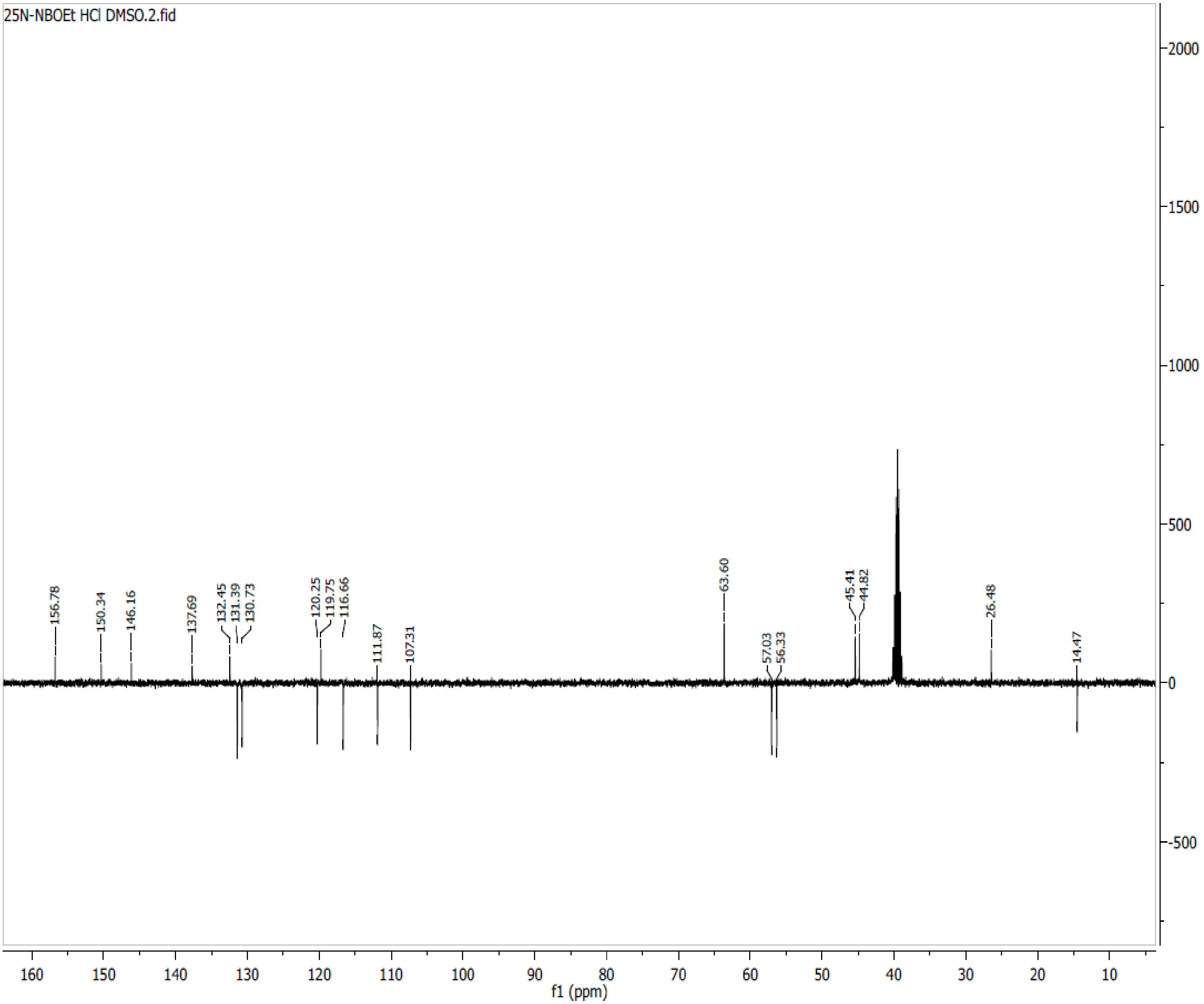

**Spectra 39. $^{13}C$ NMR of 25N-NBMe (6) HCl**

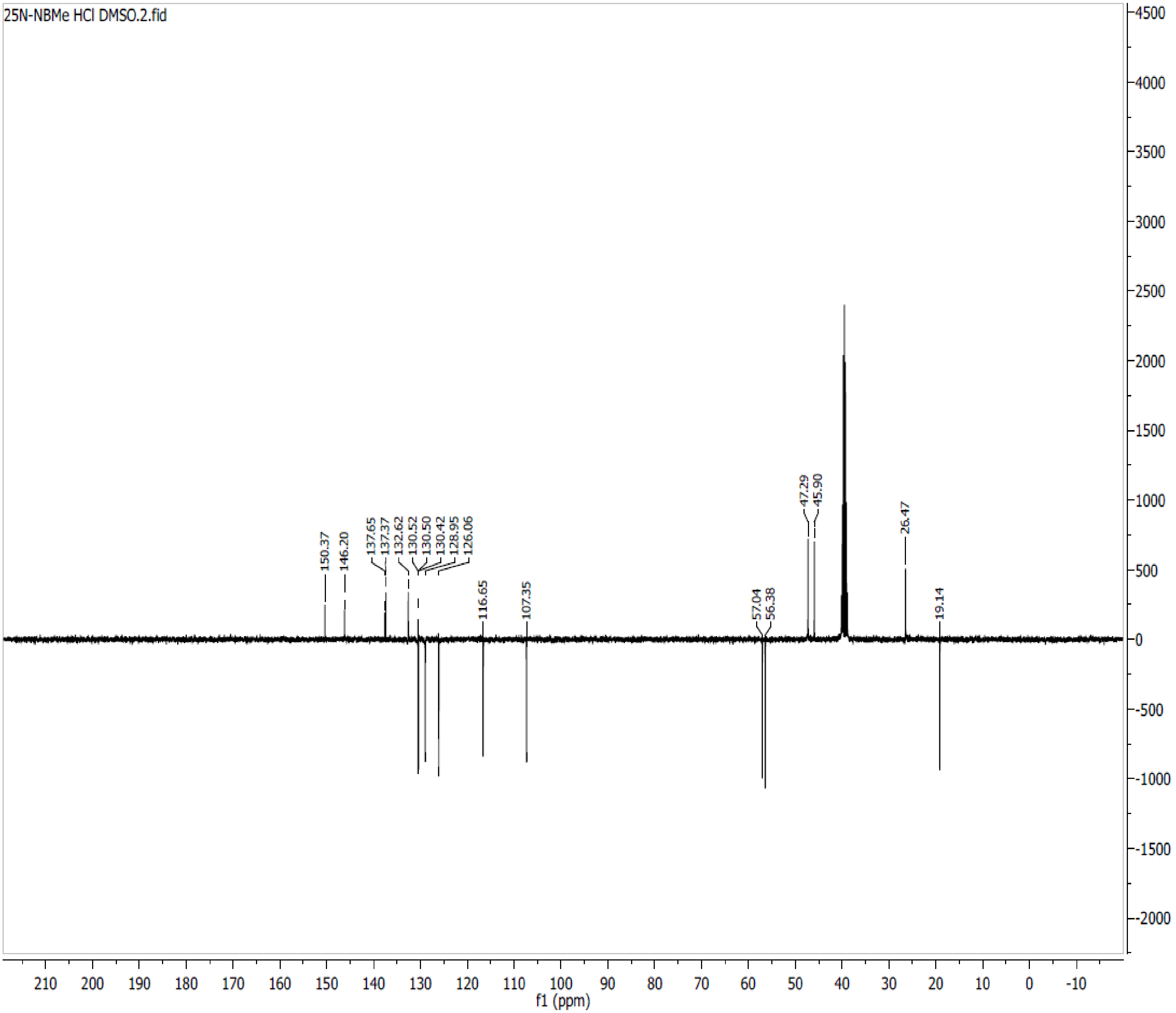

**Spectra 40. $^{13}$C NMR of 25N-NBF (7) HCl**

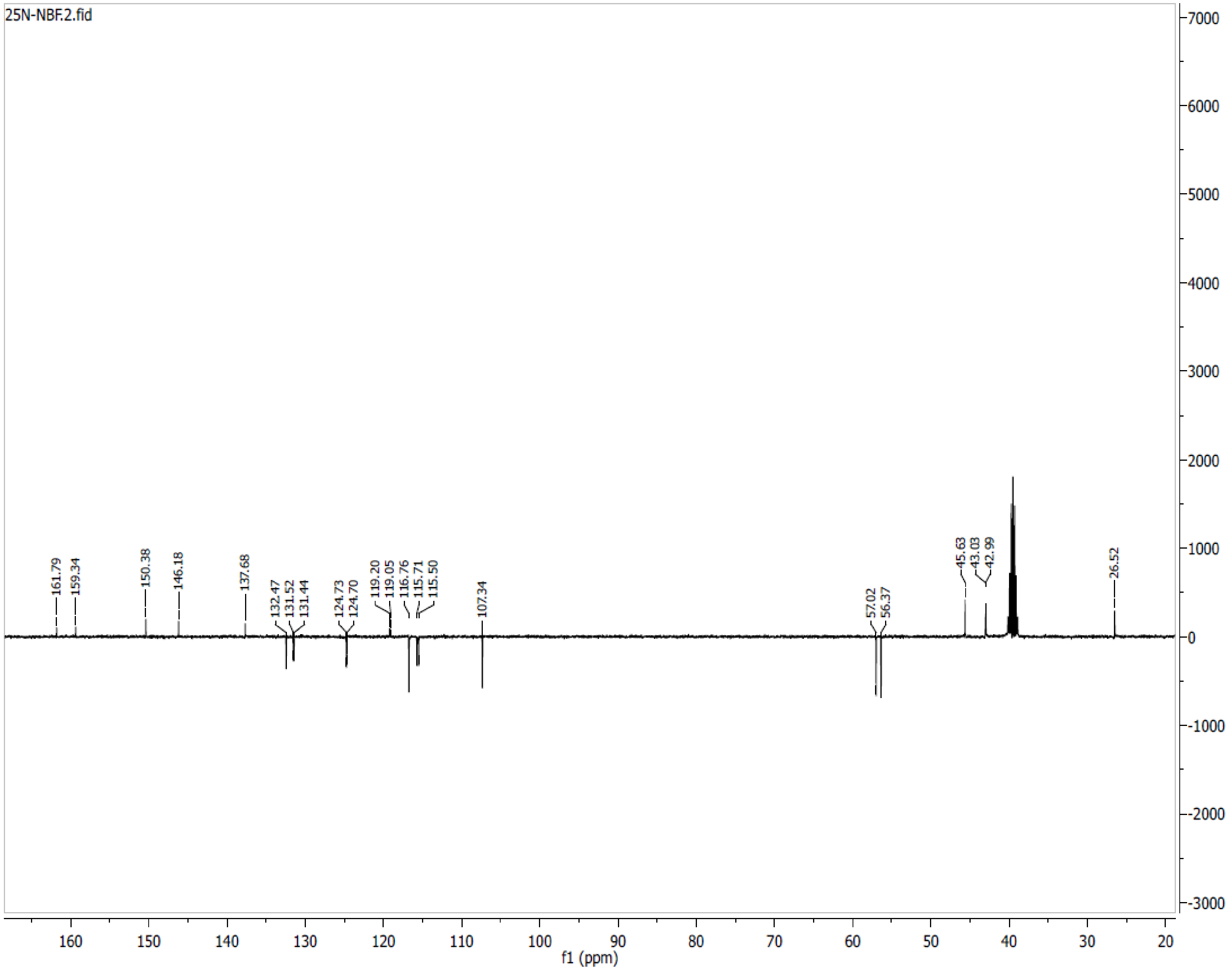

**Spectra 41. $^{13}$C NMR of 25N-NBCl (8) HCl**

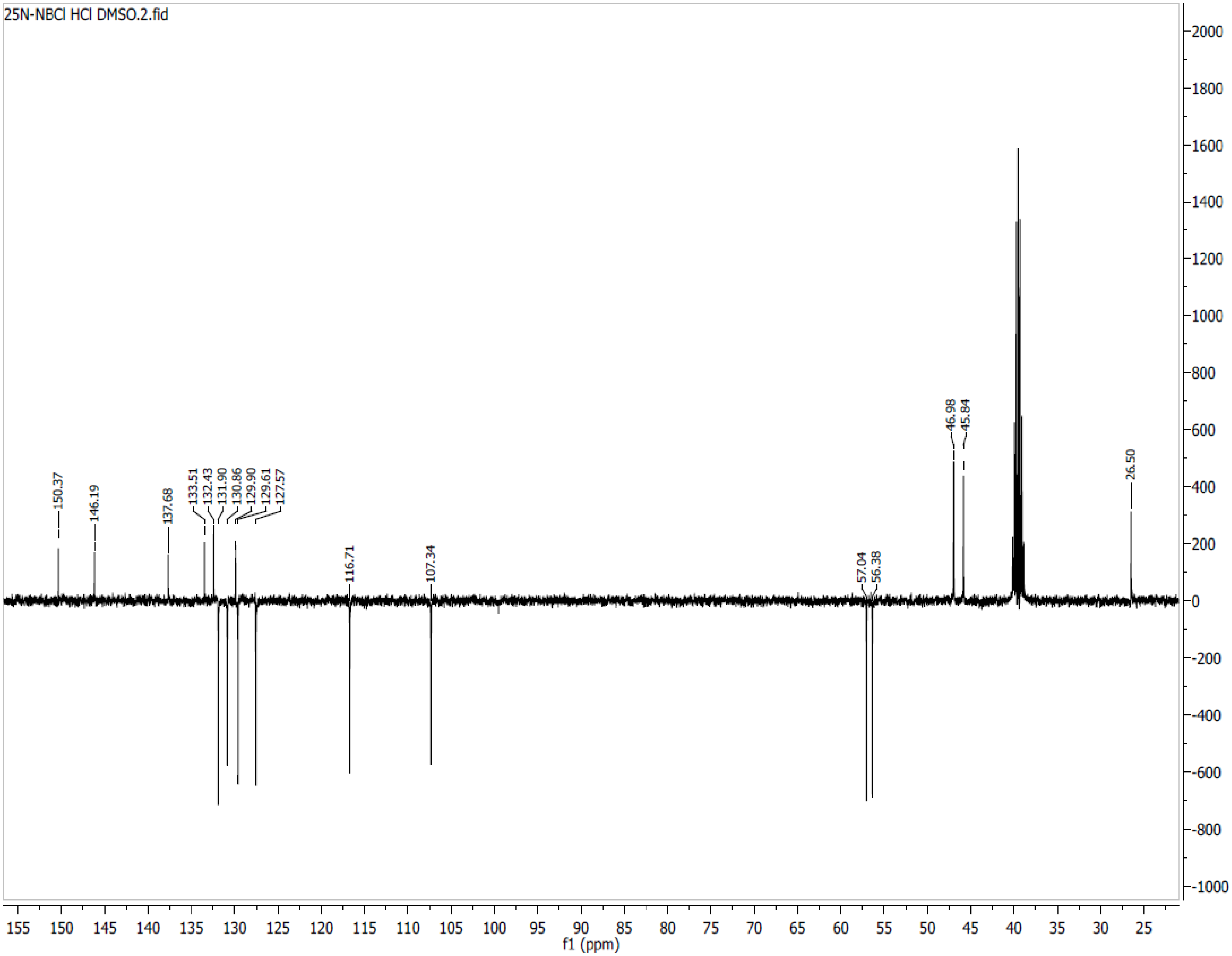

**Spectra 42. $^{13}C$ NMR of 25N-NBBr (9) HCl**

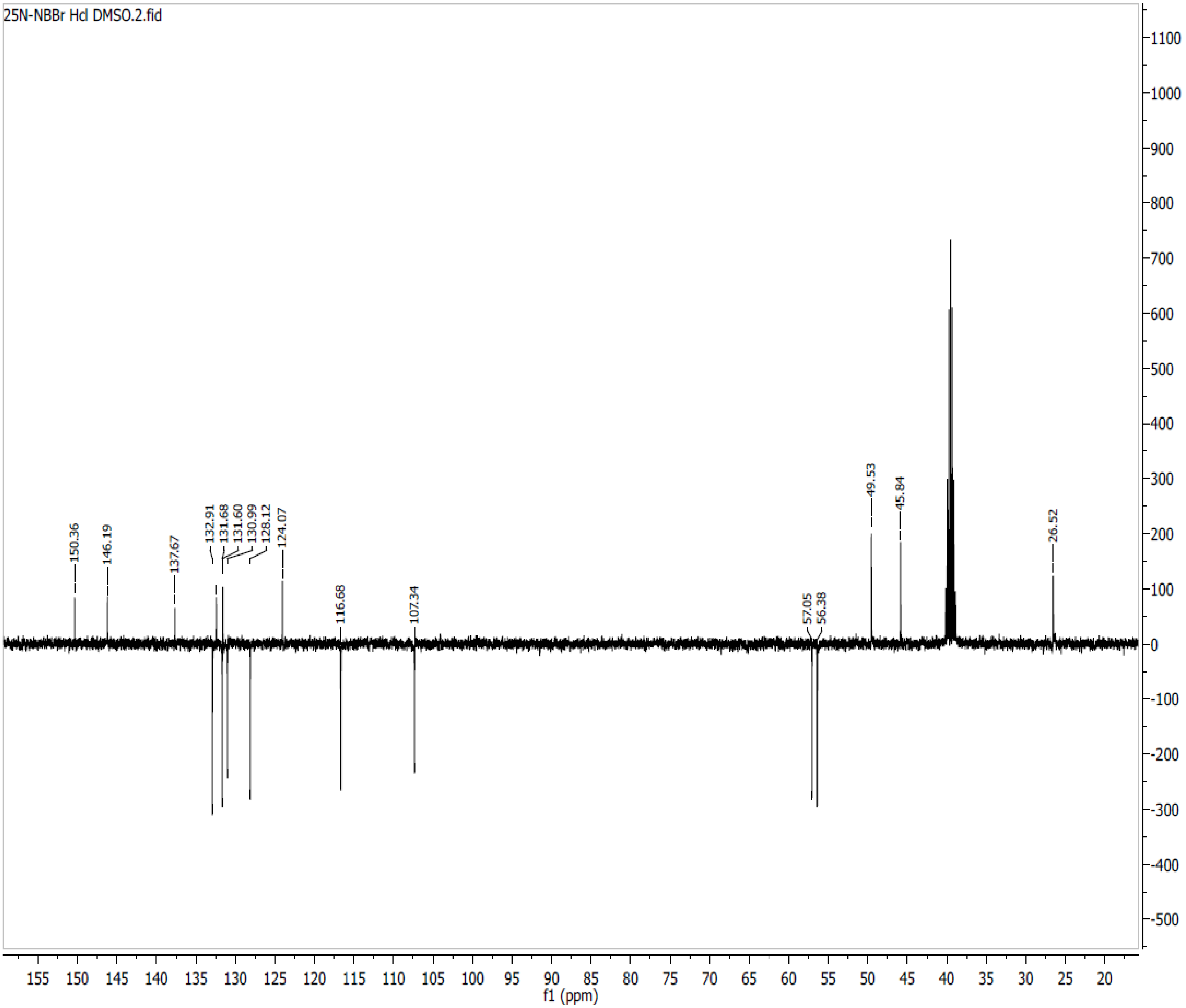

**Spectra 43. $^{13}$C NMR of 25N-NBI (10) HCl**

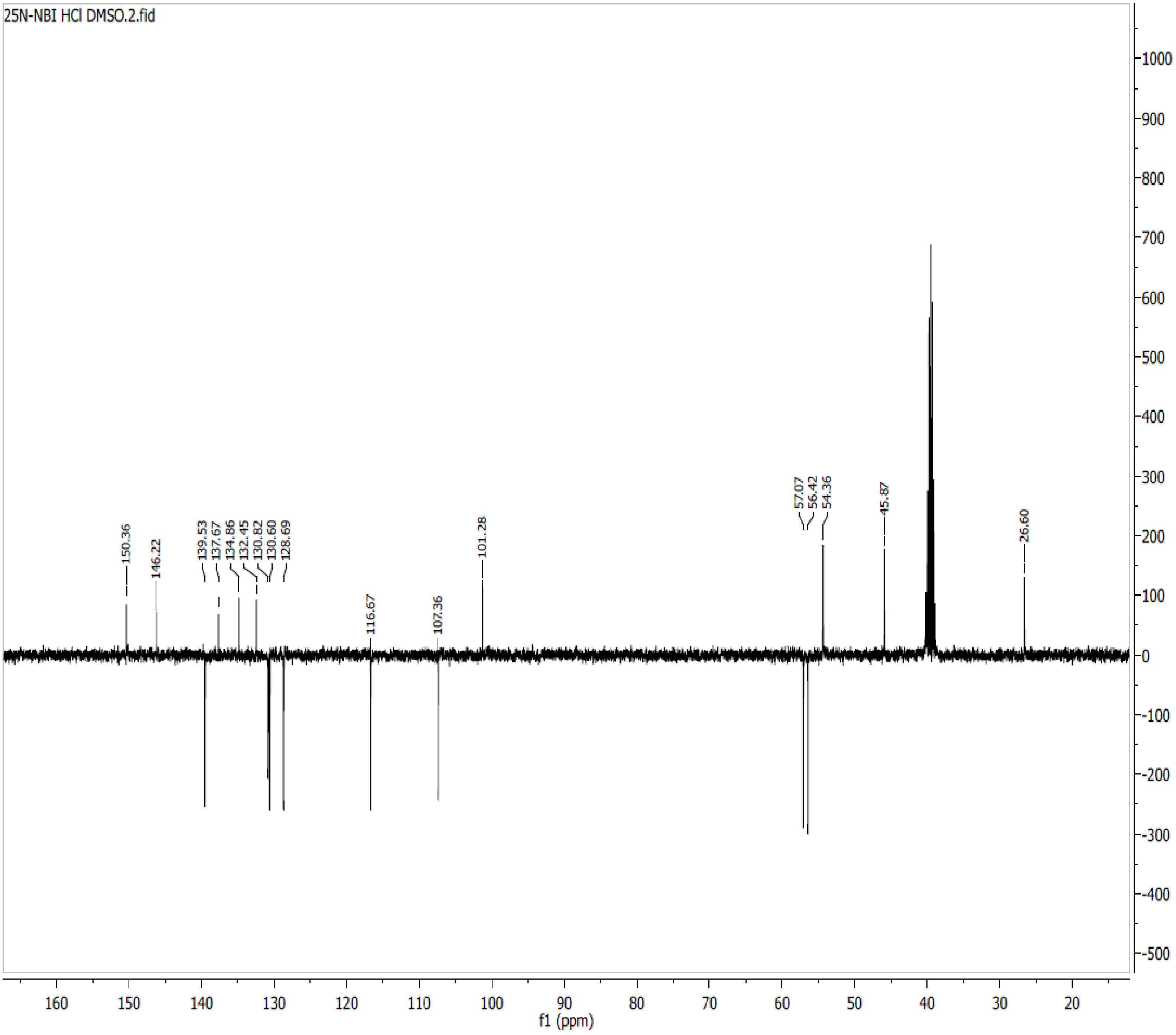

**Spectra 44. $^{13}$C NMR of 25N-NBOCF$_2$H (11) HCl**

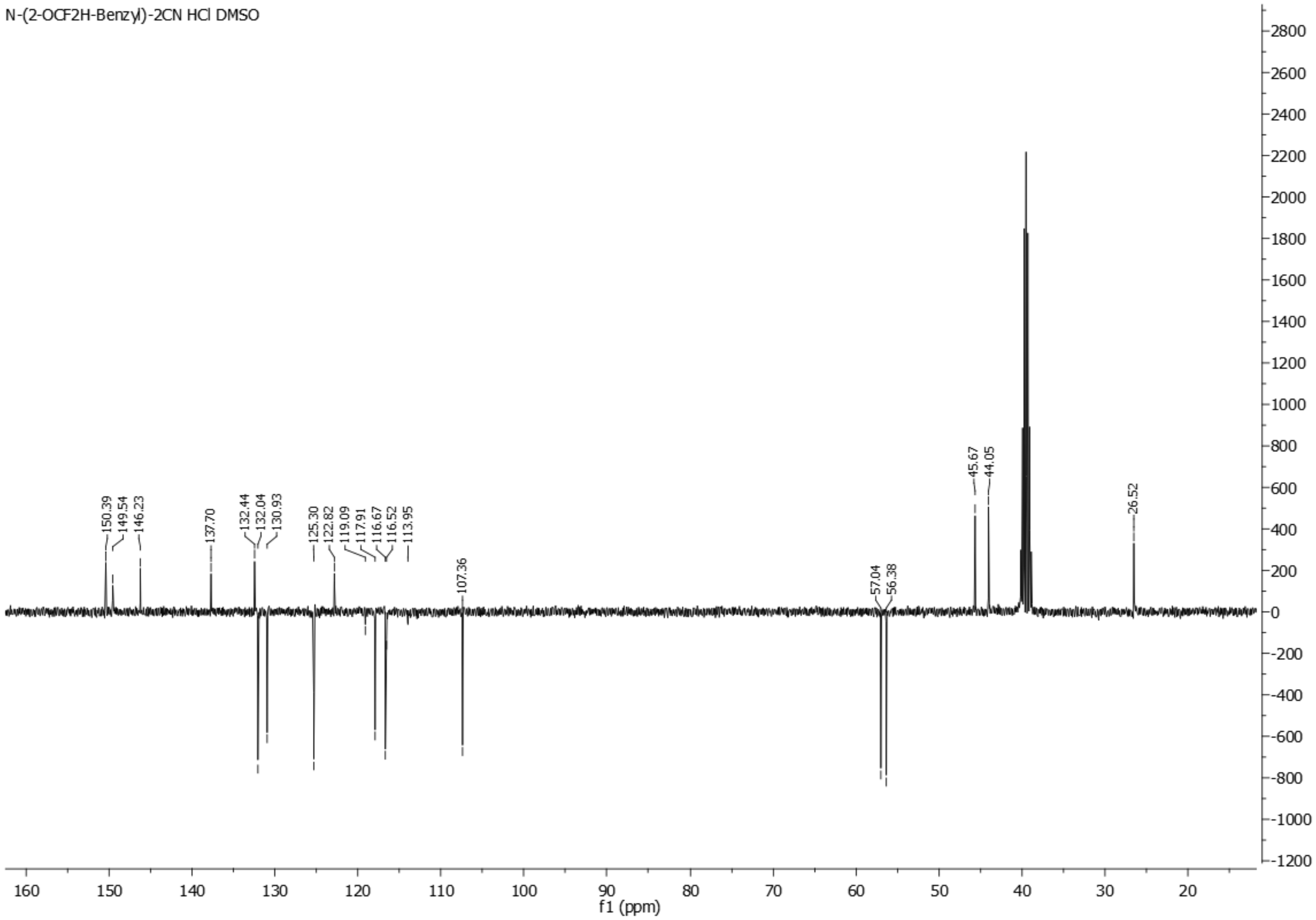

**Spectra 45. $^{13}C$ NMR of 25N-NBOCF$_3$ (12) HCl**

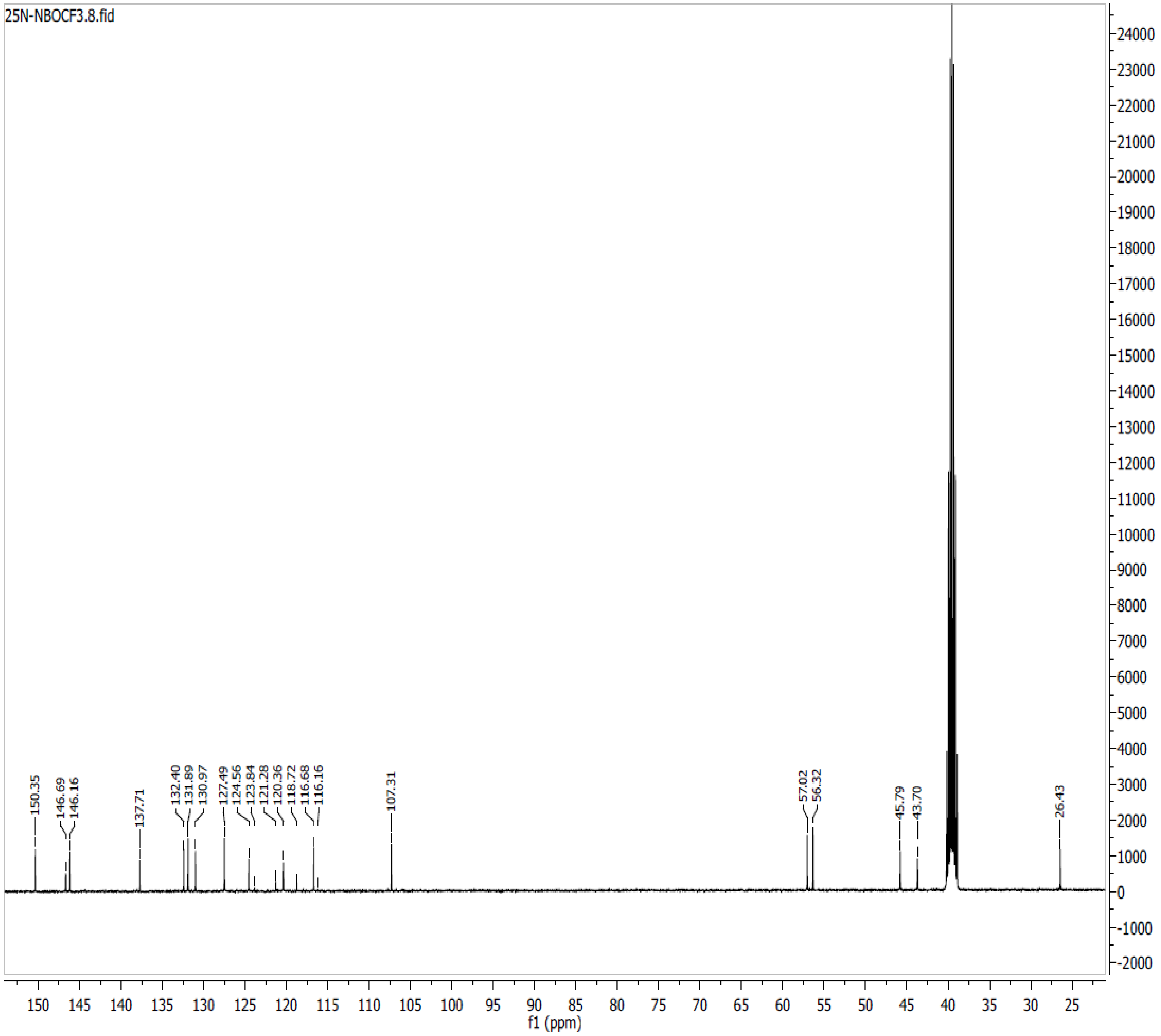

**Spectra 46. $^{13}C$ NMR of 25N-NBMDF$_2$ (13) HCl**

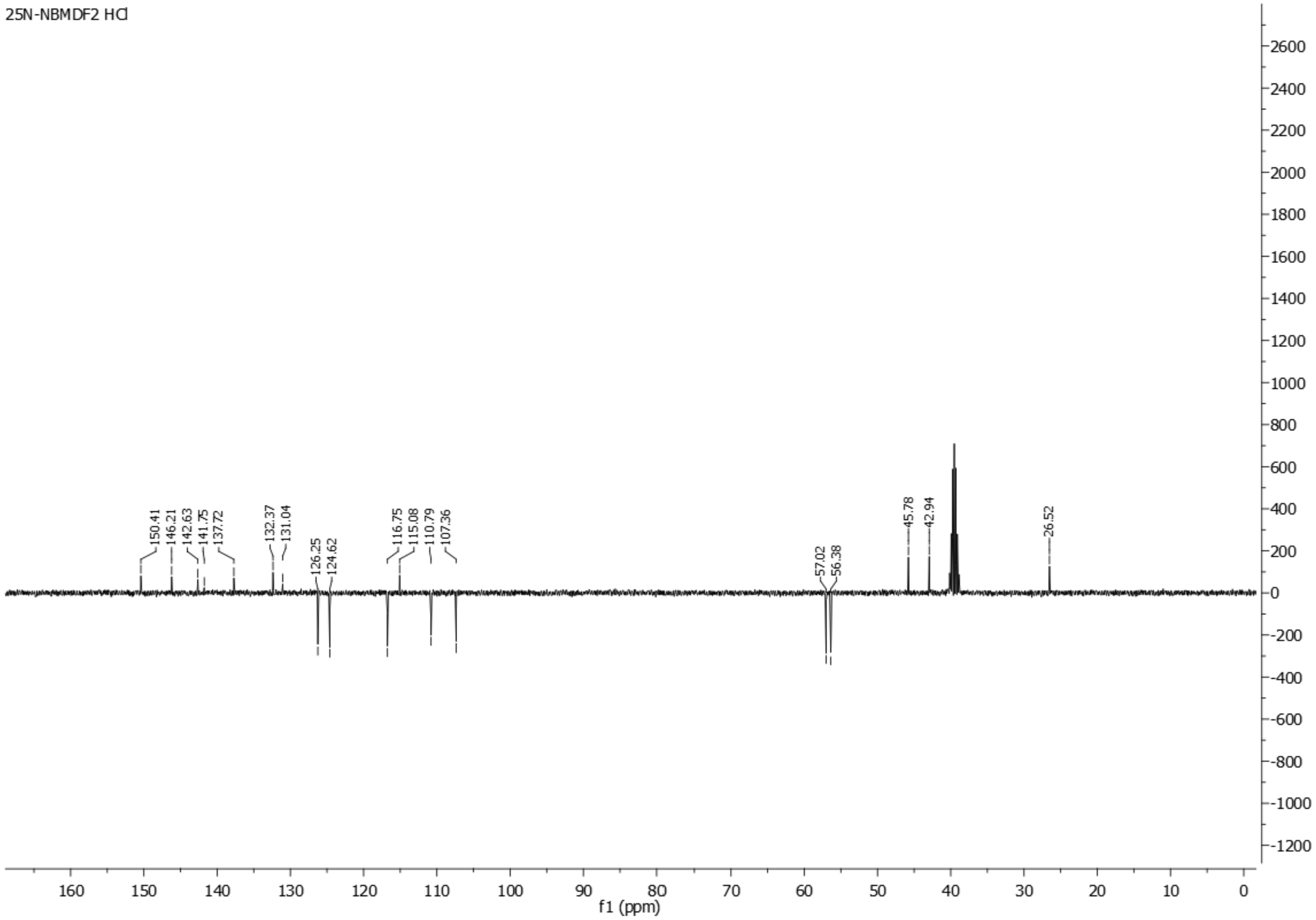

**Spectra 47. $^{13}C$ NMR of 25N-NBCF$_3$ (14) HCl**

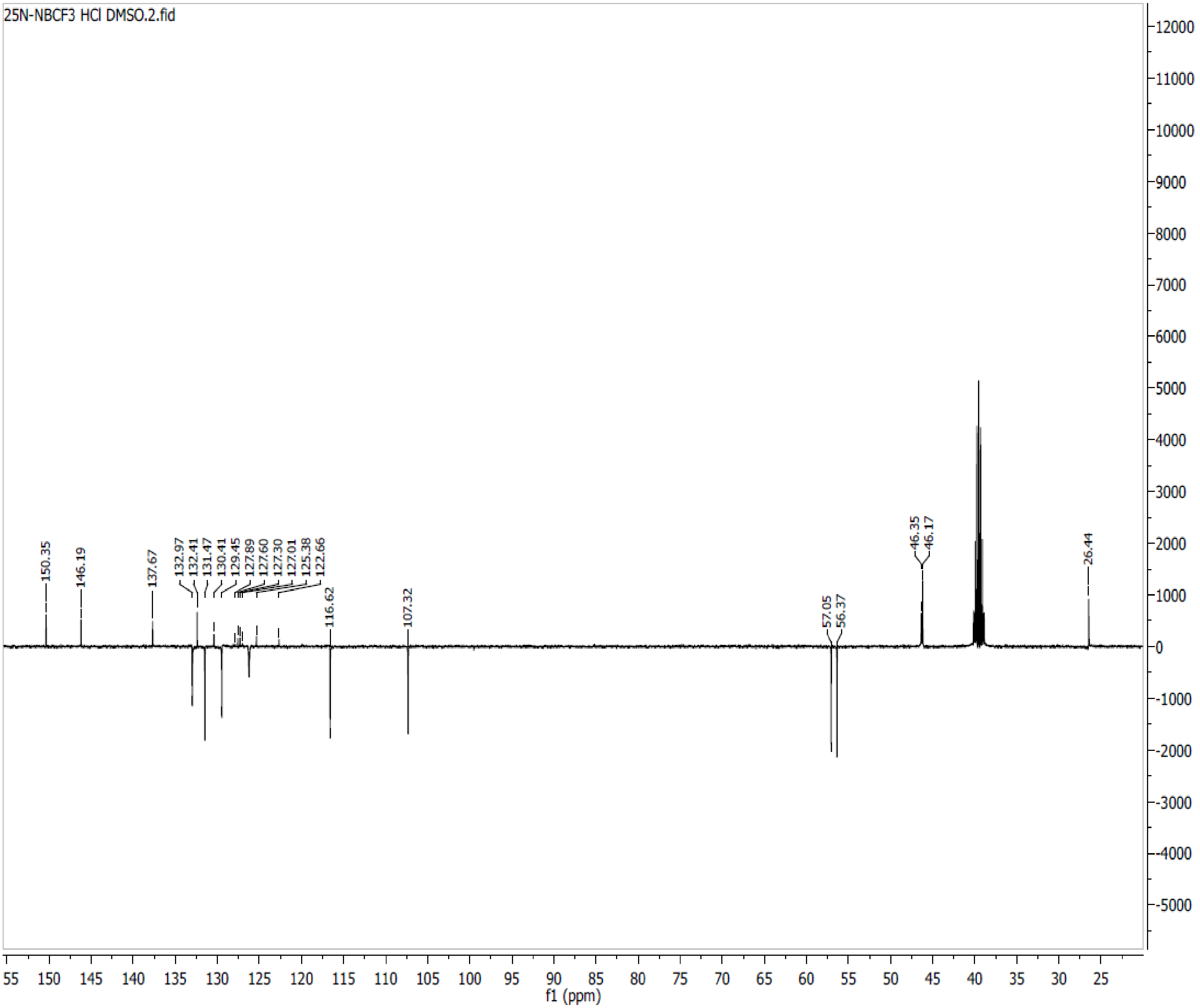

**Spectra 48. $^{13}C$ NMR of 25N-NBNO$_2$ (15) HCl**

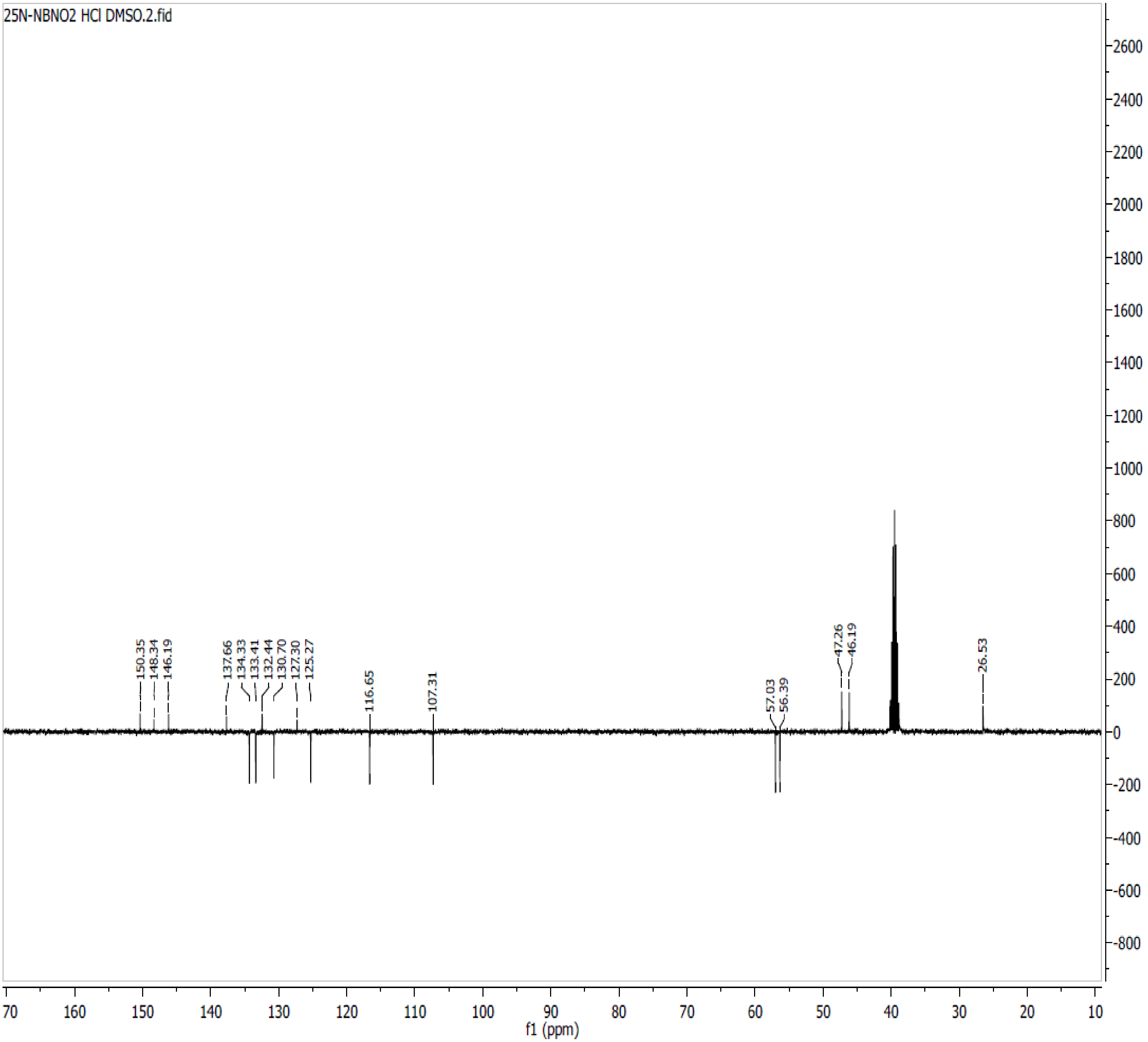

**Spectra 49. $^{13}$C NMR of 25N-N1-Nap (16) HCl**

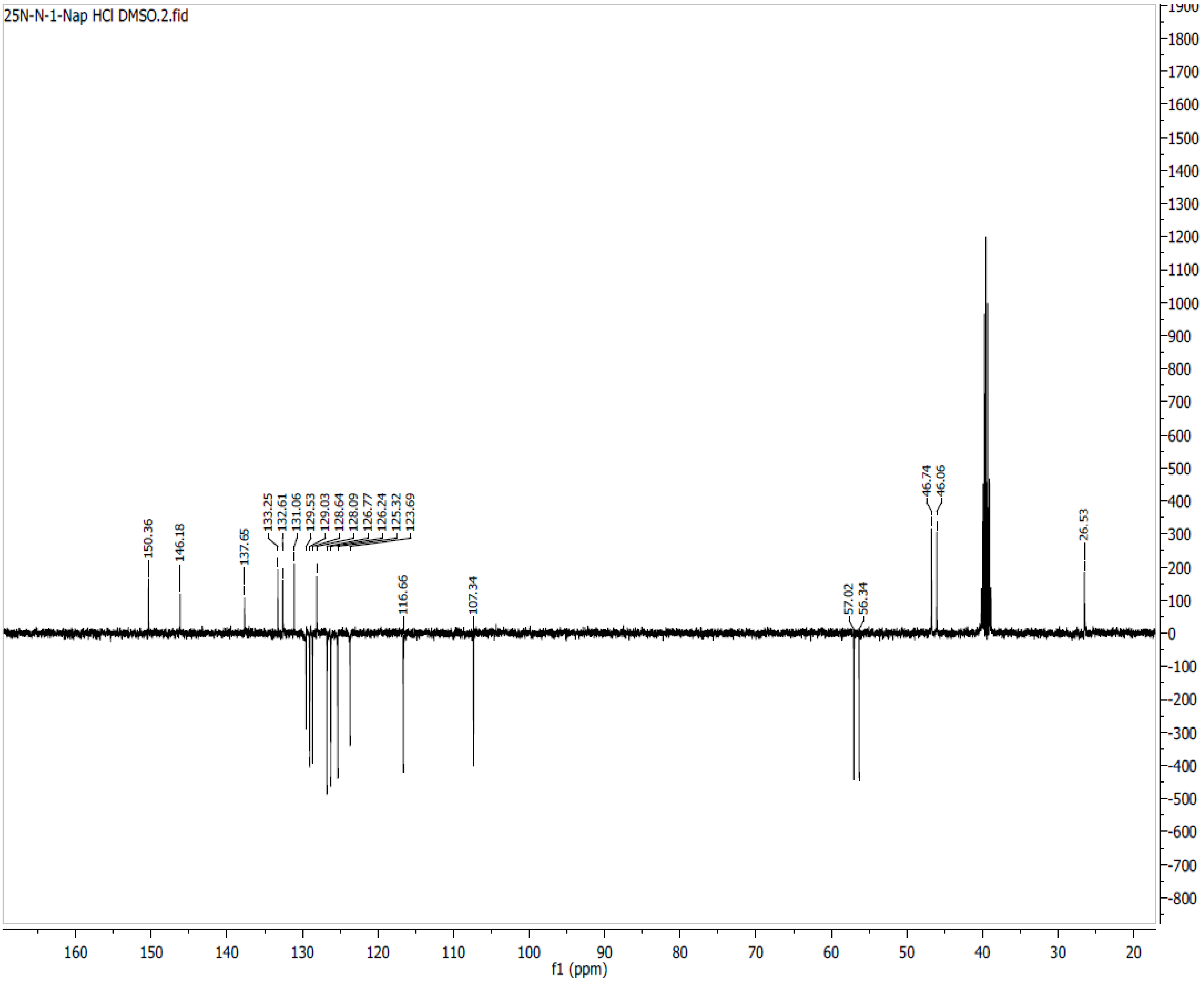

**Spectra 50. $^{13}$C NMR of 25N-NBPh (17) HCl**

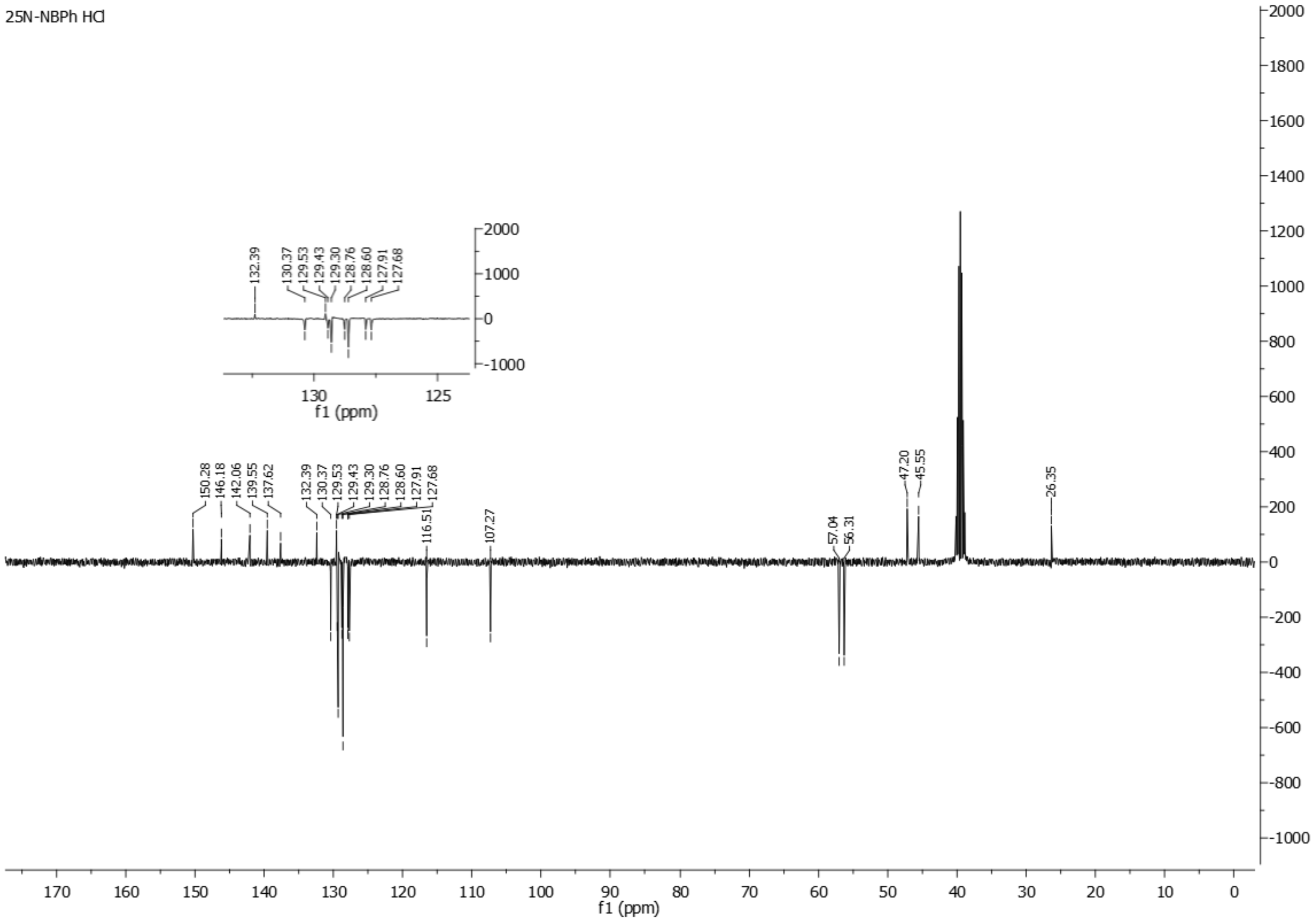

**Spectra 51. $^{13}$C NMR of 25N-NB-2-OH-3-Me (18) HCl**

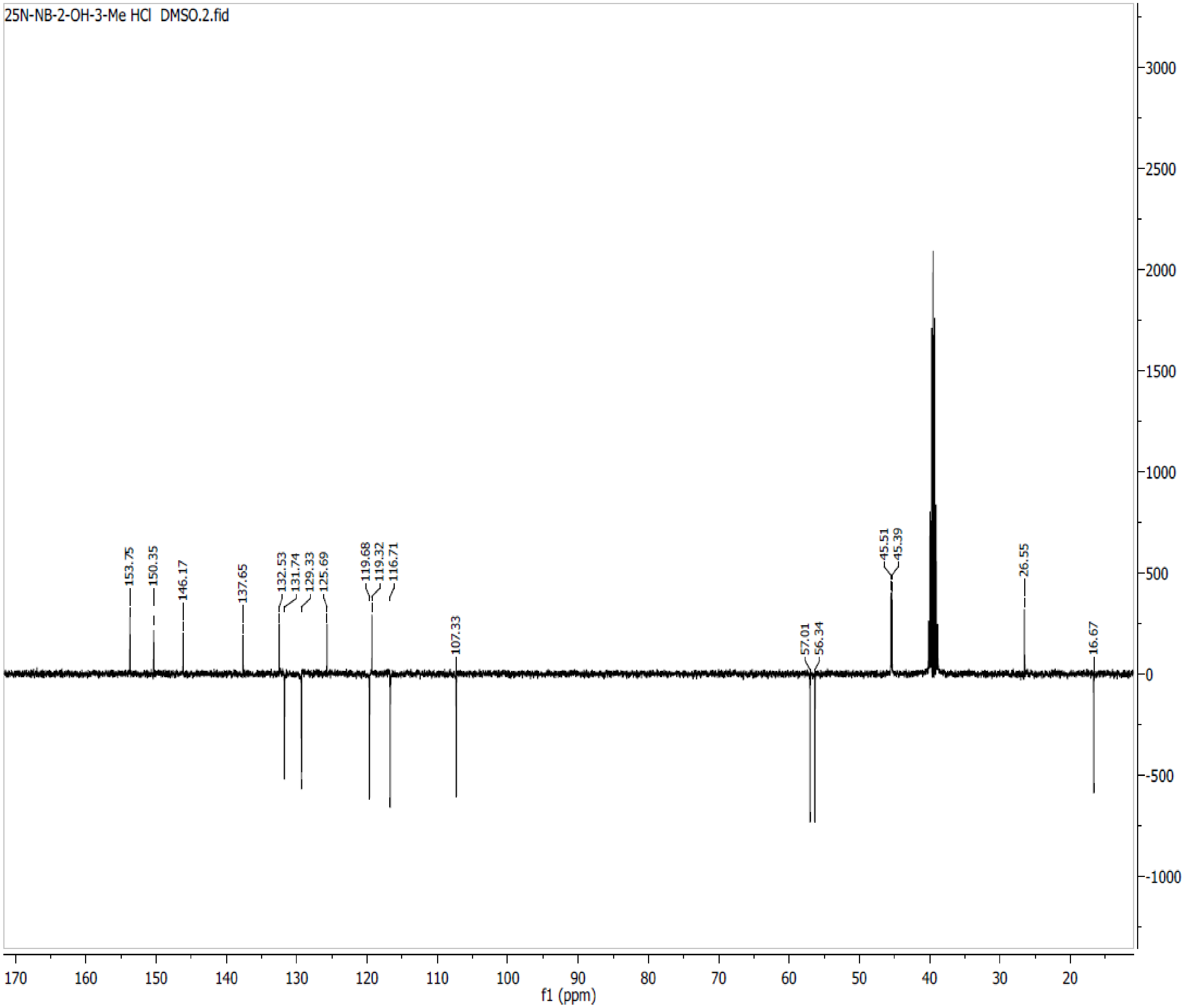

**Spectra 52. $^{13}C$ NMR of 25N-NB-2-MeO-3-F (19) HCl**

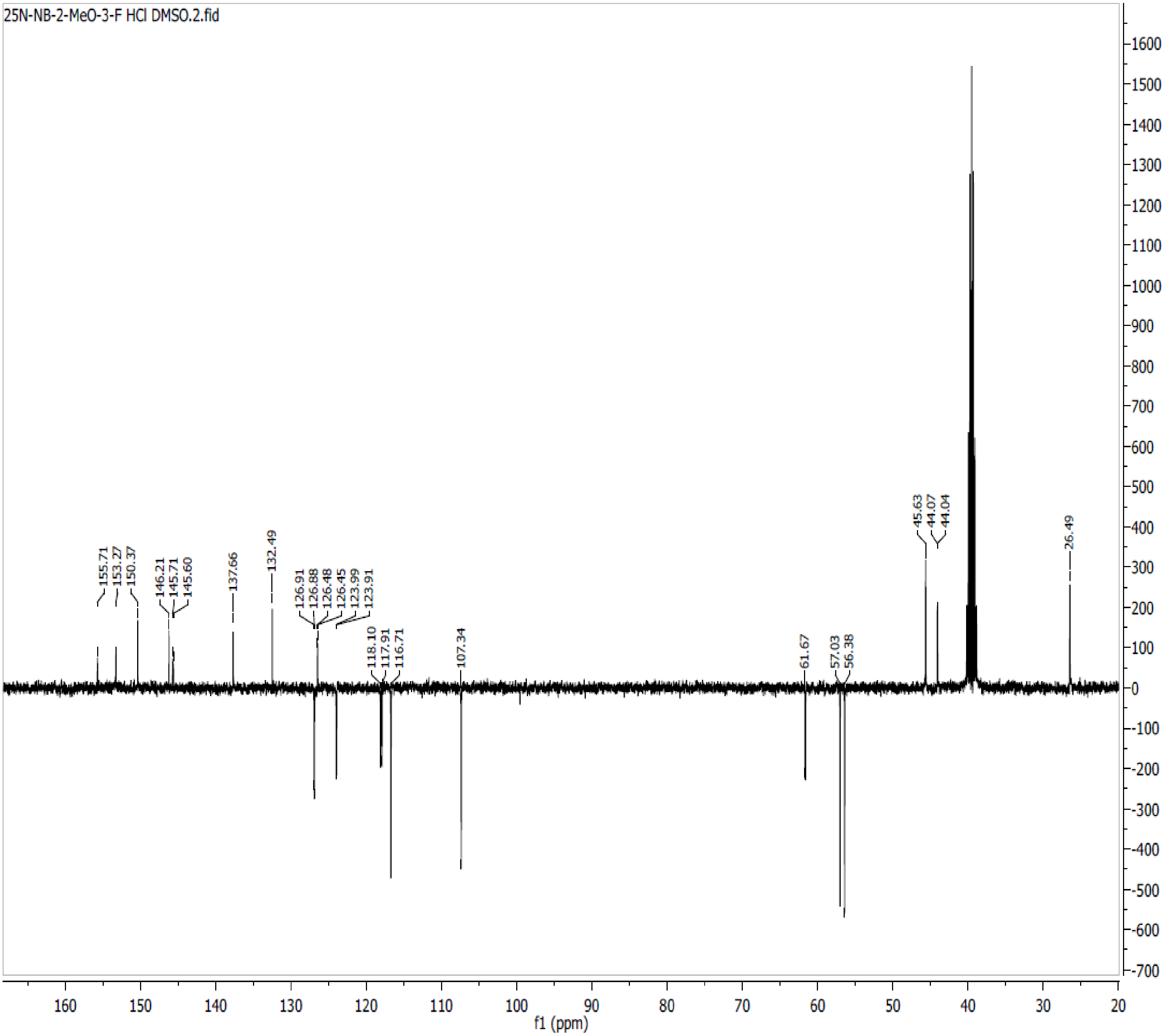

**Spectra 53. $^{13}$C NMR of 25N-NB-2,5-DiMeO (20) HCl**

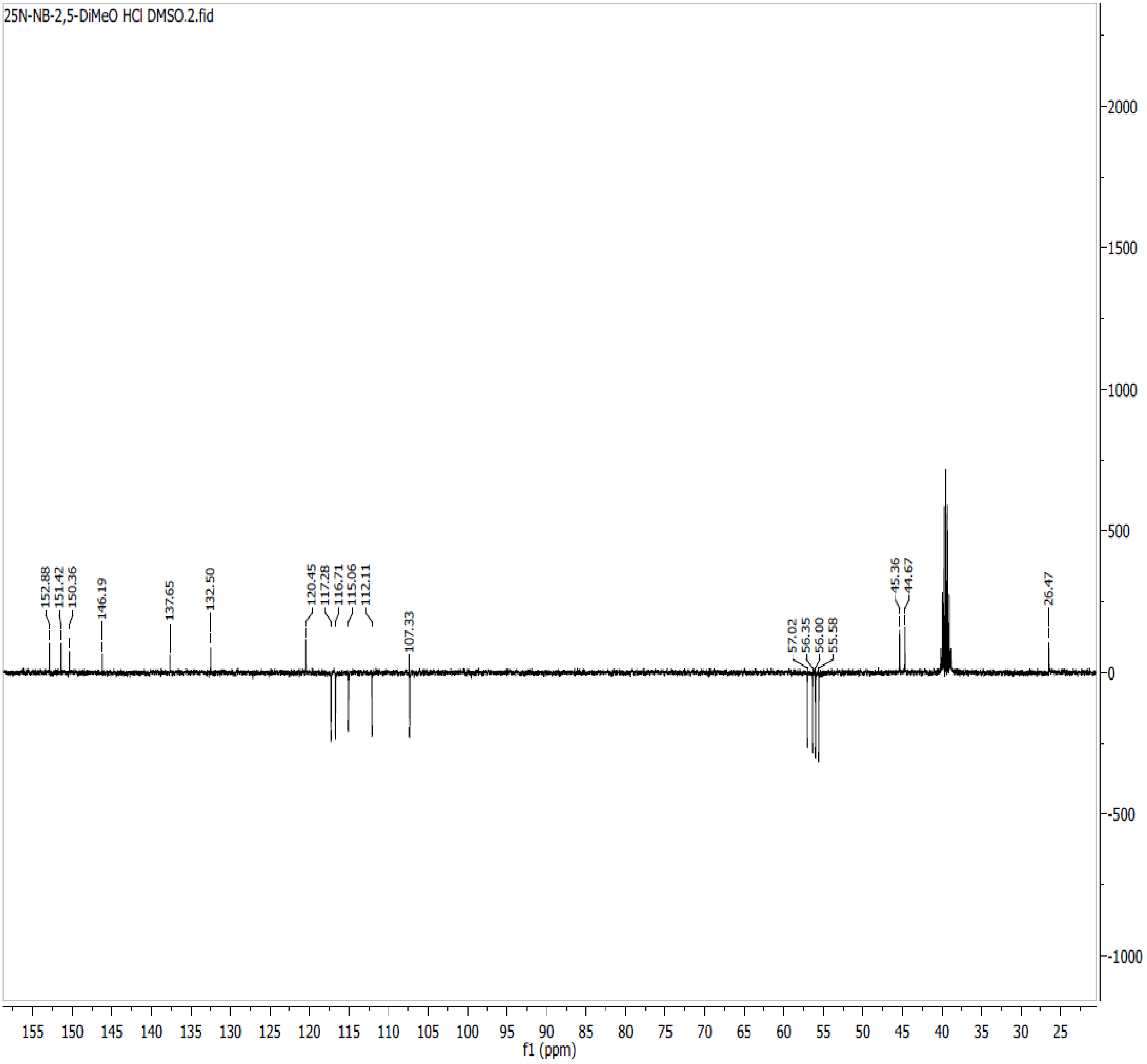

**Spectra 54. $^{13}C$ NMR of 25N-NB-3-OH (21) HCl**

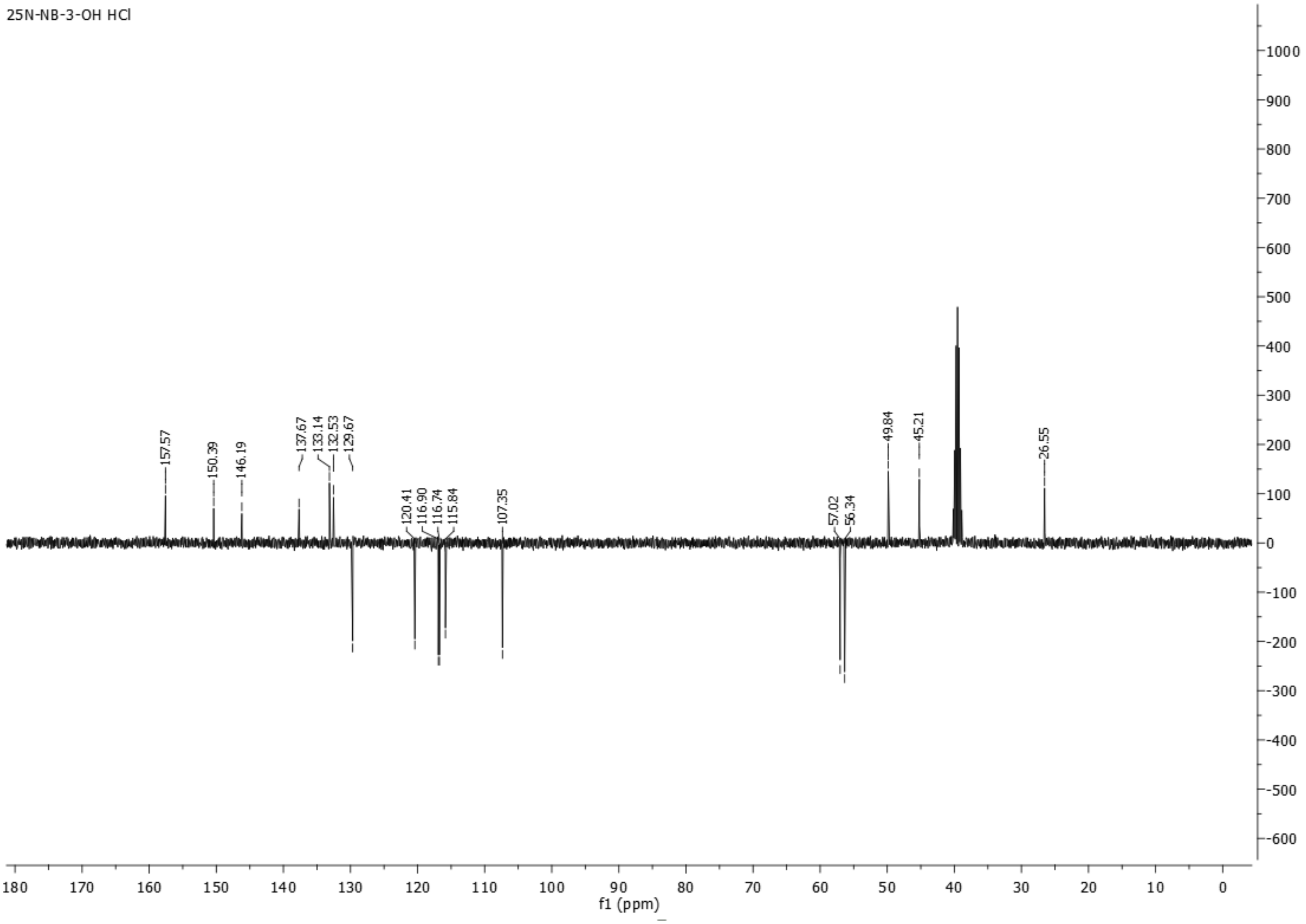

**Spectra 55. $^{13}C$ NMR of 25N-NB-3-Me (22) HCl**

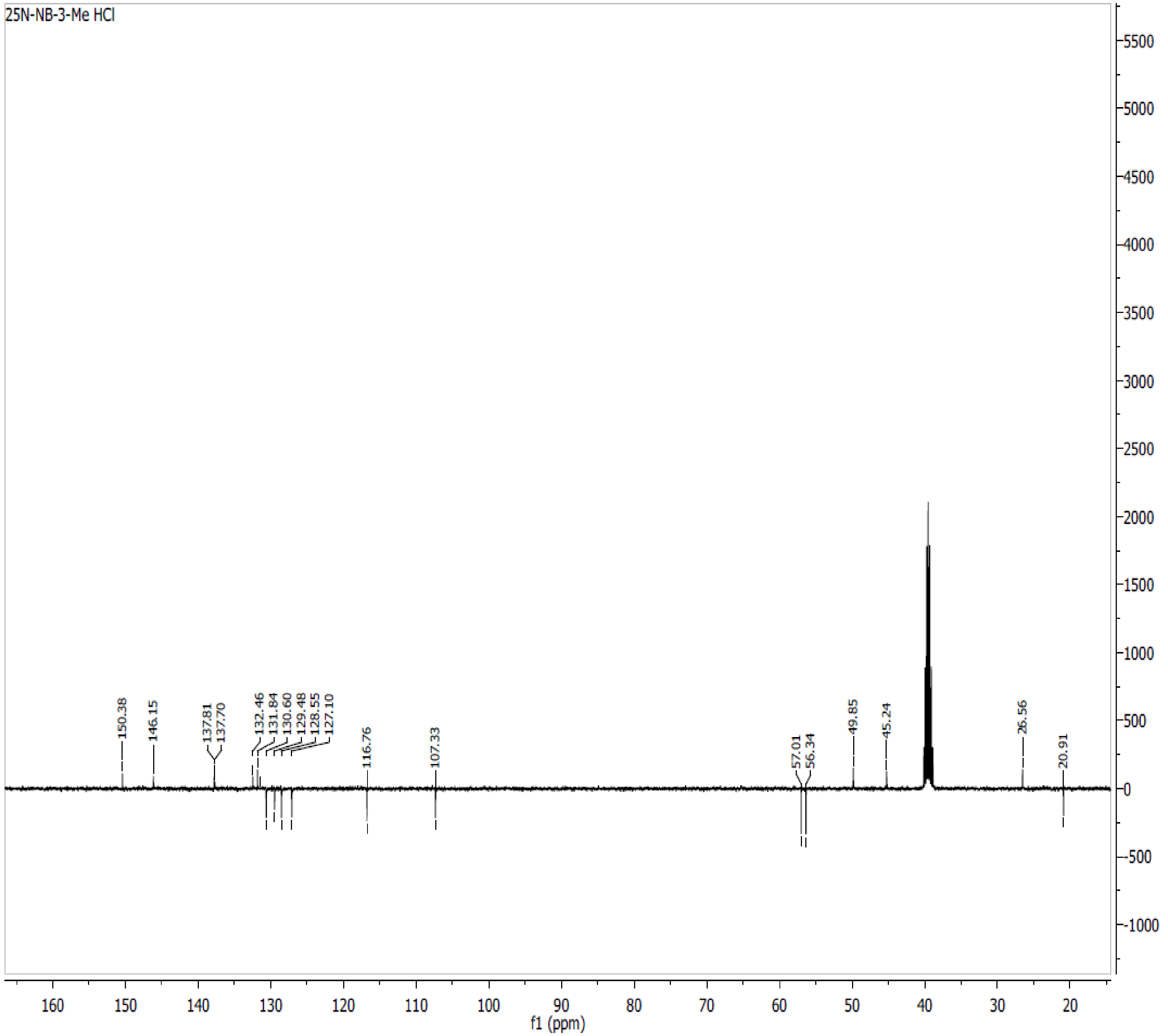

**Spectra 56. $^{13}$C NMR of 25N-NB-4-Me (23) HCl**

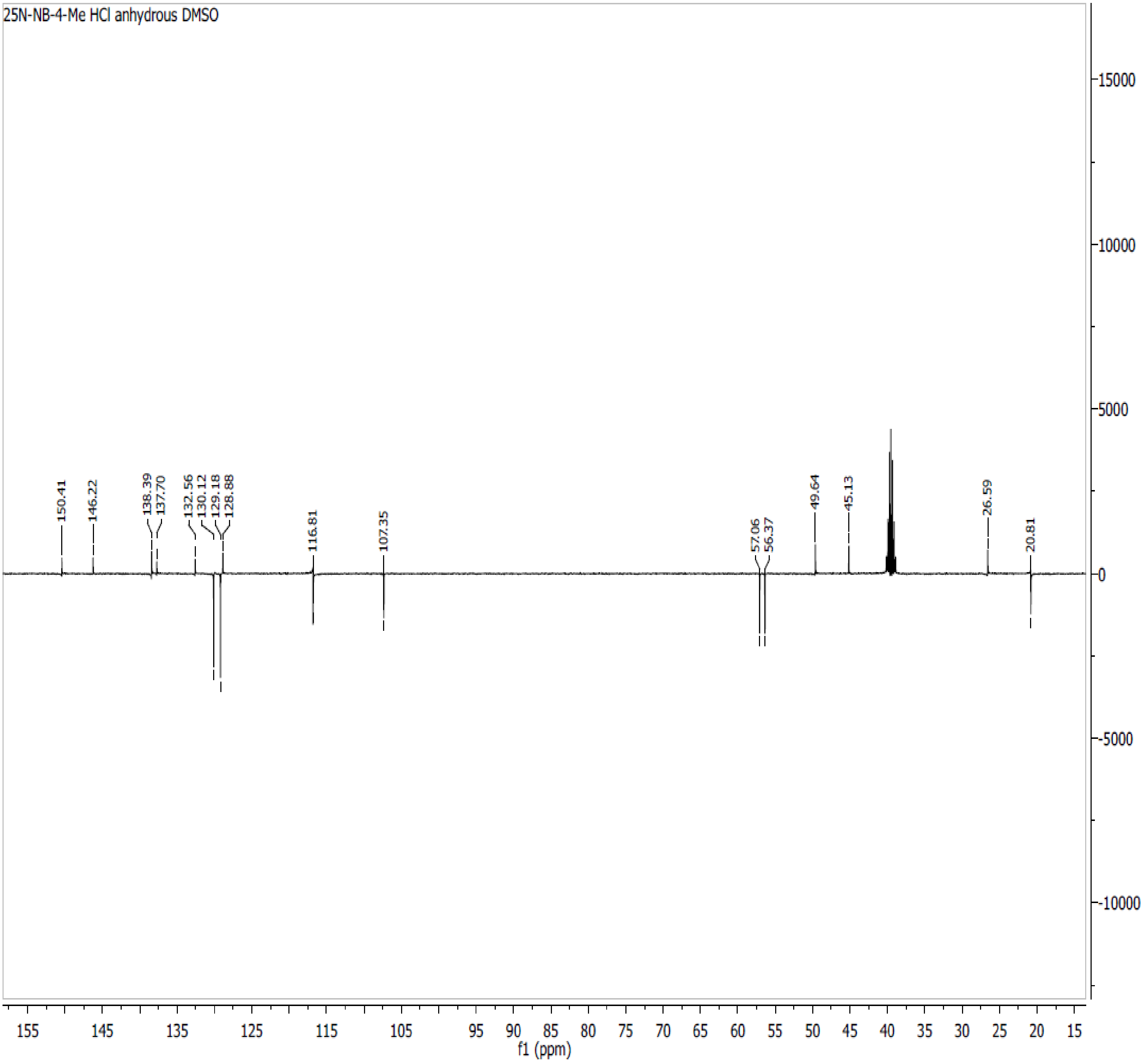

**Spectra 57. $^{13}$C NMR of 25N-NB-3-F (24) HCl**

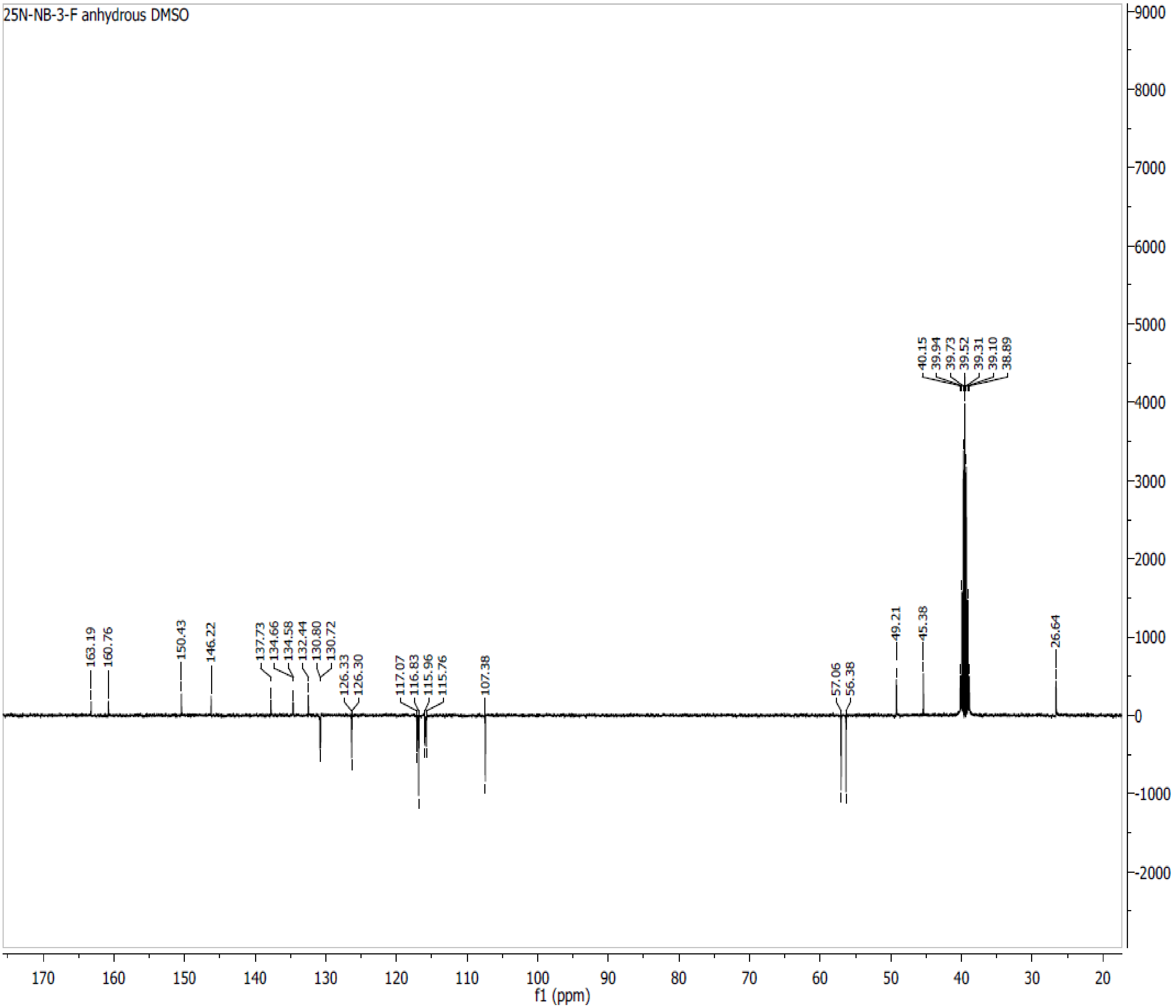

**Spectra 58. $^{13}$C NMR of 25N-NB-4-F (25) HCl**

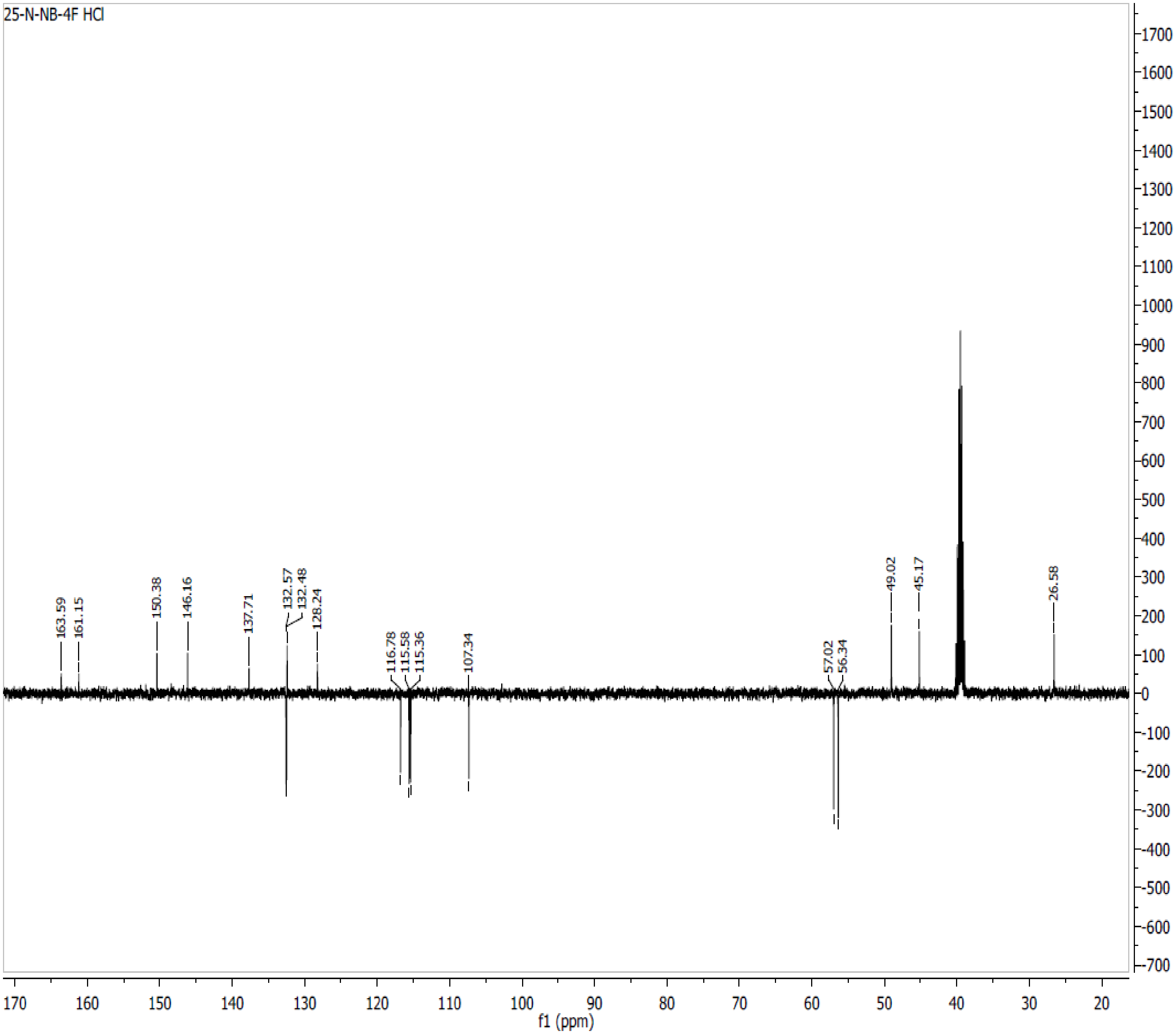

**Spectra 59. $^{13}C$ NMR of 25D-N1-Nap (26) HCl**

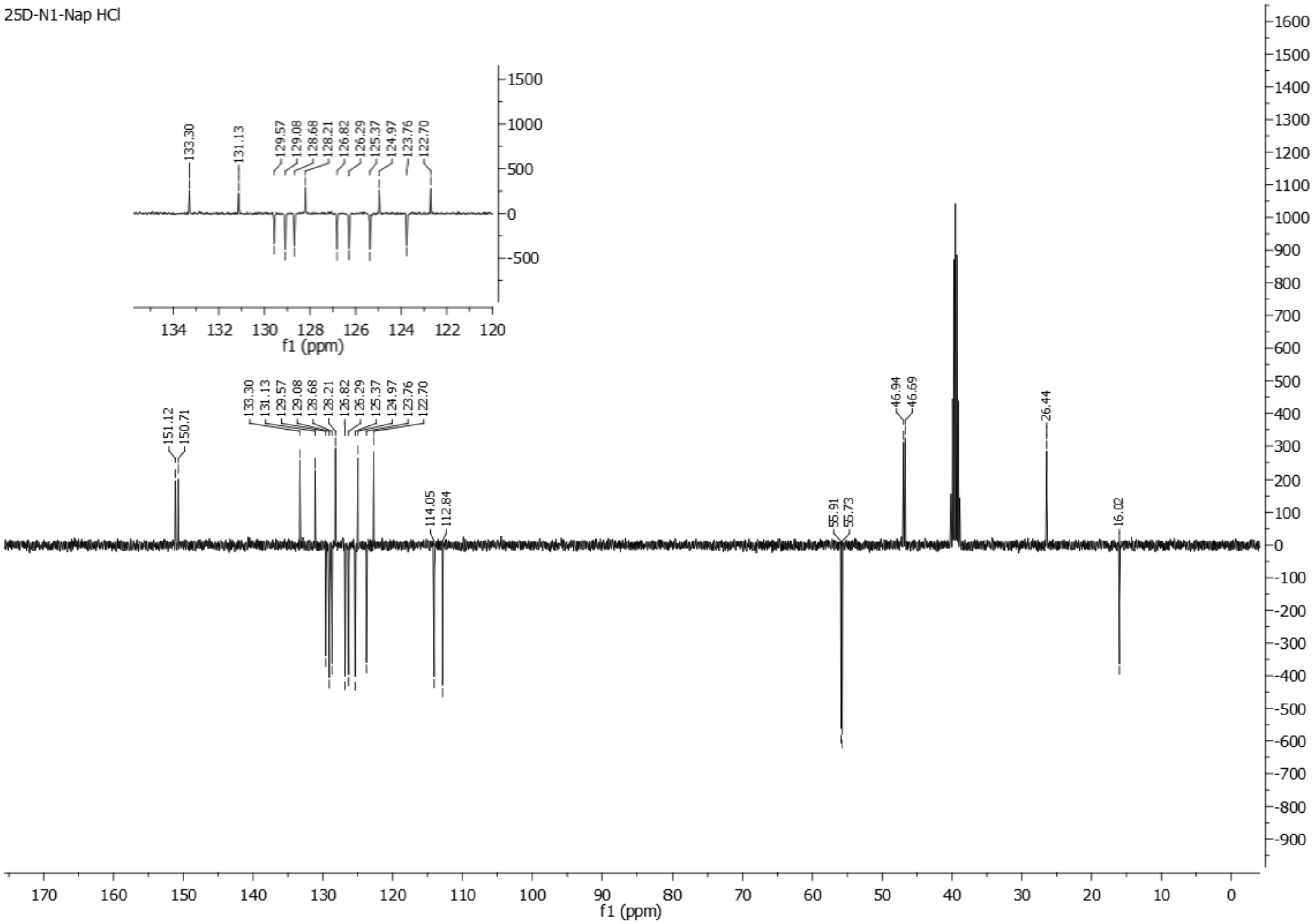

**Spectra 60. $^{13}$C NMR of 25D-NBPh (27) HCl**

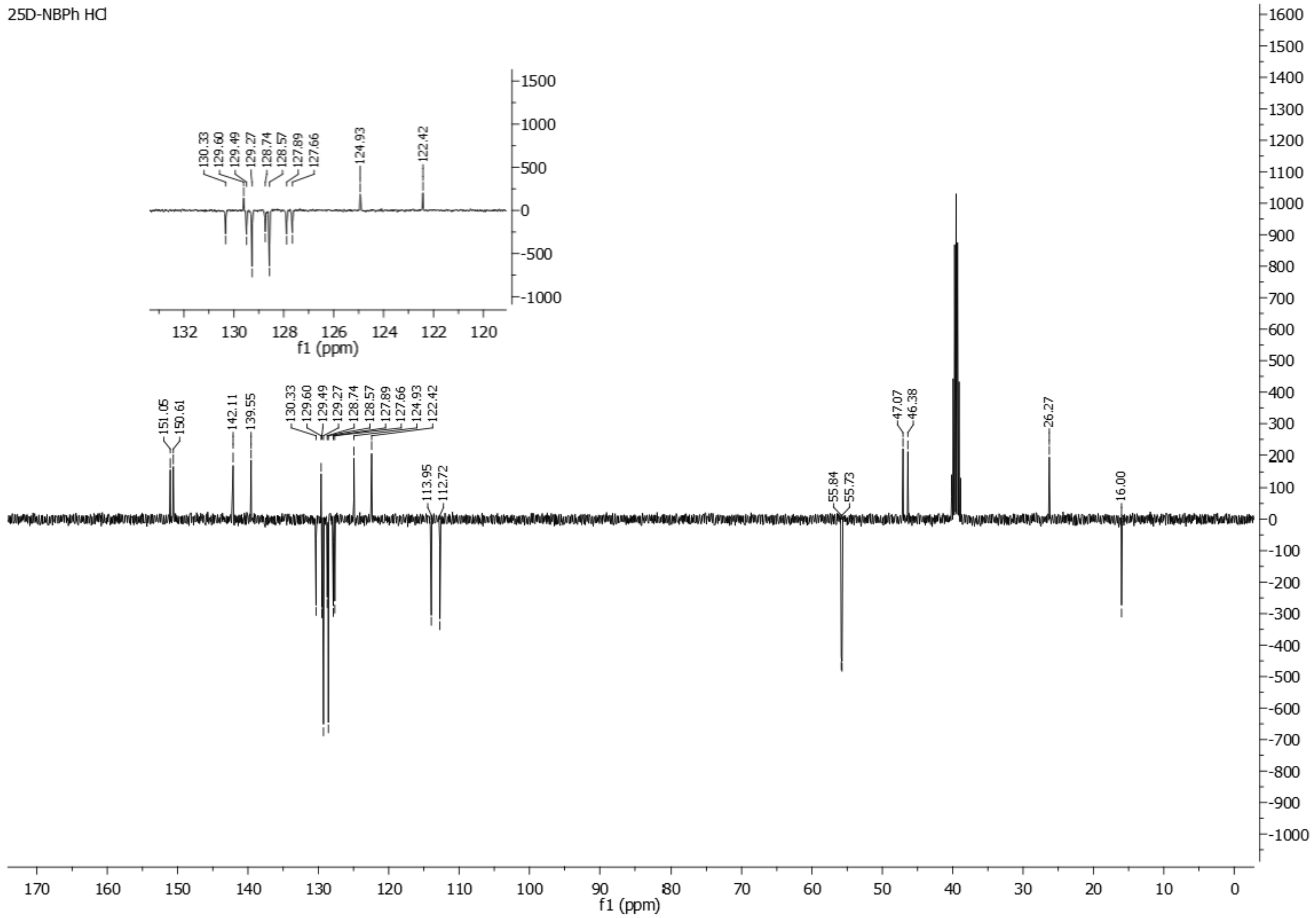

**Spectra 61. $^{13}$C NMR of 25O-N1-Nap (28) HCl**

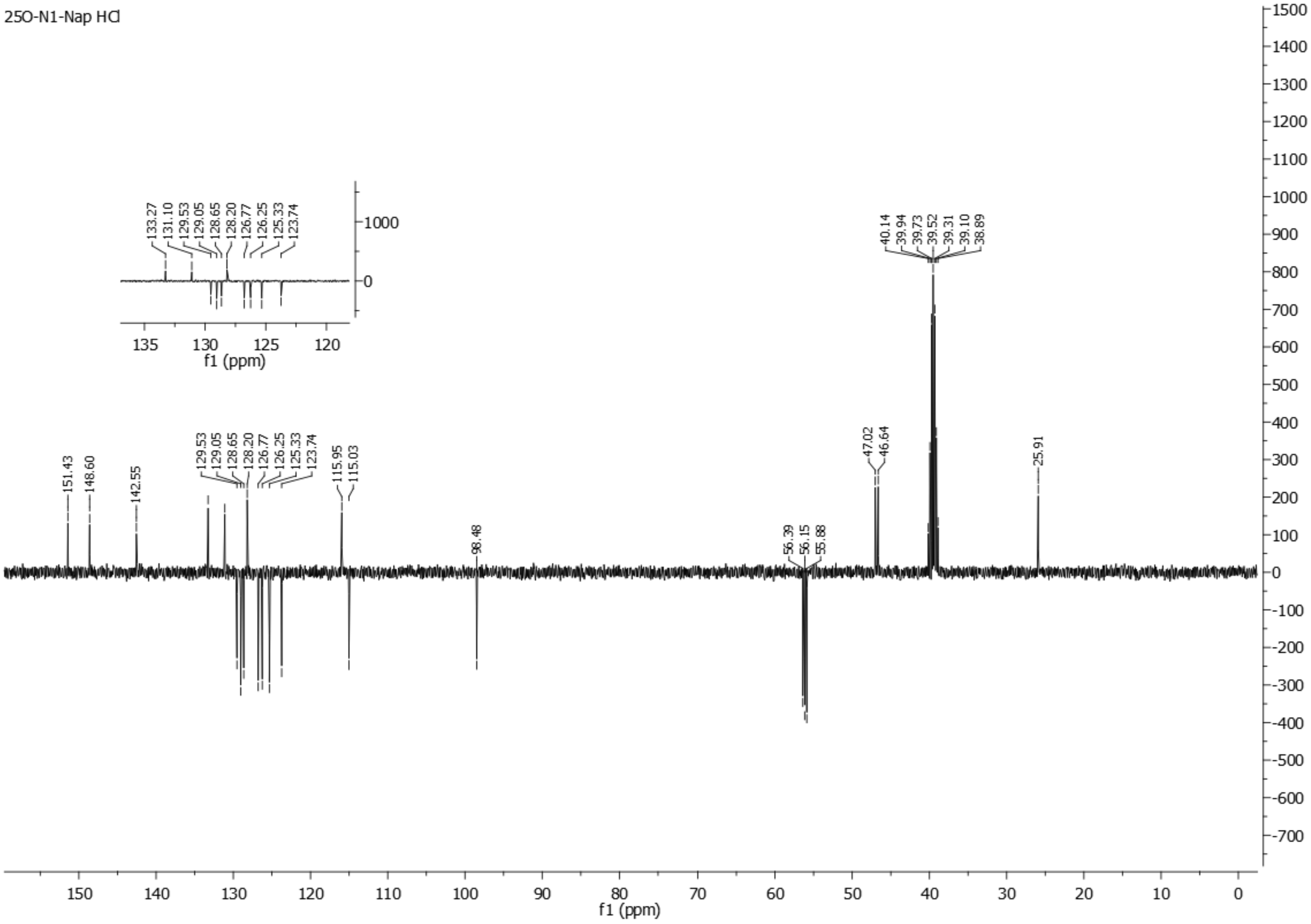

**Spectra 62.** $^{13}$C NMR spectrum of **2C2-N1-Nap (29) HCl**

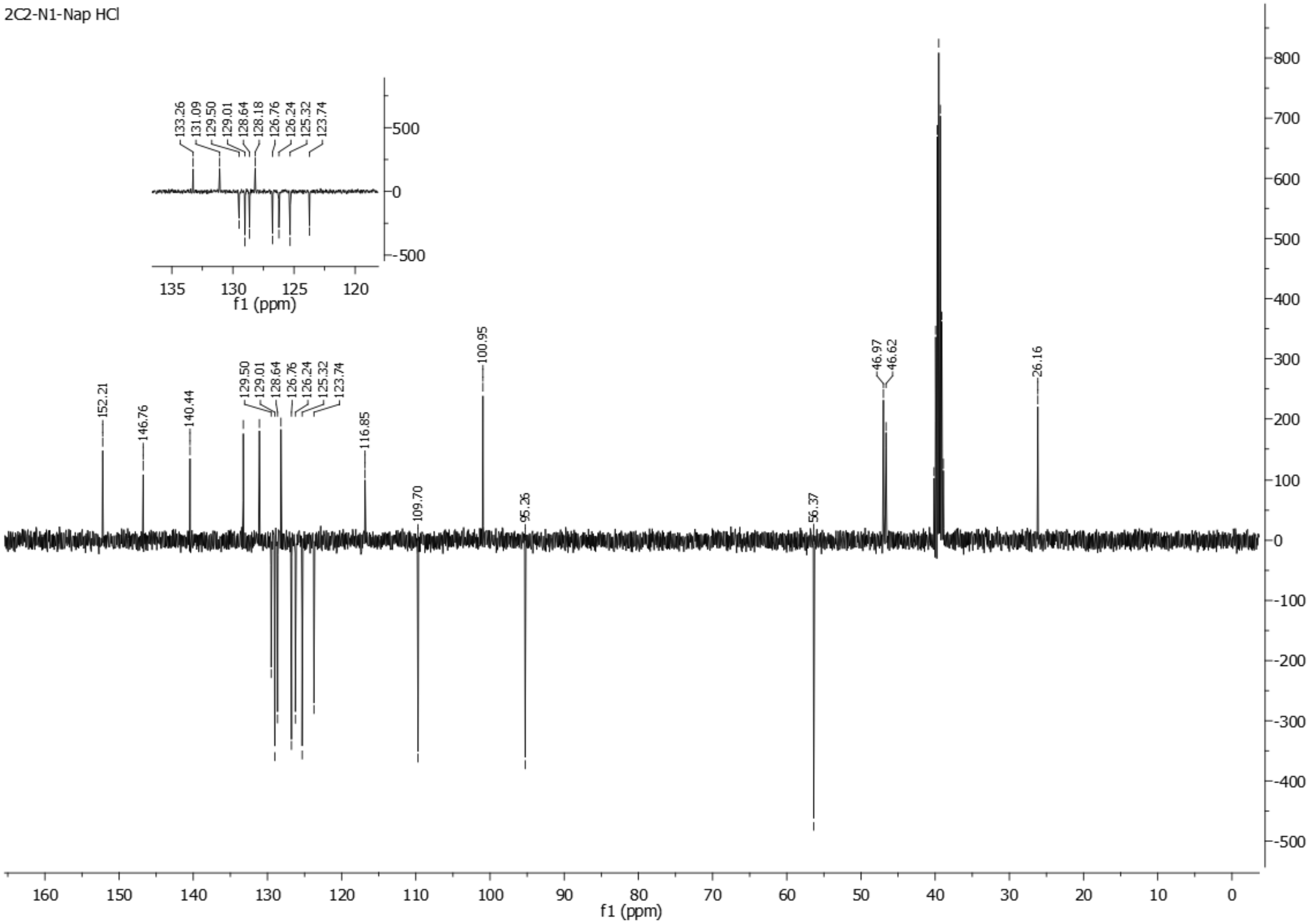

### Spectra 63. $^{13}$C NMR spectrum of Ψ-DOB-N1-Nap (30) HCl

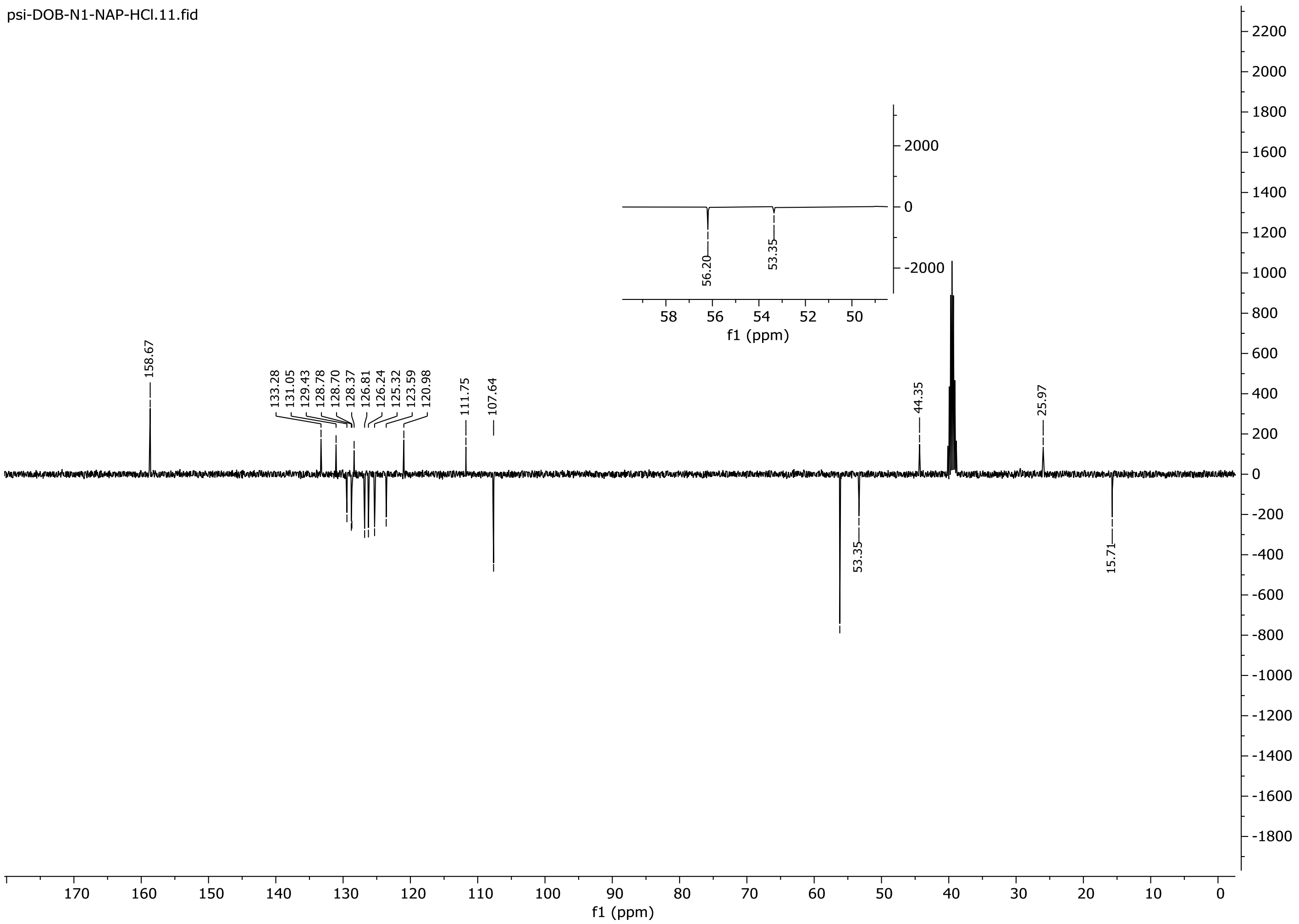

### Spectra 64. $^{13}C$ NMR of 2C2-NBOMe (31) HCl

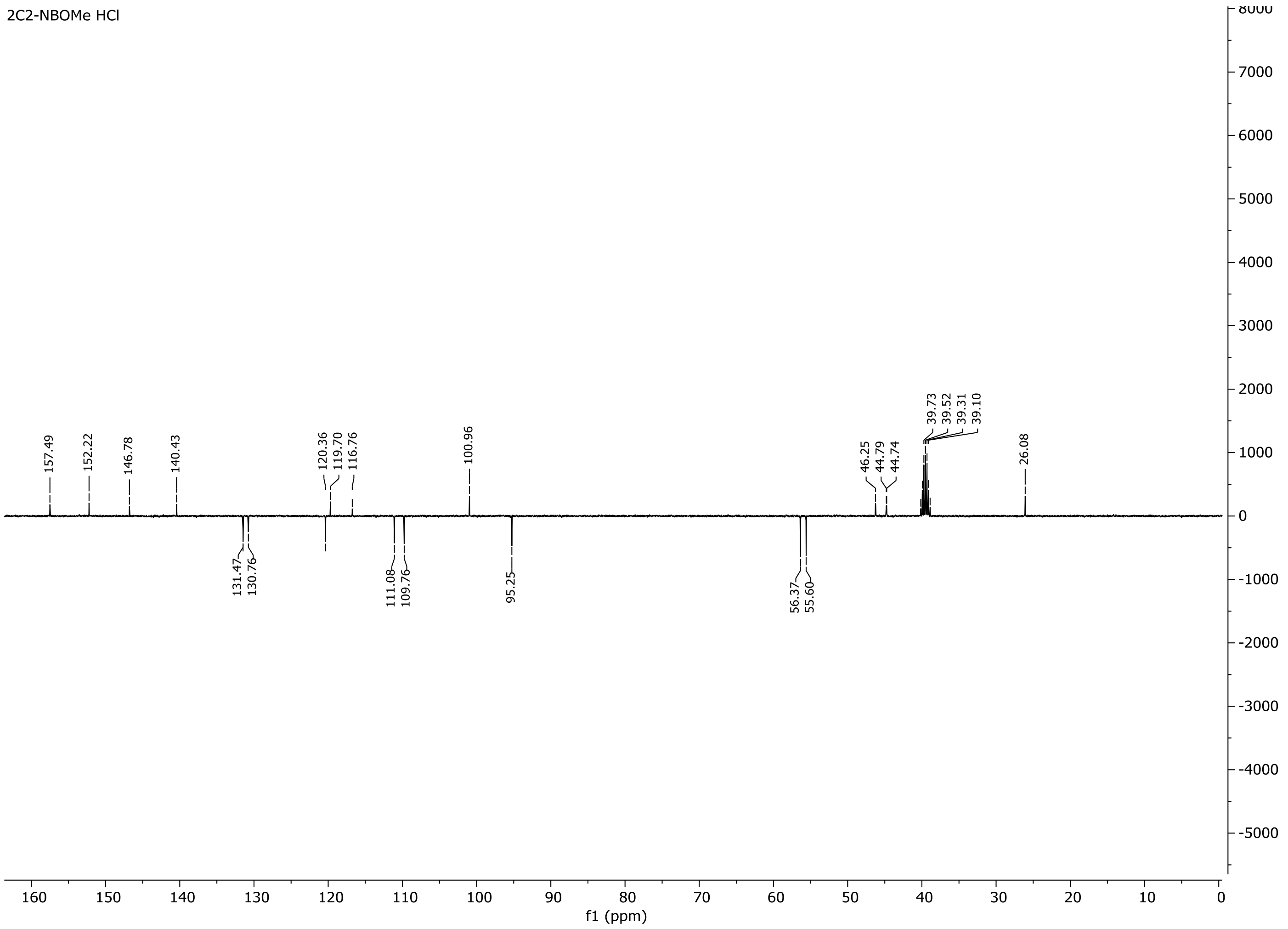

### Spectra 65. $^{13}C$ NMR of 25O-NBOMe (32) HCl

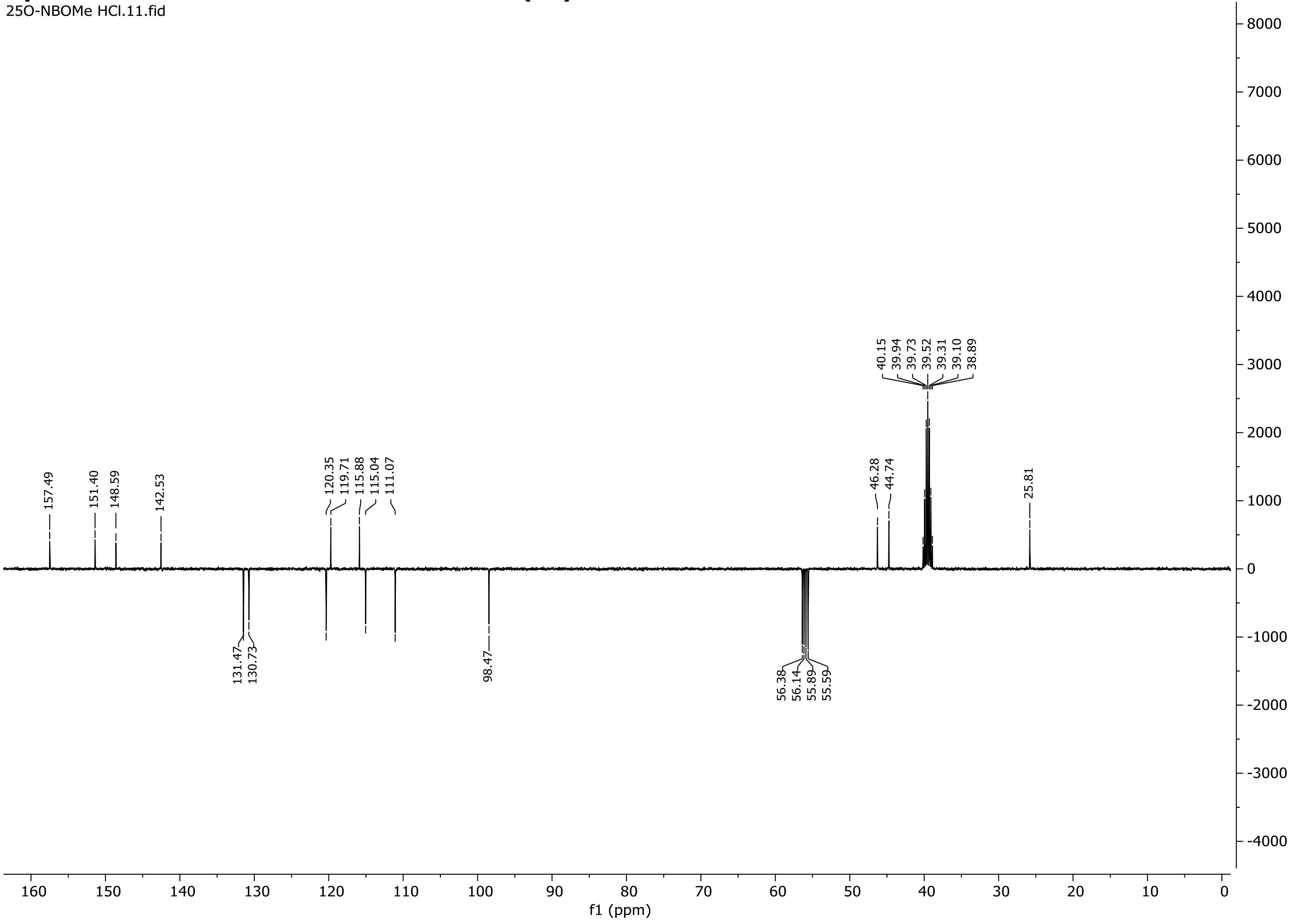

### Spectra 66. $^{13}$C NMR of 25O-NBcP (33) HCl

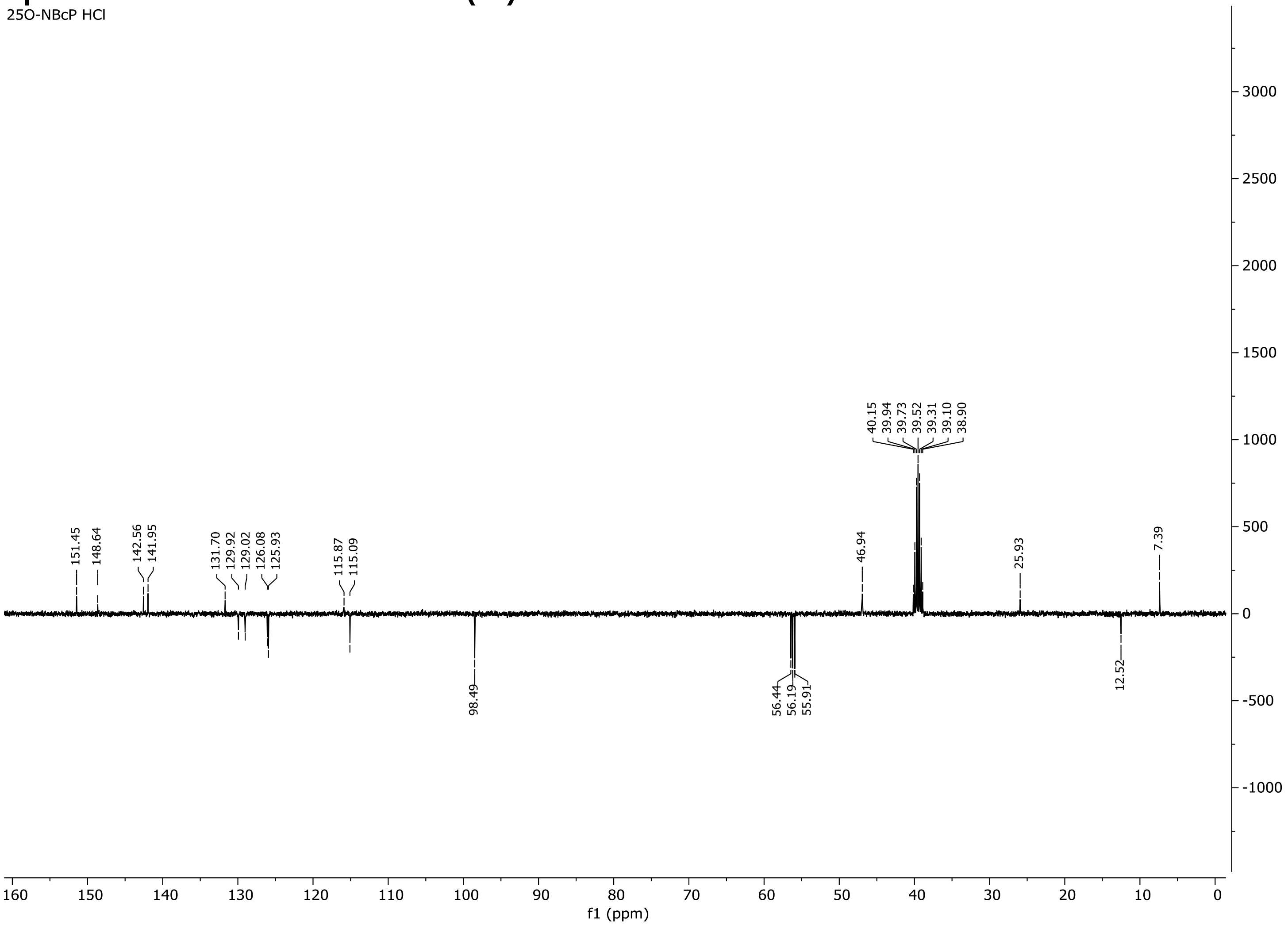

### Spectra 67. $^{13}C$ NMR of 25O-NB-3-I (34) HCl

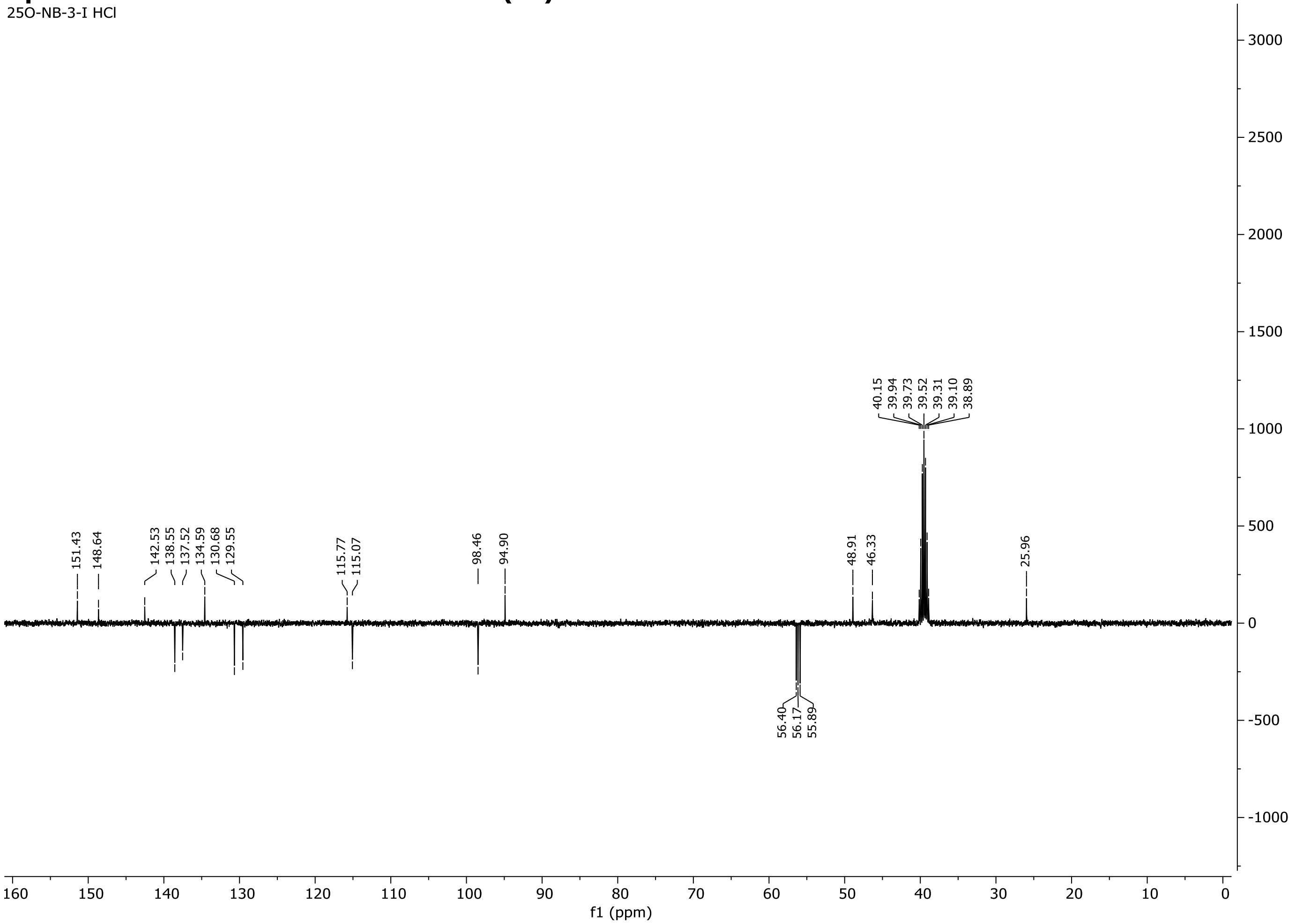

### Spectra 68. $^{13}$C NMR of 25O-NBPh-10'-OH (35) HCl

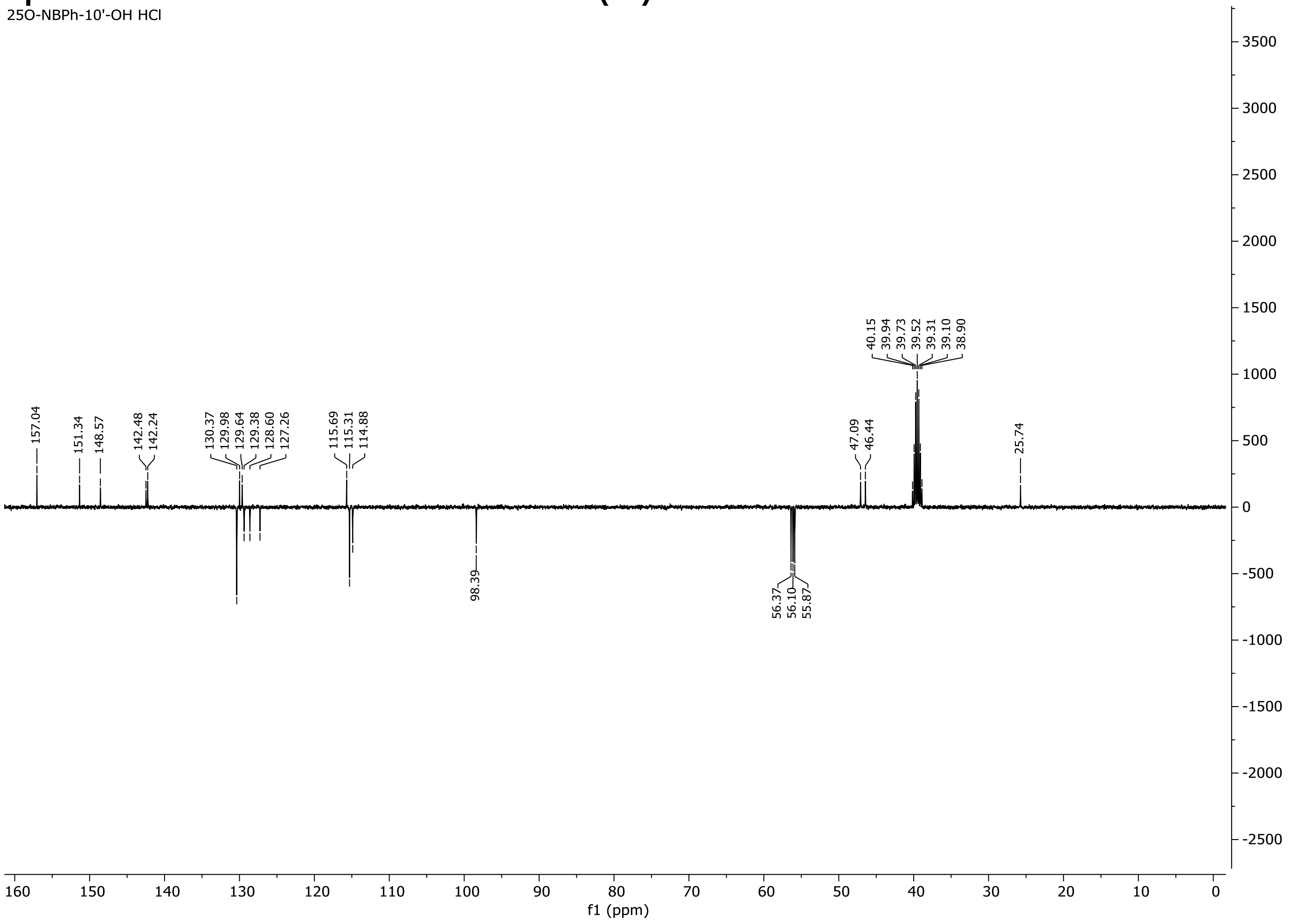

**Spectra 69. 19F NMR of 25N-NBF (7) HCl**

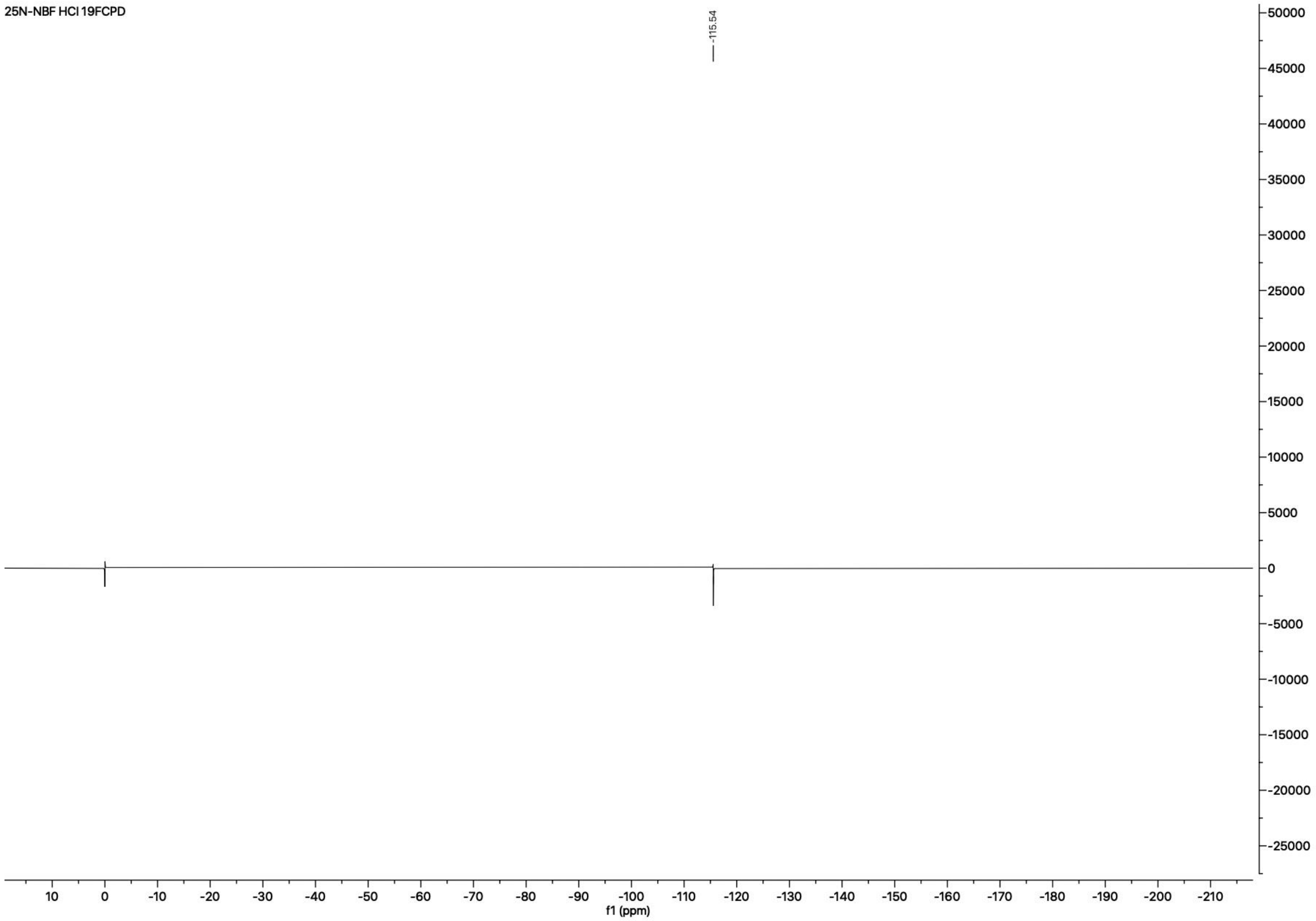

**Spectra 70. $^{19}F$ NMR of 25N-NBOCF$_2$H (11) HCl**

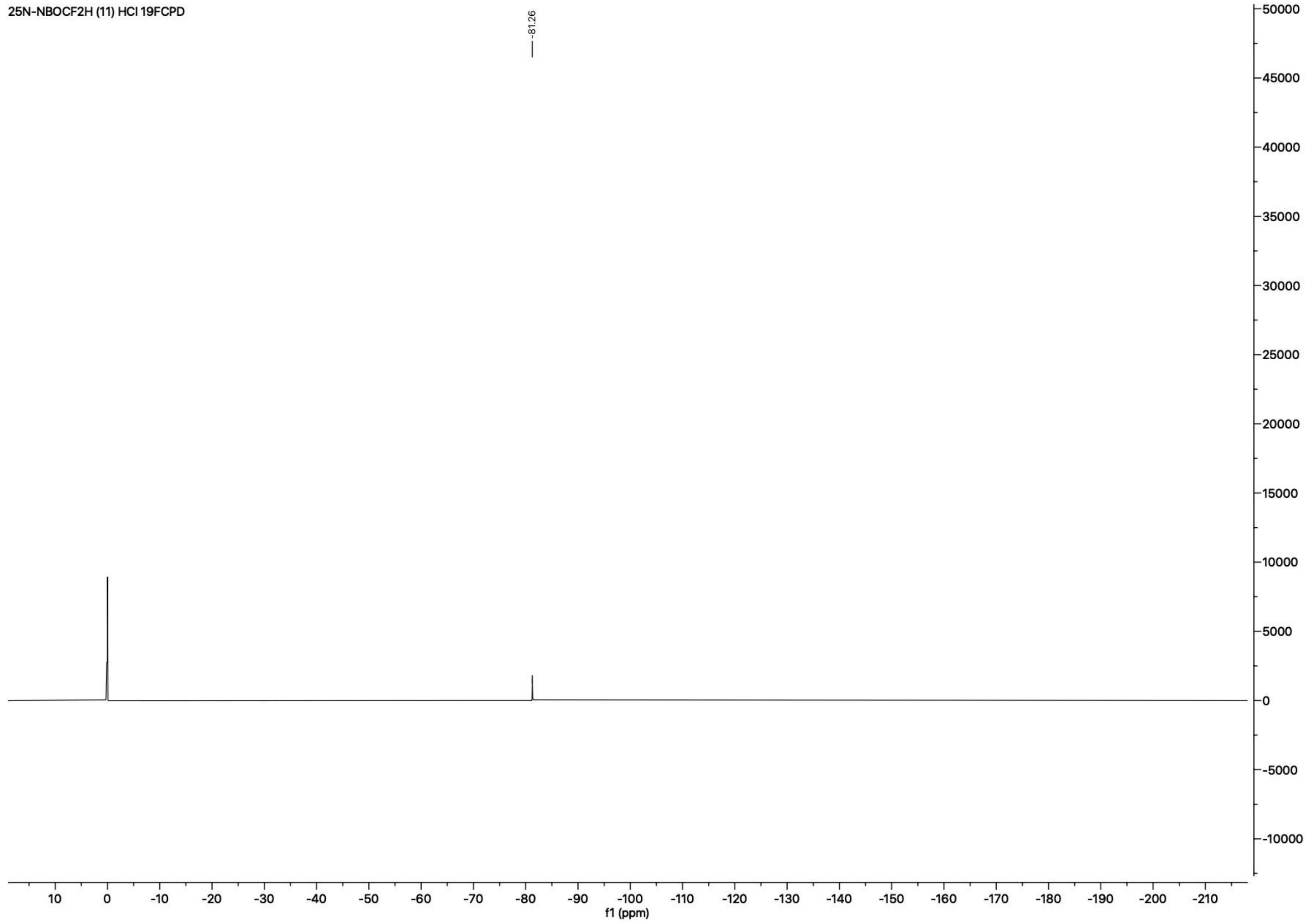

### Spectra 71. $^{19}$F NMR of 25N-NBOCF$_3$ (12) HCl

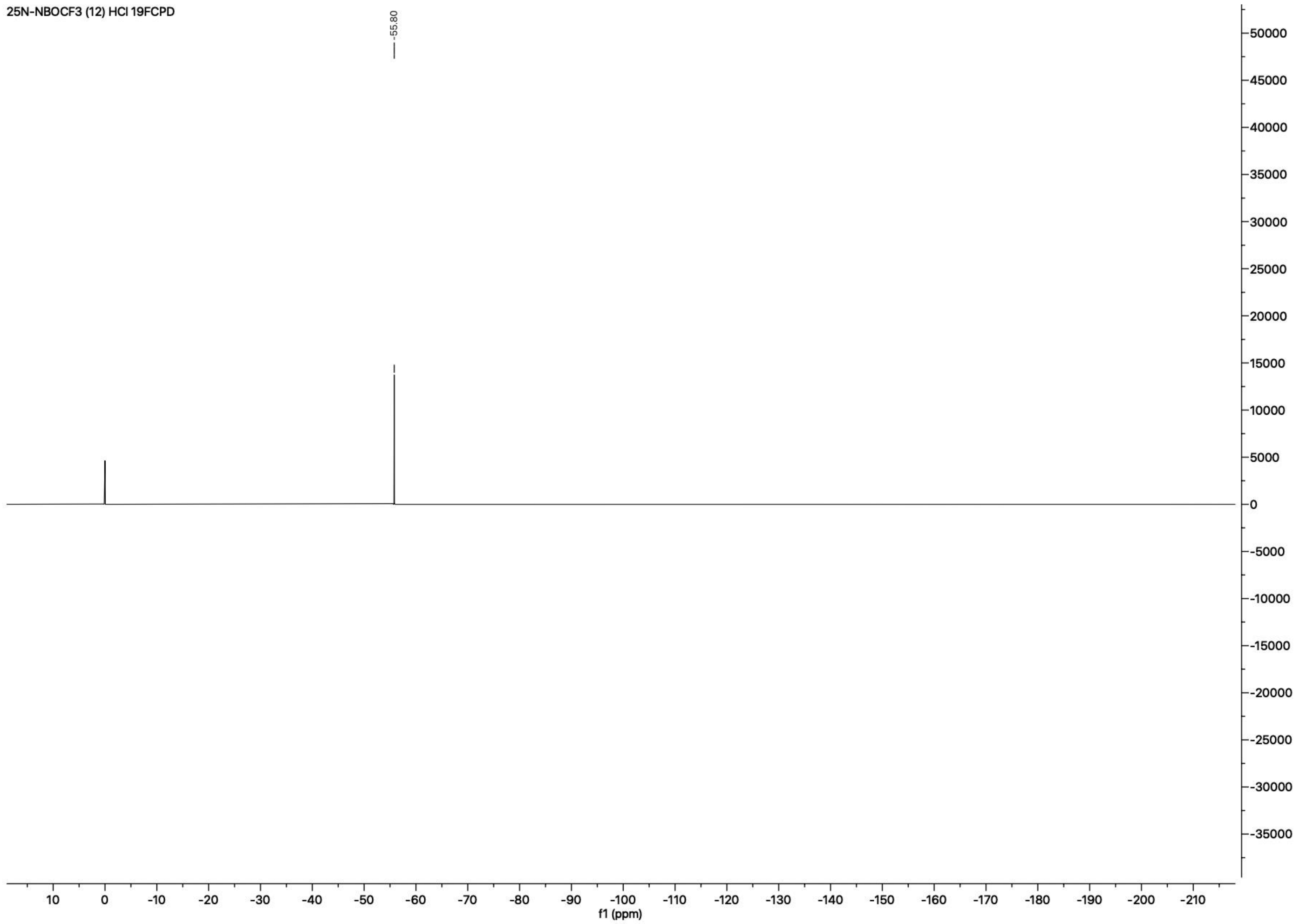

**Spectra 72. $^{19}F$ NMR of 25N-NBMDF$_2$ (13) HCl**

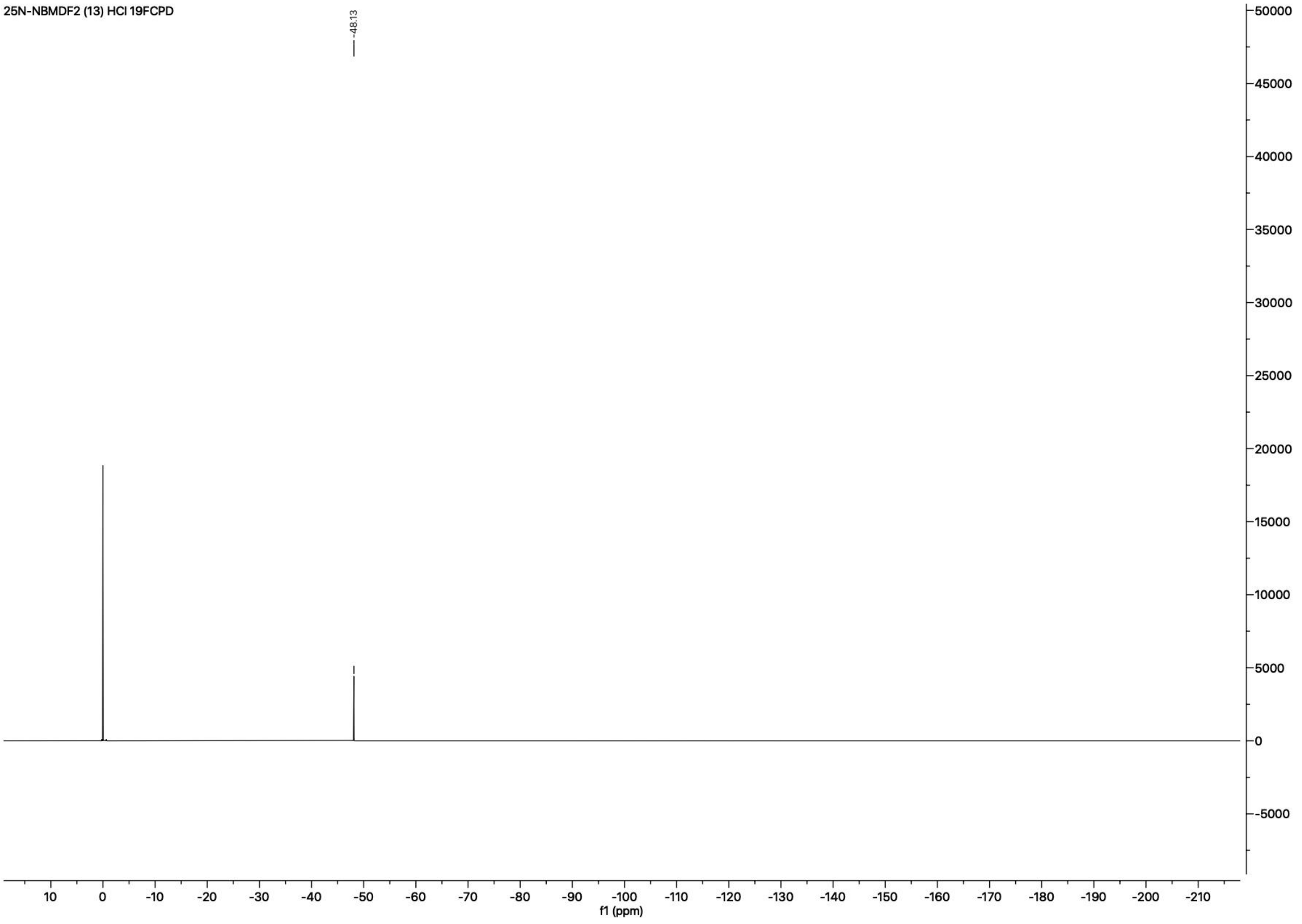

**Spectra 73. $^{19}F$ NMR of 25N-NBCF$_3$ (14) HCl**

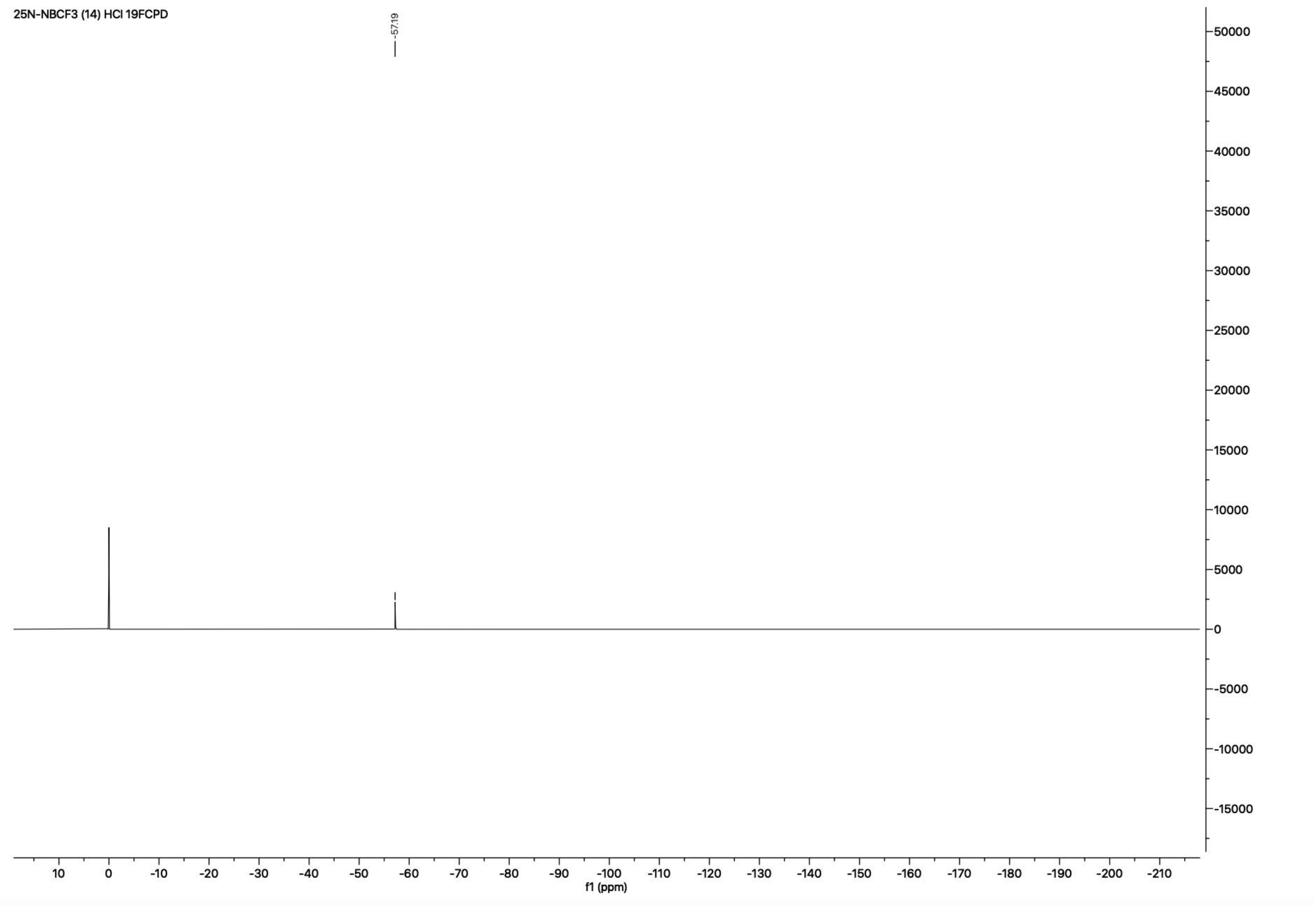

**Spectra 74. $^{19}$F NMR of 25N-NB-2-MeO-3-F (19) HCl**

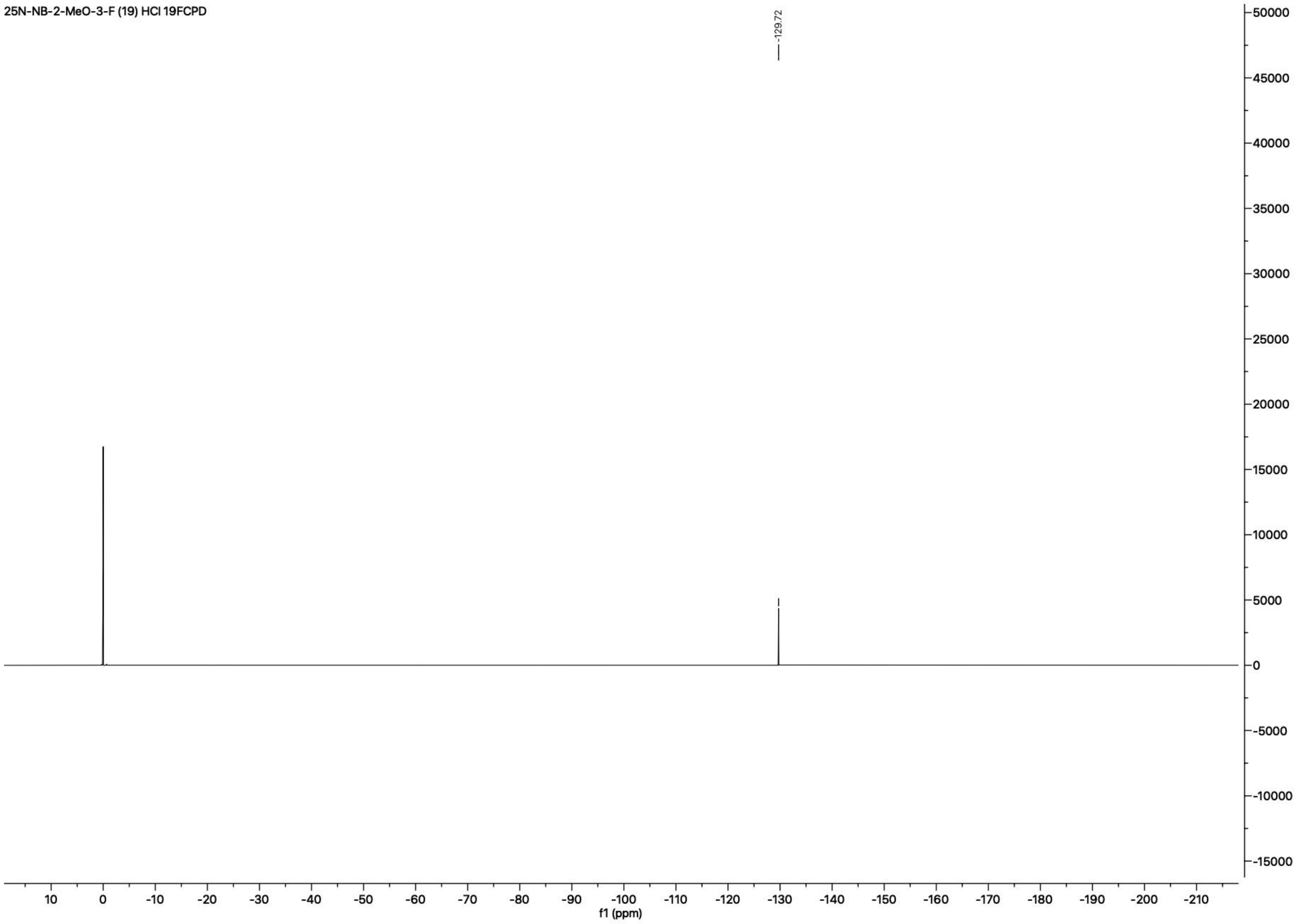

**Spectra 75. $^{19}$F NMR of 25N-NB-3-F (24) HCl**

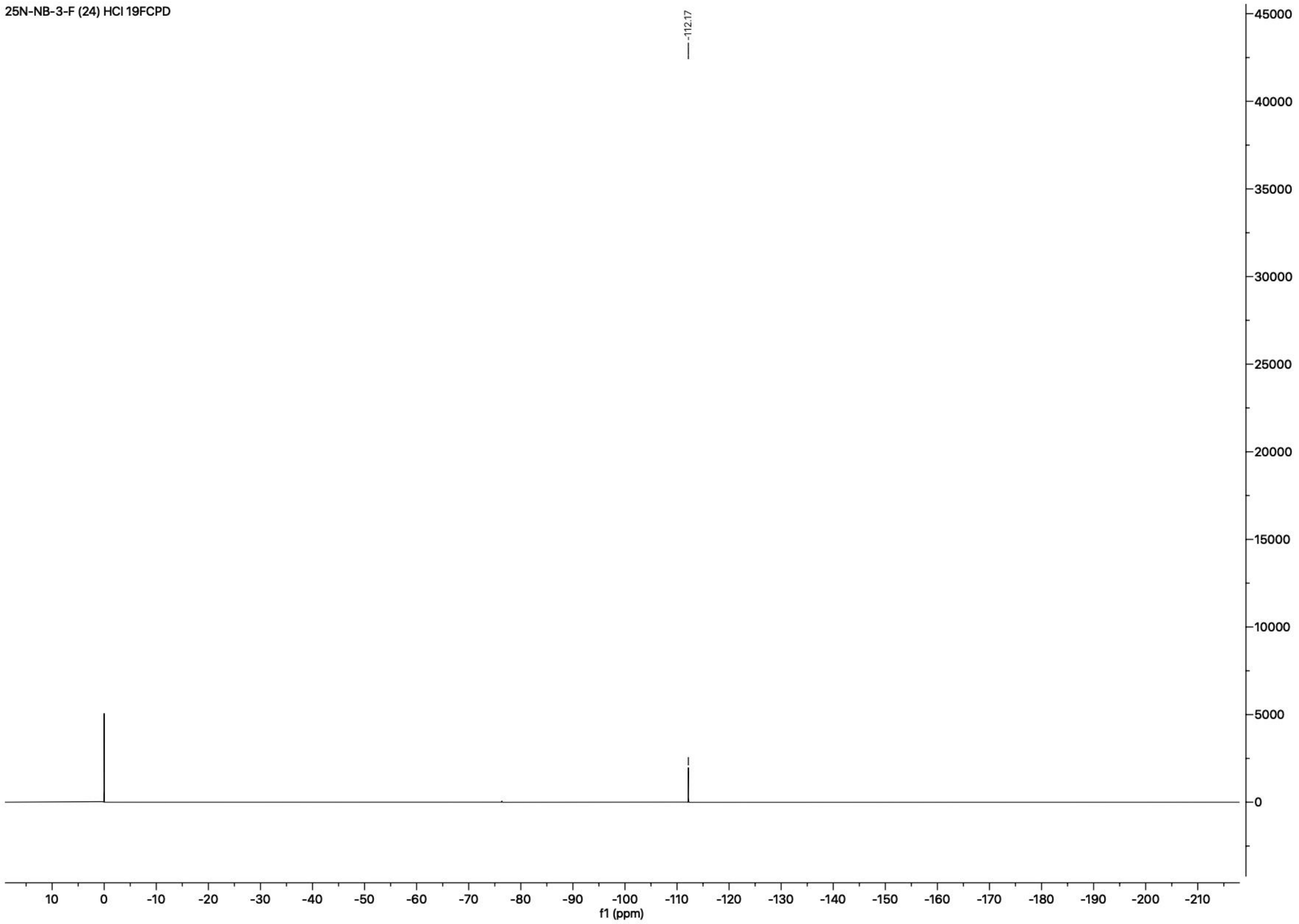

**Spectra 76. $^{19}$F NMR of 25N-NB-4-F (25) HCl**

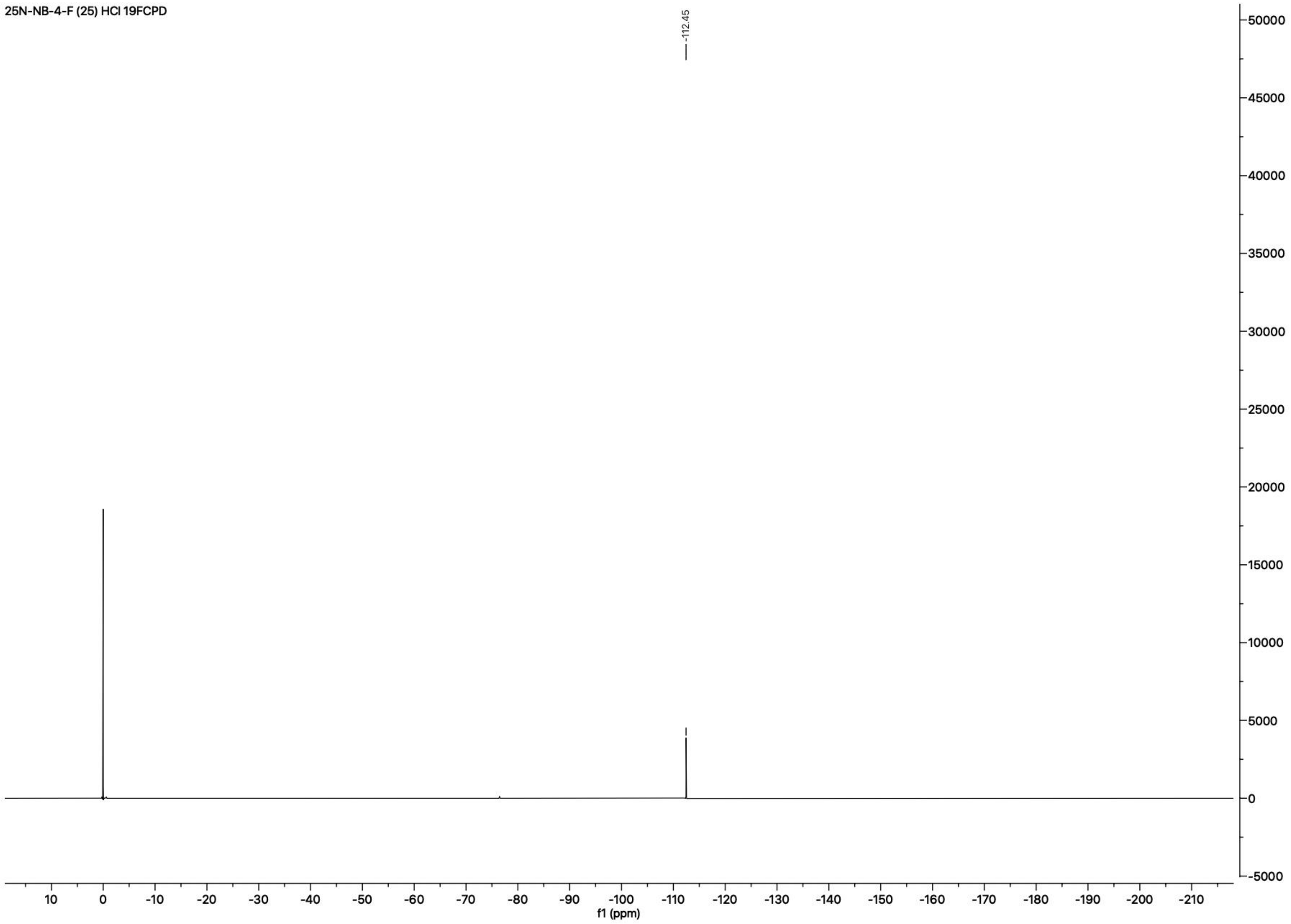

**HPLC Traces of Final *N*-Benzyl Phenethylamines.** HPLC experiments performed on HCl salts at 1 mg/mL.

**HPLC Trace of 25N-NB (2).** Purity (Peak area): 99.5%; Wavelength: 220 nm.

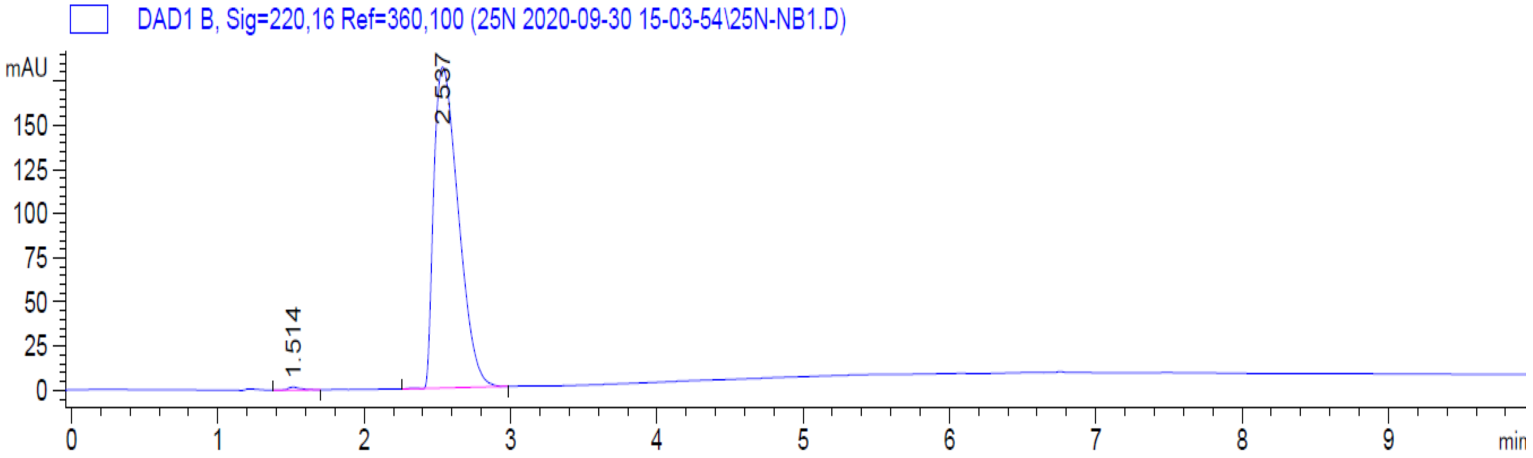

**HPLC Trace of 25N-NBOH (3).** Purity (Peak area): 97.4%; Wavelength: 220 nm.

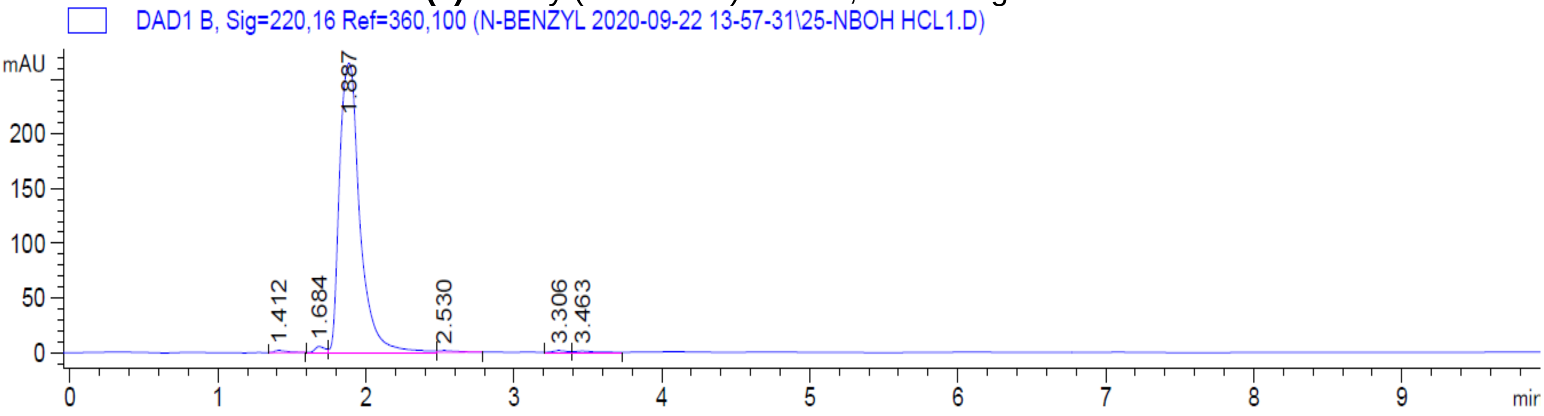

**HPLC Trace of 25N-NBOMe (4).** Purity (Peak area): 98.7%; Wavelength: 220 nm.

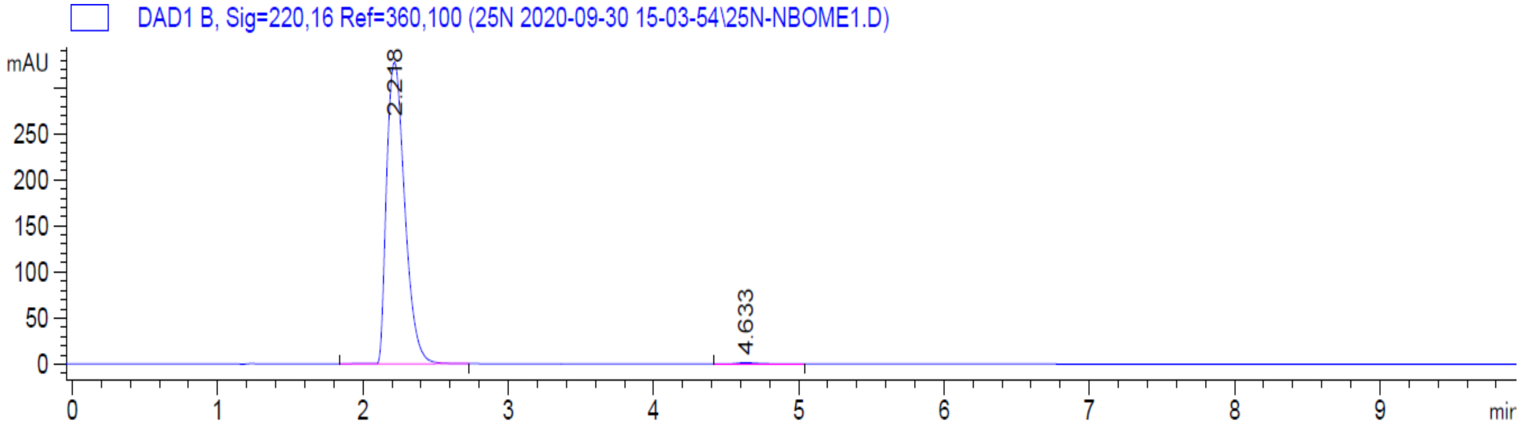

**HPLC Trace of 25N-NBOEt (5).** Purity (Peak area): 98.8%; Wavelength: 220 nm.

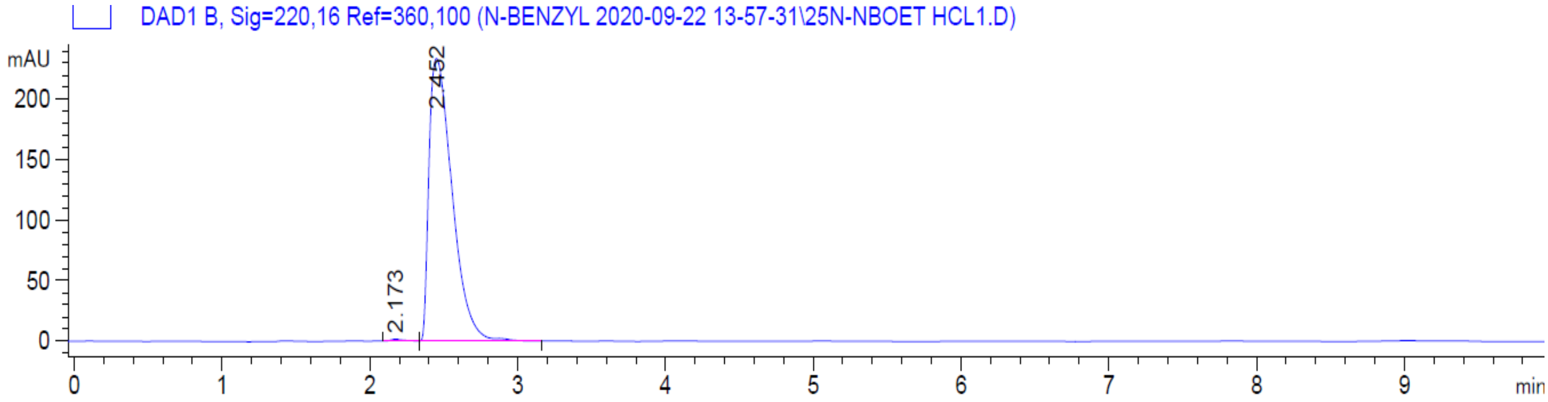

**HPLC Trace of 25N-NBMe (6).** Purity (Peak area): 98.4%; Wavelength: 220 nm.

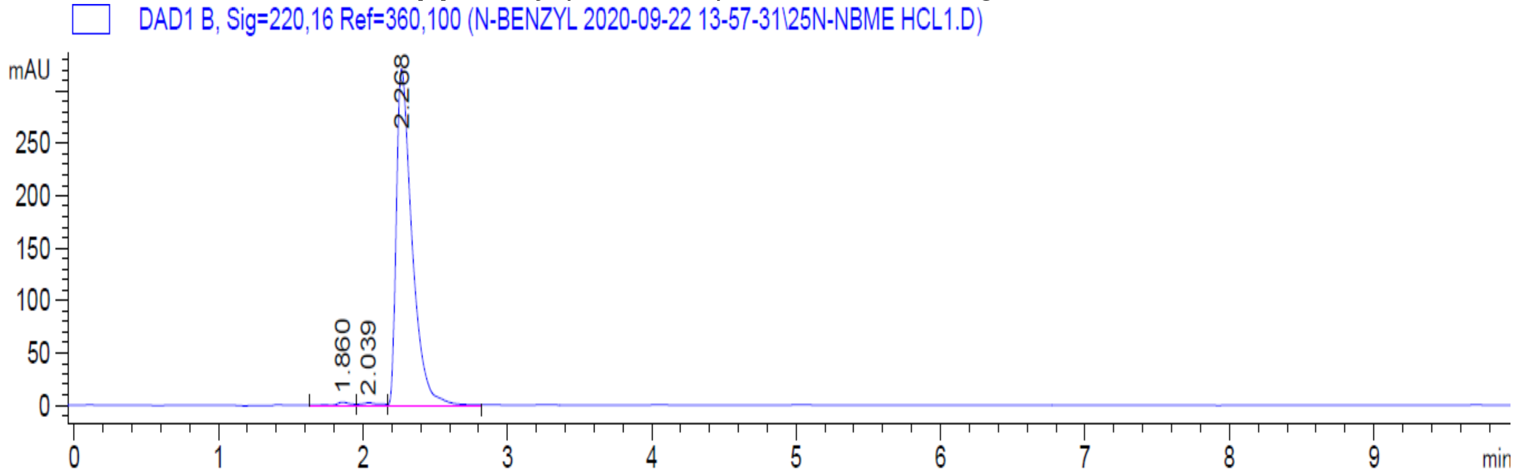

**HPLC Trace of 25N-NBF (7).** Purity (Peak area): 100%; Wavelength: 220 nm.

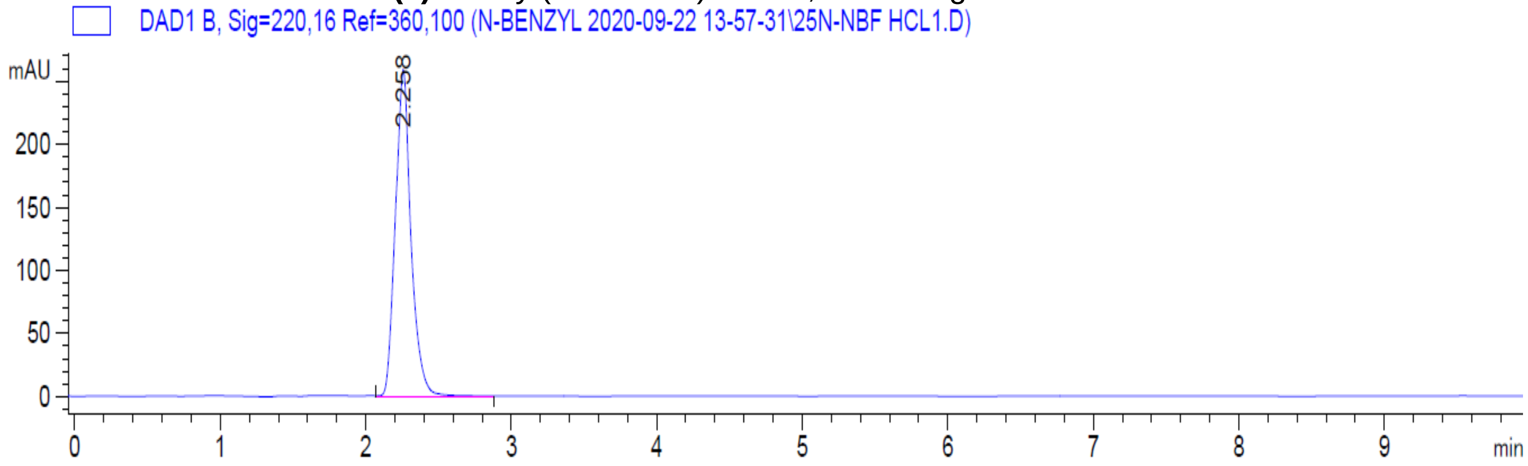

**HPLC Trace of 25N-NBCl (8).** Purity (Peak area): 98.8%; Wavelength: 220 nm.

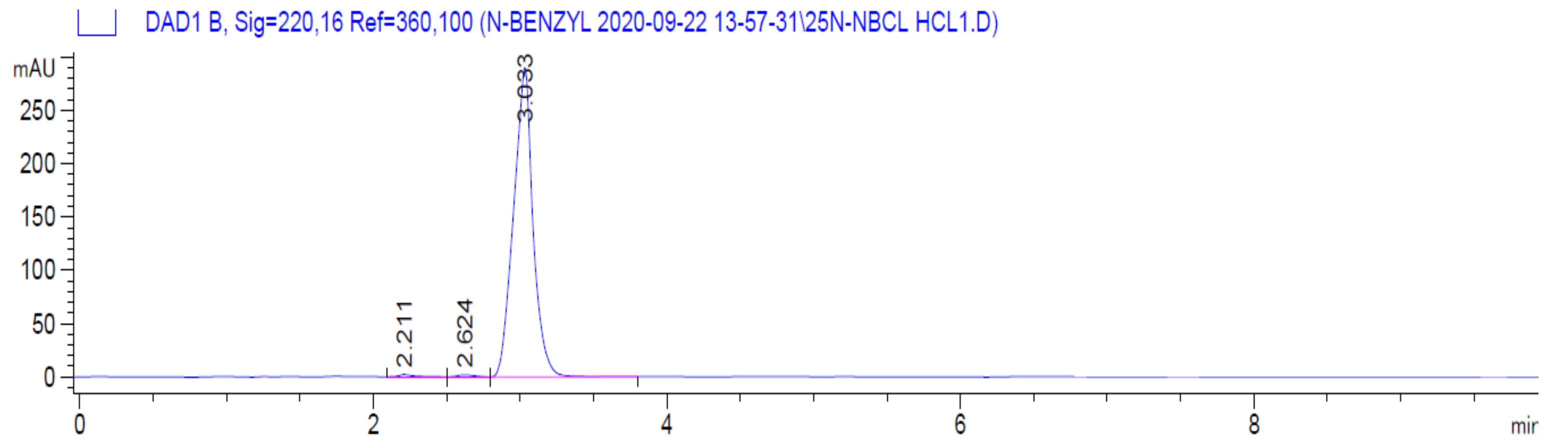

**HPLC Trace of 25N-NBBr (9).** Purity (Peak area): 99.2%; Wavelength: 220 nm.

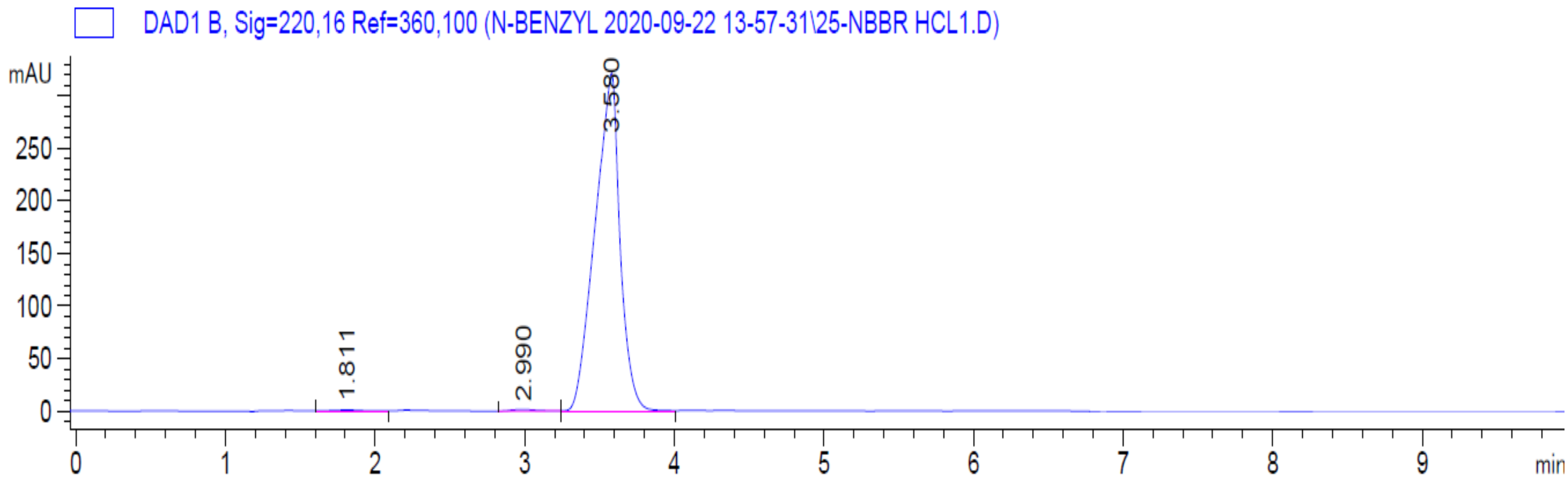

**HPLC Trace of 25N-NBI (10).** Purity (Peak area): 99.0%; Wavelength: 220 nm.

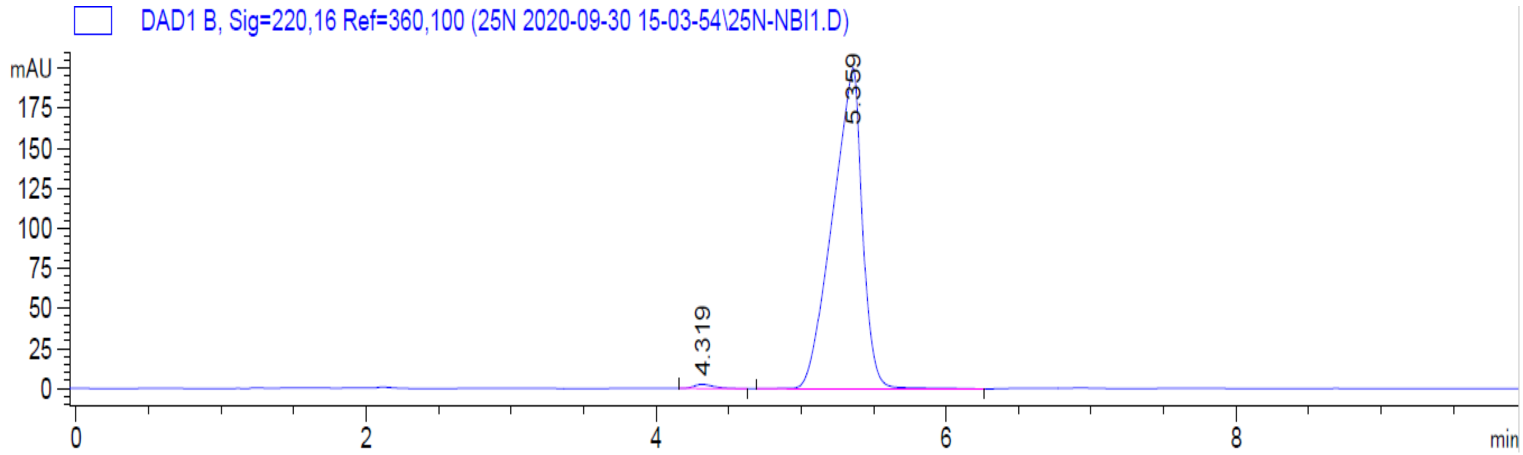

**HPLC Trace of 25N-NBOCF$_2$H (11).** Purity (Peak area): 93.1%; Wavelength: 220 nm.

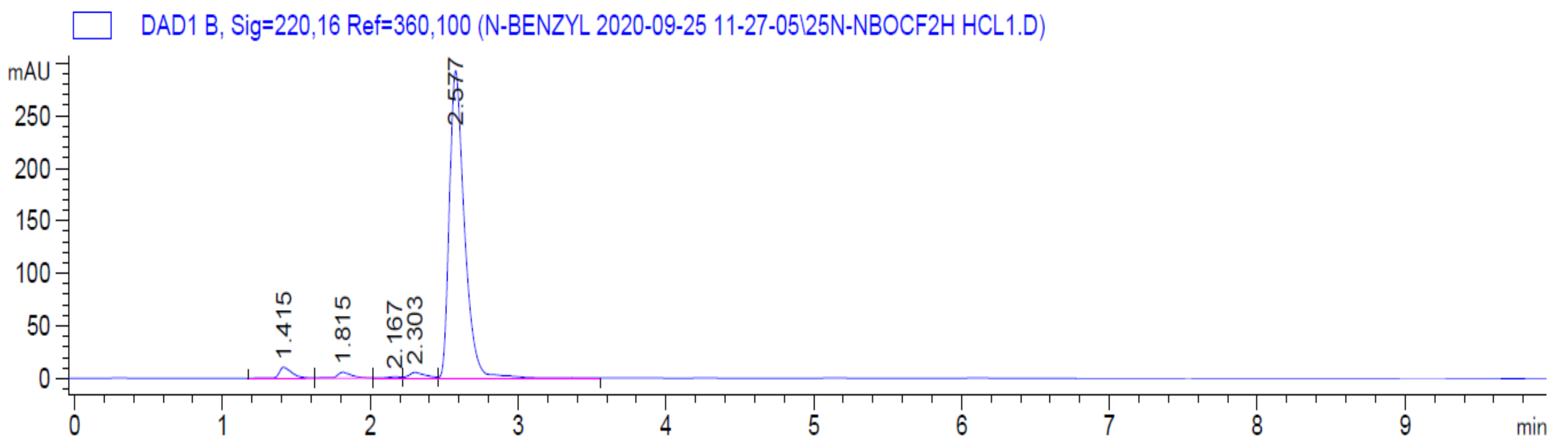

**HPLC Trace of 25N-NBOCF$_3$ (12).** Purity (Peak area): 98.7%; Wavelength: 220 nm.

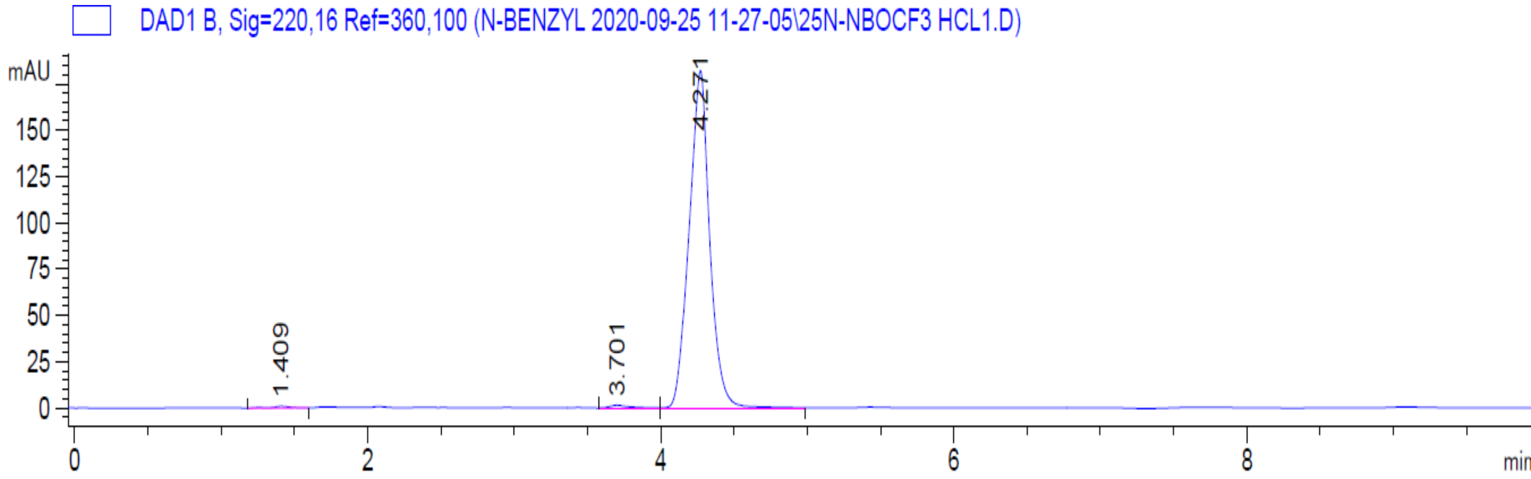

**HPLC Trace of 25N-NBMDF$_2$ (13).** Purity (Peak area): 98.9%; Wavelength: 220 nm.

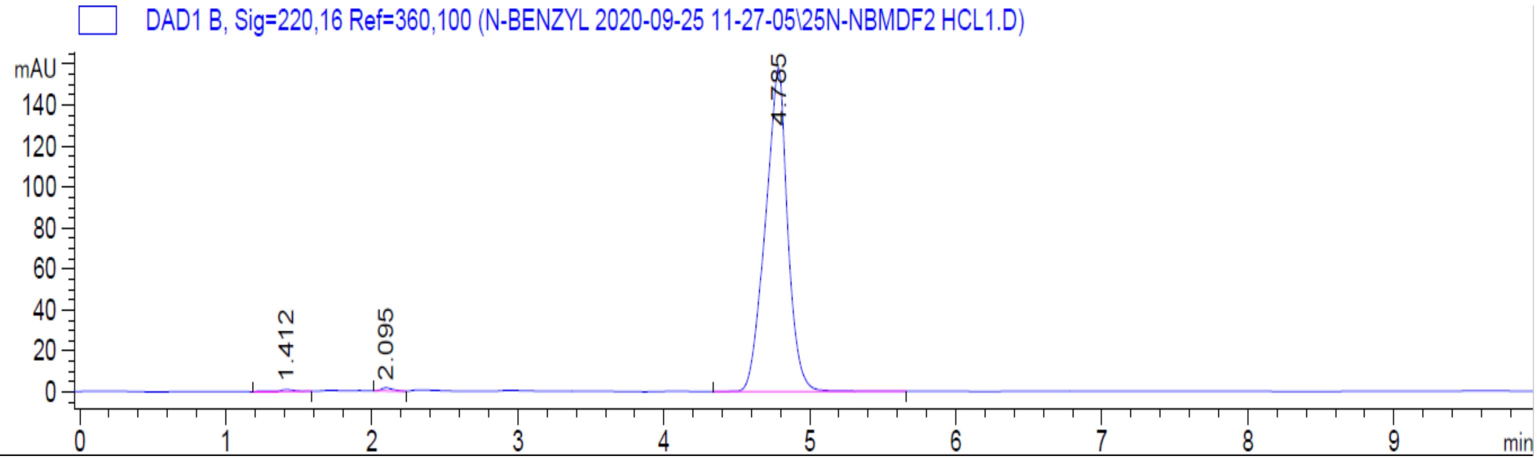

**HPLC Trace of 25N-NBCF$_3$ (14).** Purity (Peak area): 99.2%; Wavelength: 220 nm.

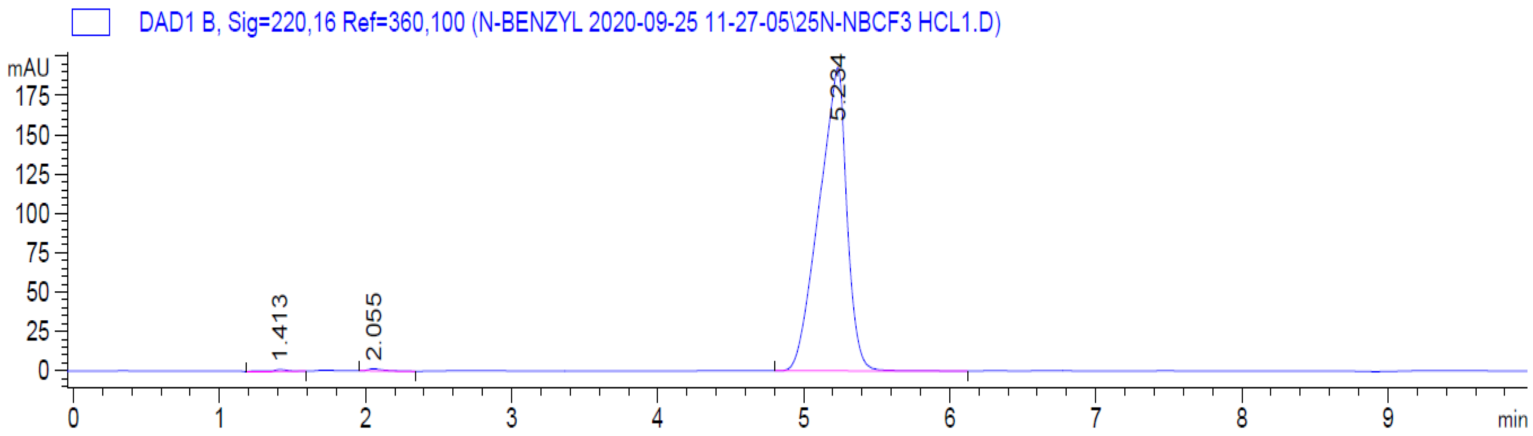

**HPLC Trace of 25N-NBNO$_2$ (15).** Purity (Peak area): 99.4%; Wavelength: 220 nm.

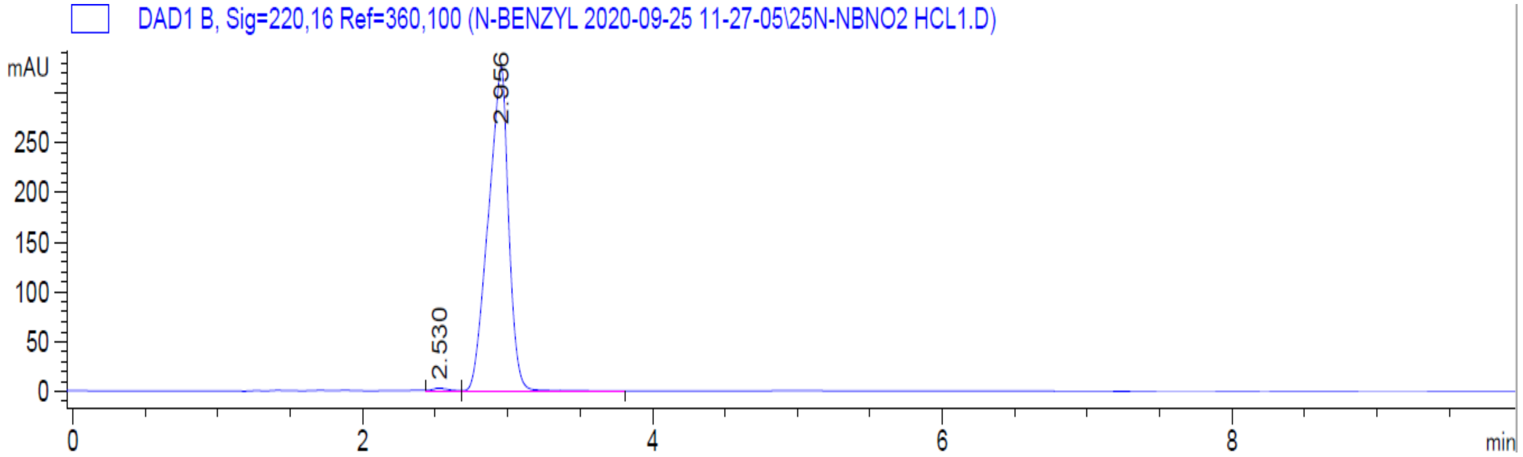

**HPLC Trace of 25N-N1-Nap (16).** Purity (Peak area): 99.2%; Wavelength: 220 nm.

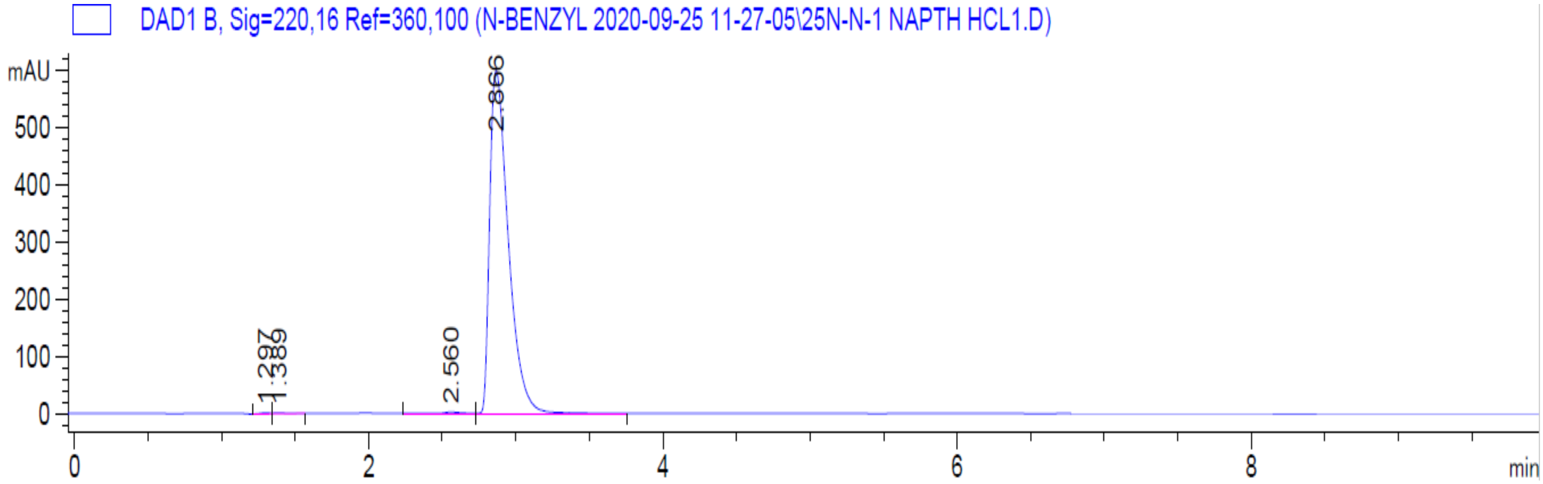

**HPLC Trace of 25N-NBPh (17).** Purity (Peak area): 98.7%; Wavelength: 220 nm.

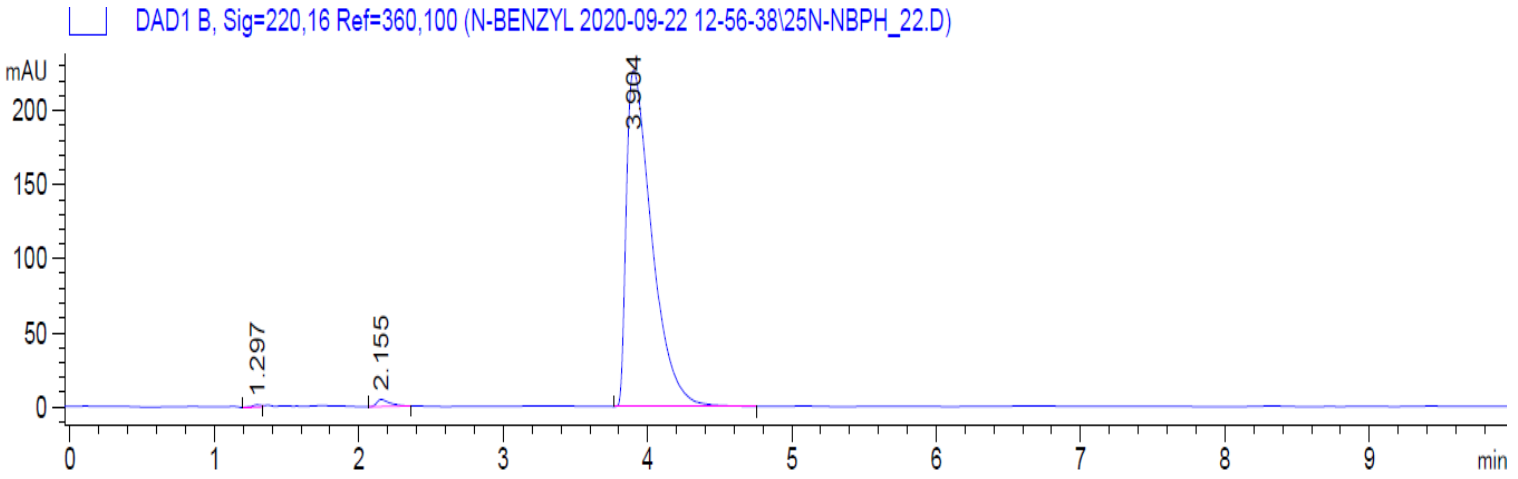

**HPLC Trace of 25N-NB-2-OH-3-Me (18).** Purity (Peak area): 100%; Wavelength: 220 nm.

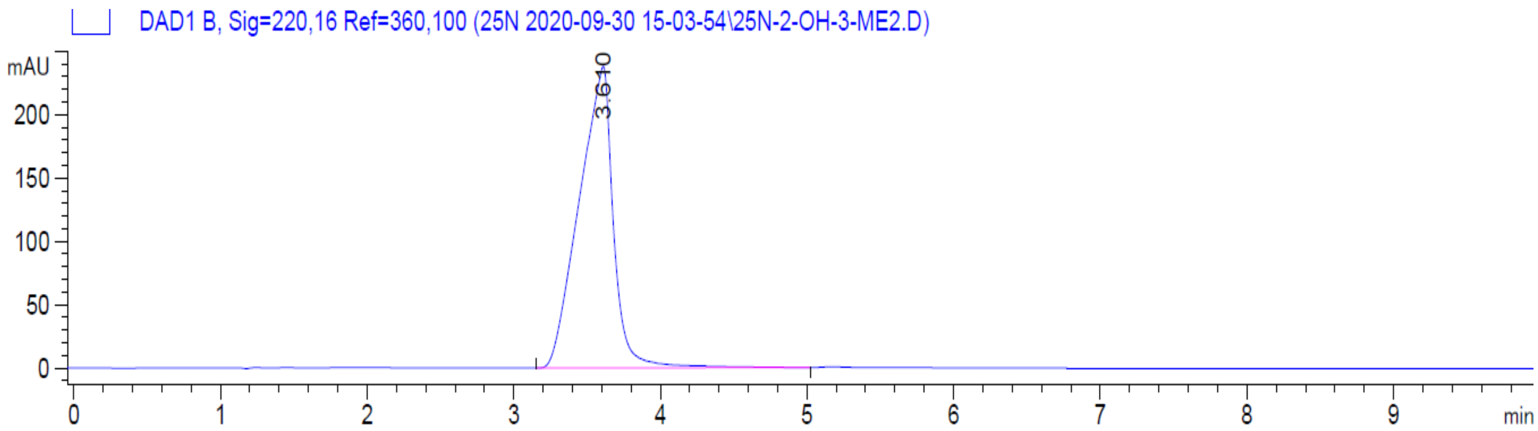

**HPLC Trace of 25N-NB-2-MeO-3-F (19).** Purity (Peak area): 96.1%; Wavelength: 220 nm.

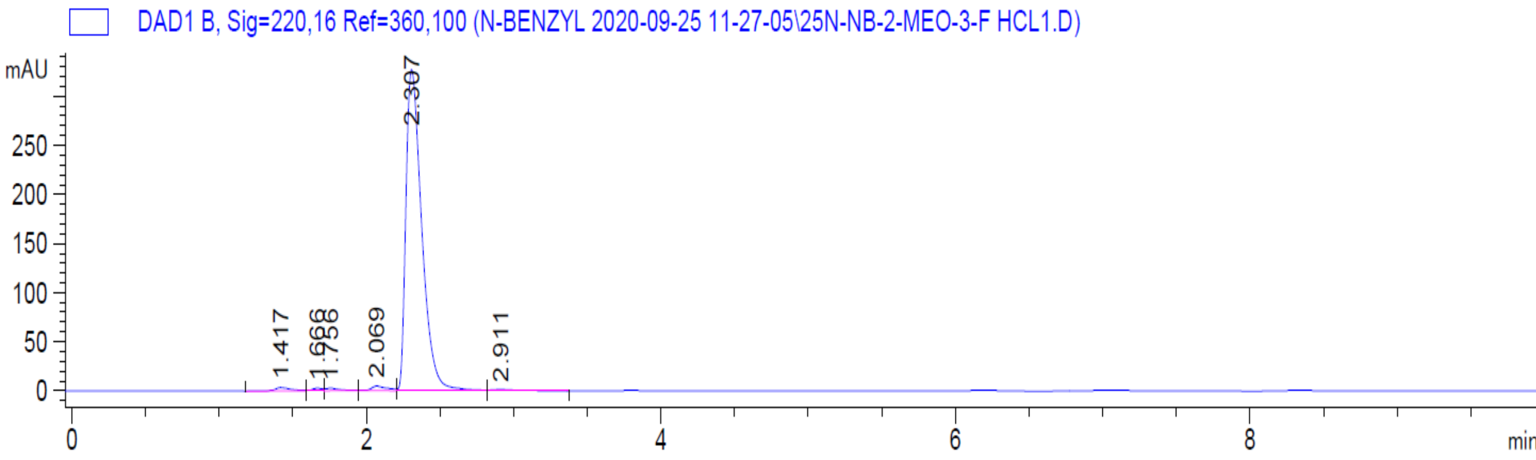

**HPLC Trace of 25N-NB-2,5-DiMeO (20).** Purity (Peak area): 95.6%; Wavelength: 220 nm.

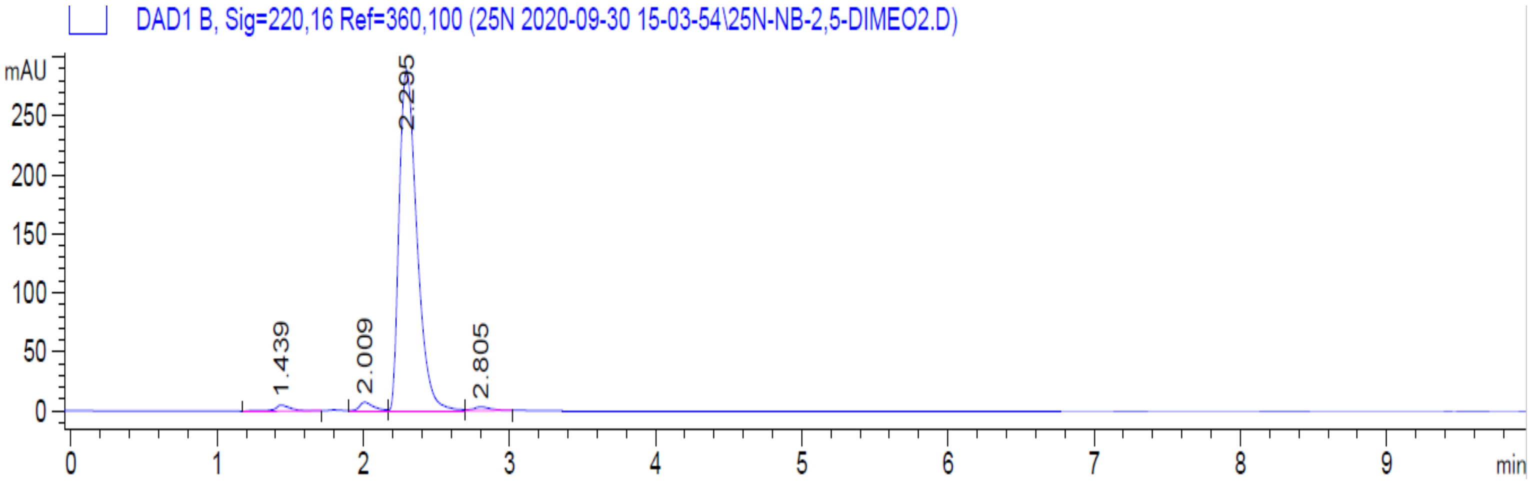

**HPLC Trace of 25N-NB-3-OH (21).** Purity (Peak area): 99.1%; Wavelength: 220 nm.

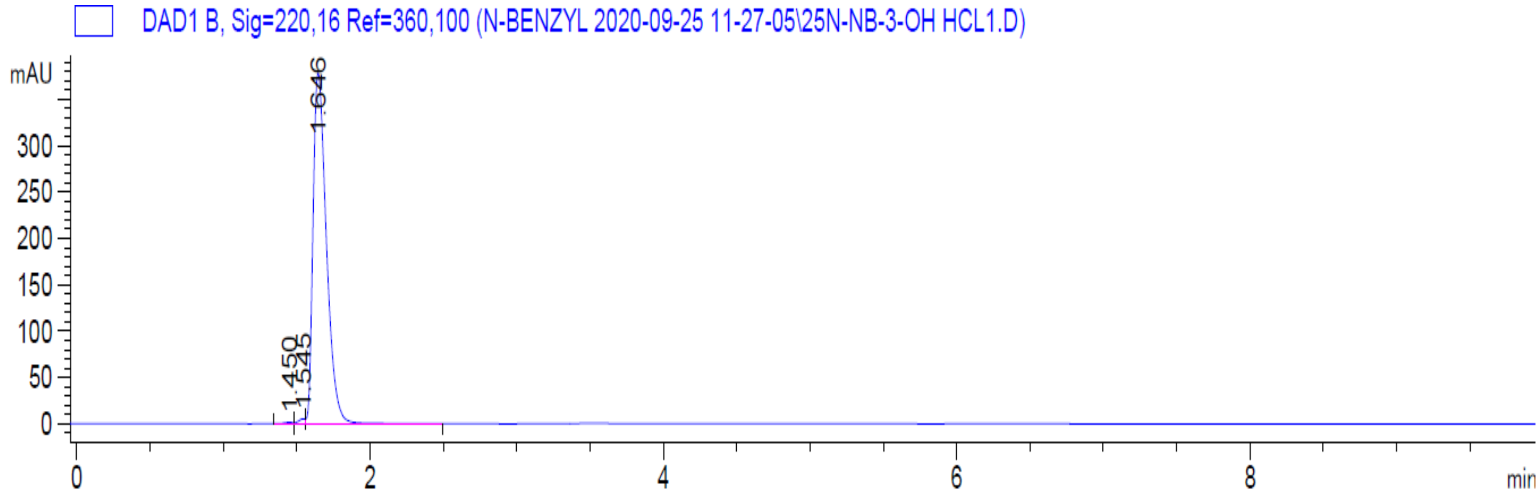

**HPLC Trace of 25N-NB-3-Me (22).** Purity (Peak area): 98.8%; Wavelength: 220 nm.

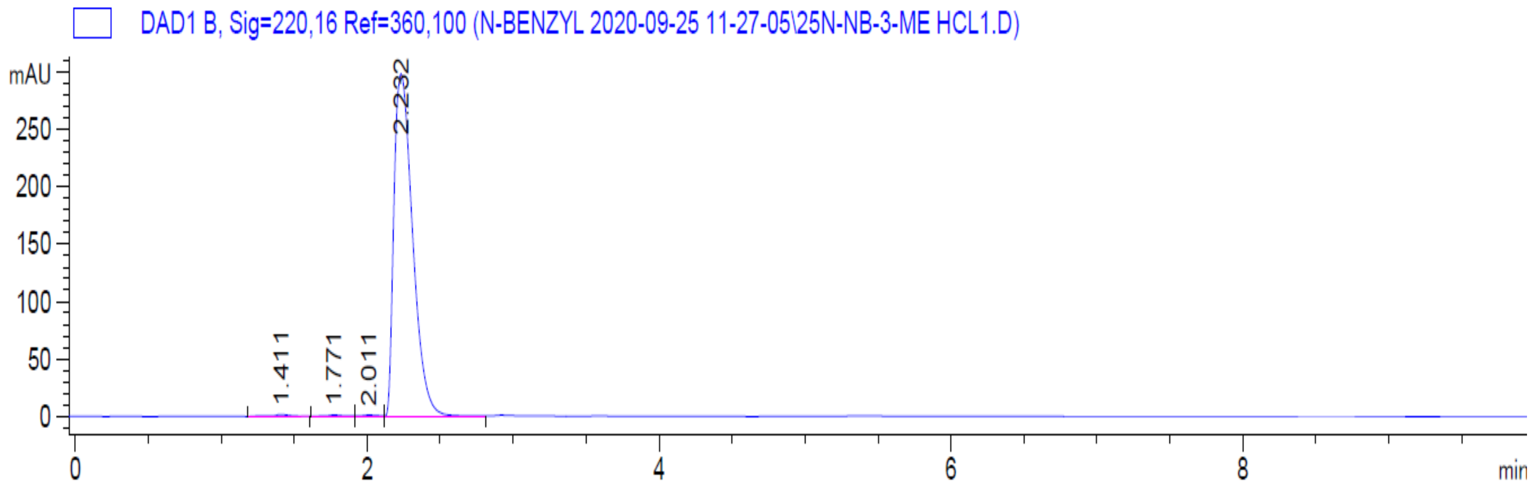

**HPLC Trace of 25N-NB-4-Me (23).** Purity (Peak area): 98.4%; Wavelength: 220 nm.

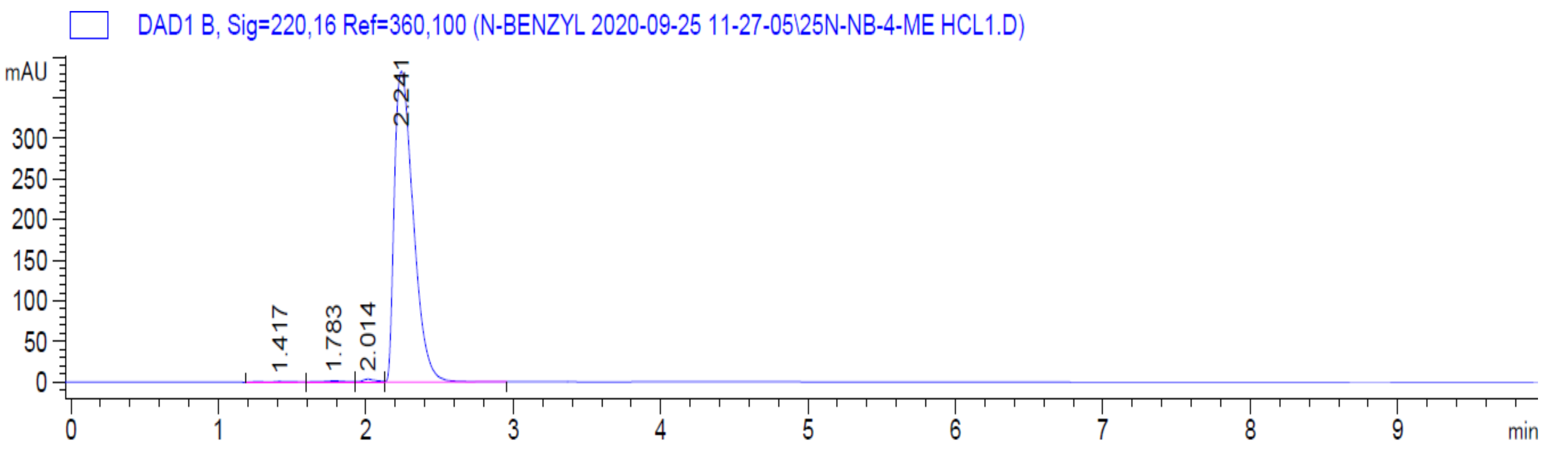

**HPLC Trace of 25N-NB-3-F (24).** Purity (Peak area): 100%; Wavelength: 220 nm.

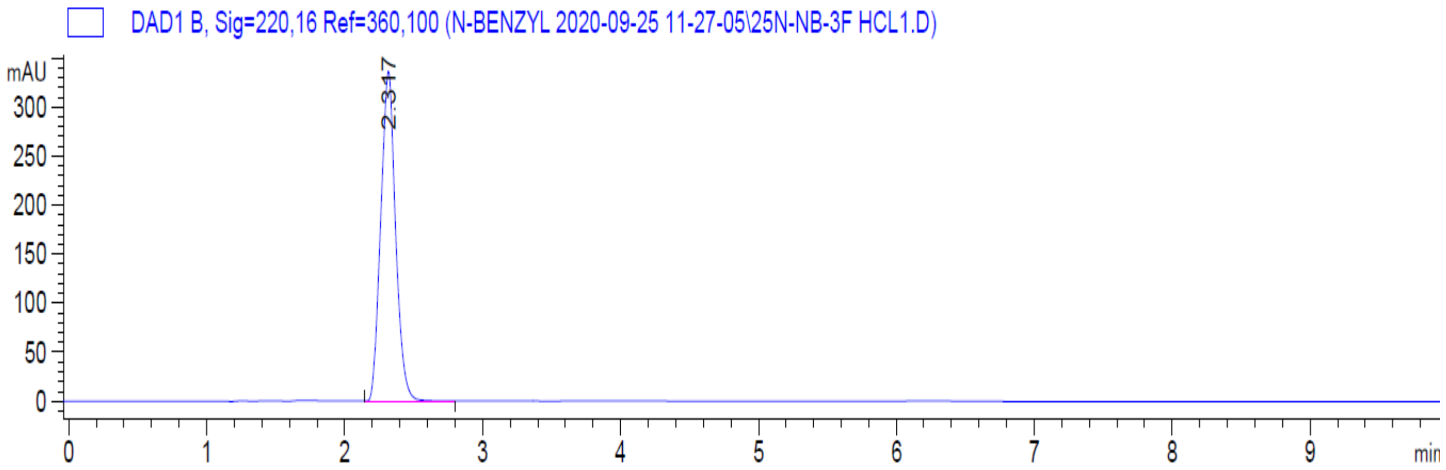

**HPLC Trace of 25N-NB-4-F (25).** Purity (Peak area): 99.4%; Wavelength: 220 nm.

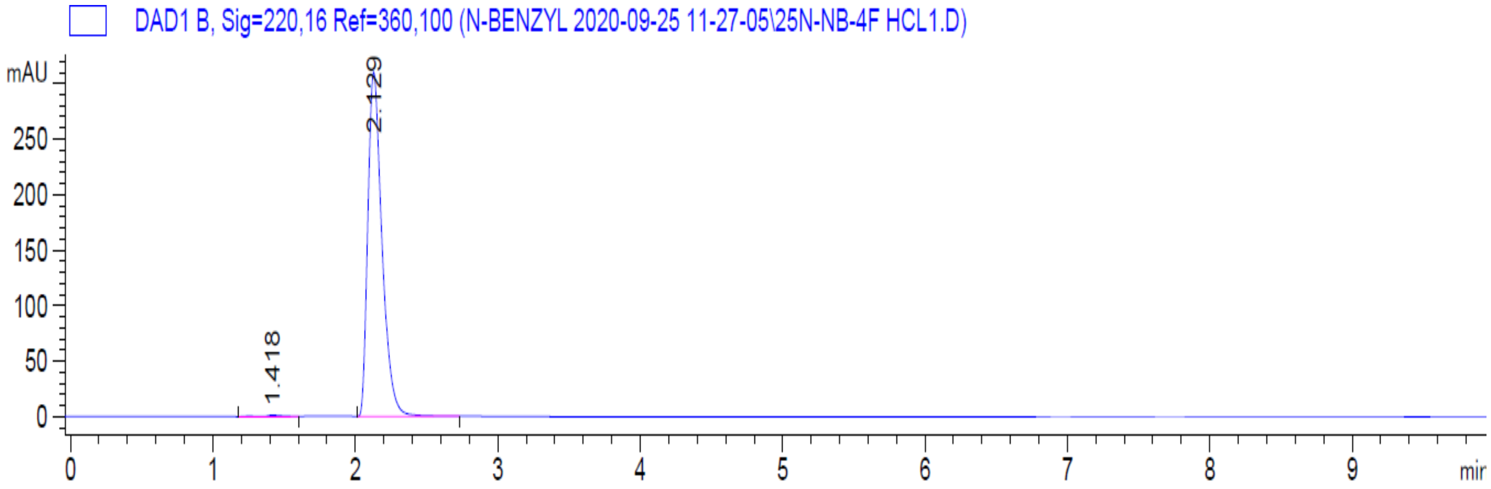

**HPLC trace of 25D-N1-Nap (26).** Purity (Peak area): 100%; Wavelength: 220 nm.

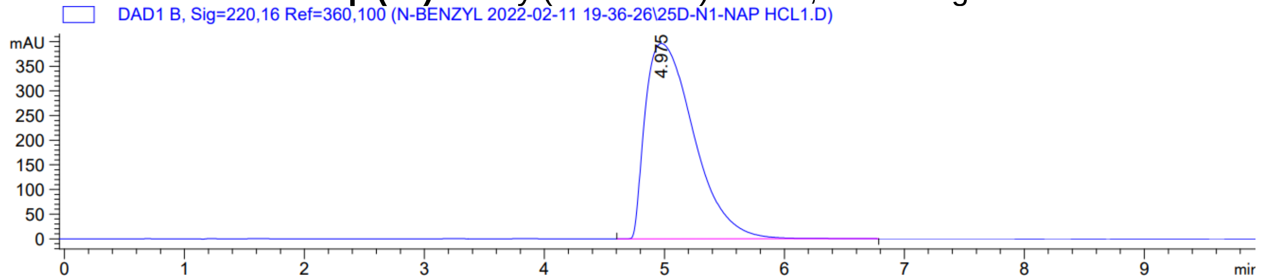

HPLC trace of **25D-NBPh (27).** Purity (Peak area): 98.9%; Wavelength: 220 nm.

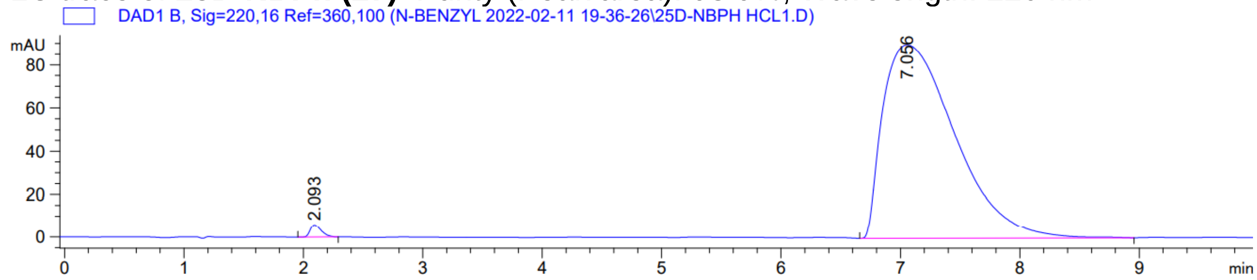

**HPLC trace of 25O-N1-Nap (28).** Purity (Peak area): 100%; Wavelength: 220 nm.

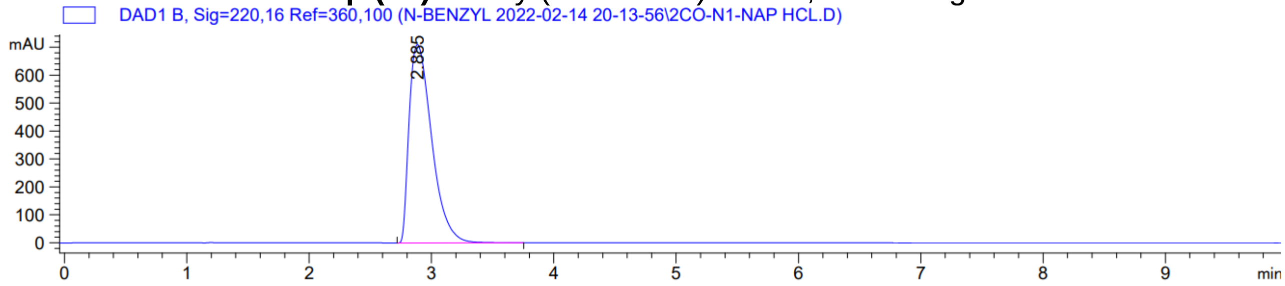

**HPLC trace of 2C2-N1-Nap (29).** Purity (Peak area): 99.8%; Wavelength: 220 nm.

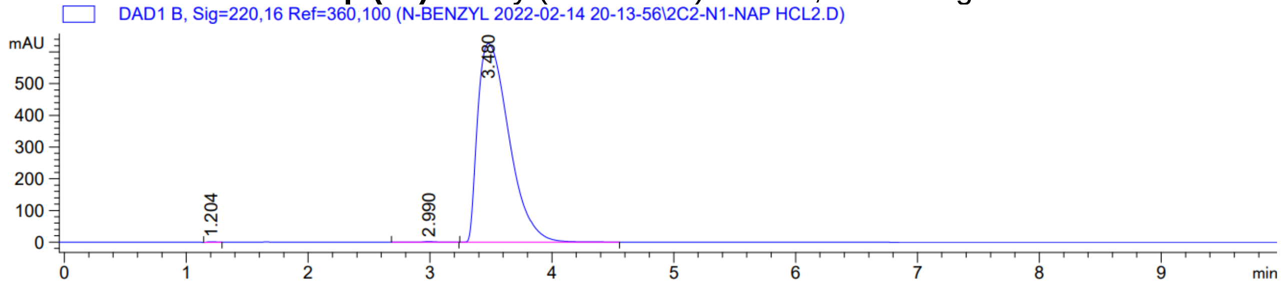

**HPLC trace of Psi-DOB-N1-Nap (30).** Purity (Peak area): 98.3%; Wavelength: 220 nm.

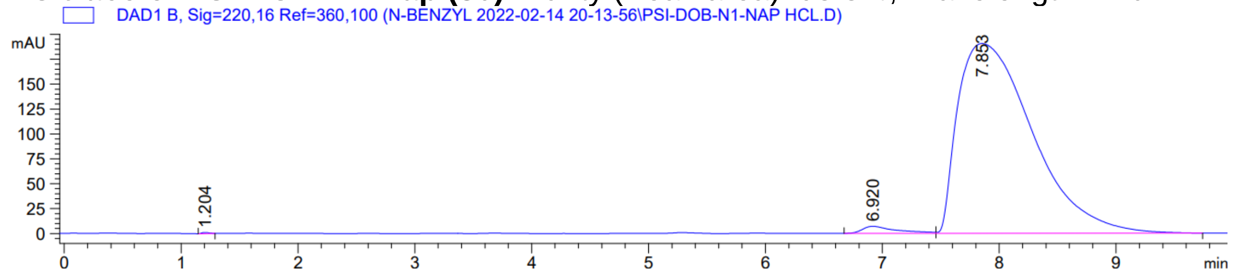

**HPLC Trace of 2C2-NBOMe (31).** Purity (Peak area): 100%; Wavelength: 220 nm.

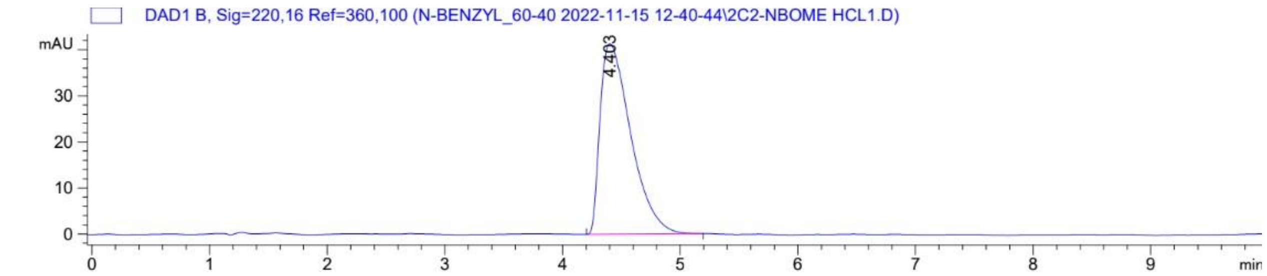

**HPLC Trace of 25O-NBOMe (32).** Purity (Peak area): 99.5%; Wavelength: 220 nm.

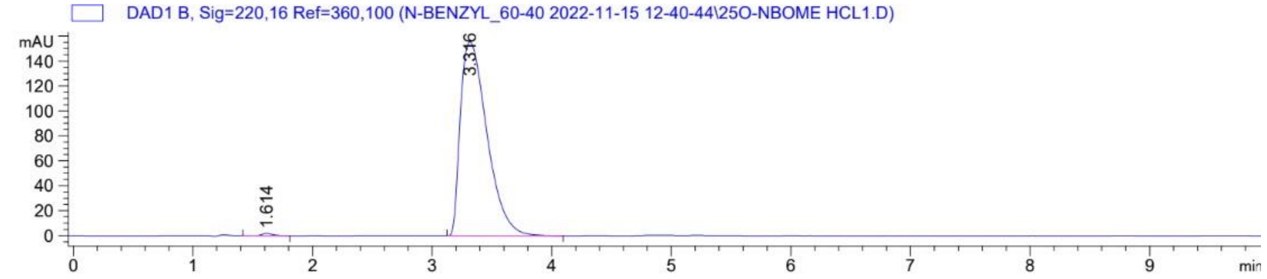

**HPLC Trace of 25O-NBcP (33).** Purity (Peak area): 100%; Wavelength: 220 nm.

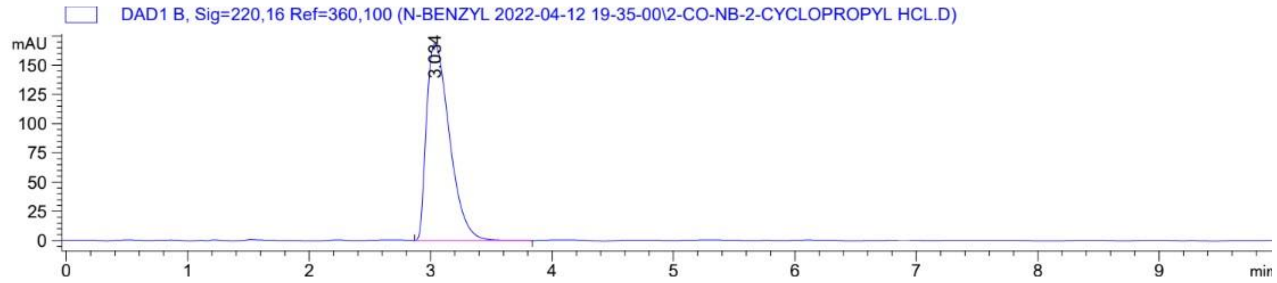

**HPLC Trace of 25O-NB-3-I (34).** Purity (Peak area): 99.3%; Wavelength: 220 nm.

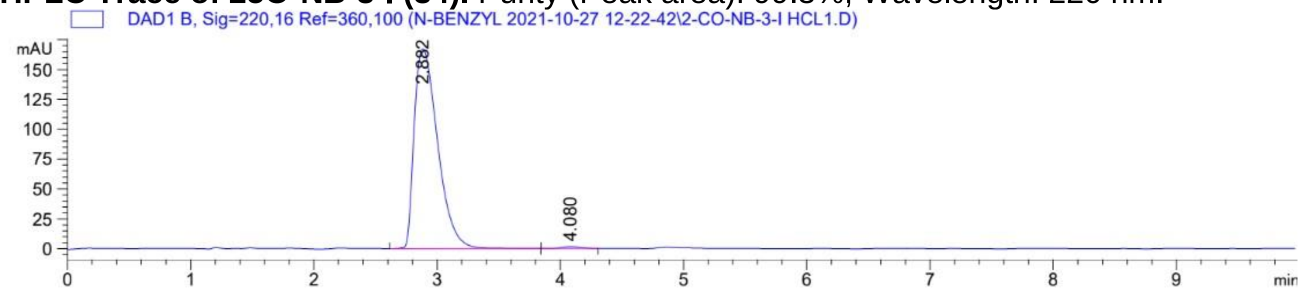

**HPLC Trace of 25O-NBPh-10'-OH (35).** Purity (Peak area): 97.8%; Wavelength: 220 nm.

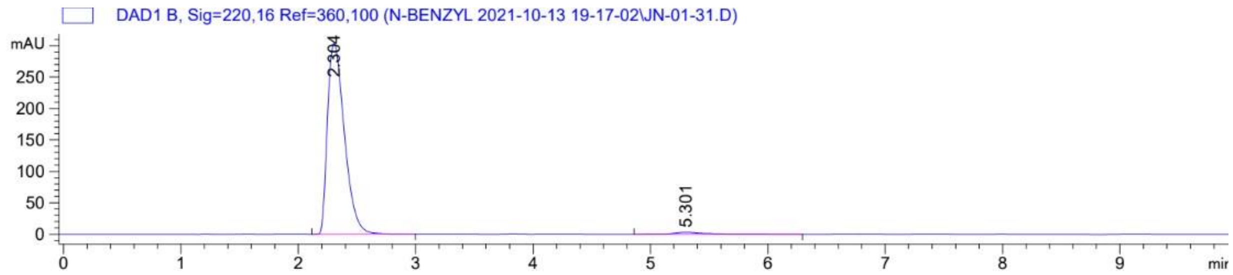
